# Conventional therapy induces tumor immunoediting and modulates the immune contexture in colorectal cancer

**DOI:** 10.1101/2024.08.21.608938

**Authors:** Georgios Fotakis, Dietmar Rieder, Zuzana Loncova, Sandro Carollo, Eckhard Klieser, Daniel Neureiter, Florian Huemer, Sandra Hoegler, Martina Tomberger, Anne Krogsdam, Lukas Kenner, Paul K. Ziegler, Richard Greil, Lukas Weiss, Zlatko Trajanoski

## Abstract

**Background:** Cancer immunotherapies for patients with colorectal cancer (CRC) continue to lag behind other solid cancer types with the exception of 4% of patients with microsatellite-instable tumors. Thus, there is an urgent need to broaden the clinical benefit of checkpoint blockers to CRC by combining conventional therapies to sensitise tumors to immunotherapy. However, the impact of conventional drugs on immunoediting, potentially promoting the positive selection of less immunogenic variants, and on the tumor immune contexture in CRC, remain elusive.

**Methods:** We performed comprehensive multimodal profiling using longitudinal samples from metastatic CRC patients undergoing neoadjuvant therapy with mFOLFOX6 and Bevacizumab. Exome-sequencing, RNA-sequencing and multiplexed immunofluorescence imaging was carried out on tumor samples obtained before and after therapy and the data was analysed using established methods. The results of the analysis were extrapolated to publicly available datasets (TCGA and CPTAC). In order to identify a surrogate marker, an explainable artificial intelligence method was developed using a transformer-based analytical pipeline for the identification of features in Hematoxylin and Eosin (H&E) images associated with specific biological processes, followed by manual evaluation of highly informative tiles by a pathologist.

**Results:** Mutational profiles were highly modified and the level of genetic intertumoral heterogeneity between patients varied following treatment. Evolutionary analysis indicated eradication of some clones and dominant clonal prevalence of others, supporting the notion of pharmacologically-induced cancer immunoeditin. Post treatment samples showed upregulation of HLA class II genes, activation of differentiation and stemness pathways, and changes in the consensus molecular subtypes. The tumor immune contexture was characterised by increased densities of CD8+ and CD4+ T cells, but reduced T cell-tumor cell interactions (and increased T cell exhaustion. The AI-guided analyses of the H&E images pinpointed extracellular mucin deposits associated with stemness genes, suggesting a surrogate marker for routine pathological evaluation.

**Conclusions:** Conventional therapy induces immunoediting and modulates the immune contexture in metastatic CRC patients.

## Background

The immunologic control of neoplasias and its implications on tumor evolutionary dynamics have been a matter of debate within the scientific community over the last century. In 1909 Paul Ehrlich introduced the idea that nascent transformed cells arise continuously in the human body and that the immune system recognises and eradicates these transformed cells before they display clinical manifestation (1). This idea served as the foundation for the development of the immunosurveillance hypothesis, which was postulated in the mid-20th century by Burnet and Thomas (2). However, the concept of immunosurveillance eluded widespread acceptance, until the early 21st century when experiments on knockout mice models validated the central role of the immune effector cells in tumor suppression (3,4). The term cancer immunoediting describes the dynamic process that consists of immunosurveillance and tumor progression, and can be broadly divided into three phases: Elimination (cancer immunosurveillance), Equilibrium (cancer persistence), and Escape (cancer progression) (5). In recent years the term intratumor heterogeneity (ITH) (6) is used to describe the different molecular and phenotypical profiles of distinct tumor cell populations. Taken together, these concepts lead to the idea that tumors are imprinted by the tumor immune contexture (TIC) (7) in which they form, and this process can often result in the emergence of tumor cell populations that are more fit to withstand immunosurveillance. Recent studies often focus on genetic mechanisms of tumor evolution, but there is increasing evidence that tumors can also evolve through mutation-independent processes (8). Multiple studies have shown that tumor initiation and dissemination can be driven by non-genetic mechanisms, (9,10), suggesting that genetic mutations are not always essential for tumor development. These findings highlight the limitations of relying solely on the somatic evolution theory, which follows the Darwinian model of mutation and selection, to fully explain the complex nature of tumor development. Despite recent efforts to characterise the tumor microenvironment (TME) and ITH, the effects of conventional therapy on immunoediting and tumor evolution remain poorly understood. This is particularly true for CRC, which exhibits high heterogeneity among patients (inter-patient), within a single patient’s cancer site (intra-patient), and within different areas of a single cancer site (intra-tumoral). Furthermore, there is a lack of understanding regarding the temporal molecular heterogeneity during conventional therapy, which is known to affect treatment response and prognosis. Additionally, It remains unclear how evolving cancer cells, with their high plasticity, shape their microenvironment under systemic treatment. We recently characterised for the first time the extent of immunoediting that tumors undergo during progression or as a consequence of checkpoint blockade using a mouse model of microsatellite instable (MSI) CRC (11). The quantification of cancer immunoediting showed that targeting the *PD-1/PD-L*1 axis renders the tumors more homogeneous and could eventually lead to a tumor not recognizable by the immune system. Thus, strong immunoediting induced by immunotherapy could represent one mechanism of acquired resistance to checkpoint blockers. However, the impact of conventional therapy on immunoediting and tumor evolution in microsatellite stable (MSS) CRC remains elusive. In the present study we analysed longitudinal samples from patients with MSS metastatic CRC under neoadjuvant therapy, in order to comprehensively characterise the impact of conventional chemotherapy on immunoediting and tumor evolution in a temporal manner. To address this, all patients in the study cohort received a combination of mFOLFOX6 and the anti-VEGF antibody Bevacizumab, a therapeutic regimen designed to enhance cytotoxic effects and potentially improve the immunological control of the cancer. Recent studies suggest that FOLFOX treatment effectively reduces tumor burden, with this effect being dependent on CD8+ T cells. FOLFOX allows tumor-infiltrating lymphocytes (TILs) to maintain a functional differentiation state characterised by lower levels of inhibitory receptors PD-1 and TIM-3. Consequently, TILs from FOLFOX-treated tumors demonstrate higher effector function (12). Additionally, research indicates that blocking VEGF can activate CD4+ and CD8+ T cells, enhance the antigen-presenting function of dendritic cells, increase the cytotoxicity of macrophages, and trigger the activation of the complement cascade (13,14). Utilising multimodal data analysis of DNA and RNA sequencing as well as multispectral immunofluorescence imaging data, we were able to pinpoint putative genetic and non-genetic mechanisms underlying the observed changes in the TIC and the ITH during conventional therapy.

## Methods

### Study cohort

The study cohort consists of five (5) patients with MSS metastatic CRC undergoing neoadjuvant therapy at the 3^rd^ Medical Department of the Paracelsus Medical University Salzburg, Austria. Analyses were approved by the Ethics Committee of the provincial government of Salzburg, Austria (415-E/2343/5-2018). Therapy consisted of chemotherapy with mFOLFOX6 and the anti-VEGF antibody Bevacizumab for 2 to 3 months before undergoing resection of the primary tumor and all metastases in curative intent. For each patient we acquired longitudinal data for two timepoints: the “Biopsy” samples correspond to the diagnostic biopsy of the primary tumor before treatment, while the “Surgery” samples correspond to the resected primary tumor following systemic treatment. For every sample we generated whole exome (WES), RNA sequencing (RNAseq), and multispectral immunofluorescence imaging (Vectra, Akoya Biosciences, Marlborough, MA, USA) data which were then further analysed at the Institute of Bioinformatics of the Medical University of Innsbruck, Austria.

### Publicly available data (TCGA and CPTAC)

All the data that were used to support the findings of this study are publicly available. The results shown in this manuscript, including histological images, are in whole or part based upon data generated by the TCGA Research Network: https://www.cancer.gov/tcga. The CPTAC (15,16) data used in this study were generated by the Clinical Proteomic Tumor Analysis Consortium (NCI/NIH) and were retrieved through the following URL: https://proteomic.datacommons.cancer.gov/. All molecular data for patients in the TCGA and CPTAC cohorts are available at https://cbioportal.org/. We used the TCGA COAD and READ cohort containing respectively 459 and 165 H&E slides for the training and validation part of the deep learning algorithm and the CPTAC cohort of 372 slides for testing purposes. All the collected H&E slides were provided in .svs format.

### DNA and RNA sequencing

DNAse-treated total RNA and RNAse-treated genomic DNA was isolated following the QiaAmp DNA FFPE tissue protocol, and the Qiagen RNeasy FFPE protocol respectively. Total genomic DNA was submitted to exome library preparation (Agilent SureSelect v7) and Illumina sequencing at Genewiz (Azenta/Genewiz, Leipzig, Germany) aiming at 60x coverage. Total RNA was submitted to library preparation using the QuantSeq 3’-mRNA protocol (Lexogen GmbH, Vienna, Austria) and Ion Proton sequencing with Hi-Q chemistry (ion torrent / Thermo Fisher, Vienna, Austria) at Innsbruck Medical University Deep Sequencing Core facility, Austria.

### Immunofluorescence Imaging (Vectra)

A panel of 6 fluorescent markers plus DAPI as a nuclear stain were used to detect the epitopes, namely the following: CD3 (Leica, NCL-L-CD3-565, clone LN10, lot no. 6057975), PD1 (Cell Marque, 315M-96, clone NAT105, lot no. 1630907C), CD8 (Part of Opal Kit from Perkin Elmer, lot no. 2380418), CD45RO (Part of Opal Kit from Perkin Elmer, lot no. 2264307), PD-L1 (DAKO, M3653, clone 22C3, lot no. 10137461), pan Cytokeratin (Abcam, ab7753, clone C11, lot no. GR3186538-14). All slides were stained with the autostainer system Bond RX (Leica Biosystems Inc., Vienna, Austria) and then scanned with the Vectra® 3 (Akoya Biosystems, Marlborough, USA, software version 3.0) microscope. Whole-slide scans were taken at 4x magnification to define regions of interest (whole tissue area) to be scanned in higher resolution (20x) using Phenochart software. InForm (Akoya Biosystems, Marlborough, USA) was used to carry out spectral unmixing and to eliminate autofluorescence from the images.

### Variant calling, Copy Number Variation and Neoantigen predictions

Somatic mutations, copy number alterations, Class I and II HLA types and neoantigens were called by running our previously published neoantigen prediction pipelines nextNEOpi (17) (v1.1) and NeoFuse (18) (v1.2.2a).

Briefly, we used the pipeline’s default options but enabled the automatic read trimming to remove adapter sequence contamination from raw WES and RNAseq reads, the NetChop and NetMHCstab features were disabled, while the “–RNA_tag_seq’’ parameter was set to true since the RNAseq data was generated from 3’ tag sequencing libraries (Lexogen QuantSeq 3’ mRNA-Seq). Finally, predicted neoantigens were filtered and prioritised using the “relaxed” filter set from nextNEOpi.

The Immunoediting scoring scheme is based on a modified reimplementation of a previously published algorithm (19).

### RNA Sequencing (RNAseq) data analysis

The raw RNAseq data was first trimmed to a maximal length of 250 base pairs and residual poly-A stretches were from the 3’ end using fastp (20). For further processing we used the nf-core (21) RNAseq (v3.8.1) pipeline (https://zenodo.org/record/7505987) and set the aligner to STAR (v2.7.4a) to map the reads to the human genome build GRCh38 and provided the gencode v33 GTF file as gene annotation for Salmon (22) (v1.10.1) to obtain the normalised gene expression values as transcript per million (TPM). Differential gene expression analysis was performed using DESeq2 (23) (v1.42.0) and heatmaps were produced using the ComplexHeatmap (24) (v2.10.0) R package. Pathway activity scores were inferred by employing the multivariate linear model available with decoupleR (25) from normalised raw counts and the PROGENy (Pathway RespOnsive GENes - v1.24.0) model (26). Transcription factor activity was inferred using the weighted mean method from decoupleR and the DoRothEA transcription factor/target regulons (A,B,C) (27). Consensus molecular subtypes (CMS) were assessed using the CMScaller (28) R package (v0.99.2) and gene set enrichment analysis (GSEA) was performed with the R package ClusterProfiler (29) (v4.10.0).

To assess the non-genetic mechanisms related to stemness and hypoxia, consensus lists of gene markers (30) were constructed and applied to the normalised gene expression (z-scores) RNAseq data. More specifically, two lists were constructed: (i) a list of hypoxia induced stemness markers (31,32), such as MMP2 and MMP17, ZEB1-2, and TWIST2, (ii) a list of gene markers related to stem-like phenotypes, such as VIM, NANOG, SOX2, and DCLK1. Consequently, the gene lists were condensed into gene signatures expression values using the GSVA (v1.50.5) R package. Once the gene signature expression values were calculated the TCGA COAD-READ cohort (n = 641) was stratified into “high” and “low” expression, using as threshold the optimal cutpoint as computed by the surv_cutpoint() function of the survminer (v0.4.9) R package. The survival analysis was carried out using the survival (v3.6-4) R package.

In order to elucidate the functional state of the infiltrated T-cells, a consensus list of T-cell exhaustion markers was constructed from the literature (33–35). The RNA sequencing expression matrix, as returned from Salmon, was used as an input file and the raw values were converted to z-scores using Stouffer’s z-score method. Finally, heatmaps were produced using the ComplexHeatmap (v2.10.0) R package.

### Tumor evolutionary trajectory inference

In order to deconvolute the mixture of clonal populations within each sample we fitted a hierarchical Bayes statistical model, as implemented by PyClone (36) (v0.13.1), on the WES data using the tumor purity estimations as returned from ASCAT (37) (v3.1.2). Once the clonal composition was deconvolved from the initial cell mixture, the clones were related to each other in terms of evolution. To that end, we used quadratic integer programming (QIP), as implemented by CITUP (38) (v0.1.0). Due to the iterative heuristic method of CITUP being highly dependent on initialization, we addressed this issue by performing 10,000 restarts with random initializations.

To asses the results of the evolutionary trajectory inference we focused on three points, (i) the dynamics of clonal prevalences during therapy, (ii) the level of ITH before (biopsy) and after therapy (surgery), and (iii) the events of clonal dominance and extinction. Since the clonal prevalence metric can take values from 0 (complete absence) to 1 (complete dominance), we define two thresholds: (1) any clonal population that exhibits clonal prevalence above 0.50 (> 0.50) is considered dominant (dominance), and (2) the clonal populations that exhibit clonal prevalence below 0.01 (< 0.01) are considered eradicated (extinction).

### Immunofluorescence Imaging (Vectra) data analysis

The raw Vectra images were obtained as a multispectral .tiff file composed of 8 channels (for CD3+, CD8+, CD4Ro+, CK+, DAPI+, PD-1+, PD-L1+ staining and autofluorescence). Each channel was extracted and tiled into smaller equally-sized images (1344 x 1008 pixels) which enabled further analysis. In order to distinguish between the tumor and stromal area and to compare its cellular composition, the original (full-scale) images were annotated by a pathologist and binary image masks for tumor and stromal regions were created. Subsequently, the binary masks were also tiled into corresponding equally-sized images and together with the image tiles for each fluorescent channel were analysed using Cellprofiler (39) software (v4.2.0). The position of each cell, its final phenotype and cell densities were calculated, as well as location of the staining within the cell (in the cytoplasm or nucleus) and the location of cells within tissue (background/tumor/stroma and inside/outside of the tumor margin).

Voronoi diagrams were generated to evaluate the spatial organisation, and all cell-cell interactions (direct neighbors) were counted. To test whether the number of direct interactions of each pairwise cell type combination is significantly different than expected by chance, a Monte Carlo simulation with 1000 iterations was performed. In this simulation, the location of the cells from the imaged slide was randomly permuted, while keeping the number of cells from each type constant and the overall cellular positions fixed. Then a z-score and p-value was calculated to assess avoidance or attraction (z-score < 2, z-score > 2, p < 0.01). A combined cohort z-score was calculated using the Stouffer’s z-score method for overall meta analysis and plotted as heatmaps. CD3+ T-cell / PDL1+ tumor cell interaction frequencies were obtained dividing the number of direct CD3+ T-cell / PDL1+ tumor cell neighbors by the total number of direct neighbors of CD3+ T-cells. Immune cell - tumor cell mixing scores were obtained by dividing the number of direct immune cell / tumor cell neighbors by the total number of direct immune cell neighbors.

### Explainable artificial intelligence (AI) model

To further support and validate our findings, we employed H&E stained whole-slide images (WSIs) from the TCGA and CPTAC cohorts. Using a deep learning-based method, we aimed to predict the stemness signature directly from the WSIs for computational pathology. The STAMP workflow that we applied allows the prediction of a patient’s biomarkers directly from WSIs using deep learning (40).

Briefly, after loading the WSIs at a 10x magnification (1.14 MPP), the slides are tessellated into *n* tiles of 224×224 pixels. Tiles that do not contain enough tissue or are blurry, are removed using Canny edge detection (41). The remaining non-rejected tiles are stain-normalised using Macenko’s method (42). Subsequently, the feature extractor model runs inference to obtain a feature vector for each tile, which are then concatenated into a large feature matrix. By default, the pipeline uses CTransPath (43), a robust model pre-trained on 14 million histological patches.

For this study, we opted for a binary classification of the stemness signature, dividing it into “High” and “Low” categories based on the mean value, as this method provided the most informative results.

After preprocessing the TCGA and CPTAC cohorts, we generated feature matrices with dimensions of *n* x 768, where *n* is the number of tiles varying for each slide. We then created two tables linking the patient IDs to the biomarkers and to the filename of the feature matrix and the classification task was carried out by employing a weakly-supervised Transformer model that was trained on the TCGA COAD-READ cohort. Subsequently, the trained model was deployed on the CPTAC cohort. The final model’s performance was evaluated by generating metrics and statistics. The predictions were sampled 1,000 times to recalculate the metrics, yielding a 95% Confidence Interval (CI) for the validation cohort model performance.

To explain the model’s decisions, a prediction heatmap was generated using gradient class activation maps (gradCAM), highlighting the top influential tiles in the model’s decision-making for predicting High or Low stemness. We selected the 10 most accurately predicted WSIs for High and Low stemness signature and generated the 5 tiles that most positively contributed to these predictions. A higher prediction score value indicates that a stronger signal was captured by the model, helping the human interpretability of the heatmaps. The pipeline was implemented using Python (v3.10), torch (v2.0.1) and cuda (v11.7) and was run on an HPC cluster using a Nvidia RTX 8000 GPU for the deep learning tasks.

The five most influential tiles taken from the 10 top-ranked cases for “high” and “low” stemness (totaling 100 individual images) were then analysed by a histopathologist (PKZ). Upon manual review of top-tier images, 17 histopathological features have been commonly occuring in CRC (44) have been identified. Six features describe nuclear and cellular morphology, six tumor cell architecture and five microenvironmental properties (Supplementary Table S4). These features have been scored semiquantitatively using a 4-tier scoring system (0=absent/none, 1=weak/few, 2=moderate/frequent, 3=strong/abundant) per tile image.

## Results

### The tumor mutational profiles are substantially altered after therapy

It is generally accepted that the level of ITH can predispose patients to diverse clinical outcomes. The selective pressure introduced by conventional therapies and the immune system can lead to treatment resistance as a result of the expansion of pre-existing sub-clonal populations or from the evolution of newly formed clones that exhibit some evolutionary trait that renders them less immunogenic or immunoevasive. The observed genetic instability of tumors provides the raw material for the generation of ITH and thus the characterization and quantification of genetic variants in tumor samples plays an important role in understanding the genetic mechanisms underlying tumor evolution (Figure 1B). With regards to known CRC driver genes, the most common mutation in the cohort was *TP53*, with four out of five patients in the study cohort exhibiting missense mutations on *TP53* in the biopsy samples (Figure 1C). Another interesting finding is that only one of the patients (Patient 5) retains the same *TP53* mutation post treatment, which indicates its important role in tumor initiation but raises a question regarding its role in cancer progression. The fact that 4 out of 5 patients’ tumors did not retain the *TP53* mutation post-treatment indicates that the clones harboring the specific mutation are eradicated during treatment. Other CRC driver genes found mutated in the study cohort include *APC*, *KRAS*, *FAT*, *ATM*, *NRAS*, and *LRP1B*. Although the number of acquired mutations and the number of Neoantigens appear to be increasing post-treatment (Figure 1D), there is no statistically significant difference between the pre and post-treatment samples. The observed increasing trend can be explained by two factors. First, two patients (Patient 5 and Patient 8) exhibit a hypermutated phenotype post-treatment, aligning with the observed intratumoral heterogeneity evident in the tumor evolutionary trajectory results. Second, the pre-treatment biopsy samples contain significantly less tumor material compared to the post-treatment surgical specimens. These findings highlight the dynamic changes in tumor mutational profiles post-therapy, emphasising the role of intratumoral heterogeneity in treatment response and the potential implications for targeted therapeutic strategies in colorectal cancer.

**Fig. 1.**
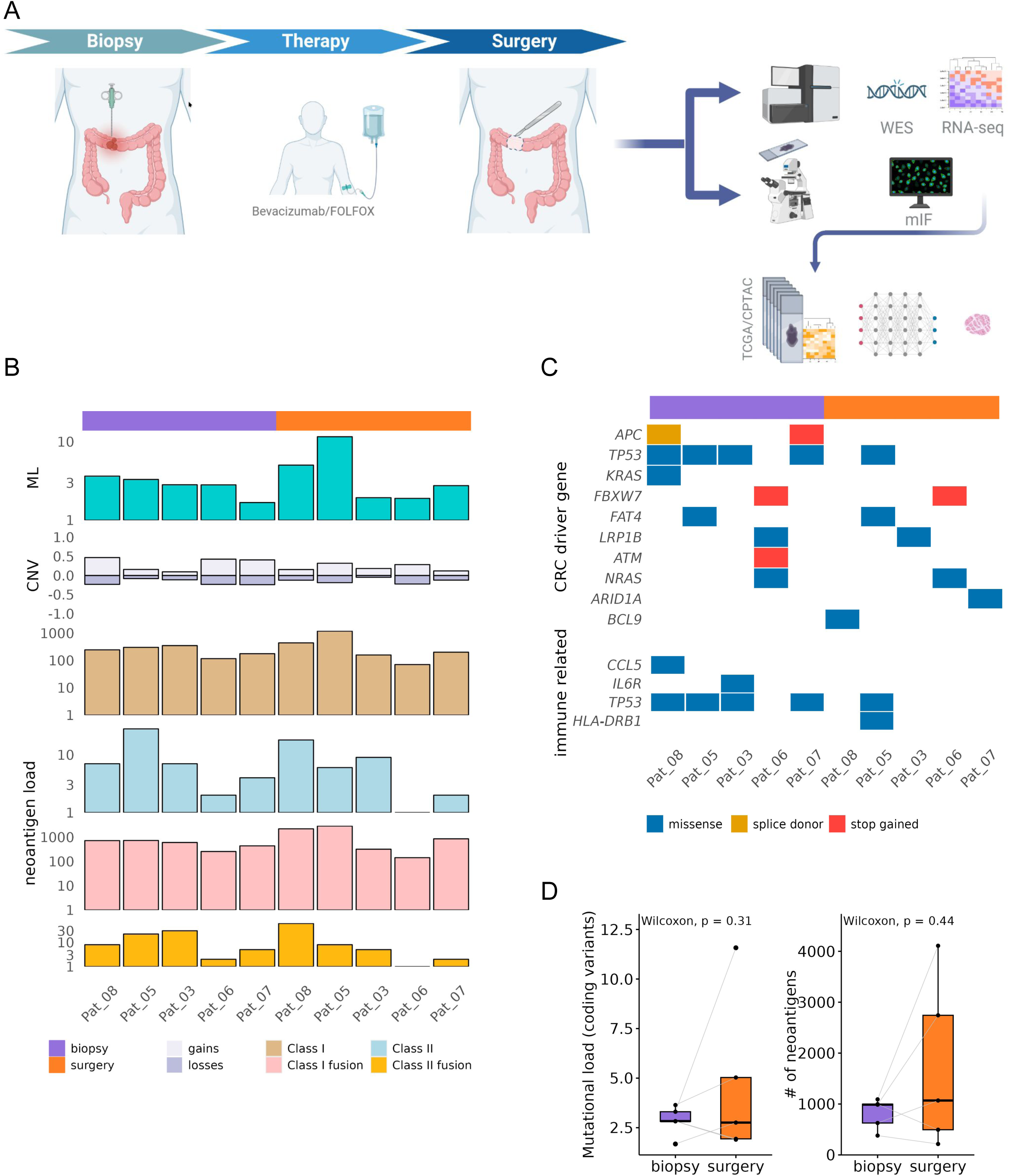
Study design and genomic data analysis. **A** CRC samples were collected before (biopsy) and after (surgery) the therapy with mFOLFOX6/Bevacizumab. For each one of the samples we generated DNA/RNA sequencing data coupled with immunofluorescence images. Stemness signature identified from gene expression was used together with TCG/CPTAC histopathology images to train an AI model. **B** Results of the WES data analysis. From top to bottom: ML, Mutational Load; CNV, Copy Number Variations; Neoantigen load for class I and II canonical and fusion Neoantigens. **C** Most common driver gene and immune related gene mutations. **D** Box and whiskers plots for the Mutational Load of coding variants (left) and the total number of Neoantigens (right) before and after therapy.

### Neo-adjuvant therapy drives non-neutral clonal evolution

To better understand the effects of the combined chemo- and anti-VEGF therapy on immunoediting and tumor evolution, we studied the dynamics of the cellular prevalences of the clonal populations. According to our hypothesis, if the combined chemo- and anti-VEGF therapy does not induce an anti-tumor immune response, the clonal populations will undergo neutral evolution since the clones will be eradicated irrespective of their immunogenic potential. During neutral evolution, the clonal prevalence of all clonal populations will increase in a stable manner leading eventually to a high level of ITH since all tumor clones are represented to some extent in the post-treatment (surgery) samples. On the other hand, if conventional treatment induces an anti-tumor immune response, the immune system will target and eradicate the more immunogenic clonal populations allowing the less immunogenic clones to increase their relative proportion within the tumor. This process is referred to as positive selection and leads to decreased levels of ITH since the immune mediated selective pressure provides an evolutionary advantage to the less immunogenic or the immunoevasive clones that will eventually become dominant.

Although there’s a high level of inter-patient heterogeneity, there is a common pattern that appears in all samples (Figure 2A). On the onset and during the time period of systemic therapy, specific clones exhibit an increase in clonal prevalence and become dominant over the rest. Furthermore, in most patients, certain clones are eradicated and completely replaced by dominant clones, a process that resembles clonal sweeping. A clonal sweep occurs when a branch driver event enhances the fitness of a subclone to such an extent that it outcompetes all other subclones in the tumor. Given the occurrence of clones becoming dominant while other clones are eradicated leads to the conclusion that the level of ITH appears to be lower in most samples following therapy when compared to the samples before therapy for all patients. Despite the fact that the evolutionary trajectory analysis indicates an immune mediated selective pressure during treatment, there is an evident decreasing trend in the immunoediting score for all patients post-treatment (Figure 2B), the statistical analysis showed that the difference between the pre and post-treatment samples was not statistically significant (p-value = 0.09). The non-significant result of the pairwise comparison test may be attributable to the low statistical power of the analysis due to the size of the study cohort. However, the observation that immunoediting scores are lower in post-treatment samples compared to pre-treatment samples raises questions about the functional state of the immune system following neoadjuvant therapy. Taking all the above into consideration, we conclude that conventional treatment induces an anti-tumor immune response which imposes a selective pressure to the clonal populations and shapes the final clonal composition.

**Fig. 2.**
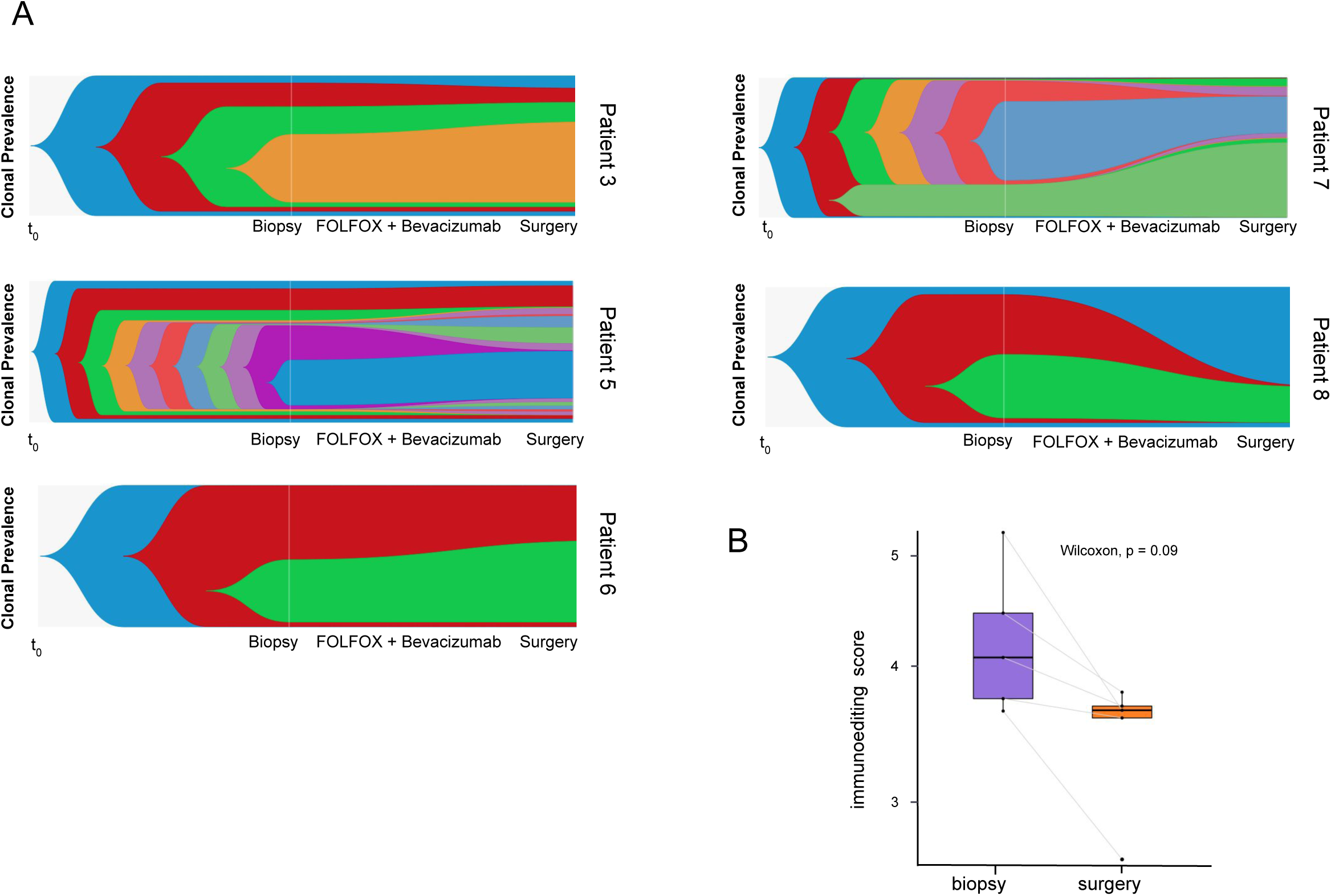
Tumor evolutionary trajectory inference and immunoediting scores. **A** Fishtail plots illustrating the evolutionary trajectories of each tumor. The y-axis represents the clonal prevalences, while the x-axis represents time. **B** Box and whiskers plot representing the immunoediting scores as calculated for each patient before and after therapy.

In summary, neo-adjuvant therapy induces non-neutral clonal evolution characterized by dynamic changes in clonal prevalence and a potential impact on immune-mediated selective pressures, highlighting implications for therapeutic strategies in colorectal cancer.(9

### Transcriptional profiling shows upregulation of HLA II and of anti-inflammatory genes

The differential gene expression analysis returned 1718 statistically significant differentially expressed genes in total (Figure 3A). Among the most interesting findings was the upregulation of genes belonging to the HLA II family, namely *HLA-DOA*, *HLA-DOB*, and *HLA-DRA* in the post-treatment samples. Similarly, the genes *CCL5*, *CCL2*, and *CXCL13* were found to be upregulated in the post treatment samples (Figure 3B). The overexpression of the inflammatory chemokine *CCL5* has been positively correlated with a reduction in T-regulatory (Treg) infiltration and CD8+ apoptosis in human and murine CRC models (45). Both *CCL5* and *CCL2* increase the attraction of tumor-associated macrophages (TAMs) within the TME while simultaneously inhibiting potential anti-tumor T cell activities. Moreover, *CCL2* has been suggested to also promote angiogenesis (46). The inflammatory factor *CXCL13* has been proposed as a biomarker and a putative therapeutic target in CRC, as it is involved in tumor growth, infiltration and migration.

**Fig. 3.**
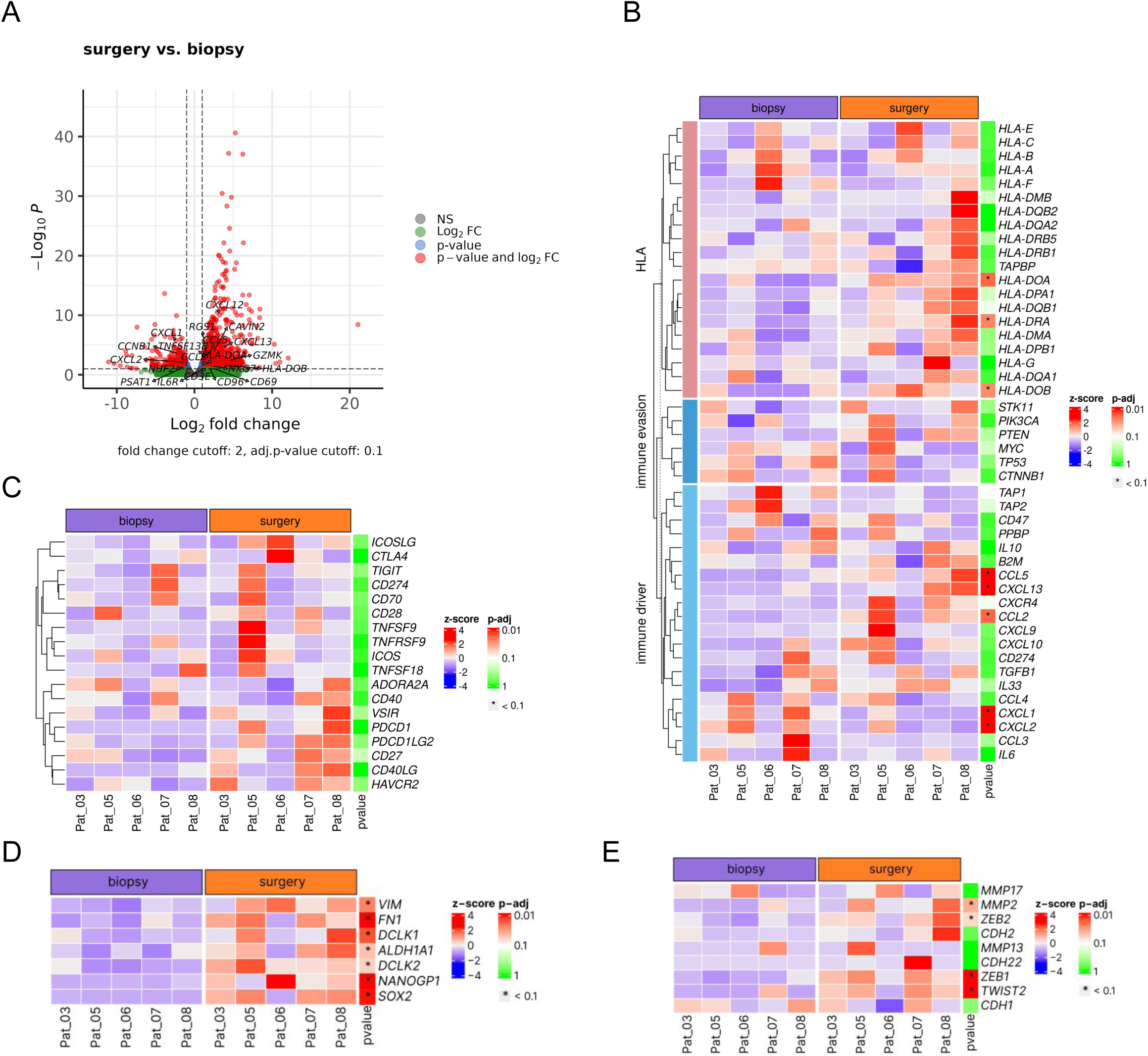
Transcriptomic data analyses. **A** Volcano plot showing the differentially expressed genes as returned from DESeq2. **B** Heatmap showing the normalised expression values (z-scores) of HLA genes (upper group), Immune evasion related genes (mid group) and immune driver genes (lower group). **C** Heatmap showing the normalised expression values (z-scores) of genes related to T-cell exhaustion, before and after therapy. **D** Heatmap showing the normalized expression values (z-scores) of genes related to cancer stemness (“stemness signature”), before and after therapy. **E** Heatmap showing the normalized expression values (z-scores) of genes related to hypoxia induced stemness, before and after therapy.

These transcriptional changes highlight a significant upregulation of HLA II and anti-inflammatory genes post-treatment, suggesting potential immune modulation and therapeutic targets in CRC.

### Neo-adjuvant therapy ultimately leads to T-cell exhaustion

In order to elucidate the functional status of the tumor infiltrated immune components we applied a consensus T-cell exhaustion gene marker list on the expression data. The comparison of gene expression levels between pre-treatment and post-treatment samples shows that genes associated with T-cell exhaustion were upregulated in the tumors of most patients. These genes include *ICOSLG*, *CTL4*, *TIGIT*, *CD274*, *CD70*, *CD28*, *CD40*, *CD27*, *CD40LG*, *TNSF9*, *TNFSF18*, *ADORA2A*, *VSIR*, *PDCD1*, *PDCD1LG2*, and *HAVCR2* (Figure 3C). The increased expression of exhaustion markers indicates that the T-cells within the TME are in a dysfunctional state and might thereby explain the fact that we observed decreased immunoediting post-treatment.

Overall, neo-adjuvant therapy induces upregulation of T-cell exhaustion markers in colorectal tumors, suggesting a potential mechanism for decreased immune function post-treatment.

### Cancer stemness emerges following treatment

It has been suggested that the major differences between cancer stem cells (CSCs) and non-CSCs is largely attributed to the activation of the biological programme termed Epithelial-to-Mesenchymal Transition (EMT) (47,48). The EMT activation in CSCs causes them to lose their epithelial characteristics and acquire mesenchymal traits, which include an increased ability to migrate and invade (49), as well as increased resistance to various types of conventional therapies (50,51). In order to assess the difference in stem-like phenotypes between the pre and post-treatment tumor samples, we compiled a list of stemness markers from the literature and applied it to the transcriptomic data. The expression of EMT associated genes like *VIM* and *FN1* was found to be significantly upregulated in the post treatment samples. Transcription factors that have been linked to stem-like phenotypes and promoting stemness in neoplastic cells (52,53), like *SOX2*, *NANOG*, *ALDH1A1* and *NANOGP1* are also upregulated in the post-treatment samples (Figure 3D). It is important to note that *NANOG* is upregulated only in one of the patients post-treatment but *NANOGP1* is strongly upregulated in all patients. *NANOGP1* is a tandem duplication of *NANOG* and it is suggested to be a strong inducer of cell pluripotency (54,55). Additionally, the expression of both isoforms of *DCLK* (*DCLK1* and *DCLK2*) was found to be upregulated in the post-treatment samples, which are considered as putative CSC markers specifically for CRC (56,57).

This indicates that cancer stemness markers, including EMT-associated genes and pluripotency factors like NANOGP1 and DCLK isoforms, are significantly upregulated following treatment, suggesting a potential role in treatment resistance and tumor progression.

### Hypoxia plays a crucial role in the activation of the EMT programme

All patients of the study cohort received combined neoadjuvant therapy of mFOLFOX6 and Bevacizumab, which is a humanised monoclonal antibody (immunoglobulin G1) that has been recombinantly engineered to target and bind with vascular endothelial growth factor (*VEGF*). Since Bevacizumab is targeting *VEGF* we expect that VEGF-mediated angiogenesis in the tumor region will be inhibited causing poorly vascularized regions of the carcinoma and leading to a hypoxic environment. Hypoxia can activate the EMT programme through the transcriptional regulator *HIF-1* that responds to cellular hypoxia. The impact of hypoxia on EMT induction via *HIF-1* has been demonstrated in clear-cell renal-cell carcinoma. In this context, *HIF*–*1* induces the expression of *ZEB1*, *ZEB2*, *TWIST1*, *TWIST2*, and *TCF3* thus indirectly repressing the expression of E–cadherin and conferring mesenchymal attributes on the tumor cells (58). Although we did not observe *HIF-1* to be upregulated, the genes *ZEB1*, *ZEB2*, *TWIST2*, and *CDH2* were upregulated in the post-treatment samples (Figure 3E) suggesting that the hypoxic conditions possibly activated these genes of the *HIF-1* pathway.

These results suggest that despite *HIF-1* not being upregulated, the observed upregulation of genes such as *ZEB1*, *ZEB2*, *TWIST2*, and *CDH2* in post-treatment samples indicates a potential role of hypoxia in activating the EMT programme via alternative pathways in colorectal carcinoma.

### Therapy activates pathways connected to cancer stemness

The GSEA results further support the hypothesis of EMT activation during therapy (Figure 4A). DNA replication and cell cycle pathways were found to be suppressed which is typical in CSCs. Additionally, signalling pathways such as the Hedgehog, *cAMP*, *cGMP-PKG* and the *PPAR* were found to be activated in the post-treatment samples with a high statistical significance (p-values < 0.05). The majority of CRCs exhibit aberrant regulation in the Hedgehog signalling pathway but its role in maintenance of cancer cell stemness extends to a wide range of tumors (59,60), while *cAMP*, *cGMP-PKG* and *PPAR*s have also been reported to regulate stem cell-like properties of CSCs via the activation of the EMT programme (61–63). Interestingly, post-treatment samples showed activation of the extracellular matrix (ECM)-receptor interaction signalling pathway. This pathway is closely related to the cell and focal adhesion molecules pathways, which were also upregulated following treatment. A growing body of evidence suggests that ECM proteins provide structural and biochemical support, thereby regulating the proliferation and self-renewal of CSCs in CRC (64,65). Finally, our results show that the CSCs exert an immunosuppressive phenotype via the activation of the *PD-1/PDL-1* expression signalling pathway (p-value = 0.04).

**Fig. 4.**
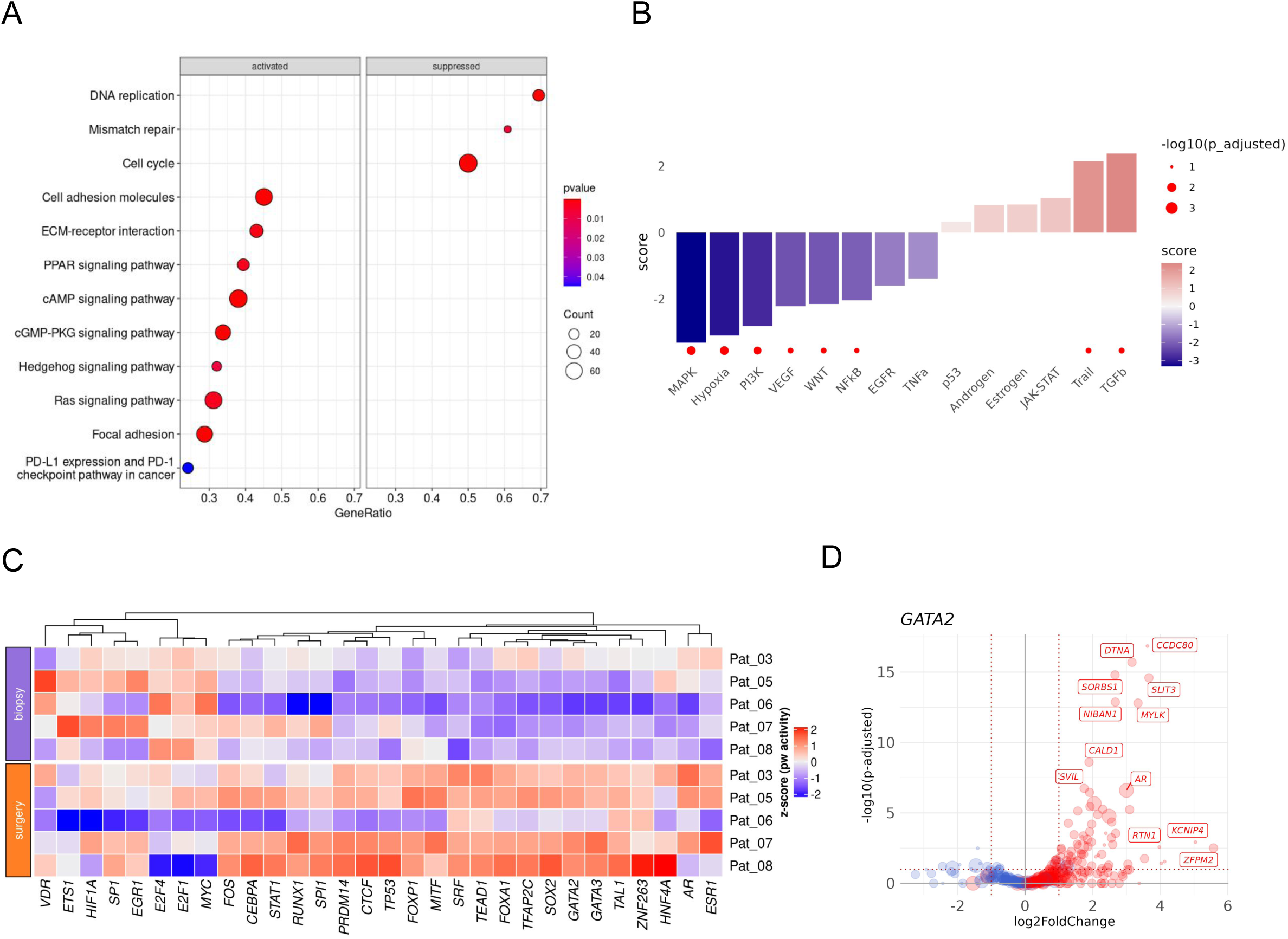
Gene set enrichment, pathway and transcription factor activity analyses. **A** Dotplot showing the activated (left) and suppressed (right) signaling pathways in the tumor samples with their respective p-values, as returned from the GSEA analysis. **B** Barplot showing the activity scores of downregulated (blue) and upregulated (red) pathways along with their respective -log_10_(adjusted p-values), as returned from the PROGENy analysis. **C** Heatmap representing the normalised activity scores of the top 30 (by standard deviation) transcription factors (TFs), before and after treatment. **D** Volcano plot showing the upregulated (red) and downregulated (blue) targets (regulons) of *GATA2*.

As we anticipated, VEGF and its downstream pathway targets, such as the MAPK and PI3K pathways (66), were found to be downregulated since all the patients were treated with Bevacizumab (Figure 4B) . Although the *MAPK* and *PI3K-Akt* pathways are known to be essential in CSCs, recent studies indicate that primary *BRAF*^mut^ and *KRAS*^mut^ colorectal cancer spheroids acquire stemness and activate compensatory pathways when treated with *MAPK* pathway inhibitors (67). One potential compensatory mechanism could be the observed upregulation of the *TGF-β* pathway, which has been associated with promoting EMT, facilitating angiogenesis, suppressing the anti-tumor activity of immune cells in the TME, and enhancing the stemness of CRC cells (68). Another interesting finding is the upregulation of the *TRAIL* pathway, which has been extensively studied as a novel strategy for tumor elimination. Cancer cells that overexpress *TRAIL* death receptors (like *DR4* and *DR5*) are induced to undergo apoptosis and inhibit blood vessel formation (69). However, recent research has revealed that CSCs can readily develop mechanisms to evade *TRAIL*-induced apoptosis by altering their properties, thereby enhancing their survival and promoting angiogenesis (70).

These findings underscore the activation of multiple pathways linked to cancer stemness following therapy, including *TGF-β*, Hedgehog, *cAMP*, *PPAR*, ECM-receptor interaction, and the immunosuppressive *PD-1/PDL-1* signaling, highlighting their potential roles in promoting therapy resistance and maintaining stem-like properties in colorectal cancer.

### Treatment-induced upregulation of transcription factors is associated with EMT activation

Transcription factor (TF) activity analysis revealed that several TFs associated with cancer stemness were upregulated. Transcription factors like *TFAP2C*, *SOX2*, *TAED1*, *GATA2*, *PRDM14*, *MITF*, *HNF4A*, *CTCF*, and *EGR1* were found to exhibit increased activity with noticeably elevated gene expression values of their targets (Figures 4C). *TFAP2C* has been found to be robustly upregulated in CRC tissues and cells, and its upregulation correlates with the expression of stem cell factors and promotes stemness and chemoresistance of CRC cells in vivo and in vitro (71). *SOX2* drives CSCs properties, and contributes to tumor aggressiveness in CRC (72). The transcriptional cofactors *YAP* and the transcriptional co-activator *TAZ* of the Hippo pathway, regulate the *TEAD* proteins, which have been suggested to be important regulators of cancer cell stemness and dedifferentiation in CRC (73–75). Both *GATA1* and *GATA2* (Figure 4D) have been reported to play key roles in CRC cell proliferation, invasion, EMT, and cancer stemness (76,77). A recent study, involving single-cell RNAseq data from 81 samples, demonstrated that the upregulation of the *HNF4A* expression is connected with CRC stemness (78), while the upregulation of *CTCF* has been linked to EMT activation (79). Finally, *EGR1* has been extensively researched due to its capacity to translate various stimuli into appropriate alterations in the cell’s transcriptional programme. This transcription factor can be considered as a diverse “stress sensor” for EMT-inducing agents and is crucial for hypoxia-induced tumor progression, survival, and angiogenesis (80).

These results emphasise the significant upregulation of transcription factors associated with cancer stemness following treatment. These factors are implicated in promoting EMT, enhancing cancer cell stemness, and potentially contributing to therapeutic resistance in colorectal cancer.

### The tumor shapes its microenvironment during therapy

The activation of the EMT programme during treatment can also be corroborated by the results of the consensus molecular subtype inference. With regards to the pre-treatment samples, the tumor of Patient 7 is categorised as CMS1, the tumors of Patients 3 and 8 are categorised as CMS2, while the tumors of Patients 5 and 6 are categorised as CMS3.

Although none of the patients’ tumors were categorised as CMS4 pre-treatment, four out of five patients’ tumors are categorised as CMS4 post-treatment (Supplementary Figure 3F), while for one patient the CMS inference returned inconclusive results (FDR > 0.01).

The activation of the EMT program during treatment aligns with CMS subtype shifts post-treatment, with four out of five patients’ tumors transitioning to CMS4, which is characterised by activated *TGF-β* pathway, activation of the EMT programme, and it is associated with a worse prognosis.

### The tumor modifies the immune contexture during therapy

In order to evaluate the TIC in terms of spatial organisation, we produced Voronoi tessellations of the immunofluorescence images (Figures 5A-B). The analysis of the TIC demonstrated increased cell densities of CD8+ and CD4+ T-cells within the tumor region for 4 patients following treatment (Figure 5C), which is indicative of increased T-cell infiltration within the core of the TME. Out of five patients, only Patient 8 exhibits decreased T-cell densities within the tumor region following therapy, suggesting T-cell exclusion from the TME. The analysis of the cell neighbourhoods and cell-cell interactions revealed a decrease in the mixing scores in four out of five patients, indicating reduced interactions between the tumor and the immune cells (Figures 5G-H). Taken together, we observed an increased infiltration of T-cells within the TME during treatment. These findings align with our hypothesis of therapy-induced immune-mediated selective pressure, as demonstrated by the non-neutral patterns of tumor evolution, while the observed decrease in the mixing scores can be explained by the upregulation of T-cell exhaustion gene markers.

**Fig. 5.**
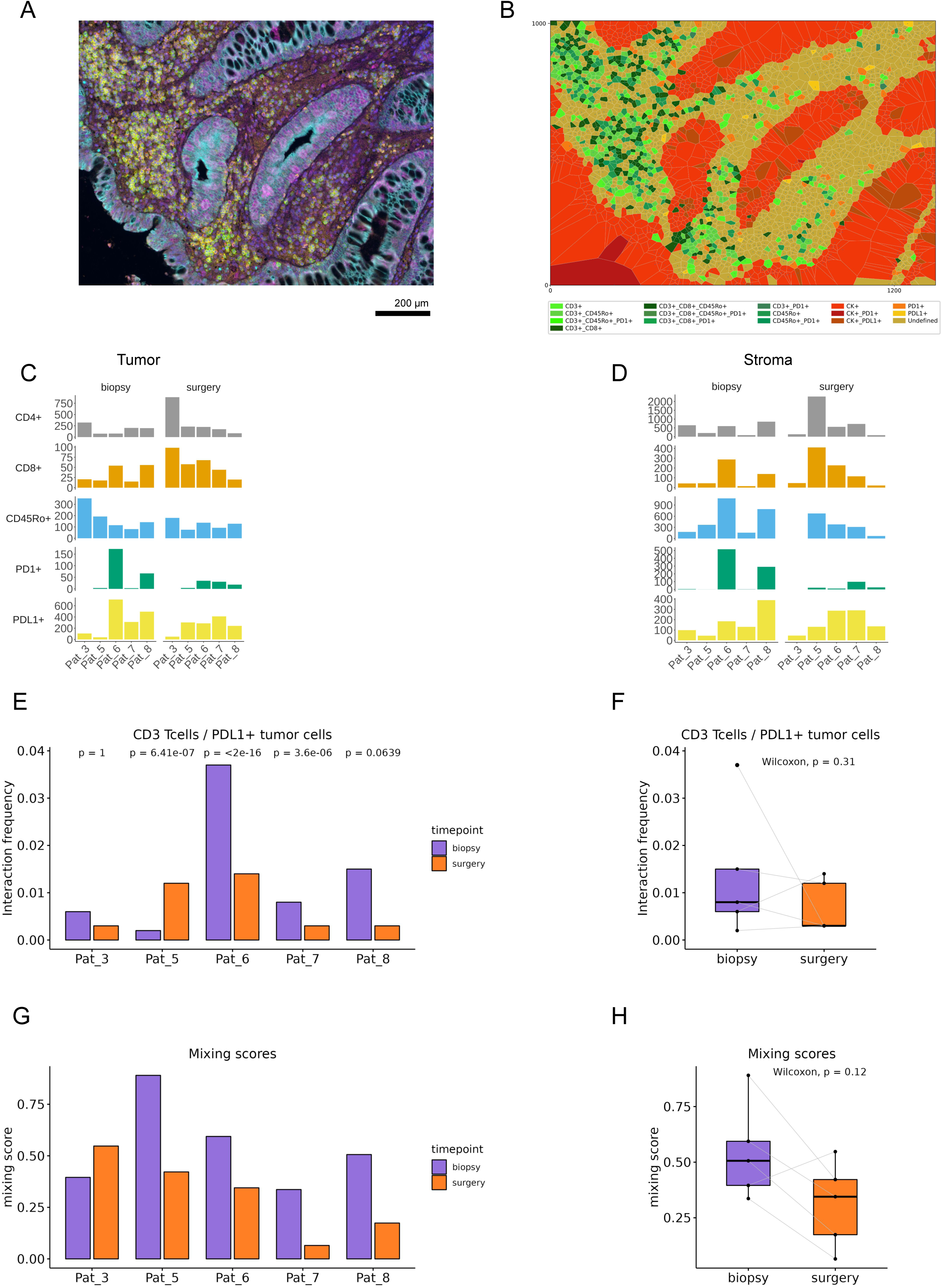
Immunofluorescence image analysis. **A** Typical multispectral immunofluorescence tile (left) and its respective Voronoi tessellation with cell types colored according to the color code tabulated below (right) (tile dimensions: 1344 x 1344 µm). **B-C** Barplots showing cell densities (cells/mm^2^) within the tumor and stroma regions, before and after therapy. **D-E** Barplot and boxplot showing the interaction frequencies (%) between CD3+ T-cells and PDL1+ tumor cells for each patient, before and after therapy **G-H** Barplot and boxplot showing the mixing scores (immune/tumor interactions vs immune any cell interaction) for each patient, before and after therapy.

### Expression of the cancer stemness signature is associated with worse overall survival

Given the small size of our study cohort, we sought to extrapolate our findings to publicly available datasets. As a first step, we employed the stemness signature in order to perform a survival analysis using data retrieved from the TCGA COAD-READ cohort (n = 641). The Kaplan-Meier curves in Figure 6A show that high expression of the stemness signature is significantly associated with worse overall survival (p = 0.0027). The survival analysis results highlight the clinical significance of the stemness gene signature, indicating its potential role in treatment resistance and tumor progression.

**Fig. 6.**
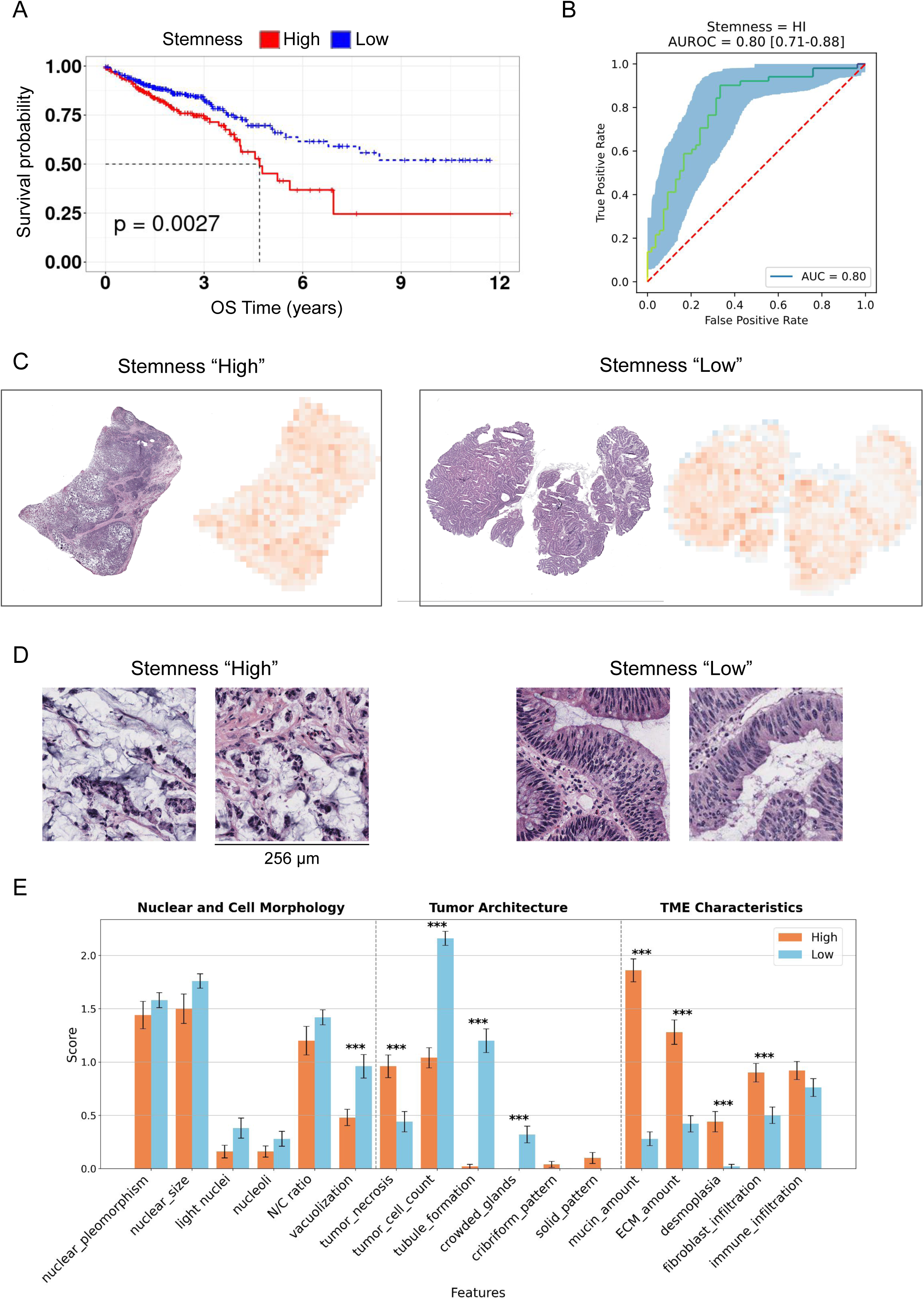
Survival and AI-guided histopathology analyses. **A** Kaplan-Meier curve for the TCGA COAD-READ cohort shows a significant association between high expression of the stemness signature and worse overall survival. **B** AUROC curves evaluating the performance of the final model on the external validation cohort CPTAC-CRC. The bootstrapped 95% confidence interval is shown in light blue, while the dotted red line represents the performance of a random prediction model. **C** Heatmaps for CPTAC slides classified as Stemness “High” (left panel) and Stemness “Low” (right panel). Each panel includes the H&E slide alongside the Gradient class activation maps (gradCAM), that highlight the most influential tiles in the model’s decision-making for predicting Stemness high or low. **D** Top tiles for the Stemness signature: the two tiles on the left correspond to “Stemness high” predictions, while the latter two on the right correspond to “Stemness low” predictions. **E** Bar plot of 17 predefined histopathological patterns commonly occurring in CRC, assessed using a 4-tier scoring system: 0=absent/none, 1=weak/few, 2=moderate/frequent, 3=strong/abundant. Scores are averaged from the top 5 tiles of the 10 most accurately predicted WSIs.

### Explainable AI reveals clinically relevant cancer stemness phenotypes

After preprocessing the WSIs and excluding low quality scans, 599 slides of the TCGA and 372 of the CPTAC cohorts remained for the analysis. For training and deployment, we used data from 583 TCGA patients and validated the model on the external cohort of 105 patients from CPTAC. The deployed model demonstrates accurate performance on the external CPTAC cohort, achieving an area under the receiver-operator characteristic (AUROC) with a bootstrapped 95% CI of 0.80 [0.71-0.88] for predicting the stemness signature (Figure 6B). To further evaluate the model’s ability to learn clinically relevant concepts, heatmaps of the external CPTAC cohort were generated. The top tiles represent the key influential tiles used to predict “High” and “Low” stemness (Figure 6C). Expert evaluation by a trained pathologist revealed that high-confidence slides differ significantly predominantly in their tumor architectural and microenvironmental characteristics. While only one out of six (17%) items of the evaluated nuclear and cellular components have been found to be different (vacuolization), four out of six (67%) and four out of five (80%) characteristics in the architectural and TME categories have been found to differ significantly (Fig. 6E). In particular, “high” stemness tiles tend to have more extracellular mucin, while “low” stemness tiles showed more intracellular mucin vacuoles (Figure 6D-E). Mucins are a family of glycoproteins (MUC1–MUC24) expressed by many epithelial tissues that have been implicated in the promotion of tumor progression of various malignancies, including colon cancer (81,82), suggesting the model’s correct identification of the phenotype corresponding to a high stemness in CRC. In addition, fibroblast infiltration, ECM production and stromal desmoplasia are associated with high stemness phenotype, confirming the complex changes in tumor growth induced by the different transcriptional programs. These results underscore the AI model’s potential to provide clinically meaningful insights into cancer stemness phenotypes, enhancing our understanding and improving therapeutic strategies for colorectal cancer patients.

## Discussion

Despite advances in sequencing technologies, immuno-oncology, and tumor evolution research, the impact of conventional therapies on immunoediting and tumor evolution in MSS CRC remains unclear. In this study we successfully characterised variations among cancer cell subgroups arising from genetic and non-genetic mechanisms within a single tumor.

Our DNA sequencing analysis shows that driver mutations initiate tumors, but the clonal populations with these mutations are likely eradicated during neoadjuvant systemic therapy, as they are mostly absent in post-treatment samples. RNA sequencing reveals that all patients’ tumors exhibit a stem-like phenotype, likely due to the activation of the EMT program. This is supported by the observed transition from various consensus molecular subtypes pre-treatment to CMS4 post-treatment. Conventional therapy induces immunoediting, indicated by the non-neutral evolution pattern during treatment and increased T-cell infiltration in the TME. However, post-treatment, cell-cell interactions between immune and tumor cells decrease, likely due to T-cell exhaustion and the immunosuppressive phenotype of CSCs indicated by the PD-1/PDL-1 axis activation. One of the major limitations of the present study is the size of the cohort. To mitigate this limitation, we successfully extrapolated our results to publicly available datasets. We integrated a stemness signature derived from gene expression data of our study cohort with TCG/CPTAC molecular data and histopathology images to develop an AI model, which provided clinically meaningful insights into cancer stemness phenotypes. Histopathologic analysis of the most influential images identified by AI reveals divergent morphologic characteristics between high and low stemness cases, particularly in tumor architecture and TIME-associated features, while nuclear morphology remains largely unchanged. High stemness is associated with extracellular mucin deposition, ECM production, and stromal desmoplasia, likely due to upregulation of the TGFb-signalling pathway and downregulation of hypoxia and VEGF-signalling. Some histologic features were frequently detected, but features such as cribriform or solid tumor growth were infrequent, likely due to limitations imposed by tillation (tile size 256×256 µm).

Finally, we comprehensively summarised our findings through a model of immunoediting and tumor evolution under neoadjuvant therapy (Figure 7). Initially, the chemotherapeutic agents target tumor cells regardless of immunogenicity, leading to the release of tumor-specific antigens and initiating immunoediting. During the elimination phase, the immune system targets the more immunogenic tumor clones, causing less immunogenic clones to increase their prevalence. This results in positive selection and reduced ITH as fitter, less immunogenic clones dominate. The tumor microenvironment’s inflammatory and hypoxic conditions activate the EMT programme, conferring a stem-like phenotype to specific tumor cells, therefore enhancing their self-renewal and proliferative capabilities. This leads to the dominance of CSCs, a transition to consensus molecular subtype CMS4, ECM formation and increased extracellular mucin deposits. Overexposure of immune components to the antigenic TME causes T-cell exhaustion, and the immunosuppressive phenotype of CSCs further impairs immunoediting, rendering the tumor immune “cold”.

**Fig. 7.**
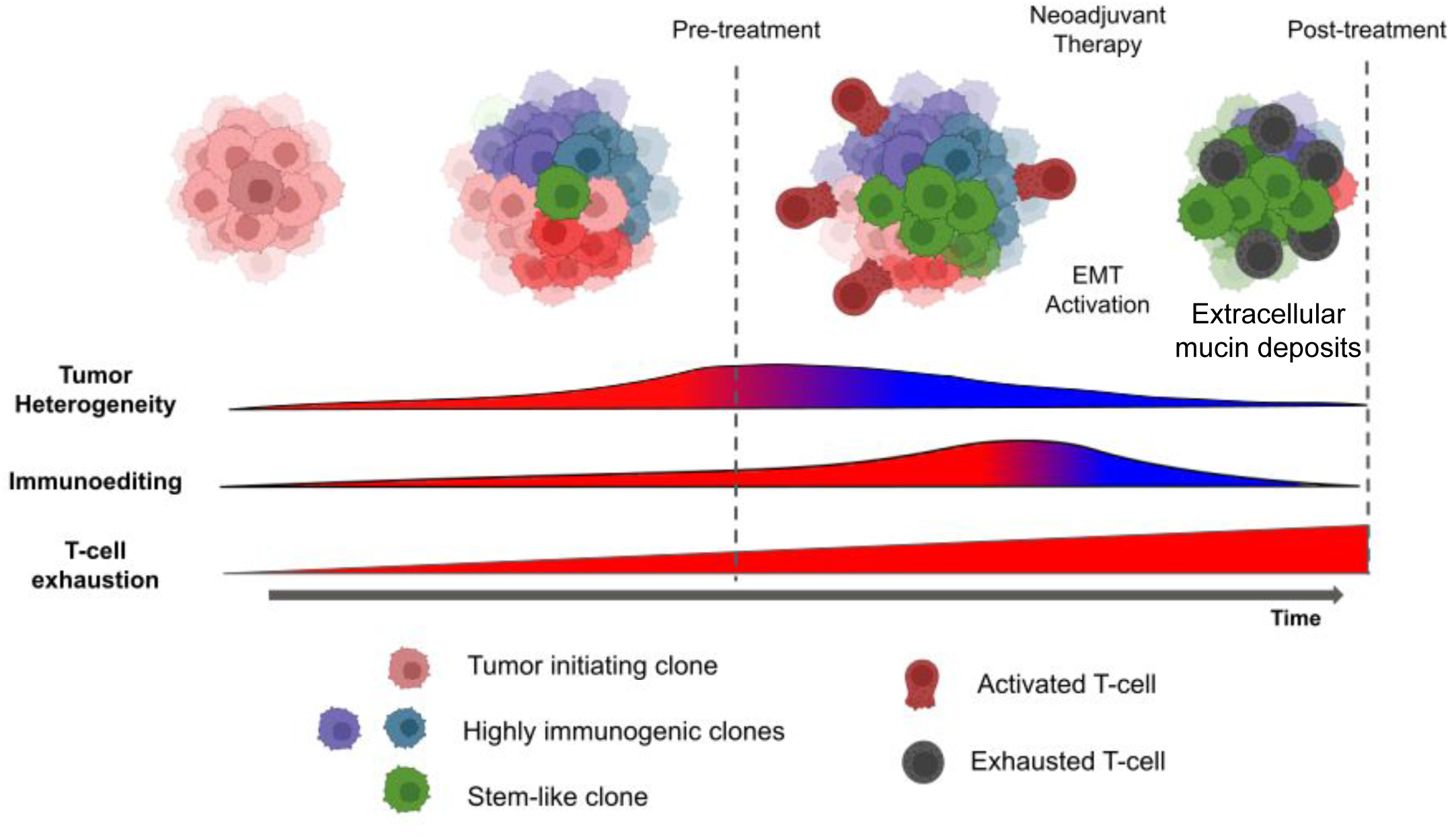
Schematic representing the suggested model of tumor evolution under therapy-induced immune-mediated selective pressure. At the onset of neoadjuvant therapy, chemotherapeutic agents eradicate tumor cells regardless of their immunogenicity, leading to a massive release of neoantigens. This attracts immune system components to the TME. The immune system then targets highly immunogenic tumor cells, reducing the level of the ITH and favoring less immunogenic tumor cells. Over time, T-cells become exhausted, while the inflammatory and hypoxic conditions in the TME activate the EMT programme. The EMT activation results in a stem-like phenotype, enhancing the self-renewal and proliferative capabilities of the CSCs, eventually leading to their dominance over the rest of the clonal populations.

## Conclusions

By performing multimodal data analysis and integration, we were able to pinpoint the impact of conventional therapy on immunoediting and on the genetic and non-genetic mechanisms attributing to the observed ITH following conventional therapy in MSS CRC. We expect that the results of this project will provide a deeper understanding of the tumor-immune interactions and will allow the development of more robust combinatorial therapeutic strategies for MSS CRC.

## Supporting information

Supplemental Tables Description

## Declarations

### Ethics approval and consent to participate

Sample collection and data analyses were approved by the Ethics Committee of the provincial government of Salzburg, Austria (415-E/2343/5-2018).

### Consent for publication

Not applicable.

### Availability of data and materials

The processed data are available through Zenodo (DOI: 10.5281/zenodo.11655540). The raw (DNA/RNA) sequencing and immunofluorescence imaging files are restricted to protect patient privacy.

## Authors’ contributions

Conception, design, funding: ZT, LW; Manuscript writing: GF, ZT, DR; Figures: DR, GF, SC. DNA and RNA sequencing: AK; Imaging data generation: SH, MT, LK; Bioinformatics analyses: DR, GF; Imaging data analysis: DR, ZL; Explainable AI: SC; Histopathology analysis: PKZ, EK; Manuscript reviewing & editing: LW, PKZ, RG, LK, DN, FH; All the authors read and approved the final manuscript.

## Acknowledgements

We thank Dr. Mihaela Angelova, PhD, for her support and for providing the code used to compute the immunoediting scores.

Additional file 1: Supplementary Figures S1-S6 (.pdf format). Additional file 2: Supplementary Tables S1-S4 (.xlsx format). Additional file 3: Descriptions of supplementary tables (.pdf format).

## List of abbreviations

AI: Artificial Intelligence
AUROC: Area Under the Receiver-Operator Characteristic
CI: Confidence Interval
CNV: Copy Number Variation
COAD: Colon Adenocarcinoma
CPTAC: Clinical Proteomic Tumor Analysis Consortium
CRC: Colorectal Cancer
CSC: Cancer Stem Cell
ECM: Extracellular Matrix
EMT: Epithelial-to-Mesenchymal Transition
FFPE: Formalin-Fixed Paraffin-Embedded
GSEA: Gene Set Enrichment Analysis
H&E: Hematoxylin and Eosin
ITH: Intra-Tumoral Heterogeneity
ML: Mutational Load
MSI: Microsatellite Instable
MSS: Microsatellite Stable
READ: Rectal Adenocarcinoma
TAM: Tumor Associated Macrophage
TCGA: The Cancer Genome Atlas Program
TF: Transcription Factor
TIC: Tumor Immune Microenvironment
TIL: Tumor-Infiltrating Lymphocyte
TME: Tumor Microenvironment

