## Supplemental Figures S1-S6 for "Conventional therapy induces tumor immunoediting and modulates the immune contexture in colorectal cancer"

#### Supplementary Figures and Legends

**Figure S1. Variants' concordance**

Stacked barplots showing the concordance between biopsy and surgery samples of **A** all variants and **B** coding variants for each patient. The numbers represent the shared variants between biopsy-surgery samples (red), variants found exclusive in biopsy samples (blue), and variants found exclusive in surgery samples (green).

**Figure S2. Tumor evolutionary trajectory and clonal prevalences**

**A** Fishtail plot displaying the tumor evolution for patient 7. The y-axis represents the clonal prevalences, while the x-axis represents time. **B** Fishtail plot showing the increase in clonal prevalence of a specific subclonal population (green), from 0.23 before therapy to 0.53 (dominance) after therapy. **C** Fishtail plot showing the decrease in clonal prevalence of a specific subclonal population (red), from 0.18 before therapy to < 0.01 (extinction) after therapy.

**Figure S3. Transcriptomic data analysis and consensus molecular subtype (CMS) predictions**

**A** Dotplot showing the results of the principal component analysis (PCA). The samples cluster together in groups according to the timepoints (biopsy-surgery), indicating that no batch effects are present. The PCA shows also clear distinction between surgery and biopsy for each patient. **B** Heatmap showing the transcriptomic profiles (top 100 variable genes) for all patients, before and after therapy. Heatmaps showing the differential expression ($log_2$FC) of immune modulator genes (**C**) and T-cell related genes (**D**). **E** Heatmap visualization of mRNA gene set analysis showing selected CMS-informative signatures for comparisons of patients (n =6) classified with CMScaller. Red and blue indicate relative up- and down-regulation, respectively, and color saturation reflects statistical significance. **F** Table showing the CMS predictions for each patient with their respective false discovery rate (FDR), before and after therapy.

**Figure S4. Pathway and transcription factor activities**

**A** Heatmap representing the normalised z-scores of pathway activities before and after treatment, as returned from the DoRothEA analysis. **B** Responsive genes in the PI3K pathway along their DESeq2 stat-values. The pathway is downregulated as most of the genes with negative weight are upregulated. **C** Barplot showing the activity scores of the Top 30 downregulated (blue) and upregulated (red) transcription factors in the post treatment samples. **D** Volcano plot showing the upregulated (red) and downregulated (blue) targets (regulons) of *NFKB1* following therapy.

**Figure S5. Cell densities and cell-to-cell interactions**

**A-B** Barplots showing cell densities (% of all cells) within the tumor and stroma regions. **C-D** Interaction analysis showing cell-cell attraction (red) and avoidance (blue). Tiles are showing the number of patient in which significant (p < 0.01) interactions (red) or avoidances (blue) between each cell-type pair occur.

.

**Figure S6. AI-guided most informative histopathological slides**

**A** Top 5 tiles for the stemness “High” signature predictions for a specific patient’s slide. The score of these tiles ranges from 0.80 to 0.82. **B** Top 5 tiles for the stemness “Low” predictions for a specific patient’s slide. The score of these tiles ranges from 0.79 to 0.76.

### Fig. S1

A

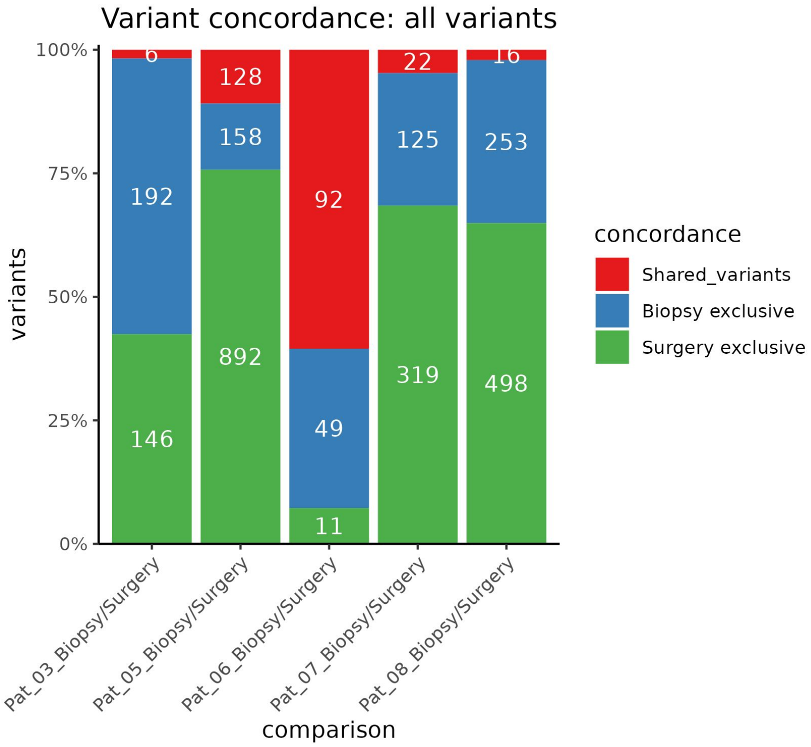
Variant concordance: all variants
100%
75%
50%
25%
0%
variants
6
192
146
128
158
892
92
49
11
22
125
319
16
253
498
Pat_03_Biopsy/Surgery
Pat_05_Biopsy/Surgery
Pat_06_Biopsy/Surgery
Pat_07_Biopsy/Surgery
Pat_08_Biopsy/Surgery
comparison
concordance
Shared_variants
Biopsy exclusive
Surgery exclusive

B

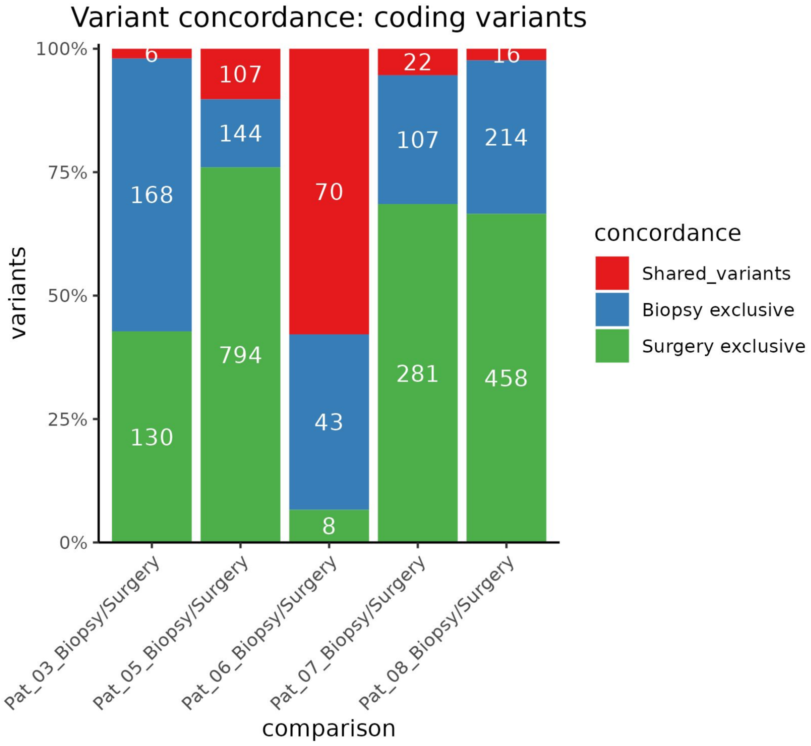
Variant concordance: coding variants
100%
75%
50%
25%
0%
variants
6
168
130
107
144
794
70
43
8
22
107
281
16
214
458
Pat_03_Biopsy/Surgery
Pat_05_Biopsy/Surgery
Pat_06_Biopsy/Surgery
Pat_07_Biopsy/Surgery
Pat_08_Biopsy/Surgery
comparison
concordance
Shared_variants
Biopsy exclusive
Surgery exclusive

Fig. S2

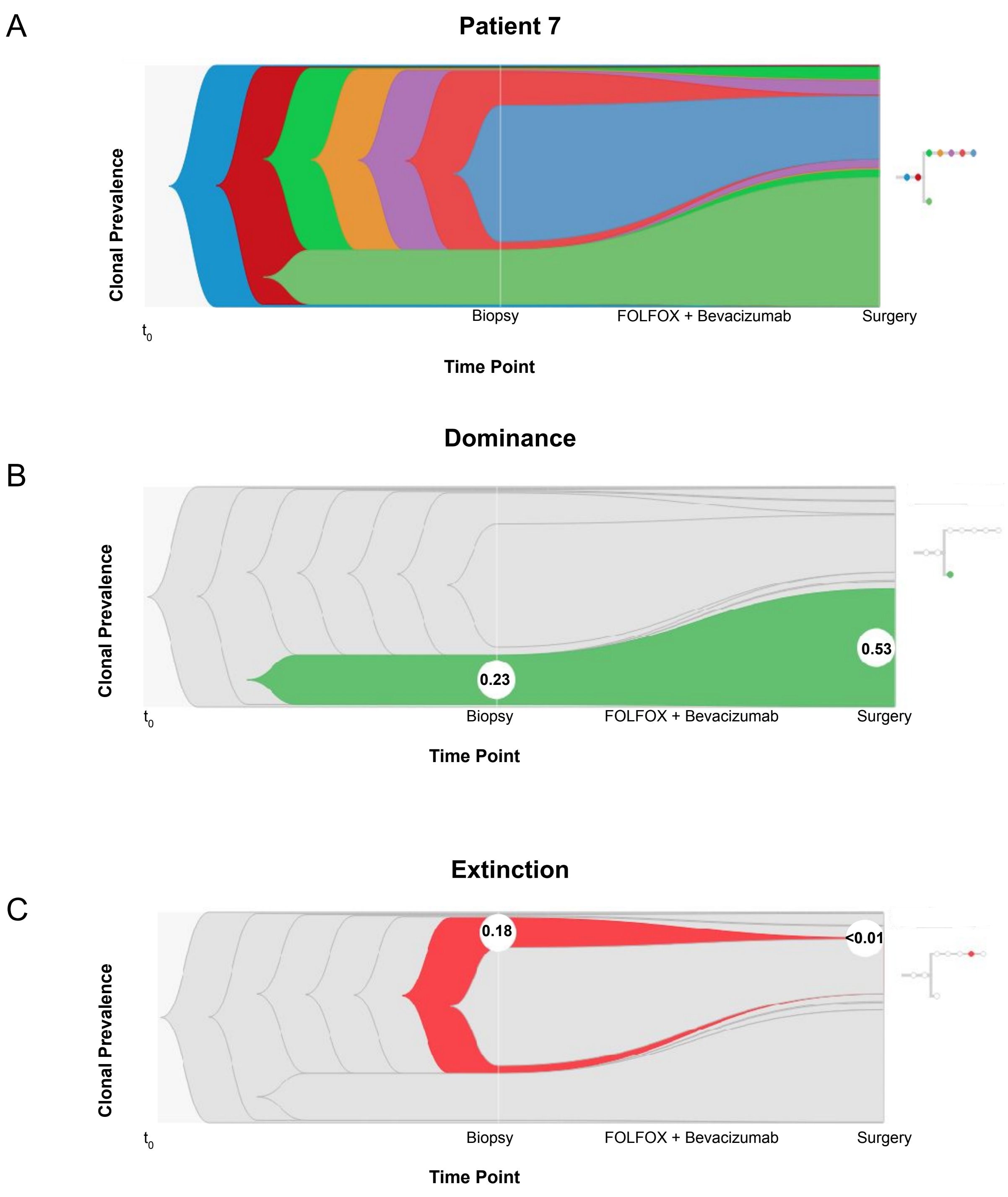
A
Patient 7
Clonal Prevalence
t0
Biopsy
FOLFOX + Bevacizumab
Surgery
Time Point
Dominance
B
Clonal Prevalence
0.23
0.53
t0
Biopsy
FOLFOX + Bevacizumab
Surgery
Time Point
Extinction
C
Clonal Prevalence
0.18
<0.01
t0
Biopsy
FOLFOX + Bevacizumab
Surgery
Time Point

Fig. S3

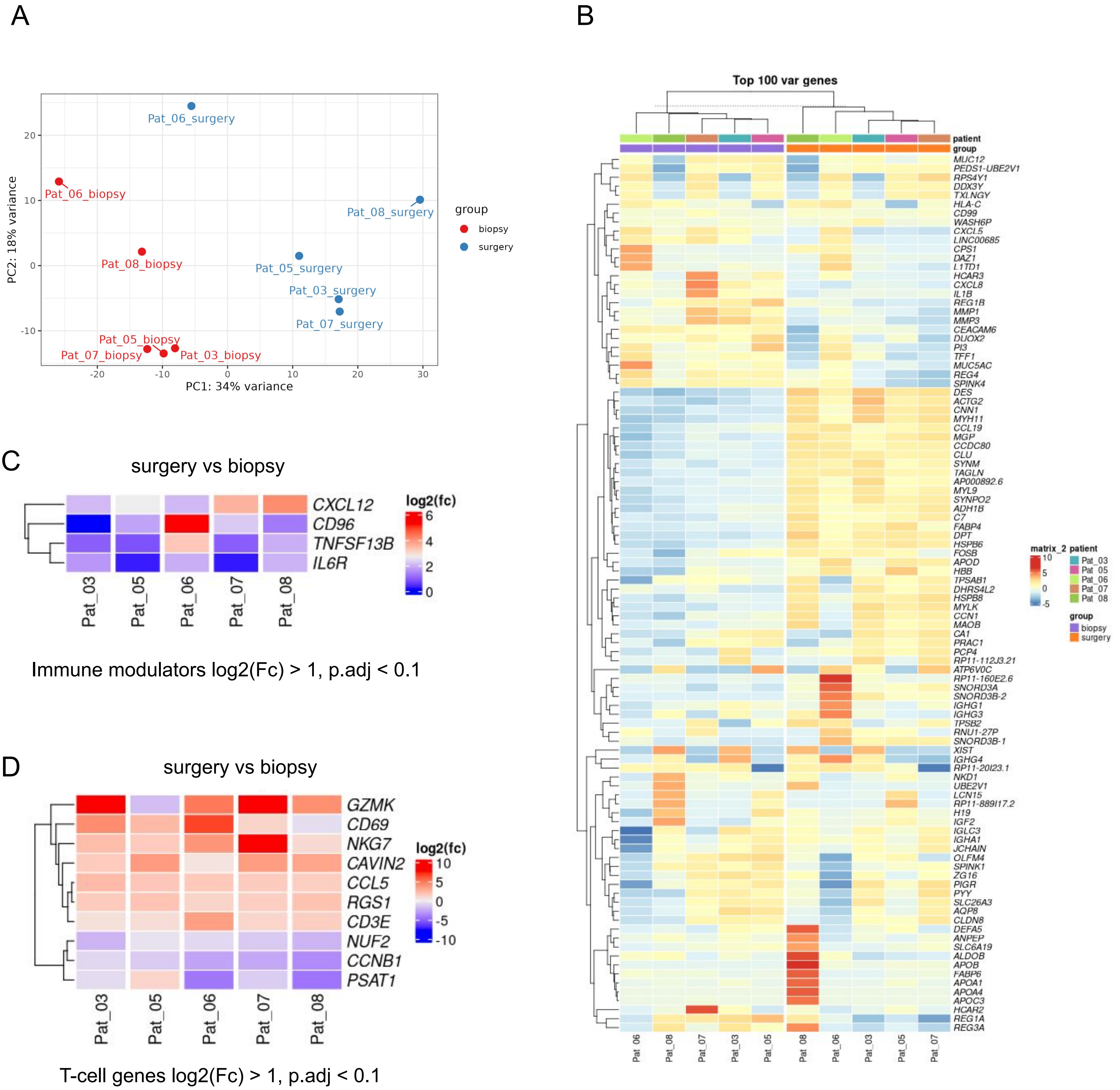


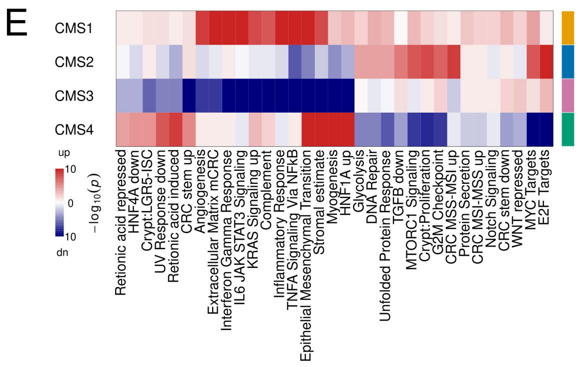


F

| Patient | prediction biopsy | prediction surgery | FDR biopsy | FDR surgery |
|---|---|---|---|---|
| Pat_03 | CMS3 | CMS4 | 0.001 | 0.001 |
| Pat_05 | CMS2 | CMS4 | 0.001 | 0.001 |
| Pat_06 | CMS3 | NA | 0.002 | 0.344 |
| Pat_07 | CMS1 | CMS4 | 0.002 | 0.001 |
| Pat_08 | CMS2 | CMS4 | 0.001 | 0.001 |

#### Fig. S4

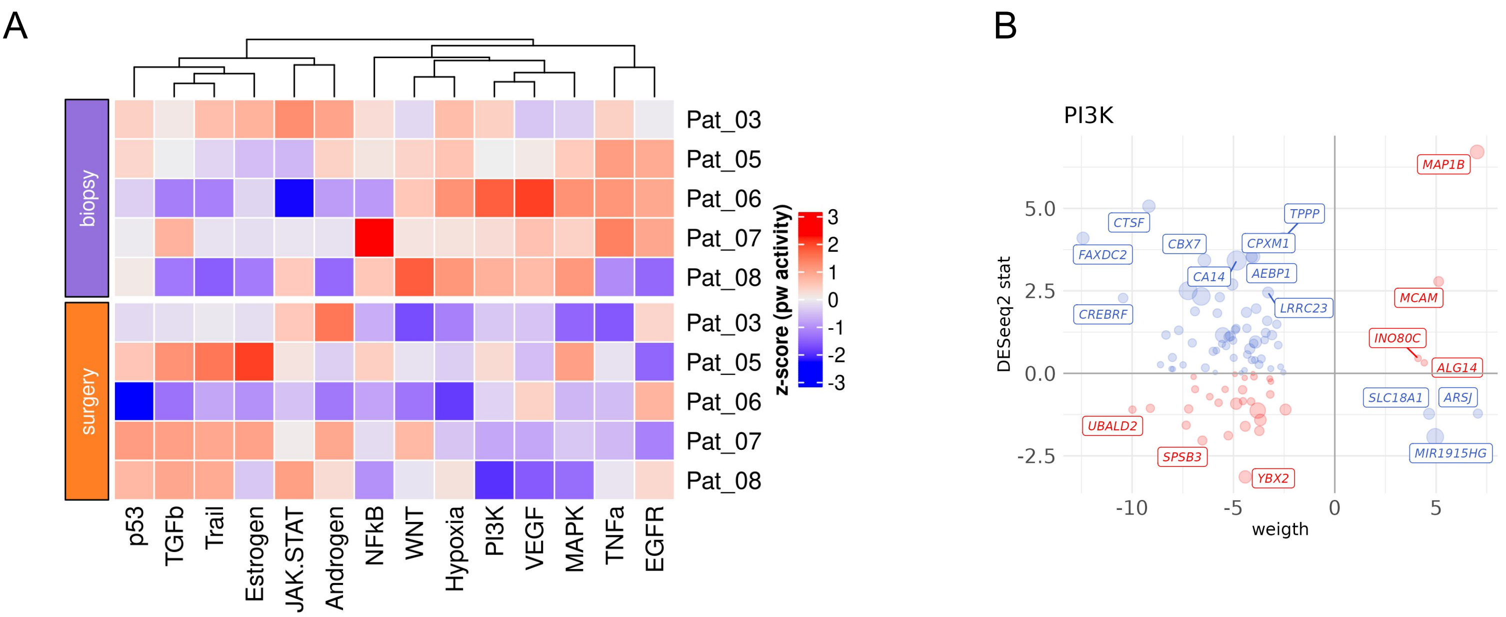


Pathway activities: surgery vs biopsy

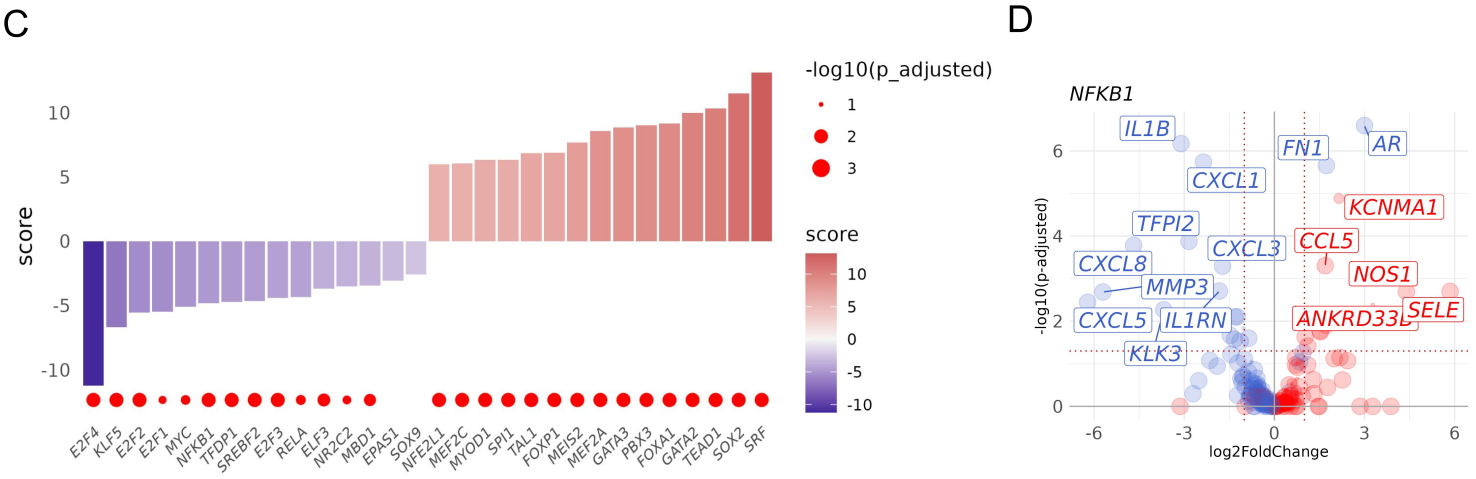


Transcription factor activities: surgery vs biopsy

### Fig. S5

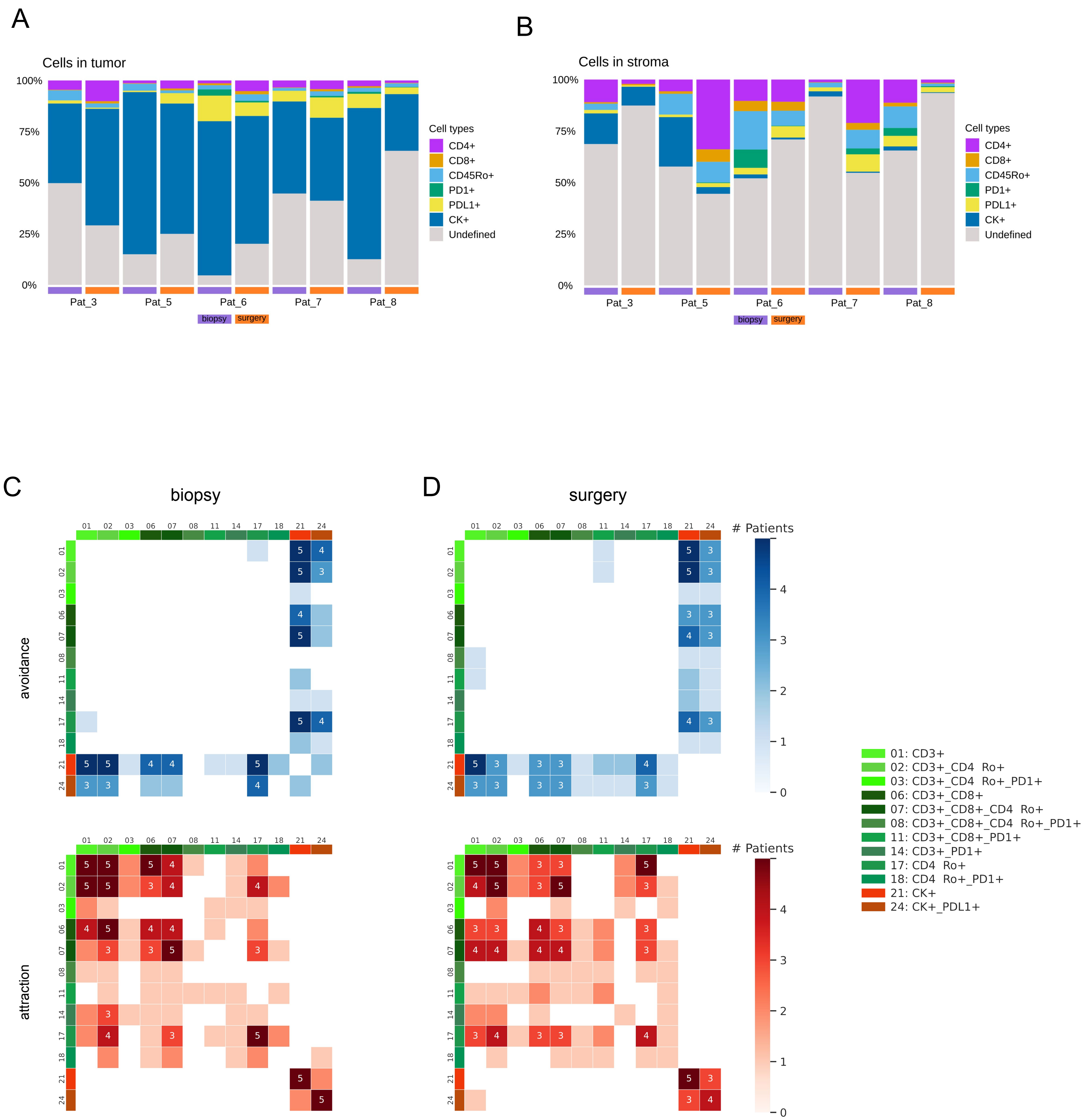
A
Cells in tumor
100%
75%
50%
25%
0%
Pat_3
Pat_5
Pat_6
Pat_7
Pat_8
biopsy
surgery
Cell types
CD4+
CD8+
CD45Ro+
PD1+
PDL1+
CK+
Undefined
B
Cells in stroma
C
biopsy
avoidance
attraction
D
surgery
### Patients
01: CD3+
02: CD3+_CD4 Ro+
03: CD3+_CD4 Ro+_PD1+
06: CD3+_CD8+
07: CD3+_CD8+_CD4 Ro+
08: CD3+_CD8+_CD4 Ro+_PD1+
11: CD3+_CD8+_PD1+
14: CD3+_PD1+
17: CD4 Ro+
18: CD4 Ro+_PD1+
21: CK+
24: CK+_PDL1+

#### Fig. S6

A

Stemness “High”

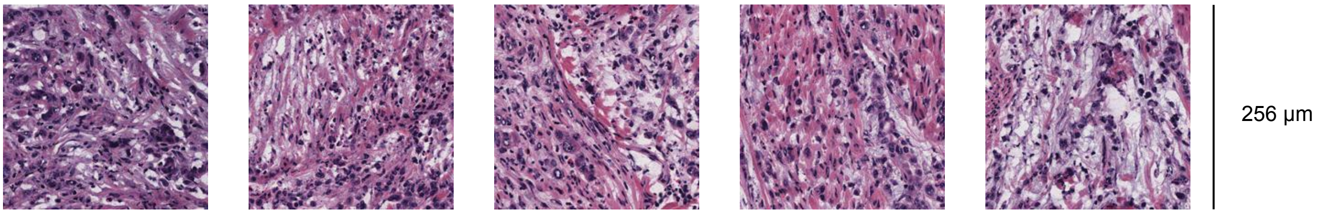


B

Stemness “Low”

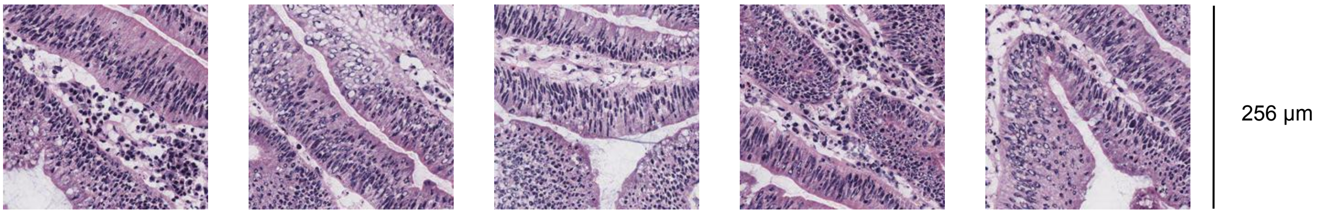
