## Supplemental Tables S1-S5 for "Conventional therapy induces tumor immunoediting and modulates the immune contexture in colorectal cancer"

| patient | timepoint | clone_id | clonal_prev |
|---|---|---|---|
| Pat_03 | Biopsy | 0 | 0.0835191 |
| Pat_03 | Biopsy | 1 | 0.201139 |
| Pat_03 | Biopsy | 2 | 0.230537 |
| Pat_03 | Biopsy | 3 | 0.484805 |
| Pat_03 | Surgery | 0 | 0.12163 |
| Pat_03 | Surgery | 1 | 0.131216 |
| Pat_03 | Surgery | 2 | 0.170884 |
| Pat_03 | Surgery | 3 | 0.576271 |
| Pat_05 | Biopsy | 0 | 0.0849675 |
| Pat_05 | Biopsy | 1 | 0.182607 |
| Pat_05 | Biopsy | 2 | 0.102763 |
| Pat_05 | Biopsy | 3 | 0.0331252 |
| Pat_05 | Biopsy | 4 | 7.68E-10 |
| Pat_05 | Biopsy | 5 | 5.97E-10 |
| Pat_05 | Biopsy | 6 | 4.44E-10 |
| Pat_05 | Biopsy | 7 | 4.15E-10 |
| Pat_05 | Biopsy | 8 | 8.83E-10 |
| Pat_05 | Biopsy | 9 | 0.272971 |
| Pat_05 | Biopsy | 10 | 0.323567 |
| Pat_05 | Surgery | 0 | 0.0612499 |
| Pat_05 | Surgery | 1 | 0.176811 |
| Pat_05 | Surgery | 2 | 7.87E-09 |
| Pat_05 | Surgery | 3 | 7.84E-09 |
| Pat_05 | Surgery | 4 | 0.0725107 |
| Pat_05 | Surgery | 5 | 0.0285134 |
| Pat_05 | Surgery | 6 | 0.10824 |
| Pat_05 | Surgery | 7 | 0.141444 |
| Pat_05 | Surgery | 8 | 0.0778165 |
| Pat_05 | Surgery | 9 | 3.60E-09 |
| Pat_05 | Surgery | 10 | 0.333415 |
| Pat_06 | Biopsy | 0 | 0.00489879 |
| Pat_06 | Biopsy | 1 | 0.554547 |
| Pat_06 | Biopsy | 2 | 0.440554 |
| Pat_06 | Surgery | 0 | 0.00378225 |
| Pat_06 | Surgery | 1 | 0.419722 |
| Pat_06 | Surgery | 2 | 0.576496 |
| Pat_07 | Biopsy | 0 | 0.0223506 |
| Pat_07 | Biopsy | 1 | 1.50E-09 |
| Pat_07 | Biopsy | 2 | 9.79E-10 |
| Pat_07 | Biopsy | 3 | 6.30E-10 |
| Pat_07 | Biopsy | 4 | 7.80E-10 |
| Pat_07 | Biopsy | 5 | 0.181081 |
| Pat_07 | Biopsy | 6 | 0.567449 |
| Pat_07 | Biopsy | 7 | 0.229119 |
| Pat_07 | Surgery | 0 | 0.00578721 |
| Pat_07 | Surgery | 1 | 7.43E-09 |
| Pat_07 | Surgery | 2 | 0.0907203 |
| Pat_07 | Surgery | 3 | 0.0112434 |
| Pat_07 | Surgery | 4 | 0.0960699 |

| | | | |
|---|---|---|---|
| Pat_07 | Surgery | 5 | 7.46E-09 |
| Pat_07 | Surgery | 6 | 0.261586 |
| Pat_07 | Surgery | 7 | 0.534593 |
| Pat_08 | Biopsy | 0 | 0.0851841 |
| Pat_08 | Biopsy | 1 | 0.460132 |
| Pat_08 | Biopsy | 2 | 0.454684 |
| Pat_08 | Surgery | 0 | 0.742028 |
| Pat_08 | Surgery | 1 | 3.96E-09 |
| Pat_08 | Surgery | 2 | 0.257972 |

| patient | gene | chromosome | coord | variation | clone_id | timepoint |
|---|---|---|---|---|---|---|
| Pat_03 | MTMR9 | chr8 | 11331413 | T/C | 3 | Biopsy |
| Pat_03 | MTMR9 | chr8 | 11331413 | T/C | 3 | Surgery |
| Pat_03 | SERF1A | chr5 | 70905146 | C/A | 1 | Biopsy |
| Pat_03 | SERF1A | chr5 | 70905146 | C/A | 1 | Surgery |
| Pat_03 | TSPEAR | chr21 | 44682221 | G/A | 2 | Biopsy |
| Pat_03 | TSPEAR | chr21 | 44682221 | G/A | 2 | Surgery |
| Pat_03 | ZDHHC8 | chr22 | 20149799 | A/G | 1 | Biopsy |
| Pat_03 | ZDHHC8 | chr22 | 20149799 | A/G | 1 | Surgery |
| Pat_05 | ADAMTSL2 | chr9 | 133568739 | C/A | 1 | Biopsy |
| Pat_05 | ADAMTSL2 | chr9 | 133568739 | C/A | 1 | Surgery |
| Pat_05 | ALPP | chr2 | 232379906 | T/C | 1 | Biopsy |
| Pat_05 | ALPP | chr2 | 232379906 | T/C | 1 | Surgery |
| Pat_05 | APOB | chr2 | 21019013 | T/G | 1 | Biopsy |
| Pat_05 | APOB | chr2 | 21019013 | T/G | 1 | Surgery |
| Pat_05 | CCL4 | chr17 | 36105096 | C/G | 1 | Biopsy |
| Pat_05 | CCL4 | chr17 | 36105096 | C/G | 1 | Surgery |
| Pat_05 | CD300LF | chr17 | 74705156 | G/T | 1 | Biopsy |
| Pat_05 | CD300LF | chr17 | 74705156 | G/T | 1 | Surgery |
| Pat_05 | CERS3 | chr15 | 100501729 | C/T | 1 | Biopsy |
| Pat_05 | CERS3 | chr15 | 100501729 | C/T | 1 | Surgery |
| Pat_05 | CR1 | chr1 | 207614451 | T/C | 1 | Biopsy |
| Pat_05 | CR1 | chr1 | 207614451 | T/C | 1 | Surgery |
| Pat_05 | DCLK1 | chr13 | 35822754 | C/T | 1 | Biopsy |
| Pat_05 | DCLK1 | chr13 | 35822754 | C/T | 1 | Surgery |
| Pat_05 | DOCK2 | chr5 | 170041060 | T/C | 1 | Biopsy |
| Pat_05 | DOCK2 | chr5 | 170041060 | T/C | 1 | Surgery |
| Pat_05 | GABRQ | chrX | 152652829 | C/T | 1 | Biopsy |
| Pat_05 | GABRQ | chrX | 152652829 | C/T | 1 | Surgery |
| Pat_05 | H2BC1 | chr6 | 25727000 | G/A | 1 | Biopsy |
| Pat_05 | H2BC1 | chr6 | 25727000 | G/A | 1 | Surgery |
| Pat_05 | HLA-B | chr6 | 31355089 | C/T | 1 | Biopsy |
| Pat_05 | HLA-B | chr6 | 31355089 | C/T | 1 | Surgery |
| Pat_05 | IGHV3-49 | chr14 | 106557029 | A/G | 1 | Biopsy |
| Pat_05 | IGHV3-49 | chr14 | 106557029 | A/G | 1 | Surgery |
| Pat_05 | INTU | chr4 | 127706911 | G/A | 1 | Biopsy |
| Pat_05 | INTU | chr4 | 127706911 | G/A | 1 | Surgery |
| Pat_05 | IRX2 | chr5 | 2749412 | C/T | 1 | Biopsy |
| Pat_05 | IRX2 | chr5 | 2749412 | C/T | 1 | Surgery |
| Pat_05 | KCNK13 | chr14 | 90184434 | G/A | 1 | Biopsy |
| Pat_05 | KCNK13 | chr14 | 90184434 | G/A | 1 | Surgery |
| Pat_05 | KRTAP19-8 | chr21 | 31038363 | G/A | 1 | Biopsy |
| Pat_05 | KRTAP19-8 | chr21 | 31038363 | G/A | 1 | Surgery |
| Pat_05 | LRRK1 | chr15 | 101065999 | G/T | 1 | Biopsy |
| Pat_05 | LRRK1 | chr15 | 101065999 | G/T | 1 | Surgery |
| Pat_05 | LTN1 | chr21 | 28958530 | A/T | 1 | Biopsy |
| Pat_05 | LTN1 | chr21 | 28958530 | A/T | 1 | Surgery |
| Pat_05 | LZTS1 | chr8 | 20250295 | G/A | 1 | Biopsy |
| Pat_05 | LZTS1 | chr8 | 20250295 | G/A | 1 | Surgery |
| Pat_05 | MSANTD1 | chr4 | 3249290 | GGCGGC/- | 1 | Biopsy |

| | | | | | | |
|---|---|---|---|---|---|---|
| Pat_05 | MSANTD1 | chr4 | 3249290 | GGCGGC/- | 1 | Surgery |
| Pat_05 | NARF | chr17 | 82483649 | C/T | 1 | Biopsy |
| Pat_05 | NARF | chr17 | 82483649 | C/T | 1 | Surgery |
| Pat_05 | NOMO2 | chr16 | 18538509 | C/G | 1 | Biopsy |
| Pat_05 | NOMO2 | chr16 | 18538509 | C/G | 1 | Surgery |
| Pat_05 | NPAS4 | chr11 | 66422176 | G/A | 1 | Biopsy |
| Pat_05 | NPAS4 | chr11 | 66422176 | G/A | 1 | Surgery |
| Pat_05 | PANK3 | chr5 | 168561428 | T/A | 1 | Biopsy |
| Pat_05 | PANK3 | chr5 | 168561428 | T/A | 1 | Surgery |
| Pat_05 | PTPRT | chr20 | 42678150 | G/T | 1 | Biopsy |
| Pat_05 | PTPRT | chr20 | 42678150 | G/T | 1 | Surgery |
| Pat_05 | RYR1 | chr19 | 38523031 | G/A | 1 | Biopsy |
| Pat_05 | RYR1 | chr19 | 38523031 | G/A | 1 | Surgery |
| Pat_05 | SAA2 | chr11 | 18245442 | T/C | 1 | Biopsy |
| Pat_05 | SAA2 | chr11 | 18245442 | T/C | 1 | Surgery |
| Pat_05 | SPATA5L1 | chr15 | 45402592 | G/A | 1 | Biopsy |
| Pat_05 | SPATA5L1 | chr15 | 45402592 | G/A | 1 | Surgery |
| Pat_05 | SYTL2 | chr11 | 85726258 | G/A | 1 | Biopsy |
| Pat_05 | SYTL2 | chr11 | 85726258 | G/A | 1 | Surgery |
| Pat_05 | TMED7-TICAM2 | chr5 | 115580995 | C/A | 1 | Biopsy |
| Pat_05 | TMED7-TICAM2 | chr5 | 115580995 | C/A | 1 | Surgery |
| Pat_05 | YOD1 | chr1 | 207049529 | G/T | 1 | Biopsy |
| Pat_05 | YOD1 | chr1 | 207049529 | G/T | 1 | Surgery |
| Pat_05 | ZNF398 | chr7 | 149179464 | G/A | 1 | Biopsy |
| Pat_05 | ZNF398 | chr7 | 149179464 | G/A | 1 | Surgery |
| Pat_05 | ZNRF3 | chr22 | 29044883 | G/A | 1 | Biopsy |
| Pat_05 | ZNRF3 | chr22 | 29044883 | G/A | 1 | Surgery |
| Pat_05 | ALAS1 | chr3 | 52204000 | A/G | 2 | Biopsy |
| Pat_05 | ALAS1 | chr3 | 52204000 | A/G | 2 | Surgery |
| Pat_05 | ANKRD28 | chr3 | 15713630 | C/T | 2 | Biopsy |
| Pat_05 | ANKRD28 | chr3 | 15713630 | C/T | 2 | Surgery |
| Pat_05 | BABAM1 | chr19 | 17282969 | G/A | 2 | Biopsy |
| Pat_05 | BABAM1 | chr19 | 17282969 | G/A | 2 | Surgery |
| Pat_05 | BTBD18 | chr11 | 57751070 | G/A | 2 | Biopsy |
| Pat_05 | BTBD18 | chr11 | 57751070 | G/A | 2 | Surgery |
| Pat_05 | CENPJ | chr13 | 24909879 | G/A | 2 | Biopsy |
| Pat_05 | CENPJ | chr13 | 24909879 | G/A | 2 | Surgery |
| Pat_05 | CTTNBP2 | chr7 | 117756537 | C/G | 2 | Biopsy |
| Pat_05 | CTTNBP2 | chr7 | 117756537 | C/G | 2 | Surgery |
| Pat_05 | DDX11 | chr12 | 31085044 | C/T | 2 | Biopsy |
| Pat_05 | DDX11 | chr12 | 31085044 | C/T | 2 | Surgery |
| Pat_05 | EVC | chr4 | 5708554 | C/T | 2 | Biopsy |
| Pat_05 | EVC | chr4 | 5708554 | C/T | 2 | Surgery |
| Pat_05 | FAT4 | chr4 | 125452472 | G/A | 2 | Biopsy |
| Pat_05 | FAT4 | chr4 | 125452472 | G/A | 2 | Surgery |
| Pat_05 | KCNA5 | chr12 | 5045909 | G/A | 2 | Biopsy |
| Pat_05 | KCNA5 | chr12 | 5045909 | G/A | 2 | Surgery |
| Pat_05 | LRIG1 | chr3 | 66462361 | C/T | 2 | Biopsy |
| Pat_05 | LRIG1 | chr3 | 66462361 | C/T | 2 | Surgery |
| Pat_05 | MRAS | chr3 | 138397401 | G/A | 2 | Biopsy |

| | | | | | | |
|---|---|---|---|---|---|---|
| Pat_05 | MRAS | chr3 | 138397401 | G/A | 2 | Surgery |
| Pat_05 | MUC16 | chr19 | 8904806 | C/A | 2 | Biopsy |
| Pat_05 | MUC16 | chr19 | 8904806 | C/A | 2 | Surgery |
| Pat_05 | NKX6-2 | chr10 | 132785947 | A/C | 2 | Biopsy |
| Pat_05 | NKX6-2 | chr10 | 132785947 | A/C | 2 | Surgery |
| Pat_05 | NLRP5 | chr19 | 56027880 | C/T | 2 | Biopsy |
| Pat_05 | NLRP5 | chr19 | 56027880 | C/T | 2 | Surgery |
| Pat_05 | PLCXD1 | chrX | 299233 | G/T | 2 | Biopsy |
| Pat_05 | PLCXD1 | chrX | 299233 | G/T | 2 | Surgery |
| Pat_05 | PTBP1 | chr19 | 812935 | C/T | 2 | Biopsy |
| Pat_05 | PTBP1 | chr19 | 812935 | C/T | 2 | Surgery |
| Pat_05 | PTPN14 | chr1 | 214376307 | C/T | 2 | Biopsy |
| Pat_05 | PTPN14 | chr1 | 214376307 | C/T | 2 | Surgery |
| Pat_05 | PZP | chr12 | 9202351 | C/T | 2 | Biopsy |
| Pat_05 | PZP | chr12 | 9202351 | C/T | 2 | Surgery |
| Pat_05 | RPS6KA4 | chr11 | 64368382 | C/T | 2 | Biopsy |
| Pat_05 | RPS6KA4 | chr11 | 64368382 | C/T | 2 | Surgery |
| Pat_05 | RXFP1 | chr4 | 158607942 | T/A | 2 | Biopsy |
| Pat_05 | RXFP1 | chr4 | 158607942 | T/A | 2 | Surgery |
| Pat_05 | SLC39A4 | chr8 | 144416010 | A/T | 2 | Biopsy |
| Pat_05 | SLC39A4 | chr8 | 144416010 | A/T | 2 | Surgery |
| Pat_05 | SPTBN1 | chr2 | 54631243 | C/A | 2 | Biopsy |
| Pat_05 | SPTBN1 | chr2 | 54631243 | C/A | 2 | Surgery |
| Pat_05 | THBD | chr20 | 23047903 | C/T | 2 | Biopsy |
| Pat_05 | THBD | chr20 | 23047903 | C/T | 2 | Surgery |
| Pat_05 | TIAL1 | chr10 | 119588278 | A/T | 2 | Biopsy |
| Pat_05 | TIAL1 | chr10 | 119588278 | A/T | 2 | Surgery |
| Pat_05 | TP53 | chr17 | 7674220 | C/T | 2 | Biopsy |
| Pat_05 | TP53 | chr17 | 7674220 | C/T | 2 | Surgery |
| Pat_05 | TTC22 | chr1 | 54781761 | C/T | 2 | Biopsy |
| Pat_05 | TTC22 | chr1 | 54781761 | C/T | 2 | Surgery |
| Pat_05 | ZNF682 | chr19 | 20006623 | C/T | 2 | Biopsy |
| Pat_05 | ZNF682 | chr19 | 20006623 | C/T | 2 | Surgery |
| Pat_05 | ESX1 | chrX | 104254872 | C/T | 3 | Biopsy |
| Pat_05 | ESX1 | chrX | 104254872 | C/T | 3 | Surgery |
| Pat_05 | GFPT2 | chr5 | 180330820 | G/T | 4 | Biopsy |
| Pat_05 | GFPT2 | chr5 | 180330820 | G/T | 4 | Surgery |
| Pat_05 | HRH3 | chr20 | 62216267 | C/T | 5 | Biopsy |
| Pat_05 | HRH3 | chr20 | 62216267 | C/T | 5 | Surgery |
| Pat_05 | NOC2L | chr1 | 960655 | G/A | 6 | Biopsy |
| Pat_05 | NOC2L | chr1 | 960655 | G/A | 6 | Surgery |
| Pat_05 | GPAT2 | chr2 | 96022586 | G/T | 7 | Biopsy |
| Pat_05 | GPAT2 | chr2 | 96022586 | G/T | 7 | Surgery |
| Pat_05 | SLITRK4 | chrX | 143628944 | C/A | 8 | Biopsy |
| Pat_05 | SLITRK4 | chrX | 143628944 | C/A | 8 | Surgery |
| Pat_05 | TEX14 | chr17 | 58599049 | C/A | 9 | Biopsy |
| Pat_05 | TEX14 | chr17 | 58599049 | C/A | 9 | Surgery |
| Pat_05 | ACBD3 | chr1 | 226152367 | C/A | 10 | Biopsy |
| Pat_05 | ACBD3 | chr1 | 226152367 | C/A | 10 | Surgery |
| Pat_05 | SELE | chr1 | 169727691 | T/A | 11 | Biopsy |

| | | | | | | |
|---|---|---|---|---|---|---|
| Pat_05 | SELE | chr1 | 169727691 | T/A | 11 | Surgery |
| Pat_06 | ABTB2 | chr11 | 34171045 | C/T | 1 | Biopsy |
| Pat_06 | ABTB2 | chr11 | 34171045 | C/T | 1 | Surgery |
| Pat_06 | AC011484.1 | chr19 | 46494830 | G/A | 1 | Biopsy |
| Pat_06 | AC011484.1 | chr19 | 46494830 | G/A | 1 | Surgery |
| Pat_06 | ACTRT2 | chr1 | 3022383 | C/T | 1 | Biopsy |
| Pat_06 | ACTRT2 | chr1 | 3022383 | C/T | 1 | Surgery |
| Pat_06 | AK5 | chr1 | 77558673 | -/CA | 1 | Biopsy |
| Pat_06 | AK5 | chr1 | 77558673 | -/CA | 1 | Surgery |
| Pat_06 | ANO1 | chr11 | 70111759 | C/A | 1 | Biopsy |
| Pat_06 | ANO1 | chr11 | 70111759 | C/A | 1 | Surgery |
| Pat_06 | C4orf54 | chr4 | 99653453 | A/T | 1 | Biopsy |
| Pat_06 | C4orf54 | chr4 | 99653453 | A/T | 1 | Surgery |
| Pat_06 | CA8 | chr8 | 60232313 | C/T | 1 | Biopsy |
| Pat_06 | CA8 | chr8 | 60232313 | C/T | 1 | Surgery |
| Pat_06 | CDH8 | chr16 | 61713946 | C/G | 1 | Biopsy |
| Pat_06 | CDH8 | chr16 | 61713946 | C/G | 1 | Surgery |
| Pat_06 | CEACAM7 | chr19 | 41687138 | C/A | 1 | Biopsy |
| Pat_06 | CEACAM7 | chr19 | 41687138 | C/A | 1 | Surgery |
| Pat_06 | CSDE1 | chr1 | 114713909 | G/T | 1 | Biopsy |
| Pat_06 | CSDE1 | chr1 | 114713909 | G/T | 1 | Surgery |
| Pat_06 | CWC22 | chr2 | 179950617 | C/G | 1 | Biopsy |
| Pat_06 | CWC22 | chr2 | 179950617 | C/G | 1 | Surgery |
| Pat_06 | CYB5D2 | chr17 | 4156933 | C/T | 1 | Biopsy |
| Pat_06 | CYB5D2 | chr17 | 4156933 | C/T | 1 | Surgery |
| Pat_06 | DCD | chr12 | 54648330 | C/A | 1 | Biopsy |
| Pat_06 | DCD | chr12 | 54648330 | C/A | 1 | Surgery |
| Pat_06 | DES | chr2 | 219420592 | G/A | 1 | Biopsy |
| Pat_06 | DES | chr2 | 219420592 | G/A | 1 | Surgery |
| Pat_06 | DNAH1 | chr3 | 52368769 | C/T | 1 | Biopsy |
| Pat_06 | DNAH1 | chr3 | 52368769 | C/T | 1 | Surgery |
| Pat_06 | ESYT3 | chr3 | 138460655 | G/A | 1 | Biopsy |
| Pat_06 | ESYT3 | chr3 | 138460655 | G/A | 1 | Surgery |
| Pat_06 | FBXW7 | chr4 | 152326013 | G/T | 1 | Biopsy |
| Pat_06 | FBXW7 | chr4 | 152326013 | G/T | 1 | Surgery |
| Pat_06 | FREM2 | chr13 | 38857970 | C/T | 1 | Biopsy |
| Pat_06 | FREM2 | chr13 | 38857970 | C/T | 1 | Surgery |
| Pat_06 | GRIK2 | chr6 | 102035516 | A/T | 1 | Biopsy |
| Pat_06 | GRIK2 | chr6 | 102035516 | A/T | 1 | Surgery |
| Pat_06 | H1-5 | chr6 | 27867382 | C/T | 1 | Biopsy |
| Pat_06 | H1-5 | chr6 | 27867382 | C/T | 1 | Surgery |
| Pat_06 | HS3ST2 | chr16 | 22915026 | C/T | 1 | Biopsy |
| Pat_06 | HS3ST2 | chr16 | 22915026 | C/T | 1 | Surgery |
| Pat_06 | IGHV3-7 | chr14 | 106062229 | T/C | 1 | Biopsy |
| Pat_06 | IGHV3-7 | chr14 | 106062229 | T/C | 1 | Surgery |
| Pat_06 | INPPL1 | chr11 | 72241182 | A/T | 1 | Biopsy |
| Pat_06 | INPPL1 | chr11 | 72241182 | A/T | 1 | Surgery |
| Pat_06 | JAKMIP2 | chr5 | 147644990 | G/A | 1 | Biopsy |
| Pat_06 | JAKMIP2 | chr5 | 147644990 | G/A | 1 | Surgery |
| Pat_06 | KRTAP19-4 | chr21 | 30501968 | G/A | 1 | Biopsy |

| | | | | | | |
|---|---|---|---|---|---|---|
| Pat_06 | KRTAP19-4 | chr21 | 30501968 | G/A | 1 | Surgery |
| Pat_06 | LRRC55 | chr11 | 57182339 | G/A | 1 | Biopsy |
| Pat_06 | LRRC55 | chr11 | 57182339 | G/A | 1 | Surgery |
| Pat_06 | MGAT4C | chr12 | 85979674 | A/C | 1 | Biopsy |
| Pat_06 | MGAT4C | chr12 | 85979674 | A/C | 1 | Surgery |
| Pat_06 | MROH2B | chr5 | 41007305 | G/A | 1 | Biopsy |
| Pat_06 | MROH2B | chr5 | 41007305 | G/A | 1 | Surgery |
| Pat_06 | MS4A12 | chr11 | 60497392 | TT/AA | 1 | Biopsy |
| Pat_06 | MS4A12 | chr11 | 60497392 | TT/AA | 1 | Surgery |
| Pat_06 | MYF5 | chr12 | 80717395 | C/T | 1 | Biopsy |
| Pat_06 | MYF5 | chr12 | 80717395 | C/T | 1 | Surgery |
| Pat_06 | NECTIN1 | chr11 | 119677683 | T/C | 1 | Biopsy |
| Pat_06 | NECTIN1 | chr11 | 119677683 | T/C | 1 | Surgery |
| Pat_06 | NEU2 | chr2 | 233034762 | G/A | 1 | Biopsy |
| Pat_06 | NEU2 | chr2 | 233034762 | G/A | 1 | Surgery |
| Pat_06 | PCDHGA3 | chr5 | 141419761 | C/T | 1 | Biopsy |
| Pat_06 | PCDHGA3 | chr5 | 141419761 | C/T | 1 | Surgery |
| Pat_06 | PLEC | chr8 | 143918224 | C/T | 1 | Biopsy |
| Pat_06 | PLEC | chr8 | 143918224 | C/T | 1 | Surgery |
| Pat_06 | PNISR | chr6 | 99401039 | C/T | 1 | Biopsy |
| Pat_06 | PNISR | chr6 | 99401039 | C/T | 1 | Surgery |
| Pat_06 | PODNL1 | chr19 | 13933257 | G/A | 1 | Biopsy |
| Pat_06 | PODNL1 | chr19 | 13933257 | G/A | 1 | Surgery |
| Pat_06 | POTEC | chr18 | 14542891 | G/T | 1 | Biopsy |
| Pat_06 | POTEC | chr18 | 14542891 | G/T | 1 | Surgery |
| Pat_06 | PRKAG3 | chr2 | 218827915 | C/T | 1 | Biopsy |
| Pat_06 | PRKAG3 | chr2 | 218827915 | C/T | 1 | Surgery |
| Pat_06 | PRKCG | chr19 | 53897966 | G/A | 1 | Biopsy |
| Pat_06 | PRKCG | chr19 | 53897966 | G/A | 1 | Surgery |
| Pat_06 | PSME2 | chr14 | 24148748 | G/A | 1 | Biopsy |
| Pat_06 | PSME2 | chr14 | 24148748 | G/A | 1 | Surgery |
| Pat_06 | RUNX1T1 | chr8 | 91986878 | A/T | 1 | Biopsy |
| Pat_06 | RUNX1T1 | chr8 | 91986878 | A/T | 1 | Surgery |
| Pat_06 | SERPINA4 | chr14 | 94569462 | C/T | 1 | Biopsy |
| Pat_06 | SERPINA4 | chr14 | 94569462 | C/T | 1 | Surgery |
| Pat_06 | SH2D3A | chr19 | 6754157 | G/A | 1 | Biopsy |
| Pat_06 | SH2D3A | chr19 | 6754157 | G/A | 1 | Surgery |
| Pat_06 | SH2D3A | chr19 | 6755075 | G/A | 1 | Biopsy |
| Pat_06 | SH2D3A | chr19 | 6755075 | G/A | 1 | Surgery |
| Pat_06 | SLC27A5 | chr19 | 58500470 | T/A | 1 | Biopsy |
| Pat_06 | SLC27A5 | chr19 | 58500470 | T/A | 1 | Surgery |
| Pat_06 | SLC6A20 | chr3 | 45759118 | C/T | 1 | Biopsy |
| Pat_06 | SLC6A20 | chr3 | 45759118 | C/T | 1 | Surgery |
| Pat_06 | SNX19 | chr11 | 130903296 | C/A | 1 | Biopsy |
| Pat_06 | SNX19 | chr11 | 130903296 | C/A | 1 | Surgery |
| Pat_06 | SORCS2 | chr4 | 7726844 | C/T | 1 | Biopsy |
| Pat_06 | SORCS2 | chr4 | 7726844 | C/T | 1 | Surgery |
| Pat_06 | STARD13 | chr13 | 33129281 | G/A | 1 | Biopsy |
| Pat_06 | STARD13 | chr13 | 33129281 | G/A | 1 | Surgery |
| Pat_06 | TBX18 | chr6 | 84748038 | C/T | 1 | Biopsy |

| | | | | | | |
|---|---|---|---|---|---|---|
| Pat_06 | TBX18 | chr6 | 84748038 | C/T | 1 | Surgery |
| Pat_06 | TGFA | chr2 | 70456401 | G/A | 1 | Biopsy |
| Pat_06 | TGFA | chr2 | 70456401 | G/A | 1 | Surgery |
| Pat_06 | TMEM121 | chr14 | 105529161 | G/A | 1 | Biopsy |
| Pat_06 | TMEM121 | chr14 | 105529161 | G/A | 1 | Surgery |
| Pat_06 | TRGC1 | chr7 | 38262190 | A/T | 1 | Biopsy |
| Pat_06 | TRGC1 | chr7 | 38262190 | A/T | 1 | Surgery |
| Pat_06 | USP17L2 | chr8 | 12138537 | G/T | 1 | Biopsy |
| Pat_06 | USP17L2 | chr8 | 12138537 | G/T | 1 | Surgery |
| Pat_06 | WSCD2 | chr12 | 108210260 | C/T | 1 | Biopsy |
| Pat_06 | WSCD2 | chr12 | 108210260 | C/T | 1 | Surgery |
| Pat_06 | ST18 | chr8 | 52214207 | G/A | 2 | Biopsy |
| Pat_06 | ST18 | chr8 | 52214207 | G/A | 2 | Surgery |
| Pat_07 | PIAS3 | chr1 | 145859052 | C/A | 1 | Surgery |
| Pat_07 | PIAS3 | chr1 | 145859052 | C/A | 1 | Biopsy |
| Pat_07 | CASZ1 | chr1 | 10639655 | G/A | 2 | Surgery |
| Pat_07 | CASZ1 | chr1 | 10639655 | G/A | 2 | Biopsy |
| Pat_07 | ANKRD36C | chr2 | 95962232 | A/T | 3 | Surgery |
| Pat_07 | ANKRD36C | chr2 | 95962232 | A/T | 3 | Biopsy |
| Pat_07 | ZDHHC8 | chr22 | 20149799 | A/G | 3 | Surgery |
| Pat_07 | ZDHHC8 | chr22 | 20149799 | A/G | 3 | Biopsy |
| Pat_07 | ITGA2 | chr5 | 52989422 | A/G | 4 | Surgery |
| Pat_07 | ITGA2 | chr5 | 52989422 | A/G | 4 | Biopsy |
| Pat_07 | KCMF1 | chr2 | 84971469 | T/A | 5 | Surgery |
| Pat_07 | KCMF1 | chr2 | 84971469 | T/A | 5 | Biopsy |
| Pat_07 | TYRO3 | chr15 | 41565109 | G/A | 6 | Surgery |
| Pat_07 | TYRO3 | chr15 | 41565109 | G/A | 6 | Biopsy |
| Pat_07 | KCNG2 | chr18 | 79899362 | G/A | 7 | Surgery |
| Pat_07 | KCNG2 | chr18 | 79899362 | G/A | 7 | Biopsy |
| Pat_08 | PCDHGA3 | chr5 | 141361662 | G/A | 1 | Surgery |
| Pat_08 | PCDHGA3 | chr5 | 141361662 | G/A | 1 | Biopsy |
| Pat_08 | SLC24A2 | chr9 | 19785969 | C/T | 1 | Surgery |
| Pat_08 | SLC24A2 | chr9 | 19785969 | C/T | 1 | Biopsy |
| Pat_08 | SGF29 | chr16 | 28592475 | T/C | 2 | Surgery |
| Pat_08 | SGF29 | chr16 | 28592475 | T/C | 2 | Biopsy |

| VAF |
|---|
| 0.27027027 |
| 0.17692307 |
| 0.10169491 |
| 0.10729613 |
| 0.23636363 |
| 0.203125 |
| 0.29629629 |
| 0.16 |
| 0.11578947 |
| 0.10738255 |
| 0.04878048 |
| 0.08771929 |
| 0.11009174 |
| 0.06451612 |
| 0.08163265 |
| 0.111111111 |
| 0.08849557 |
| 0.1 |
| 0.07438016 |
| 0.07964601 |
| 0.05508474 |
| 0.04363636 |
| 0.15384615 |
| 0.08108108 |
| 0.06153846 |
| 0.07865168 |
| 0.16091954 |
| 0.08 |
| 0.08823529 |
| 0.06976744 |
| 0.08163265 |
| 0.09302325 |
| 0.07575757 |
| 0.07594936 |
| 0.16197183 |
| 0.11038961 |
| 0.04040404 |
| 0.12334801 |
| 0.06617647 |
| 0.11392405 |
| 0.07017543 |
| 0.12578616 |
| 0.04048582 |
| 0.10196078 |
| 0.09638554 |
| 0.11428571 |
| 0.14634146 |
| 0.06956521 |
| 0.09411764 |

| |
|---|
| 0.09794988 |
| 0.10294117 |
| 0.11538461 |
| 0.08558558 |
| 0.06802721 |
| 0.08154506 |
| 0.11803278 |
| 0.07329842 |
| 0.06451612 |
| 0.10714285 |
| 0.05714285 |
| 0.111111111 |
| 0.10588235 |
| 0.07741935 |
| 0.04761904 |
| 0.06034482 |
| 0.13450292 |
| 0.07142857 |
| 0.09467455 |
| 0.08396946 |
| 0.05154639 |
| 0.05487804 |
| 0.06122448 |
| 0.04207920 |
| 0.12378640 |
| 0.13385826 |
| 0.23214285 |
| 0.16393442 |
| 0.17808219 |
| 0.17355371 |
| 0.20325203 |
| 0.18571428 |
| 0.2 |
| 0.08823529 |
| 0.18446601 |
| 0.28947368 |
| 0.24827586 |
| 0.17808219 |
| 0.09589041 |
| 0.15789473 |
| 0.14634146 |
| 0.13636363 |
| 0.15652173 |
| 0.14814814 |
| 0.13970588 |
| 0.16666666 |
| 0.13475177 |
| 0.17857142 |
| 0.12820512 |
| 0.14035087 |

| |
|---|
| 0.15671641 |
| 0.17073170 |
| 0.1625 |
| 0.28571428 |
| 0.14173228 |
| 0.13709677 |
| 0.17054263 |
| 0.13666666 |
| 0.17588932 |
| 0.13207547 |
| 0.20224719 |
| 0.09821428 |
| 0.19475655 |
| 0.13725490 |
| 0.18897637 |
| 0.33333333 |
| 0.21568627 |
| 0.20338983 |
| 0.27450980 |
| 0.17333333 |
| 0.21311475 |
| 0.20918367 |
| 0.16597510 |
| 0.27722772 |
| 0.17674418 |
| 0.17142857 |
| 0.17821782 |
| 0.20121951 |
| 0.17266187 |
| 0.13114754 |
| 0.22488038 |
| 0.11818181 |
| 0.14840989 |
| 0.12 |
| 0.13333333 |
| 0.12871287 |
| 0.12173913 |
| 0.23580786 |
| 0.28712871 |
| 0.3 |
| 0.08450704 |
| 0.09523809 |
| 0.15254237 |
| 0.19444444 |
| 0.32894736 |
| 0.18390804 |
| 0.05970149 |
| 0.07382550 |
| 0.18652849 |
| 0.12068965 |

| |
|---|
| 0.16666666 |
| 0.38709677 |
| 0.24822695 |
| 0.53846153 |
| 0.45294117 |
| 0.83333333 |
| 0.39473684 |
| 0.31707317 |
| 0.22222222 |
| 0.31372549 |
| 0.21951219 |
| 0.375 |
| 0.19178082 |
| 0.61818181 |
| 0.22641509 |
| 0.26315789 |
| 0.13186813 |
| 0.69387755 |
| 0.36423841 |
| 0.59722222 |
| 0.375 |
| 0.31428571 |
| 0.2 |
| 0.66666666 |
| 0.35964912 |
| 0.28 |
| 0.30693069 |
| 0.4 |
| 0.37719298 |
| 0.5 |
| 0.15909090 |
| 0.4 |
| 0.33587786 |
| 0.37931034 |
| 0.20987654 |
| 0.33333333 |
| 0.15217391 |
| 0.36923076 |
| 0.2875 |
| 0.66153846 |
| 0.16470588 |
| 0.23333333 |
| 0.15476190 |
| 0.15254237 |
| 0.18272425 |
| 0.34482758 |
| 0.23529411 |
| 0.52272727 |
| 0.20183486 |
| 0.51851851 |

| |
|---|
| 0.38926174 |
| 0.45833333 |
| 0.17687074 |
| 0.40350877 |
| 0.25698324 |
| 0.46666666 |
| 0.19318181 |
| 0.31707317 |
| 0.23076923 |
| 0.35 |
| 0.15882352 |
| 0.41463414 |
| 0.31325301 |
| 0.39130434 |
| 0.26666666 |
| 0.30769230 |
| 0.28333333 |
| 0.71641791 |
| 0.38596491 |
| 0.40540540 |
| 0.17592592 |
| 0.2 |
| 0.12994350 |
| 0.26785714 |
| 0.23825503 |
| 0.45714285 |
| 0.16346153 |
| 1 |
| 0.22641509 |
| 0.47826086 |
| 0.25 |
| 0.57142857 |
| 0.28571428 |
| 0.55172413 |
| 0.30405405 |
| 0.60377358 |
| 0.33333333 |
| 0.54716981 |
| 0.32110091 |
| 0.22826086 |
| 0.12953367 |
| 0.2 |
| 0.17361111 |
| 0.60465116 |
| 0.29931972 |
| 0.26923076 |
| 0.17560975 |
| 0.38461538 |
| 0.33333333 |
| 0.19444444 |

| |
|---|
| 0.22641509 |
| 0.33333333 |
| 0.25 |
| 0.40476190 |
| 0.22047244 |
| 0.17142857 |
| 0.15789473 |
| 0.41666666 |
| 0.17010309 |
| 0.42857142 |
| 0.29770992 |
| 0.10169491 |
| 0.15625 |
| 0.05487804 |
| 0.15384615 |
| 0.04040404 |
| 0.11392405 |
| 0.64788732 |
| 0.72727272 |
| 0.22222222 |
| 0.19512195 |
| 0.03663003 |
| 0.24299065 |
| 0.05185185 |
| 0.2 |
| 0.08602150 |
| 0.05660377 |
| 0.07272727 |
| 0.16666666 |
| 0.02024922 |
| 0.38829787 |
| 0.03083700 |
| 0.15306122 |
| 0.07456140 |
| 0.11450381 |

| gene_id | baseMean | log2FoldChange | lfcSE | stat | pvalue | padj | weight |
|---|---|---|---|---|---|---|---|
| ENSG00000143 | 829.392049 | 5.264764326 | 0.371896573 | 14.15652821 | 1.70E-45 | 2.48E-41 | 1.597450288 |
| ENSG00000133 | 7149.80236 | 4.399850431 | 0.3249676919 | 13.53934727 | 9.16E-42 | 6.64E-38 | 1.602205449 |
| ENSG00000175 | 2399.439396 | 6.229820958 | 0.4618975275 | 13.48745249 | 1.85E-41 | 8.99E-38 | 1.597450288 |
| ENSG00000172 | 2119.985142 | 3.563595445 | 0.2898378649 | 12.2951342 | 9.62E-35 | 3.56E-31 | 1.572476891 |
| ENSG00000182 | 467.2973867 | 4.762597633 | 0.3922651969 | 12.14126991 | 6.38E-34 | 1.63E-30 | 1.825593782 |
| ENSG00000130 | 5128.353239 | 4.178081485 | 0.3524245096 | 11.85525232 | 2.02E-32 | 4.79E-29 | 1.634803475 |
| ENSG00000163 | 4332.518506 | 4.516380037 | 0.406827291 | 11.10146772 | 1.23E-28 | 2.61E-25 | 1.572476891 |
| ENSG00000170 | 276.8942531 | 6.303002735 | 0.5889927481 | 10.70132486 | 1.00E-26 | 6.99E-23 | 0.3706356727 |
| ENSG00000111 | 5888.214211 | 3.792810308 | 0.3590340085 | 10.56393049 | 4.38E-26 | 6.99E-23 | 1.634803475 |
| ENSG00000004 | 781.9519427 | 4.547344267 | 0.4435748287 | 10.25158321 | 1.16E-24 | 1.69E-21 | 1.597450288 |
| ENSG00000149 | 9243.48496 | 3.10070567 | 0.3079779175 | 10.06794804 | 7.66E-24 | 8.50E-21 | 1.902376403 |
| ENSG00000101 | 3655.741398 | 3.188987305 | 0.3175693832 | 10.04186006 | 9.98E-24 | 1.02E-20 | 1.902376403 |
| ENSG00000163 | 348.8408688 | 3.450296211 | 0.3466343363 | 9.95370582 | 2.43E-23 | 3.50E-20 | 1.242301963 |
| ENSG00000072 | 388.5217595 | 3.690542187 | 0.3757011112 | 9.823080308 | 8.96E-23 | 1.20E-19 | 1.242301963 |
| ENSG00000059 | 153.1386444 | 5.400128073 | 0.5514269889 | 9.793006474 | 1.21E-22 | 1.56E-19 | 1.198705341 |
| ENSG00000188 | 331.7957387 | 3.989809676 | 0.4083378722 | 9.770853864 | 1.50E-22 | 1.65E-19 | 1.32213648 |
| ENSG00000154 | 251.4079586 | 5.175064627 | 0.5369687461 | 9.63755277 | 5.55E-22 | 1.93E-18 | 0.3939394882 |
| ENSG00000120 | 1328.687438 | 4.111217958 | 0.4355287438 | 9.439601902 | 3.74E-21 | 3.03E-18 | 1.597450288 |
| ENSG00000145 | 172.7044146 | 4.033961301 | 0.4279242329 | 9.426812018 | 4.23E-21 | 4.31E-18 | 1.198705341 |
| ENSG00000137 | 401.8509142 | 3.85758126 | 0.4148618437 | 9.298472055 | 1.42E-20 | 1.25E-17 | 1.32213648 |
| ENSG00000091 | 1526.535878 | 3.621140149 | 0.3918462472 | 9.241227076 | 2.44E-20 | 1.42E-17 | 1.902376403 |
| ENSG00000128 | 284.3347252 | 3.981512002 | 0.4342540271 | 9.168624245 | 4.79E-20 | 4.07E-17 | 1.242301963 |
| ENSG00000198 | 141.7613133 | 4.126263258 | 0.4562157951 | 9.044542741 | 1.50E-19 | 1.22E-16 | 1.242301963 |
| ENSG00000134 | 378.2126367 | 3.162922158 | 0.3522300918 | 8.97970455 | 2.71E-19 | 2.03E-16 | 1.29680518 |
| ENSG00000162 | 318.1480898 | 2.923957181 | 0.327956037 | 8.915698604 | 4.85E-19 | 3.40E-16 | 1.327567552 |
| ENSG00000124 | 154.3002624 | 3.732988235 | 0.4276744948 | 8.728573436 | 2.58E-18 | 1.79E-15 | 1.198705341 |
| ENSG00000095 | 469.570829 | 2.661136418 | 0.3054013939 | 8.713569981 | 2.94E-18 | 1.58E-15 | 1.602205449 |
| ENSG00000280 | 742.6856402 | 3.432332971 | 0.3939357151 | 8.712926601 | 2.96E-18 | 1.53E-15 | 1.729500602 |
| ENSG00000184 | 136.8067063 | 3.665410659 | 0.4222873672 | 8.679896542 | 3.96E-18 | 2.56E-15 | 1.242301963 |
| ENSG00000198 | 723.5596933 | 2.648899786 | 0.3107412109 | 8.524456021 | 1.54E-17 | 7.57E-15 | 1.572476891 |
| ENSG00000103 | 343.9028239 | -3.842714647 | 0.4567656941 | -8.412879288 | 4.00E-17 | 2.26E-14 | 1.327567552 |
| ENSG00000167 | 96.49980897 | 7.061259511 | 0.8458442205 | 8.348179652 | 6.93E-17 | 4.05E-14 | 1.22159078 |

| | | | | | | | |
|---|---|---|---|---|---|---|---|
| ENSG00000135[illegible] | 130.6170213 | 6.226752895 | 0.7460298751 | 8.346519494 | 7.03E-17 | 4.05E-14 | 1.22159078 |
| ENSG00000079[illegible] | 2301.069799 | 2.699378868 | 0.329797426 | 8.18496039 | 2.72E-16 | 1.18E-13 | 1.572476891 |
| ENSG00000135[illegible] | 462.8219621 | 2.670367987 | 0.3270791862 | 8.164285897 | 3.23E-16 | 1.34E-13 | 1.602205449 |
| ENSG00000104[illegible] | 238.2637904 | 3.860200705 | 0.4733660702 | 8.154789597 | 3.50E-16 | 1.83E-13 | 1.198705341 |
| ENSG00000166[illegible] | 241.7637107 | 4.309785989 | 0.5293206505 | 8.142108163 | 3.88E-16 | 1.98E-13 | 1.198705341 |
| ENSG00000065[illegible] | 4432.637179 | 3.341548463 | 0.4104740103 | 8.140706547 | 3.93E-16 | 1.58E-13 | 1.602205449 |
| ENSG00000196[illegible] | 620.6023996 | 3.77645693 | 0.4676199196 | 8.075911165 | 6.70E-16 | 2.55E-13 | 1.564848717 |
| ENSG00000154[illegible] | 542.5897042 | 2.486101126 | 0.308988115 | 8.04594418 | 8.56E-16 | 3.16E-13 | 1.572476891 |
| ENSG00000237[illegible] | 138.6934532 | 3.348173805 | 0.4238518907 | 7.899395707 | 2.80E-15 | 8.76E-13 | 1.814765822 |
| ENSG00000181[illegible] | 81.33213316 | 6.66147481 | 0.8479776741 | 7.855719571 | 3.97E-15 | 1.58E-12 | 1.393563518 |
| ENSG00000196[illegible] | 4493.02216 | 2.316328143 | 0.2968178926 | 7.803869646 | 6.00E-15 | 2.06E-12 | 1.572476891 |
| ENSG00000132[illegible] | 200.655382 | 3.288815227 | 0.4219382397 | 7.794541754 | 6.46E-15 | 7.95E-12 | 0.3939394882 |
| ENSG00000048[illegible] | 164.4242416 | 4.884444327 | 0.6355658552 | 7.685189956 | 1.53E-14 | 6.50E-12 | 1.242301963 |
| ENSG00000131[illegible] | 429.0870128 | 3.856481331 | 0.5044696108 | 7.644625659 | 2.10E-14 | 7.95E-12 | 1.29680518 |
| ENSG00000189[illegible] | 565.4034956 | 3.694424539 | 0.4848318197 | 7.620012527 | 2.54E-14 | 7.41E-12 | 1.729500602 |
| ENSG00000106[illegible] | 193.0397767 | 3.91503033 | 0.5140233394 | 7.616444682 | 2.61E-14 | 7.41E-12 | 1.814765822 |
| ENSG00000150[illegible] | 65.23903819 | 7.059693026 | 0.9283828299 | 7.604290814 | 2.86E-14 | 9.81E-12 | 1.357988055 |
| ENSG00000164[illegible] | 47.45430428 | 8.427580729 | 1.108954912 | 7.599570223 | 2.97E-14 | 9.59E-12 | 1.470104381 |
| ENSG00000113[illegible] | 56.05722026 | 7.366318445 | 0.9840520494 | 7.48570002 | 7.12E-14 | 2.43E-11 | 1.337480197 |
| ENSG00000101[illegible] | 62.56661644 | 5.247673446 | 0.7043243486 | 7.450648918 | 9.29E-14 | 3.06E-11 | 1.357988055 |
| ENSG00000107[illegible] | 293.3204399 | 2.912623089 | 0.3912437456 | 7.444523067 | 9.73E-14 | 3.23E-11 | 1.32213648 |
| ENSG00000280[illegible] | 176.1554748 | 3.514711886 | 0.4727434954 | 7.434712311 | 1.05E-13 | 3.76E-11 | 1.198705341 |
| ENSG00000215[illegible] | 150.1460483 | 3.226635587 | 0.4422055211 | 7.2966877 | 2.95E-13 | 1.00E-10 | 1.198705341 |
| ENSG00000182[illegible] | 61.12009589 | 5.480572258 | 0.7513542259 | 7.294258911 | 3.00E-13 | 9.35E-11 | 1.357988055 |
| ENSG00000174[illegible] | 71.99487487 | 6.0527207 | 0.8323535336 | 7.271814746 | 3.55E-13 | 9.72E-11 | 1.514054369 |
| ENSG00000101[illegible] | 571.2940009 | -2.309198502 | 0.3201075298 | -7.213821255 | 5.44E-13 | 1.19E-10 | 1.825593782 |
| ENSG00000156[illegible] | 751.4002255 | 2.307734932 | 0.3219429828 | 7.16814795 | 7.60E-13 | 1.87E-10 | 1.602205449 |
| ENSG00000118[illegible] | 34.94972893 | 8.390884622 | 1.175108349 | 7.14051996 | 9.30E-13 | 2.69E-10 | 1.319539707 |
| ENSG00000249[illegible] | 2014.77271 | 2.465494126 | 0.3452869819 | 7.140420157 | 9.30E-13 | 2.25E-10 | 1.602205449 |
| ENSG00000198[illegible] | 783.2687346 | 2.499721358 | 0.3532775236 | 7.075800728 | 1.49E-12 | 3.48E-10 | 1.602205449 |
| ENSG00000149[illegible] | 368.3026729 | 2.583896277 | 0.3672413825 | 7.035961631 | 1.98E-12 | 5.46E-10 | 1.29680518 |
| ENSG00000185[illegible] | 336.1128218 | 2.820861695 | 0.4013276334 | 7.028824981 | 2.08E-12 | 5.55E-10 | 1.32213648 |
| ENSG00000279[illegible] | 80.36475477 | 4.0963723 | 0.5833309061 | 7.022381733 | 2.18E-12 | 5.23E-10 | 1.514054369 |

| | | | | | | | |
|---|---|---|---|---|---|---|---|
| ENSG00000147( | 164.0888433 | 3.984285115 | 0.5675597758 | 7.02002729 | 2.22E-12 | 6.23E-10 | 1.198705341 |
| ENSG00000162' | 709.8926414 | 2.225331906 | 0.3170115304 | 7.019719136 | 2.22E-12 | 5.23E-10 | 1.564848717 |
| ENSG00000163' | 42.07918625 | 7.046250942 | 1.005380605 | 7.00854075 | 2.41E-12 | 5.68E-10 | 1.470104381 |
| ENSG00000068( | 74.28276 | 4.171479291 | 0.5955236731 | 7.004724546 | 2.47E-12 | 6.07E-10 | 1.393563518 |
| ENSG00000167' | 883.1601823 | 2.775942405 | 0.3991384325 | 6.954836164 | 3.53E-12 | 7.45E-10 | 1.572476891 |
| ENSG00000077! | 3978.728853 | 2.474665746 | 0.3564206087 | 6.943105102 | 3.84E-12 | 7.68E-10 | 1.634803475 |
| ENSG00000122: | 135.0791959 | 4.150161154 | 0.5982330493 | 6.937365227 | 3.99E-12 | 1.04E-09 | 1.242301963 |
| ENSG00000167: | 52.76173177 | 4.646584083 | 0.6745527968 | 6.88839199 | 5.64E-12 | 1.32E-09 | 1.357988055 |
| ENSG00000118: | 341.4758104 | 2.086146457 | 0.3028923412 | 6.887418972 | 5.68E-12 | 1.43E-09 | 1.249985076 |
| ENSG00000018( | 62.71469904 | 4.855818927 | 0.710511246 | 6.834260477 | 8.24E-12 | 1.68E-09 | 1.516424702 |
| ENSG00000058( | 725.1189866 | 2.017800697 | 0.2957033833 | 6.823732196 | 8.87E-12 | 1.73E-09 | 1.572476891 |
| ENSG00000174( | 312.7395781 | 2.145411179 | 0.315227732 | 6.805908751 | 1.00E-11 | 2.38E-09 | 1.242301963 |
| ENSG00000217: | 52.36397281 | 7.224337832 | 1.062339845 | 6.800401838 | 1.04E-11 | 2.32E-09 | 1.337480197 |
| ENSG00000145! | 140.5679528 | 2.649138241 | 0.390299223 | 6.787454561 | 1.14E-11 | 1.90E-09 | 1.814765822 |
| ENSG00000122' | 1954.796671 | 1.891937138 | 0.279977989 | 6.757449558 | 1.40E-11 | 2.50E-09 | 1.634803475 |
| ENSG00000183: | 124.0740051 | 4.152443765 | 0.6160271655 | 6.740682875 | 1.58E-11 | 3.60E-09 | 1.242301963 |
| ENSG00000095' | 114.5824119 | -5.087924655 | 0.7553890037 | -6.735502675 | 1.63E-11 | 3.68E-09 | 1.242301963 |
| ENSG00000110: | 424.8234382 | 21.05916836 | 3.132402864 | 6.723007632 | 1.78E-11 | 3.73E-09 | 1.32213648 |
| ENSG00000077: | 1182.802 | 1.945126486 | 0.2897286268 | 6.713615106 | 1.90E-11 | 3.47E-09 | 1.572476891 |
| ENSG00000131' | 259.3917137 | 2.485819273 | 0.3704333622 | 6.710570718 | 1.94E-11 | 1.24E-08 | 0.3939394882 |
| ENSG00000113( | 586.6864948 | 1.966566041 | 0.2950611767 | 6.664943395 | 2.65E-11 | 4.52E-09 | 1.602205449 |
| ENSG00000274: | 39.12871863 | 6.240996433 | 0.9369653998 | 6.660861152 | 2.72E-11 | 5.45E-09 | 1.319539707 |
| ENSG00000135- | 1027.911927 | 2.170536654 | 0.3264127864 | 6.64966798 | 2.94E-11 | 4.96E-09 | 1.602205449 |
| ENSG00000278' | 29.56371404 | 8.091630747 | 1.219797634 | 6.633584555 | 3.28E-11 | 6.40E-09 | 1.337480197 |
| ENSG00000183! | 752.7767089 | 2.089192759 | 0.3151279359 | 6.629665353 | 3.36E-11 | 5.20E-09 | 1.729500602 |
| ENSG00000121! | 148.7658513 | 3.224848668 | 0.4920781545 | 6.553529432 | 5.62E-11 | 1.16E-08 | 1.242301963 |
| ENSG00000243- | 41.33053268 | -7.393048099 | 1.129366703 | -6.546189186 | 5.90E-11 | 1.04E-08 | 1.470104381 |
| ENSG00000125' | 602.5767167 | 2.77931923 | 0.4284694368 | 6.486621894 | 8.78E-11 | 1.27E-08 | 1.729500602 |
| ENSG00000107' | 4355.639725 | 1.823886177 | 0.2815208033 | 6.478690581 | 9.25E-11 | 1.42E-08 | 1.597450288 |
| ENSG00000197' | 1037.729827 | 2.461353765 | 0.3813220923 | 6.454789309 | 1.08E-10 | 1.41E-08 | 1.902376403 |
| ENSG00000143! | 177.2310353 | 2.737117596 | 0.4260316357 | 6.424681563 | 1.32E-10 | 2.62E-08 | 1.22159078 |
| ENSG00000139! | 131.2896126 | 2.737715279 | 0.4283108627 | 6.391888503 | 1.64E-10 | 3.21E-08 | 1.22159078 |
| ENSG00000105- | 328.3753944 | -2.713018003 | 0.4255376845 | -6.375505865 | 1.82E-10 | 3.34E-08 | 1.29680518 |

| | | | | | | | |
|---|---|---|---|---|---|---|---|
| ENSG00000072 | 671.4329681 | 2.169284591 | 0.3434984206 | 6.315268021 | 2.70E-10 | 3.47E-08 | 1.825593782 |
| ENSG00000173 | 223.1214101 | 2.962447296 | 0.4699673315 | 6.303517495 | 2.91E-10 | 1.60E-07 | 0.3939394882 |
| ENSG00000113 | 81.73647091 | 4.887590593 | 0.7793268035 | 6.271554592 | 3.57E-10 | 6.15E-08 | 1.337480197 |
| ENSG00000166 | 187.0020525 | 2.479912622 | 0.3962445088 | 6.258541297 | 3.89E-10 | 4.98E-08 | 1.814765822 |
| ENSG00000100 | 406.7955535 | 2.282706631 | 0.3657416107 | 6.241309613 | 4.34E-10 | 7.83E-08 | 1.249985076 |
| ENSG00000181 | 158.1644019 | 2.925079367 | 0.4717515511 | 6.200465817 | 5.63E-10 | 7.07E-08 | 1.814765822 |
| ENSG00000144 | 276.7932847 | 1.906436982 | 0.3087475918 | 6.174742841 | 6.63E-10 | 3.49E-07 | 0.3706356727 |
| ENSG00000159 | 1594.74918 | 2.11025645 | 0.3431435715 | 6.149777018 | 7.76E-10 | 1.08E-07 | 1.602205449 |
| ENSG00000005 | 150.1931136 | -2.77390533 | 0.4516980972 | -6.141060472 | 8.20E-10 | 1.50E-07 | 1.198705341 |
| ENSG00000011 | 1388.556722 | 1.92696252 | 0.3160405457 | 6.097200331 | 1.08E-09 | 1.50E-07 | 1.597450288 |
| ENSG00000165 | 38.76615945 | 5.988783746 | 0.9836548692 | 6.088297769 | 1.14E-09 | 1.81E-07 | 1.357988055 |
| ENSG00000270 | 27.82967839 | 7.147473793 | 1.175241417 | 6.081706863 | 1.19E-09 | 1.82E-07 | 1.393563518 |
| ENSG00000262 | 216.9593824 | -2.62766476 | 0.4330315186 | -6.068068136 | 1.29E-09 | 2.26E-07 | 1.198705341 |
| ENSG00000197 | 507.8493061 | 1.748976643 | 0.2885799199 | 6.060631812 | 1.36E-09 | 1.82E-07 | 1.572476891 |
| ENSG00000197 | 207.9304443 | 2.410964889 | 0.3980587428 | 6.056806772 | 1.39E-09 | 6.86E-07 | 0.3706356727 |
| ENSG00000266 | 106.7575917 | 3.581168225 | 0.5918056556 | 6.051257185 | 1.44E-09 | 2.44E-07 | 1.22159078 |
| ENSG00000113 | 165.2609971 | 2.97804083 | 0.4931606668 | 6.038682787 | 1.55E-09 | 2.62E-07 | 1.198705341 |
| ENSG00000169 | 66.76397448 | 2.998597266 | 0.4985399196 | 6.014758594 | 1.80E-09 | 2.53E-07 | 1.449168615 |
| ENSG00000166 | 45.7322501 | -5.144504415 | 0.8553993402 | -6.014155229 | 1.81E-09 | 2.45E-07 | 1.516424702 |
| ENSG00000136 | 43.76621554 | 5.793888538 | 0.9661234775 | 5.997047658 | 2.01E-09 | 2.74E-07 | 1.470104381 |
| ENSG00000115 | 325.964405 | 2.008776729 | 0.3361072019 | 5.976595315 | 2.28E-09 | 3.38E-07 | 1.327567552 |
| ENSG00000168 | 31.03613702 | 6.211169617 | 1.040562421 | 5.969050481 | 2.39E-09 | 3.38E-07 | 1.393563518 |
| ENSG00000123 | 83.38787701 | -3.79175854 | 0.6362542303 | -5.959502286 | 2.53E-09 | 3.66E-07 | 1.337480197 |
| ENSG00000170 | 261.3453661 | 2.05274361 | 0.3450795824 | 5.948609291 | 2.70E-09 | 3.91E-07 | 1.327567552 |
| ENSG00000147 | 125.6968406 | 2.548163882 | 0.4288053321 | 5.942472472 | 2.81E-09 | 4.06E-07 | 1.318051683 |
| ENSG00000167 | 228.0987865 | 2.233585787 | 0.3760929932 | 5.938918905 | 2.87E-09 | 4.08E-07 | 1.327567552 |
| ENSG00000184 | 44.66954024 | 5.156818848 | 0.8719618016 | 5.91404215 | 3.34E-09 | 4.49E-07 | 1.393563518 |
| ENSG00000126 | 205.0051052 | 3.469549888 | 0.5874357778 | 5.90626247 | 3.50E-09 | 1.41E-06 | 0.3939394882 |
| ENSG00000125 | 426.8886131 | -3.108007717 | 0.5300194495 | -5.863950313 | 4.52E-09 | 6.71E-07 | 1.242301963 |
| ENSG00000106 | 150.15325 | 8.1514673 | 1.39566986 | 5.840541187 | 5.20E-09 | 7.49E-07 | 1.242301963 |
| ENSG00000076 | 445.8994277 | 1.935155268 | 0.3317892063 | 5.832484092 | 5.46E-09 | 6.46E-07 | 1.572476891 |
| ENSG00000239 | 201.7850598 | 2.532589713 | 0.4351912531 | 5.819486707 | 5.90E-09 | 2.33E-06 | 0.3706356727 |
| ENSG00000120 | 1204.765175 | -1.595341113 | 0.2750346847 | -5.800508815 | 6.61E-09 | 7.46E-07 | 1.597450288 |

| | | | | | | | |
|---|---|---|---|---|---|---|---|
| ENSG000002549 | 133.5918079 | 2.881069388 | 0.4973262498 | 5.793117475 | 6.91E-09 | 9.79E-07 | 1.242301963 |
| ENSG000001233 | 289.061069 | 2.320161765 | 0.4005171745 | 5.792914543 | 6.92E-09 | 9.79E-07 | 1.249985076 |
| ENSG000001837 | 670.2463327 | 1.646662283 | 0.2845270663 | 5.787366046 | 7.15E-09 | 7.46E-07 | 1.729500602 |
| ENSG000001731 | 98.40390356 | 2.954280217 | 0.5111061626 | 5.780169431 | 7.46E-09 | 1.12E-06 | 1.133466638 |
| ENSG000001318 | 109.5814003 | 2.657075829 | 0.4602127978 | 5.773580921 | 7.76E-09 | 1.14E-06 | 1.133466638 |
| ENSG000002764 | 253.630177 | 2.148792223 | 0.3724609946 | 5.769173832 | 7.97E-09 | 2.88E-06 | 0.3939394882 |
| ENSG000001310 | 339.7533379 | 1.928862592 | 0.3346782132 | 5.763334796 | 8.25E-09 | 1.12E-06 | 1.249985076 |
| ENSG000001976 | 270.0267624 | 2.312124965 | 0.4016602379 | 5.756419846 | 8.59E-09 | 1.17E-06 | 1.198705341 |
| ENSG000001098 | 124.8961024 | 2.76631715 | 0.4809783161 | 5.751438386 | 8.85E-09 | 1.17E-06 | 1.242301963 |
| ENSG000002218 | 515.1502318 | 2.538030072 | 0.4416230927 | 5.747050174 | 9.08E-09 | 9.91E-07 | 1.602205449 |
| ENSG000000222 | 525.487595 | 2.551667471 | 0.4454346008 | 5.728489584 | 1.01E-08 | 1.11E-06 | 1.572476891 |
| ENSG000001584 | 37.75554754 | 4.208219551 | 0.7352494749 | 5.723526088 | 1.04E-08 | 1.21E-06 | 1.393563518 |
| ENSG000002151 | 430.9087261 | -5.518782804 | 0.9648879272 | -5.719610172 | 1.07E-08 | 1.12E-06 | 1.602205449 |
| ENSG000001387 | 459.284821 | 2.122438773 | 0.3710976347 | 5.719354085 | 1.07E-08 | 1.02E-06 | 1.825593782 |
| ENSG000001642 | 1137.80158 | -1.79940961 | 0.3146728245 | -5.718350839 | 1.08E-08 | 1.13E-06 | 1.597450288 |
| ENSG000002134 | 80.09649356 | 4.337765634 | 0.7598634956 | 5.70861169 | 1.14E-08 | 1.21E-06 | 1.514054369 |
| ENSG000001651 | 141.246849 | 2.314953578 | 0.405564169 | 5.707983483 | 1.14E-08 | 1.87E-06 | 0.9247574895 |
| ENSG000001355 | 50.75455527 | 3.877229995 | 0.6813330731 | 5.690652851 | 1.27E-08 | 1.34E-06 | 1.516424702 |
| ENSG000000506 | 100.4615191 | 3.386956749 | 0.596920701 | 5.674048066 | 1.39E-08 | 1.79E-06 | 1.22159078 |
| ENSG000000642 | 179.8983765 | 7.39010907 | 1.30466664 | 5.664365781 | 1.48E-08 | 1.85E-06 | 1.22159078 |
| ENSG000001963 | 28.81058251 | 6.60213376 | 1.167838123 | 5.653295288 | 1.57E-08 | 1.81E-06 | 1.357988055 |
| ENSG000001868 | 50.15448955 | 3.253766138 | 0.5761233952 | 5.647689653 | 1.63E-08 | 1.81E-06 | 1.393563518 |
| ENSG000001004 | 41.43241285 | 4.463741772 | 0.7907090133 | 5.645239521 | 1.65E-08 | 1.87E-06 | 1.337480197 |
| ENSG000001201 | 688.0173364 | 2.102559073 | 0.3726684994 | 5.641901788 | 1.68E-08 | 1.54E-06 | 1.729500602 |
| ENSG000001660 | 201.0332283 | 2.044846983 | 0.3628305918 | 5.635817456 | 1.74E-08 | 2.15E-06 | 1.198705341 |
| ENSG000001199 | 43.73258897 | 3.629904626 | 0.6455547931 | 5.622922585 | 1.88E-08 | 2.09E-06 | 1.337480197 |
| ENSG000001637 | 726.4943744 | -2.361150255 | 0.4200977822 | -5.620477791 | 1.90E-08 | 1.83E-06 | 1.602205449 |
| ENSG000001562 | 43.90819985 | 3.496504461 | 0.6230949192 | 5.611511751 | 2.01E-08 | 1.98E-06 | 1.516424702 |
| ENSG000000872 | 136.090642 | 2.330972455 | 0.4167214762 | 5.593598093 | 2.22E-08 | 2.45E-06 | 1.318051683 |
| ENSG000001525 | 426.1973386 | 1.823302339 | 0.3263762114 | 5.586505007 | 2.32E-08 | 2.66E-06 | 1.249985076 |
| ENSG000001154 | 2611.345754 | 1.730895579 | 0.3101595382 | 5.580662097 | 2.40E-08 | 2.20E-06 | 1.602205449 |
| ENSG000001815 | 231.5719666 | 2.507131446 | 0.449753565 | 5.57445597 | 2.48E-08 | 7.75E-06 | 0.3939394882 |
| ENSG000001545 | 107.187522 | 2.822336893 | 0.5071821312 | 5.564740395 | 2.63E-08 | 3.01E-06 | 1.22159078 |

| | | | | | | | |
|---|---|---|---|---|---|---|---|
| ENSG000001088 | 132.6048946 | 2.94461824 | 0.5298574821 | 5.55737786 | 2.74E-08 | 2.95E-06 | 1.318051683 |
| ENSG000001757 | 86.44542301 | 4.317785913 | 0.7779671129 | 5.550087968 | 2.86E-08 | 3.01E-06 | 1.337480197 |
| ENSG000001341 | 148.7989171 | 3.062141327 | 0.5520603362 | 5.546751191 | 2.91E-08 | 4.28E-06 | 0.9247574895 |
| ENSG000001059 | 736.5360979 | 1.611382307 | 0.2911307804 | 5.534908761 | 3.11E-08 | 2.46E-06 | 1.825593782 |
| ENSG000001643 | 176.8343259 | 6.081048901 | 1.100065819 | 5.527895508 | 3.24E-08 | 3.74E-06 | 1.198705341 |
| ENSG000001989 | 180.2798865 | 2.24377188 | 0.4077092709 | 5.503362419 | 3.73E-08 | 5.29E-06 | 0.9247574895 |
| ENSG000001448 | 157.8624936 | 2.737698983 | 0.497653878 | 5.501210989 | 3.77E-08 | 4.22E-06 | 1.22159078 |
| ENSG000001879 | 578.3520051 | 1.919530132 | 0.3497080942 | 5.488949679 | 4.04E-08 | 3.58E-06 | 1.572476891 |
| ENSG000002879 | 74.63775206 | 2.944734083 | 0.5374134476 | 5.479457383 | 4.27E-08 | 4.21E-06 | 1.393563518 |
| ENSG000001285 | 131.3412295 | 2.785557654 | 0.5098832968 | 5.46312788 | 4.68E-08 | 4.74E-06 | 1.318051683 |
| ENSG000001870 | 125.9002602 | 2.29863445 | 0.4212089851 | 5.457230333 | 4.84E-08 | 5.56E-06 | 1.130447911 |
| ENSG000001527 | 72.6947627 | 3.424783058 | 0.6277994573 | 5.455218252 | 4.89E-08 | 4.37E-06 | 1.514054369 |
| ENSG000001636 | 367.8078336 | 1.77227207 | 0.3249698868 | 5.453650143 | 4.93E-08 | 5.25E-06 | 1.242301963 |
| ENSG000002744 | 182.5283329 | 2.480051886 | 0.4549624656 | 5.451113166 | 5.01E-08 | 6.88E-06 | 0.9247574895 |
| ENSG000001345 | 50.54111816 | 3.647498256 | 0.670031185 | 5.443773868 | 5.22E-08 | 4.97E-06 | 1.393563518 |
| ENSG000001333 | 565.7822358 | 1.95671074 | 0.3598036319 | 5.438274009 | 5.38E-08 | 4.51E-06 | 1.602205449 |
| ENSG000001099 | 269.1853507 | 3.09282212 | 0.569642195 | 5.429411913 | 5.65E-08 | 5.88E-06 | 1.242301963 |
| ENSG000001282 | 74.25288526 | 3.054011883 | 0.5642102501 | 5.412896845 | 6.20E-08 | 5.35E-06 | 1.514054369 |
| ENSG000002657 | 109.1867976 | 2.508903467 | 0.4644911896 | 5.401401627 | 6.61E-08 | 7.31E-06 | 1.130447911 |
| ENSG000000534 | 16.2879599 | 7.428405202 | 1.377344715 | 5.393279635 | 6.92E-08 | 6.51E-06 | 1.357988055 |
| ENSG000000349 | 16.35937197 | 7.411550906 | 1.374255619 | 5.393138513 | 6.92E-08 | 6.51E-06 | 1.357988055 |
| ENSG000001891 | 89.19650346 | 2.492999414 | 0.463259066 | 5.381436861 | 7.39E-08 | 6.98E-06 | 1.337480197 |
| ENSG000001692 | 90.79290473 | 3.903728474 | 0.7274325992 | 5.366446979 | 8.03E-08 | 7.24E-06 | 1.393563518 |
| ENSG000001541 | 110.1944264 | 2.582034319 | 0.4836016698 | 5.339175774 | 9.34E-08 | 9.25E-06 | 1.22159078 |
| ENSG000002620 | 492.5044379 | 10.2841866 | 1.927179466 | 5.336392786 | 9.48E-08 | 7.49E-06 | 1.564848717 |
| ENSG000001627 | 76.35167296 | 4.046776545 | 0.7596818046 | 5.326936252 | 9.99E-08 | 9.14E-06 | 1.337480197 |
| ENSG000001571 | 518.0248143 | 1.559436113 | 0.2929578374 | 5.323073541 | 1.02E-07 | 7.33E-06 | 1.729500602 |
| ENSG000001542 | 146.4590232 | 2.341174628 | 0.4413970931 | 5.304010073 | 1.13E-07 | 1.13E-05 | 1.198705341 |
| ENSG000002416 | 75.83217414 | 2.696907385 | 0.5092563745 | 5.295775409 | 1.19E-07 | 1.06E-05 | 1.337480197 |
| ENSG000002552 | 121.448406 | 2.512035876 | 0.4745622176 | 5.293375205 | 1.20E-07 | 1.17E-05 | 1.22159078 |
| ENSG000001370 | 70.69488919 | 3.209473837 | 0.6076226419 | 5.282018173 | 1.28E-07 | 1.06E-05 | 1.449168615 |
| ENSG000001601 | 107.2831058 | 3.322705024 | 0.6303539922 | 5.271173127 | 1.36E-07 | 1.39E-05 | 1.130447911 |
| ENSG000001492 | 180.4971689 | 2.515341187 | 0.4775200068 | 5.267509531 | 1.38E-07 | 9.25E-06 | 1.814765822 |

| | | | | | | | |
|---|---|---|---|---|---|---|---|
| ENSG00000197 | 288.8706652 | 1.90041363 | 0.3613193448 | 5.259650937 | 1.44E-07 | 1.29E-05 | 1.32213648 |
| ENSG00000229 | 76.40096639 | 3.195634632 | 0.6086502194 | 5.250363066 | 1.52E-07 | 1.33E-05 | 1.337480197 |
| ENSG00000236 | 211.6560112 | -2.692634879 | 0.5139053267 | -5.239554328 | 1.61E-07 | 1.54E-05 | 1.198705341 |
| ENSG00000116 | 134.2349103 | 2.021403226 | 0.3861414846 | 5.234877129 | 1.65E-07 | 1.45E-05 | 1.318051683 |
| ENSG00000171 | 296.0770609 | 2.025214982 | 0.3878857586 | 5.221163544 | 1.78E-07 | 1.54E-05 | 1.32213648 |
| ENSG00000156 | 812.1398883 | 2.148391774 | 0.4115228955 | 5.22058869 | 1.78E-07 | 1.31E-05 | 1.602205449 |
| ENSG00000205 | 65.22297481 | 3.464352262 | 0.6650171687 | 5.209417779 | 1.89E-07 | 1.59E-05 | 1.357988055 |
| ENSG00000154 | 61.59609977 | 3.421036585 | 0.6569550961 | 5.207413118 | 1.91E-07 | 1.60E-05 | 1.357988055 |
| ENSG00000213 | 195.9750072 | 2.484059031 | 0.4776043396 | 5.201081367 | 1.98E-07 | 1.68E-05 | 1.327567552 |
| ENSG00000276 | 31.03381653 | 4.706256494 | 0.9062772078 | 5.192954708 | 2.07E-07 | 1.76E-05 | 1.319539707 |
| ENSG00000101 | 145.0469935 | 2.463885093 | 0.4750060235 | 5.187060734 | 2.14E-07 | 1.91E-05 | 1.242301963 |
| ENSG00000203 | 25.77072155 | -6.445012008 | 1.25221947 | -5.146870947 | 2.65E-07 | 2.16E-05 | 1.357988055 |
| ENSG00000165 | 56.44712594 | 3.7783036 | 0.7352838348 | 5.138564758 | 2.77E-07 | 2.28E-05 | 1.337480197 |
| ENSG00000156 | 71.10504972 | 3.730081221 | 0.7283692948 | 5.121140125 | 3.04E-07 | 2.45E-05 | 1.357988055 |
| ENSG00000167 | 394.1046241 | 2.01289074 | 0.3934086118 | 5.116539597 | 3.11E-07 | 2.55E-05 | 1.327567552 |
| ENSG00000101 | 1035.862918 | -1.84970562 | 0.3617124713 | -5.113745771 | 3.16E-07 | 1.86E-05 | 1.902376403 |
| ENSG00000138 | 46.08544599 | 3.866879808 | 0.7581449356 | 5.100449302 | 3.39E-07 | 2.70E-05 | 1.357988055 |
| ENSG00000187 | 793.3912433 | 1.48727741 | 0.2921838226 | 5.09021135 | 3.58E-07 | 2.48E-05 | 1.572476891 |
| ENSG00000159 | 165.4174728 | 2.5843996 | 0.5082085037 | 5.085313569 | 3.67E-07 | 3.18E-05 | 1.242301963 |
| ENSG00000203 | 106.9211553 | 2.556372324 | 0.5029501344 | 5.082755027 | 3.72E-07 | 3.21E-05 | 1.242301963 |
| ENSG00000129 | 195.8760957 | 1.898053772 | 0.3744408907 | 5.069034442 | 4.00E-07 | 3.52E-05 | 1.198705341 |
| ENSG00000174 | 340.5550196 | 1.842652562 | 0.3636640628 | 5.066908584 | 4.04E-07 | 3.32E-05 | 1.29680518 |
| ENSG00000280 | 30.35354219 | 5.60184523 | 1.112133526 | 5.037025768 | 4.73E-07 | 3.65E-05 | 1.357988055 |
| ENSG00000091 | 26.2946929 | 4.895644985 | 0.9728645396 | 5.032195938 | 4.85E-07 | 3.71E-05 | 1.357988055 |
| ENSG00000185 | 140.6451546 | 2.212753257 | 0.4402183589 | 5.026490179 | 5.00E-07 | 5.30E-05 | 0.9247574895 |
| ENSG00000267 | 32.95510913 | 5.372462179 | 1.069732235 | 5.022249497 | 5.11E-07 | 4.00E-05 | 1.319539707 |
| ENSG00000004 | 679.2691228 | 1.789620026 | 0.3565505901 | 5.01925975 | 5.19E-07 | 3.52E-05 | 1.564848717 |
| ENSG00000127 | 22.61103047 | 4.70677585 | 0.9378641236 | 5.018611686 | 5.20E-07 | 4.06E-05 | 1.319539707 |
| ENSG00000104 | 148.5878822 | 2.563924037 | 0.5115068039 | 5.012492537 | 5.37E-07 | 4.45E-05 | 1.22159078 |
| ENSG00000280 | 49.82314288 | -4.96980481 | 0.9916670374 | -5.011566002 | 5.40E-07 | 4.13E-05 | 1.337480197 |
| ENSG00000007 | 872.7088866 | -2.431469994 | 0.4853071673 | -5.010167081 | 5.44E-07 | 3.64E-05 | 1.572476891 |
| ENSG00000086 | 3572.343921 | -2.251947413 | 0.4503061869 | -5.000924878 | 5.71E-07 | 3.71E-05 | 1.597450288 |
| ENSG00000173 | 147.1053832 | 3.213541357 | 0.6433341037 | 4.995136025 | 5.88E-07 | 6.13E-05 | 0.9247574895 |

| | | | | | | | |
|---|---|---|---|---|---|---|---|
| ENSG00000149 | 127.1799574 | 2.099964556 | 0.4212177965 | 4.98546019 | 6.18E-07 | 4.96E-05 | 1.242301963 |
| ENSG00000168 | 92.6682176 | 2.446274312 | 0.4912217883 | 4.979979248 | 6.36E-07 | 4.45E-05 | 1.449168615 |
| ENSG00000196 | 84.16977715 | 2.350266681 | 0.4724695934 | 4.974429496 | 6.54E-07 | 4.54E-05 | 1.449168615 |
| ENSG00000101 | 45.94399938 | 3.415198429 | 0.6867478553 | 4.973001958 | 6.59E-07 | 4.74E-05 | 1.393563518 |
| ENSG00000072 | 344.1114573 | 1.762334371 | 0.3546450853 | 4.969290268 | 6.72E-07 | 5.10E-05 | 1.29680518 |
| ENSG00000253 | 459.408333 | 1.892622037 | 0.3809892775 | 4.967651714 | 6.78E-07 | 4.31E-05 | 1.602205449 |
| ENSG00000071 | 36.14241682 | 4.371802703 | 0.8812401031 | 4.960966582 | 7.01E-07 | 5.00E-05 | 1.393563518 |
| ENSG00000091 | 85.74186898 | 2.915460293 | 0.5899235121 | 4.942098821 | 7.73E-07 | 5.05E-05 | 1.514054369 |
| ENSG00000166 | 326.4843029 | 2.159902083 | 0.4375025643 | 4.936890111 | 7.94E-07 | 5.82E-05 | 1.32213648 |
| ENSG00000101 | 413.0642903 | -2.116254535 | 0.4293982383 | -4.92841923 | 8.29E-07 | 6.32E-05 | 1.249985076 |
| ENSG00000145 | 168.5630865 | 1.969464429 | 0.3996944536 | 4.927424965 | 8.33E-07 | 6.34E-05 | 1.242301963 |
| ENSG00000079 | 303.6323731 | 1.595768822 | 0.3246231463 | 4.915757979 | 8.84E-07 | 6.32E-05 | 1.327567552 |
| ENSG00000145 | 1797.317805 | 1.277958061 | 0.2603006682 | 4.909545836 | 9.13E-07 | 5.45E-05 | 1.634803475 |
| ENSG00000227 | 95.67984878 | 2.417201795 | 0.4929102731 | 4.903938764 | 9.39E-07 | 7.76E-05 | 1.133466638 |
| ENSG00000206 | 203.3339647 | -2.476095287 | 0.5067810504 | -4.885927138 | 1.03E-06 | 7.99E-05 | 1.198705341 |
| ENSG00000182 | 388.1226707 | 2.376491565 | 0.4869690377 | 4.880169746 | 1.06E-06 | 7.51E-05 | 1.327567552 |
| ENSG00000269 | 97.6186532 | 2.494453057 | 0.5111563248 | 4.88001994 | 1.06E-06 | 8.55E-05 | 1.130447911 |
| ENSG00000152 | 217.9750161 | -1.894848405 | 0.3889127142 | -4.872168833 | 1.10E-06 | 7.76E-05 | 1.327567552 |
| ENSG00000174 | 899.451637 | 1.514584797 | 0.3108681511 | 4.872113119 | 1.10E-06 | 5.65E-05 | 1.902376403 |
| ENSG00000196 | 190.7157476 | 1.805274262 | 0.3714262785 | 4.860383788 | 1.17E-06 | 6.18E-05 | 1.814765822 |
| ENSG00000100 | 156.1434363 | 2.565243952 | 0.5277964317 | 4.860290442 | 1.17E-06 | 6.18E-05 | 1.814765822 |
| ENSG00000064 | 66.47328545 | 2.845448696 | 0.5858646632 | 4.856836186 | 1.19E-06 | 8.23E-05 | 1.337480197 |
| ENSG00000165 | 133.8007411 | 2.145293371 | 0.4426558844 | 4.846413314 | 1.26E-06 | 8.66E-05 | 1.318051683 |
| ENSG00000146 | 92.82742416 | 3.44319026 | 0.7106243275 | 4.845303104 | 1.26E-06 | 8.34E-05 | 1.393563518 |
| ENSG00000170 | 29.45528757 | 4.706773086 | 0.9716149884 | 4.844277973 | 1.27E-06 | 8.71E-05 | 1.319539707 |
| ENSG00000162 | 143.154265 | -8.86226766 | 1.82983619 | -4.843202747 | 1.28E-06 | 9.48E-05 | 1.198705341 |
| ENSG00000013 | 46.97317336 | 4.428958347 | 0.9167667833 | 4.831063284 | 1.36E-06 | 8.45E-05 | 1.470104381 |
| ENSG00000123 | 370.4232555 | 1.62322194 | 0.3364681704 | 4.824295677 | 1.40E-06 | 9.81E-05 | 1.242301963 |
| ENSG00000268 | 28.90389838 | -4.867948377 | 1.009067386 | -4.824205446 | 1.41E-06 | 9.29E-05 | 1.357988055 |
| ENSG00000167 | 104.9106687 | 2.272987872 | 0.4712080047 | 4.823746306 | 1.41E-06 | 9.48E-05 | 1.318051683 |
| ENSG00000102 | 5043.788353 | -1.856255683 | 0.3848581763 | -4.823220077 | 1.41E-06 | 8.00E-05 | 1.634803475 |
| ENSG00000140 | 21.93925128 | -5.428264811 | 1.126400733 | -4.819124004 | 1.44E-06 | 9.53E-05 | 1.337480197 |
| ENSG00000224 | 23.14309195 | 5.647731911 | 1.172947749 | 4.814990197 | 1.47E-06 | 9.71E-05 | 1.319539707 |

| | | | | | | | |
|---|---|---|---|---|---|---|---|
| ENSG000001842 | 37.08459904 | 3.198001675 | 0.6646411594 | 4.811621474 | 1.50E-06 | 9.67E-05 | 1.357988055 |
| ENSG000001184 | 133.5660265 | 2.480753306 | 0.5165727663 | 4.802330801 | 1.57E-06 | 1.09E-04 | 1.242301963 |
| ENSG000001328 | 62.14036456 | 2.750133818 | 0.573367047 | 4.796462986 | 1.61E-06 | 9.48E-05 | 1.516424702 |
| ENSG000001664 | 18.40319931 | 5.822656716 | 1.215054939 | 4.792093368 | 1.65E-06 | 9.26E-05 | 1.605577974 |
| ENSG000001684 | 113.5040928 | 2.491123358 | 0.520481029 | 4.786194346 | 1.70E-06 | 1.26E-04 | 1.133466638 |
| ENSG000001486 | 926.0083928 | 1.476358493 | 0.3085434978 | 4.78492823 | 1.71E-06 | 9.58E-05 | 1.572476891 |
| ENSG000002336 | 102.1458292 | 2.81650767 | 0.5891706767 | 4.780461387 | 1.75E-06 | 1.22E-04 | 1.22159078 |
| ENSG000001114 | 19.61823194 | 6.462533064 | 1.352269969 | 4.779025795 | 1.76E-06 | 1.13E-04 | 1.337480197 |
| ENSG000001410 | 150.1975366 | 1.82525832 | 0.3819441394 | 4.778861965 | 1.76E-06 | 1.24E-04 | 1.198705341 |
| ENSG000000029 | 2119.253884 | -1.17992726 | 0.2469598396 | -4.777810279 | 1.77E-06 | 9.69E-05 | 1.597450288 |
| ENSG000000184 | 295.456082 | 1.516290742 | 0.3176406791 | 4.773603767 | 1.81E-06 | 1.23E-04 | 1.249985076 |
| ENSG000001058 | 115.7022383 | -2.843863282 | 0.5958564451 | -4.772732266 | 1.82E-06 | 1.33E-04 | 1.133466638 |
| ENSG000001829 | 350.2655652 | 1.692055158 | 0.3550190788 | 4.766096413 | 1.88E-06 | 1.21E-04 | 1.32213648 |
| ENSG000001097 | 78.96749319 | 2.74784407 | 0.5771519554 | 4.761040908 | 1.93E-06 | 1.19E-04 | 1.393563518 |
| ENSG000000737 | 242.3545035 | -2.551427305 | 0.5360515989 | -4.759667373 | 1.94E-06 | 3.51E-04 | 0.3939394882 |
| ENSG000001123 | 148.7930076 | 2.029662386 | 0.4265399261 | 4.758434702 | 1.95E-06 | 1.33E-04 | 1.22159078 |
| ENSG000000737 | 388.1549864 | 1.97400881 | 0.4157762587 | 4.747767023 | 2.06E-06 | 1.36E-04 | 1.242301963 |
| ENSG000002533 | 266.6100048 | 1.932348118 | 0.4071364268 | 4.746193145 | 2.07E-06 | 1.37E-04 | 1.242301963 |
| ENSG000001608 | 50.09775124 | 3.612310014 | 0.7611974059 | 4.745562696 | 2.08E-06 | 1.26E-04 | 1.393563518 |
| ENSG000001161 | 36.56464033 | 4.164741794 | 0.8791426048 | 4.737276718 | 2.17E-06 | 1.33E-04 | 1.357988055 |
| ENSG000001176 | 171.3494559 | 1.872689101 | 0.3963730427 | 4.724562214 | 2.31E-06 | 1.54E-04 | 1.198705341 |
| ENSG000002541 | 54.61228858 | 3.865137903 | 0.8181764647 | 4.724088347 | 2.31E-06 | 1.41E-04 | 1.337480197 |
| ENSG000002803 | 38.55977125 | 3.414281532 | 0.7232572522 | 4.720701413 | 2.35E-06 | 1.42E-04 | 1.337480197 |
| ENSG000002651 | 309.0196598 | 6.574247687 | 1.393127606 | 4.719056357 | 2.37E-06 | 1.45E-04 | 1.32213648 |
| ENSG000001230 | 529.0432831 | 1.696544217 | 0.3598258486 | 4.714903678 | 2.42E-06 | 1.29E-04 | 1.564848717 |
| ENSG000001649 | 53.24122002 | 2.939692212 | 0.6240869154 | 4.710389112 | 2.47E-06 | 1.38E-04 | 1.470104381 |
| ENSG000001808 | 59.25589584 | 3.810786686 | 0.8101455761 | 4.703829532 | 2.55E-06 | 1.41E-04 | 1.470104381 |
| ENSG000001422 | 130.3156501 | 1.972328654 | 0.4204181458 | 4.691349965 | 2.71E-06 | 1.85E-04 | 1.133466638 |
| ENSG000002121 | 18.39400918 | 4.862567301 | 1.037260378 | 4.687894577 | 2.76E-06 | 1.47E-04 | 1.514850267 |
| ENSG000001607 | 406.4656397 | -1.45634348 | 0.3107322649 | -4.686811266 | 2.77E-06 | 1.69E-04 | 1.29680518 |
| ENSG000002803 | 166.1676637 | 5.594881194 | 1.194204584 | 4.685027399 | 2.80E-06 | 1.79E-04 | 1.22159078 |
| ENSG000001799 | 45.1939385 | 3.888766525 | 0.8306788421 | 4.681432014 | 2.85E-06 | 1.66E-04 | 1.357988055 |
| ENSG000001824 | 258.5663897 | 2.046351009 | 0.4379232027 | 4.672853588 | 2.97E-06 | 5.08E-04 | 0.3939394882 |

| | | | | | | | |
|---|---|---|---|---|---|---|---|
| ENSG0000023 | 56.98500706 | 2.847997345 | 0.6103968171 | 4.665812903 | 3.07E-06 | 1.77E-04 | 1.357988055 |
| ENSG0000014 | 486.0090523 | 1.369491607 | 0.2935979835 | 4.664512987 | 3.09E-06 | 1.58E-04 | 1.564848717 |
| ENSG0000010 | 50.71663565 | 3.209922159 | 0.6884682299 | 4.662411451 | 3.13E-06 | 1.64E-04 | 1.516424702 |
| ENSG0000013 | 145.9952666 | -2.150801689 | 0.4625816066 | -4.649561631 | 3.33E-06 | 2.68E-04 | 0.9247574895 |
| ENSG0000014 | 49.85035788 | 3.509332831 | 0.7549920001 | 4.648172206 | 3.35E-06 | 1.85E-04 | 1.393563518 |
| ENSG0000014 | 130.8406578 | 1.982807397 | 0.4283438119 | 4.629009086 | 3.67E-06 | 2.26E-04 | 1.22159078 |
| ENSG0000002 | 691.9940995 | 1.47554715 | 0.3189875476 | 4.625720223 | 3.73E-06 | 1.70E-04 | 1.729500602 |
| ENSG0000006 | 280.5596767 | 1.910240378 | 0.4132239874 | 4.62277224 | 3.79E-06 | 2.21E-04 | 1.29680518 |
| ENSG0000016 | 622.9925288 | -4.686275608 | 1.014081517 | -4.621202074 | 3.82E-06 | 1.66E-04 | 1.825593782 |
| ENSG0000011 | 312.4468455 | 5.214748934 | 1.128999639 | 4.618911071 | 3.86E-06 | 2.21E-04 | 1.32213648 |
| ENSG0000002 | 936.9494751 | 1.788631239 | 0.3872528055 | 4.618768963 | 3.86E-06 | 1.84E-04 | 1.634803475 |
| ENSG0000014 | 3606.249254 | 1.342944073 | 0.291688202 | 4.604039738 | 4.14E-06 | 1.98E-04 | 1.602205449 |
| ENSG0000016 | 2578.668797 | -1.919755456 | 0.4176983213 | -4.596033449 | 4.31E-06 | 2.06E-04 | 1.597450288 |
| ENSG0000007 | 147.2626404 | -1.69881292 | 0.3698242794 | -4.593567851 | 4.36E-06 | 1.85E-04 | 1.814765822 |
| ENSG0000017 | 112.7717327 | 2.681386915 | 0.5839560235 | 4.59176172 | 4.40E-06 | 2.83E-04 | 1.133466638 |
| ENSG0000012 | 430.7289028 | 1.53091279 | 0.3337860891 | 4.586508664 | 4.51E-06 | 2.14E-04 | 1.602205449 |
| ENSG0000014 | 818.5693194 | 1.237852685 | 0.2707612472 | 4.571749827 | 4.84E-06 | 2.27E-04 | 1.597450288 |
| ENSG0000011 | 97.64644398 | 2.389128629 | 0.5232501283 | 4.565939882 | 4.97E-06 | 2.79E-04 | 1.318051683 |
| ENSG0000011 | 342.9785394 | 2.08984024 | 0.4578464624 | 4.564500136 | 5.01E-06 | 2.92E-04 | 1.249985076 |
| ENSG0000021 | 285.0822611 | 1.638966176 | 0.3591331031 | 4.56367336 | 5.03E-06 | 2.79E-04 | 1.327567552 |
| ENSG0000025 | 380.9130344 | -1.829261444 | 0.4010716223 | -4.560934612 | 5.09E-06 | 2.98E-04 | 1.242301963 |
| ENSG0000015 | 78.46574213 | 2.28845684 | 0.5022219004 | 4.55666477 | 5.20E-06 | 2.81E-04 | 1.357988055 |
| ENSG0000011 | 786.2769936 | -2.11605562 | 0.4662375948 | -4.538577849 | 5.66E-06 | 2.25E-04 | 1.902376403 |
| ENSG0000007 | 659.3576296 | 1.445147635 | 0.318765367 | 4.533577939 | 5.80E-06 | 2.51E-04 | 1.729500602 |
| ENSG0000024 | 409.1604167 | -2.150563099 | 0.4744142484 | -4.533091292 | 5.81E-06 | 3.16E-04 | 1.327567552 |
| ENSG0000010 | 22.85900873 | 4.508146826 | 0.9946893738 | 4.53221573 | 5.84E-06 | 3.29E-04 | 1.278627818 |
| ENSG0000014 | 273.5511308 | 1.44786235 | 0.3197518384 | 4.528081393 | 5.95E-06 | 3.53E-04 | 1.198705341 |
| ENSG0000010 | 14.44573334 | 6.402551574 | 1.415455169 | 4.523316397 | 6.09E-06 | 2.79E-04 | 1.605577974 |
| ENSG0000010 | 568.6072355 | -1.5826627 | 0.3504510509 | -4.516073488 | 6.30E-06 | 2.71E-04 | 1.729500602 |
| ENSG0000013 | 221.3508885 | 1.946344523 | 0.4316313372 | 4.509275288 | 6.50E-06 | 9.92E-04 | 0.3939394882 |
| ENSG0000016 | 121.2468689 | -3.003099748 | 0.6660933309 | -4.508526971 | 6.53E-06 | 3.71E-04 | 1.22159078 |
| ENSG0000014 | 301.9646406 | 4.645126136 | 1.030442715 | 4.507893614 | 6.55E-06 | 3.56E-04 | 1.29680518 |
| ENSG0000006 | 55.99784136 | 2.855871313 | 0.6341430313 | 4.503512885 | 6.68E-06 | 3.54E-04 | 1.337480197 |

| | | | | | | | |
|---|---|---|---|---|---|---|---|
| ENSG00000271 | 143.9128676 | 1.690050808 | 0.3754626782 | 4.501248475 | 6.76E-06 | 4.95E-04 | 0.9247574895 |
| ENSG00000189 | 15.85201188 | 6.454218728 | 1.434038873 | 4.500727875 | 6.77E-06 | 3.05E-04 | 1.605577974 |
| ENSG00000173 | 43.63947636 | 2.599690594 | 0.5781562273 | 4.49651923 | 6.91E-06 | 3.36E-04 | 1.470104381 |
| ENSG00000117 | 241.3191175 | -1.687118937 | 0.3753555617 | -4.494722096 | 6.97E-06 | 3.66E-04 | 1.327567552 |
| ENSG00000180 | 79.2517255 | 2.076807661 | 0.4622020549 | 4.493289545 | 7.01E-06 | 3.32E-04 | 1.514054369 |
| ENSG00000151 | 110.0207979 | 2.960640169 | 0.6589769503 | 4.492782589 | 7.03E-06 | 4.28E-04 | 1.133466638 |
| ENSG00000197 | 55.44654589 | 2.716741887 | 0.6063003733 | 4.480851416 | 7.43E-06 | 3.56E-04 | 1.470104381 |
| ENSG00000152 | 23.93885311 | 4.308598511 | 0.9617122389 | 4.480132764 | 7.46E-06 | 3.71E-04 | 1.393563518 |
| ENSG00000124 | 45.39532118 | -4.436223286 | 0.9919406811 | -4.47226671 | 7.74E-06 | 3.58E-04 | 1.516424702 |
| ENSG00000144 | 221.5068164 | 1.725404927 | 0.3858416054 | 4.471795947 | 7.76E-06 | 4.29E-04 | 1.242301963 |
| ENSG00000172 | 82.51241102 | 2.157177636 | 0.4826733173 | 4.469229101 | 7.85E-06 | 3.90E-04 | 1.393563518 |
| ENSG00000172 | 252.7603044 | -8.760543166 | 1.962172133 | -4.464716943 | 8.02E-06 | 4.43E-04 | 1.242301963 |
| ENSG00000121 | 243.8825428 | 1.96373876 | 0.4399660252 | 4.463387278 | 8.07E-06 | 0.001175432083 | 0.3939394882 |
| ENSG00000108 | 110.6558184 | 2.216345588 | 0.4988986176 | 4.442476907 | 8.89E-06 | 4.60E-04 | 1.318051683 |
| ENSG00000144 | 36.20771365 | 3.775025214 | 0.852318354 | 4.429125803 | 9.46E-06 | 4.81E-04 | 1.337480197 |
| ENSG00000167 | 152.8468062 | 2.109629473 | 0.4763889176 | 4.428376469 | 9.49E-06 | 6.63E-04 | 0.9247574895 |
| ENSG00000172 | 1122.679716 | 1.723138689 | 0.3898175301 | 4.420372496 | 9.85E-06 | 3.63E-04 | 1.902376403 |
| ENSG00000122 | 37.27771859 | 3.510235364 | 0.794950064 | 4.415667754 | 1.01E-05 | 5.13E-04 | 1.319539707 |
| ENSG00000142 | 95.97069529 | 2.957920537 | 0.6699834961 | 4.414915523 | 1.01E-05 | 5.45E-04 | 1.242301963 |
| ENSG00000152 | 393.5055934 | 4.669333382 | 1.058449911 | 4.411482615 | 1.03E-05 | 5.51E-04 | 1.242301963 |
| ENSG00000160 | 2507.329822 | -1.729927485 | 0.3924008527 | -4.408572185 | 1.04E-05 | 4.45E-04 | 1.597450288 |
| ENSG00000055 | 149.8390991 | 2.177852541 | 0.4947679538 | 4.401765564 | 1.07E-05 | 5.87E-04 | 1.198705341 |
| ENSG00000103 | 375.5555001 | 1.988153735 | 0.4521299191 | 4.397306286 | 1.10E-05 | 5.60E-04 | 1.29680518 |
| ENSG00000017 | 397.2680901 | 1.9251106 | 0.4385550577 | 4.389666853 | 1.14E-05 | 5.65E-04 | 1.327567552 |
| ENSG00000163 | 1025.76772 | -1.725339538 | 0.393802526 | -4.381230246 | 1.18E-05 | 5.07E-04 | 1.572476891 |
| ENSG00000169 | 55.56745486 | -8.052164708 | 1.842347914 | -4.370599411 | 1.24E-05 | 5.94E-04 | 1.357988055 |
| ENSG00000090 | 163.9077371 | 1.583538323 | 0.3625047838 | 4.368323934 | 1.25E-05 | 8.46E-04 | 0.9247574895 |
| ENSG00000106 | 60.66120069 | 2.556684289 | 0.5854248938 | 4.367228514 | 1.26E-05 | 5.90E-04 | 1.393563518 |
| ENSG00000175 | 195.3349684 | 2.410927382 | 0.5530634722 | 4.359223676 | 1.31E-05 | 6.38E-04 | 1.327567552 |
| ENSG00000064 | 472.2309524 | -1.453699976 | 0.3337255949 | -4.355973886 | 1.32E-05 | 5.51E-04 | 1.602205449 |
| ENSG00000227 | 16.5975318 | 5.623419204 | 1.291835825 | 4.353044788 | 1.34E-05 | 5.55E-04 | 1.605577974 |
| ENSG00000240 | 503.062512 | 1.696214667 | 0.3902010732 | 4.34702717 | 1.38E-05 | 5.78E-04 | 1.572476891 |
| ENSG00000173 | 220.4097233 | 1.848321729 | 0.4255276288 | 4.343599814 | 1.40E-05 | 7.47E-04 | 1.198705341 |

| | | | | | | | |
|---|---|---|---|---|---|---|---|
| ENSG00000129 | 2607.954398 | 1.234231517 | 0.284378093 | 4.340107581 | 1.42E-05 | 5.84E-04 | 1.602205449 |
| ENSG00000163 | 290.3990869 | 2.100635242 | 0.4846142158 | 4.334654604 | 1.46E-05 | 7.11E-04 | 1.32213648 |
| ENSG00000145 | 767.594771 | -1.488653027 | 0.3447571802 | -4.317975411 | 1.57E-05 | 6.48E-04 | 1.572476891 |
| ENSG00000175 | 285.2769813 | -2.102146983 | 0.4885205652 | -4.303088003 | 1.68E-05 | 8.16E-04 | 1.29680518 |
| ENSG00000137 | 33.93284817 | 4.256698653 | 0.989535632 | 4.301713365 | 1.69E-05 | 8.09E-04 | 1.319539707 |
| ENSG00000134 | 205.0314016 | -1.829182318 | 0.4255799555 | -4.298093212 | 1.72E-05 | 0.002149198876 | 0.3939394882 |
| ENSG00000150 | 163.1495476 | 1.55858791 | 0.3627122244 | 4.29703717 | 1.73E-05 | 8.88E-04 | 1.198705341 |
| ENSG00000113 | 331.4590661 | 1.680870676 | 0.3915988538 | 4.292327875 | 1.77E-05 | 8.34E-04 | 1.327567552 |
| ENSG00000116 | 1220.286209 | 1.792420827 | 0.418146118 | 4.286589662 | 1.81E-05 | 7.39E-04 | 1.572476891 |
| ENSG00000112 | 94.36631889 | 2.119380843 | 0.4946221659 | 4.284848091 | 1.83E-05 | 9.77E-04 | 1.133466638 |
| ENSG00000183 | 117.3619089 | 1.932760083 | 0.4511562064 | 4.284015281 | 1.84E-05 | 9.04E-04 | 1.242301963 |
| ENSG00000049 | 46.9574334 | 2.547936004 | 0.5949925247 | 4.282299186 | 1.85E-05 | 7.75E-04 | 1.516424702 |
| ENSG00000183 | 62.49698778 | 2.316585611 | 0.5419997007 | 4.274145554 | 1.92E-05 | 8.81E-04 | 1.357988055 |
| ENSG00000187 | 19.32702672 | 5.551117822 | 1.298995956 | 4.27339115 | 1.93E-05 | 8.88E-04 | 1.337480197 |
| ENSG00000163 | 41.59635695 | 3.447022259 | 0.8066624035 | 4.273190672 | 1.93E-05 | 8.05E-04 | 1.516424702 |
| ENSG00000163 | 1882.377335 | -1.252283818 | 0.2930806315 | -4.272830353 | 1.93E-05 | 7.69E-04 | 1.597450288 |
| ENSG00000088 | 106.2842789 | -2.504314352 | 0.586167199 | -4.272354981 | 1.93E-05 | 0.001017126065 | 1.133466638 |
| ENSG00000241 | 73.85228292 | 2.740656368 | 0.6415524745 | 4.271913019 | 1.94E-05 | 8.82E-04 | 1.357988055 |
| ENSG00000099 | 57.03067218 | 2.532816422 | 0.59290459 | 4.271878587 | 1.94E-05 | 8.89E-04 | 1.337480197 |
| ENSG00000154 | 216.7916422 | 1.863968673 | 0.4366529519 | 4.268764622 | 1.97E-05 | 0.002487946652 | 0.3706356727 |
| ENSG00000134 | 100.7479084 | 2.068623726 | 0.4850002558 | 4.26520131 | 2.00E-05 | 0.001044911282 | 1.133466638 |
| ENSG00000124 | 114.4302196 | 2.647843187 | 0.6209951547 | 4.263870928 | 2.01E-05 | 0.001051280566 | 1.130447911 |
| ENSG00000140 | 530.9785369 | 1.37322606 | 0.3223527184 | 4.260010796 | 2.04E-05 | 8.06E-04 | 1.602205449 |
| ENSG00000171 | 62.51410772 | -3.25563223 | 0.765095799 | -4.255195538 | 2.09E-05 | 8.81E-04 | 1.470104381 |
| ENSG00000008 | 1464.362853 | 1.19234623 | 0.2806062278 | 4.249179498 | 2.15E-05 | 7.24E-04 | 1.902376403 |
| ENSG00000050 | 23.30696733 | 5.943344038 | 1.398788788 | 4.248921702 | 2.15E-05 | 9.38E-04 | 1.393563518 |
| ENSG00000133 | 189.9552405 | 1.63451814 | 0.3847095164 | 4.248707326 | 2.15E-05 | 0.001302345152 | 0.9247574895 |
| ENSG00000232 | 39.43500236 | -3.093035741 | 0.7290244378 | -4.242705156 | 2.21E-05 | 0.001002708574 | 1.319539707 |
| ENSG00000133 | 138.0380731 | 1.851645015 | 0.4372584898 | 4.2346691 | 2.29E-05 | 0.001099895089 | 1.22159078 |
| ENSG00000198 | 65.64322296 | 2.316590747 | 0.5482898749 | 4.225120421 | 2.39E-05 | 9.92E-04 | 1.449168615 |
| ENSG00000179 | 88.64451177 | 2.51471076 | 0.5952256769 | 4.224802218 | 2.39E-05 | 0.001020273005 | 1.393563518 |
| ENSG00000121 | 196.0392367 | 1.73212287 | 0.4101500593 | 4.223144263 | 2.41E-05 | 0.001132677672 | 1.242301963 |
| ENSG00000196 | 28.16921941 | -4.385970958 | 1.038616792 | -4.222896252 | 2.41E-05 | 0.001078390331 | 1.319539707 |

| | | | | | | | |
|---|---|---|---|---|---|---|---|
| ENSG0000014[illegible] | 7609.216884 | 1.441887653 | 0.3421636038 | 4.214029887 | 2.51E-05 | 9.68E-04 | 1.572476891 |
| ENSG0000012[illegible] | 156.6236941 | 1.883300393 | 0.4471375324 | 4.211904071 | 2.53E-05 | 0.001172869941 | 1.242301963 |
| ENSG0000019[illegible] | 484.7954198 | 1.688834419 | 0.4014551511 | 4.206782287 | 2.59E-05 | 8.81E-04 | 1.825593782 |
| ENSG0000006[illegible] | 131.3692904 | 1.784710416 | 0.4243573393 | 4.205678213 | 2.60E-05 | 0.001289478036 | 1.133466638 |
| ENSG0000017[illegible] | 33.98711799 | 4.926970226 | 1.171544943 | 4.205532408 | 2.60E-05 | 0.001134600019 | 1.337480197 |
| ENSG0000014[illegible] | 928.1271204 | 1.170817232 | 0.2786652273 | 4.201518947 | 2.65E-05 | 9.79E-04 | 1.634803475 |
| ENSG0000024[illegible] | 28.61839862 | 4.814350068 | 1.145980993 | 4.201073225 | 2.66E-05 | 0.001151411713 | 1.337480197 |
| ENSG0000022[illegible] | 198.554117 | -1.615798902 | 0.3848117579 | -4.198933294 | 2.68E-05 | 0.003019994869 | 0.3939394882 |
| ENSG0000026[illegible] | 140.5428206 | 2.492137653 | 0.593676477 | 4.197804275 | 2.70E-05 | 9.06E-04 | 1.814765822 |
| ENSG0000014[illegible] | 37.44757284 | 3.023321824 | 0.7208914071 | 4.193865808 | 2.74E-05 | 0.001219023221 | 1.278627818 |
| ENSG0000016[illegible] | 36.48042521 | 3.785223008 | 0.9027447805 | 4.193015667 | 2.75E-05 | 0.001169248415 | 1.357988055 |
| ENSG0000016[illegible] | 226.6156638 | 1.69699792 | 0.4048528059 | 4.191641741 | 2.77E-05 | 0.001257808032 | 1.242301963 |
| ENSG0000018[illegible] | 87.50719279 | 3.522199671 | 0.8414176682 | 4.186030082 | 2.84E-05 | 0.001209285911 | 1.337480197 |
| ENSG0000011[illegible] | 97.03917754 | 2.288243421 | 0.5475498327 | 4.179059666 | 2.93E-05 | 0.001256092873 | 1.318051683 |
| ENSG0000018[illegible] | 608.5495348 | 1.389598643 | 0.3326524386 | 4.177328894 | 2.95E-05 | 0.001083390071 | 1.602205449 |
| ENSG0000009[illegible] | 158.931096 | 1.593220351 | 0.3815453542 | 4.17570371 | 2.97E-05 | 0.001720749514 | 0.9247574895 |
| ENSG0000017[illegible] | 224.0790122 | 1.429547593 | 0.342827329 | 4.169876414 | 3.05E-05 | 0.001361043267 | 1.242301963 |
| ENSG0000015[illegible] | 78.41383732 | 1.886980411 | 0.4526839319 | 4.168428076 | 3.07E-05 | 0.00124784895 | 1.393563518 |
| ENSG0000025[illegible] | 13.24266106 | 5.246415501 | 1.259180821 | 4.166530663 | 3.09E-05 | 0.001281859602 | 1.357988055 |
| ENSG0000017[illegible] | 757.1461052 | 1.373313217 | 0.3296123823 | 4.166449111 | 3.09E-05 | 0.001130662643 | 1.602205449 |
| ENSG0000013[illegible] | 449.0012119 | -1.304560163 | 0.3131519967 | -4.16590083 | 3.10E-05 | 0.001015154085 | 1.825593782 |
| ENSG0000018[illegible] | 22.50478 | 5.042504855 | 1.211649567 | 4.161685848 | 3.16E-05 | 0.001191085725 | 1.514850267 |
| ENSG0000017[illegible] | 771.4152734 | 1.712958536 | 0.4120565465 | 4.157095792 | 3.22E-05 | 0.001163294572 | 1.602205449 |
| ENSG0000016[illegible] | 97.70391437 | 3.298067063 | 0.793864424 | 4.154446229 | 3.26E-05 | 0.00145999971 | 1.22159078 |
| ENSG0000023[illegible] | 85.57649569 | 2.079697483 | 0.5017563092 | 4.144835739 | 3.40E-05 | 0.001385974384 | 1.357988055 |
| ENSG0000012[illegible] | 14.03327465 | 5.627547018 | 1.357957229 | 4.144126852 | 3.41E-05 | 0.001386950376 | 1.357988055 |
| ENSG0000014[illegible] | 62.96726622 | 2.684227352 | 0.648006341 | 4.142285626 | 3.44E-05 | 0.0013070451 | 1.470104381 |
| ENSG0000013[illegible] | 170.1828301 | 1.608400778 | 0.3882895117 | 4.142272016 | 3.44E-05 | 0.001536110685 | 1.22159078 |
| ENSG0000027[illegible] | 63.03616568 | 2.773679663 | 0.670895117 | 4.13429699 | 3.56E-05 | 0.001404052422 | 1.393563518 |
| ENSG0000028[illegible] | 47.99321001 | 3.3569943 | 0.8123929583 | 4.132229687 | 3.59E-05 | 0.001320692108 | 1.516424702 |
| ENSG0000014[illegible] | 158.3813848 | 1.791617584 | 0.4347069125 | 4.121437991 | 3.77E-05 | 0.001187910398 | 1.814765822 |
| ENSG0000019[illegible] | 14709.48377 | -1.686704509 | 0.4092704988 | -4.12124625 | 3.77E-05 | 0.001151159195 | 1.902376403 |
| ENSG0000017[illegible] | 154.1842143 | 1.544233917 | 0.3747700612 | 4.120483671 | 3.78E-05 | 0.001649092511 | 1.242301963 |

| | | | | | | | |
|---|---|---|---|---|---|---|---|
| ENSG00000170[illegible] | 18.67611917 | 5.701561437 | 1.384391536 | 4.118460195 | 3.81E-05 | 0.001386950376 | 1.514850267 |
| ENSG00000181[illegible] | 16.02545325 | 5.49087221 | 1.333619781 | 4.11726962 | 3.83E-05 | 0.001327939999 | 1.605577974 |
| ENSG00000241[illegible] | 19.59936531 | 5.491236162 | 1.339768209 | 4.098646411 | 4.16E-05 | 0.001654365609 | 1.357988055 |
| ENSG00000170[illegible] | 105.19646 | 1.719820278 | 0.4197988733 | 4.09677202 | 4.19E-05 | 0.001799625444 | 1.22159078 |
| ENSG00000259[illegible] | 682.9148734 | -1.628192484 | 0.3974416616 | -4.096682962 | 4.19E-05 | 0.00145999971 | 1.572476891 |
| ENSG00000165[illegible] | 115.5277566 | -3.036198072 | 0.7415558902 | -4.09436175 | 4.23E-05 | 0.001925933483 | 1.130447911 |
| ENSG00000106[illegible] | 35.58820589 | 3.777021949 | 0.9226857519 | 4.093508479 | 4.25E-05 | 0.001720749514 | 1.319539707 |
| ENSG00000258[illegible] | 162.4583014 | -2.294748519 | 0.5610770861 | -4.089898832 | 4.32E-05 | 0.001818782099 | 1.242301963 |
| ENSG00000140[illegible] | 33.93110183 | 3.273760344 | 0.8005550083 | 4.089363392 | 4.33E-05 | 0.001744242157 | 1.319539707 |
| ENSG00000105[illegible] | 151.1008647 | 1.629202519 | 0.3986116809 | 4.087192116 | 4.37E-05 | 0.002277101965 | 0.9247574895 |
| ENSG00000171[illegible] | 136.6943843 | 1.745834636 | 0.4275693613 | 4.083161222 | 4.44E-05 | 0.001988460867 | 1.133466638 |
| ENSG00000141[illegible] | 134.7545678 | 2.345456915 | 0.5752967691 | 4.076951307 | 4.56E-05 | 0.001905872887 | 1.242301963 |
| ENSG00000169[illegible] | 101.5597227 | 2.01998443 | 0.4954948976 | 4.076700769 | 4.57E-05 | 0.001925933483 | 1.22159078 |
| ENSG00000105[illegible] | 612.191749 | 1.326722336 | 0.3259784136 | 4.069969913 | 4.70E-05 | 0.001631952189 | 1.564848717 |
| ENSG00000130[illegible] | 55.96596108 | 3.44091756 | 0.8456565843 | 4.068930135 | 4.72E-05 | 0.001720749514 | 1.470104381 |
| ENSG00000185[illegible] | 1218.023735 | 12.07423858 | 2.967502315 | 4.068821959 | 4.73E-05 | 0.001605525317 | 1.602205449 |
| ENSG00000165[illegible] | 61.41560649 | 2.469422205 | 0.6074921236 | 4.064945222 | 4.80E-05 | 0.001848115553 | 1.357988055 |
| ENSG00000039[illegible] | 97.42731917 | -3.123351982 | 0.7685049228 | -4.06419255 | 4.82E-05 | 0.001973527194 | 1.242301963 |
| ENSG00000135[illegible] | 15.99791156 | -5.008965302 | 1.232465603 | -4.064182637 | 4.82E-05 | 0.00185001025 | 1.357988055 |
| ENSG00000174[illegible] | 417.0278959 | -1.578051765 | 0.3886262189 | -4.06059007 | 4.89E-05 | 0.001916761549 | 1.32213648 |
| ENSG00000136[illegible] | 200.8159601 | 1.625962852 | 0.4006332185 | 4.058482365 | 4.94E-05 | 0.002054806746 | 1.198705341 |
| ENSG00000151[illegible] | 309.9761456 | 1.661986582 | 0.4098364035 | 4.055243916 | 5.01E-05 | 0.001943817688 | 1.32213648 |
| ENSG00000188[illegible] | 42.99822836 | 2.591904636 | 0.6393293118 | 4.054099489 | 5.03E-05 | 0.001757400604 | 1.516424702 |
| ENSG00000128[illegible] | 151.7778749 | -2.596703504 | 0.6407097721 | -4.052854532 | 5.06E-05 | 0.002087145823 | 1.198705341 |
| ENSG00000196[illegible] | 417.776976 | 1.332331254 | 0.3291526345 | 4.047761175 | 5.17E-05 | 0.002054806746 | 1.249985076 |
| ENSG00000105[illegible] | 6295.607924 | -1.671042508 | 0.4132114137 | -4.044037635 | 5.25E-05 | 0.001720749514 | 1.634803475 |
| ENSG00000166[illegible] | 597.6550834 | -1.406502968 | 0.3477987306 | -4.044014093 | 5.25E-05 | 0.00176135343 | 1.572476891 |
| ENSG00000227[illegible] | 440.0925184 | 1.584451059 | 0.3920261113 | 4.041697768 | 5.31E-05 | 0.001757400604 | 1.602205449 |
| ENSG00000137[illegible] | 5015.069763 | -1.32812648 | 0.3287646043 | -4.039748996 | 5.35E-05 | 0.00176135343 | 1.602205449 |
| ENSG00000123[illegible] | 1219.971381 | 1.596678765 | 0.395283467 | 4.039325949 | 5.36E-05 | 0.001744242157 | 1.634803475 |
| ENSG00000007[illegible] | 13.79355874 | 5.848337161 | 1.448061214 | 4.038736143 | 5.37E-05 | 0.001995486596 | 1.357988055 |
| ENSG00000089[illegible] | 15.5335712 | 4.393583192 | 1.088036351 | 4.038084929 | 5.39E-05 | 0.002022938963 | 1.337480197 |
| ENSG00000169[illegible] | 48.46164607 | 3.349354203 | 0.8295689853 | 4.037463144 | 5.40E-05 | 0.00200199969 | 1.357988055 |

| | | | | | | | |
|---|---|---|---|---|---|---|---|
| ENSG00000148 | 282.2165139 | 1.503298503 | 0.3726590311 | 4.033978456 | 5.48E-05 | 0.002087145823 | 1.29680518 |
| ENSG00000101 | 66.67563811 | -1.984461729 | 0.4929950584 | -4.025317688 | 5.69E-05 | 0.001988460867 | 1.449168615 |
| ENSG00000172 | 75.17368899 | 1.973910394 | 0.4905001635 | 4.024280807 | 5.71E-05 | 0.001992899473 | 1.449168615 |
| ENSG00000253 | 42.95771717 | 3.223045166 | 0.8016773679 | 4.020376894 | 5.81E-05 | 0.001957628557 | 1.516424702 |
| ENSG00000180 | 1183.8837 | -1.106881695 | 0.2756562303 | -4.01544233 | 5.93E-05 | 0.001682213517 | 1.902376403 |
| ENSG00000176 | 82.6970318 | 2.942932669 | 0.7329464478 | 4.015208311 | 5.94E-05 | 0.002164113557 | 1.337480197 |
| ENSG00000188 | 1679.070397 | -1.399002705 | 0.3485726874 | -4.013517856 | 5.98E-05 | 0.001925933483 | 1.597450288 |
| ENSG00000187 | 22.5265498 | 4.072616253 | 1.016564671 | 4.006253973 | 6.17E-05 | 0.002161844614 | 1.393563518 |
| ENSG00000243 | 467.738623 | -1.581516732 | 0.3947718562 | -4.006153699 | 6.17E-05 | 0.001855727624 | 1.729500602 |
| ENSG00000205 | 11.51056191 | 5.848592455 | 1.460002605 | 4.005878095 | 6.18E-05 | 0.00220809357 | 1.357988055 |
| ENSG00000156 | 32.34722412 | 3.342686237 | 0.8345684262 | 4.005287203 | 6.19E-05 | 0.002242880242 | 1.337480197 |
| ENSG00000094 | 15.69748185 | 5.463266337 | 1.364332948 | 4.004349776 | 6.22E-05 | 0.002052268686 | 1.514850267 |
| ENSG00000132 | 290.4623159 | 1.376571494 | 0.3440746598 | 4.000793011 | 6.31E-05 | 0.002405820611 | 1.249985076 |
| ENSG00000103 | 348.030891 | -1.4081106 | 0.3521119113 | -3.999042789 | 6.36E-05 | 0.002428714769 | 1.242301963 |
| ENSG00000120 | 3956.52499 | 1.626337907 | 0.4069733775 | 3.99617763 | 6.44E-05 | 0.001779656685 | 1.902376403 |
| ENSG00000145 | 3149.3598 | -1.12383819 | 0.2813621603 | -3.994276235 | 6.49E-05 | 0.002035144359 | 1.597450288 |
| ENSG00000108 | 80.26936633 | 3.322785982 | 0.8322171957 | 3.992690849 | 6.53E-05 | 0.002265551105 | 1.393563518 |
| ENSG00000213 | 106.3070297 | 2.880701295 | 0.7217761358 | 3.991128485 | 6.58E-05 | 0.002487946652 | 1.242301963 |
| ENSG00000106 | 363.447521 | -1.394571827 | 0.34947732 | -3.990450158 | 6.59E-05 | 0.002487946652 | 1.242301963 |
| ENSG00000157 | 25.68208222 | 3.459402274 | 0.8669768169 | 3.990190057 | 6.60E-05 | 0.002444559174 | 1.278627818 |
| ENSG00000136 | 568.298983 | -1.831499672 | 0.4591841915 | -3.988594785 | 6.65E-05 | 0.001959120753 | 1.729500602 |
| ENSG00000139 | 127.7008581 | -2.283593746 | 0.5730516683 | -3.984970068 | 6.75E-05 | 0.00271393813 | 1.133466638 |
| ENSG00000166 | 19.8430268 | 3.981603112 | 0.9993314033 | 3.984266979 | 6.77E-05 | 0.002087145823 | 1.605577974 |
| ENSG00000160 | 175.7512339 | 1.623986971 | 0.4082163905 | 3.978250282 | 6.94E-05 | 0.001957628557 | 1.814765822 |
| ENSG00000109 | 485.9153243 | 1.527346179 | 0.3839321302 | 3.978167126 | 6.94E-05 | 0.002161392037 | 1.572476891 |
| ENSG00000149 | 147.6991401 | 1.665006214 | 0.4186687496 | 3.976905885 | 6.98E-05 | 0.002686403836 | 1.198705341 |
| ENSG00000102 | 341.5720568 | 1.568266875 | 0.3944155911 | 3.976178707 | 7.00E-05 | 0.002487946652 | 1.32213648 |
| ENSG00000136 | 227.2781216 | 1.838191553 | 0.462396701 | 3.975356115 | 7.03E-05 | 0.002487946652 | 1.327567552 |
| ENSG00000225 | 920.8801258 | 1.592905059 | 0.4007247024 | 3.975060806 | 7.04E-05 | 0.002153700945 | 1.602205449 |
| ENSG00000058 | 122.9801099 | 1.858046707 | 0.4677447455 | 3.972351854 | 7.12E-05 | 0.002520356843 | 1.318051683 |
| ENSG00000198 | 982.4475216 | -1.201160607 | 0.3024665689 | -3.971217748 | 7.15E-05 | 0.00220809357 | 1.572476891 |
| ENSG00000165 | 131.0345933 | 1.875256496 | 0.4723443765 | 3.970104419 | 7.18E-05 | 0.002539150623 | 1.318051683 |
| ENSG00000198 | 21.01904598 | 4.981039159 | 1.255000257 | 3.968954691 | 7.22E-05 | 0.002520356843 | 1.337480197 |

| | | | | | | | |
|---|---|---|---|---|---|---|---|
| ENSG00000075 | 146.5910447 | 5.944349328 | 1.499586217 | 3.963993042 | 7.37E-05 | 0.00342474563 | 0.9247574895 |
| ENSG00000177 | 291.4889235 | -1.314647382 | 0.3318368109 | -3.961728595 | 7.44E-05 | 0.00271393813 | 1.249985076 |
| ENSG00000143 | 646.5131362 | 1.43781743 | 0.3632029464 | 3.958716316 | 7.54E-05 | 0.002301563895 | 1.572476891 |
| ENSG00000149 | 543.8343431 | -5.708413824 | 1.442026651 | -3.958604941 | 7.54E-05 | 0.002054806746 | 1.825593782 |
| ENSG00000185 | 458.8722206 | -1.257136407 | 0.3176343746 | -3.957809694 | 7.56E-05 | 0.00230555132 | 1.572476891 |
| ENSG00000139 | 22.72399878 | 3.973970168 | 1.004549524 | 3.955972375 | 7.62E-05 | 0.002669619385 | 1.319539707 |
| ENSG00000142 | 57.25920085 | -2.714399549 | 0.6862154166 | -3.955608521 | 7.63E-05 | 0.002402984985 | 1.516424702 |
| ENSG00000127 | 50.68343094 | 2.781537772 | 0.7036918493 | 3.952778158 | 7.72E-05 | 0.002669619385 | 1.337480197 |
| ENSG00000169 | 393.0763697 | -1.875125607 | 0.4745098882 | -3.951710289 | 7.76E-05 | 0.002691576749 | 1.32213648 |
| ENSG00000107 | 106.8371902 | 2.026699913 | 0.5130060404 | 3.950635573 | 7.79E-05 | 0.002868892316 | 1.22159078 |
| ENSG00000188 | 153.323494 | 1.614023084 | 0.4085749674 | 3.950371934 | 7.80E-05 | 0.003579320113 | 0.9247574895 |
| ENSG00000108 | 594.3912256 | 1.265587996 | 0.3204042297 | 3.949972811 | 7.82E-05 | 0.002108565195 | 1.825593782 |
| ENSG00000213 | 122.8808981 | 1.624982046 | 0.4119044281 | 3.945046315 | 7.98E-05 | 0.002753681628 | 1.318051683 |
| ENSG00000170 | 62.88806955 | 2.084086366 | 0.5291088055 | 3.938861618 | 8.19E-05 | 0.002774546225 | 1.337480197 |
| ENSG00000198 | 17.45724419 | 4.237896033 | 1.076066031 | 3.938323403 | 8.21E-05 | 0.002774546225 | 1.337480197 |
| ENSG00000114 | 12.53239227 | 5.865793815 | 1.489434483 | 3.938269109 | 8.21E-05 | 0.002694946914 | 1.393563518 |
| ENSG00000125 | 805.8587438 | -1.500631671 | 0.3813693034 | -3.934851751 | 8.32E-05 | 0.002420739019 | 1.634803475 |
| ENSG00000133 | 47.69692867 | 3.694972783 | 0.9396425487 | 3.932317441 | 8.41E-05 | 0.002833782637 | 1.337480197 |
| ENSG00000127 | 24.96527557 | -3.849588912 | 0.9798190031 | -3.928877578 | 8.53E-05 | 0.002833782637 | 1.357988055 |
| ENSG00000273 | 46.49011275 | -2.636352664 | 0.6711908103 | -3.927873599 | 8.57E-05 | 0.002622235211 | 1.516424702 |
| ENSG00000198 | 155.6639732 | 1.576299089 | 0.4015911044 | 3.925134477 | 8.67E-05 | 0.003112232505 | 1.22159078 |
| ENSG00000111 | 473.0739842 | -1.430445023 | 0.3645567009 | -3.923792979 | 8.72E-05 | 0.002589710545 | 1.564848717 |
| ENSG00000180 | 33.82188395 | 2.887872858 | 0.7368242986 | 3.919350737 | 8.88E-05 | 0.00299646147 | 1.319539707 |
| ENSG00000030 | 13.68530369 | 4.891602818 | 1.248082346 | 3.919294935 | 8.88E-05 | 0.002691576749 | 1.514850267 |
| ENSG00000005 | 63.60823389 | 2.474189357 | 0.6319640271 | 3.91507942 | 9.04E-05 | 0.003003342571 | 1.337480197 |
| ENSG00000133 | 160.7395109 | -1.482235824 | 0.3786771504 | -3.9142468 | 9.07E-05 | 0.003274958426 | 1.198705341 |
| ENSG00000111 | 456.8610561 | 1.38190297 | 0.3530653298 | 3.914014923 | 9.08E-05 | 0.0023783151 | 1.825593782 |
| ENSG00000157 | 794.9938884 | 1.464659279 | 0.3742615737 | 3.913464224 | 9.10E-05 | 0.002301563895 | 1.902376403 |
| ENSG00000123 | 19.63920394 | 3.995880756 | 1.021565691 | 3.911525994 | 9.17E-05 | 0.002936453196 | 1.393563518 |
| ENSG00000186 | 174.1946576 | 1.588745498 | 0.4064546546 | 3.908789036 | 9.28E-05 | 0.003280957335 | 1.22159078 |
| ENSG00000169 | 627.4853775 | 1.132218359 | 0.2899083045 | 3.905436105 | 9.41E-05 | 0.002691576749 | 1.602205449 |
| ENSG00000188 | 116.354117 | 1.652631772 | 0.4234527497 | 3.902753668 | 9.51E-05 | 0.003575878016 | 1.130447911 |
| ENSG00000235 | 14.41584488 | 4.522231378 | 1.159051436 | 3.901665825 | 9.55E-05 | 0.003097217463 | 1.357988055 |

| | | | | | | | |
|---|---|---|---|---|---|---|---|
| ENSG000001668 | 18.04402707 | 4.975113692 | 1.275450175 | 3.90067271 | 9.59E-05 | 0.003047876925 | 1.393563518 |
| ENSG000001699 | 10.60126142 | 5.577193159 | 1.429914215 | 3.900369058 | 9.60E-05 | 0.003107968659 | 1.357988055 |
| ENSG000001988 | 21.19081027 | 4.206034855 | 1.079089162 | 3.897763969 | 9.71E-05 | 0.003073002106 | 1.393563518 |
| ENSG000000776 | 13.21055336 | 5.729464399 | 1.470206963 | 3.89704616 | 9.74E-05 | 0.002884613341 | 1.514850267 |
| ENSG000001861 | 308.2178955 | -1.403953778 | 0.3604782733 | -3.894697356 | 9.83E-05 | 0.003217999183 | 1.327567552 |
| ENSG000001641 | 65.48954071 | 6.049909615 | 1.556590096 | 3.886642754 | 1.02E-04 | 0.003186949588 | 1.393563518 |
| ENSG000001856 | 3734.201406 | 1.073764022 | 0.2764847218 | 3.883628777 | 1.03E-04 | 0.002928207983 | 1.572476891 |
| ENSG000000889 | 513.1100674 | -1.847619981 | 0.4757723738 | -3.883411654 | 1.03E-04 | 0.002928207983 | 1.572476891 |
| ENSG000001170 | 203.0511076 | 1.493997638 | 0.3849434556 | 3.881083355 | 1.04E-04 | 0.003571303648 | 1.242301963 |
| ENSG000001565 | 52.62370238 | 2.162090967 | 0.5573112044 | 3.879503857 | 1.05E-04 | 0.003050442138 | 1.516424702 |
| ENSG000001649 | 149.5434351 | 2.286500774 | 0.5896610964 | 3.877652414 | 1.05E-04 | 0.003629303032 | 1.22159078 |
| ENSG000001380 | 471.0987689 | -1.217852509 | 0.3141519421 | -3.876635302 | 1.06E-04 | 0.002774546225 | 1.729500602 |
| ENSG000001577 | 26.20964633 | -3.12357862 | 0.8059272203 | -3.875757688 | 1.06E-04 | 0.003375747146 | 1.357988055 |
| ENSG000001987 | 225.7367086 | -1.371354633 | 0.3542102503 | -3.871583704 | 1.08E-04 | 0.009369574417 | 0.3939394882 |
| ENSG000002860 | 14.39001333 | -5.48662627 | 1.418740998 | -3.867250101 | 1.10E-04 | 0.003469920766 | 1.357988055 |
| ENSG000001637 | 307.4805073 | -6.215280336 | 1.607737802 | -3.865854449 | 1.11E-04 | 0.003571303648 | 1.32213648 |
| ENSG000001818 | 63.25058471 | 2.235865981 | 0.5784448622 | 3.865305281 | 1.11E-04 | 0.00327313481 | 1.470104381 |
| ENSG000001675 | 21.06829423 | 3.882218014 | 1.004588198 | 3.864486985 | 1.11E-04 | 0.003426153284 | 1.393563518 |
| ENSG000001093 | 299.0756722 | 1.649694866 | 0.4273819219 | 3.860001514 | 1.13E-04 | 0.003618103749 | 1.32213648 |
| ENSG000001524 | 12.20944002 | -5.376509659 | 1.393219143 | -3.859055259 | 1.14E-04 | 0.003603551786 | 1.337480197 |
| ENSG000001885 | 728.9600246 | 1.716662096 | 0.4451193381 | 3.856633377 | 1.15E-04 | 0.003188461516 | 1.572476891 |
| ENSG000001173 | 458.6530555 | -1.685251505 | 0.4370729712 | -3.855766922 | 1.15E-04 | 0.00315223439 | 1.602205449 |
| ENSG000001248 | 30.94460212 | -3.406376119 | 0.8844203285 | -3.851535304 | 1.17E-04 | 0.003705819059 | 1.319539707 |
| ENSG000001543 | 669.222775 | 1.062329534 | 0.2759066284 | 3.850322626 | 1.18E-04 | 0.003205030224 | 1.602205449 |
| ENSG000001130 | 20.79408813 | 4.526014483 | 1.177604972 | 3.843406396 | 1.21E-04 | 0.003650579824 | 1.393563518 |
| ENSG000001664 | 42.21170518 | -2.831071885 | 0.7372908478 | -3.8398305 | 1.23E-04 | 0.00368716312 | 1.393563518 |
| ENSG000001253 | 16.00336586 | 5.299306677 | 1.380827478 | 3.837776088 | 1.24E-04 | 0.003858790353 | 1.337480197 |
| ENSG000000166 | 91.84626442 | -2.417341698 | 0.6306676983 | -3.832987966 | 1.27E-04 | 0.003652372087 | 1.449168615 |
| ENSG000001621 | 334.7951177 | -1.310609661 | 0.3422257393 | -3.829664197 | 1.28E-04 | 0.004078333993 | 1.29680518 |
| ENSG000001703 | 1412.47595 | 1.217047054 | 0.3179817896 | 3.827411172 | 1.29E-04 | 0.003409983654 | 1.634803475 |
| ENSG000001680 | 119.1319682 | 1.938220427 | 0.5076615691 | 3.817938061 | 1.35E-04 | 0.004740269375 | 1.130447911 |
| ENSG000000549 | 62.00809925 | 2.11427928 | 0.5537829473 | 3.817884407 | 1.35E-04 | 0.003988055845 | 1.393563518 |
| ENSG000000064 | 493.7733377 | -1.384753079 | 0.3627390288 | -3.817491278 | 1.35E-04 | 0.003575878016 | 1.602205449 |

| | | | | | | | |
|---|---|---|---|---|---|---|---|
| ENSG0000013 | 345.592215 | 1.424228491 | 0.373364252 | 3.814581828 | 1.36E-04 | 0.004282481321 | 1.29680518 |
| ENSG0000016 | 22.02825495 | 3.271678077 | 0.8578071384 | 3.814001925 | 1.37E-04 | 0.004128069676 | 1.357988055 |
| ENSG0000017 | 3439.801269 | 1.258009122 | 0.3299590134 | 3.812622389 | 1.38E-04 | 0.00362480651 | 1.597450288 |
| ENSG0000010 | 40.57695294 | -2.656938258 | 0.6969103962 | -3.812453183 | 1.38E-04 | 0.00420289985 | 1.337480197 |
| ENSG0000010 | 608.0513268 | -1.305940321 | 0.3428747932 | -3.808796525 | 1.40E-04 | 0.003650579824 | 1.602205449 |
| ENSG0000011 | 99.10170966 | 1.587449756 | 0.4169179139 | 3.807583468 | 1.40E-04 | 0.00462259127 | 1.22159078 |
| ENSG0000018 | 91.23353771 | -1.95090954 | 0.5124726837 | -3.806855666 | 1.41E-04 | 0.003858790353 | 1.514054369 |
| ENSG0000016 | 90.13777749 | 2.243145368 | 0.5901600233 | 3.800910395 | 1.44E-04 | 0.004093002443 | 1.449168615 |
| ENSG0000018 | 37.36411452 | 2.9215907 | 0.7687605785 | 3.800390891 | 1.44E-04 | 0.004227837882 | 1.393563518 |
| ENSG0000005 | 493.7750785 | 1.10867801 | 0.2923924931 | 3.791745807 | 1.50E-04 | 0.003941934722 | 1.572476891 |
| ENSG0000015 | 25.93059801 | 3.529510859 | 0.9313768276 | 3.789562672 | 1.51E-04 | 0.004724151815 | 1.278627818 |
| ENSG0000016 | 308.106313 | 1.172459545 | 0.3094952917 | 3.788295254 | 1.52E-04 | 0.004696087747 | 1.29680518 |
| ENSG0000018 | 1298.047935 | 1.414972254 | 0.3736670501 | 3.786719364 | 1.53E-04 | 0.003952569255 | 1.597450288 |
| ENSG0000010 | 24.38547304 | 3.742740292 | 0.9884036967 | 3.786651451 | 1.53E-04 | 0.004554894289 | 1.357988055 |
| ENSG0000027 | 18.17155177 | 4.385962817 | 1.158423643 | 3.786147534 | 1.53E-04 | 0.004554894289 | 1.357988055 |
| ENSG0000017 | 149.343877 | 1.632934411 | 0.4316871465 | 3.782680175 | 1.55E-04 | 0.005108209176 | 1.198705341 |
| ENSG0000024 | 194.5363244 | 1.532559662 | 0.4051657358 | 3.782550019 | 1.55E-04 | 0.005023663059 | 1.22159078 |
| ENSG0000024 | 504.8688735 | -1.128068142 | 0.298327783 | -3.781304345 | 1.56E-04 | 0.003611856737 | 1.825593782 |
| ENSG0000016 | 797.4475505 | 1.257609137 | 0.3331800016 | 3.774563693 | 1.60E-04 | 0.004108795058 | 1.602205449 |
| ENSG0000005 | 15.38928922 | 4.669784468 | 1.237324416 | 3.774098697 | 1.61E-04 | 0.004724151815 | 1.357988055 |
| ENSG0000016 | 122.846741 | -1.756105245 | 0.4653259165 | -3.773925292 | 1.61E-04 | 0.005520495622 | 1.133466638 |
| ENSG0000015 | 819.7699963 | 1.070032893 | 0.2838005783 | 3.770368966 | 1.63E-04 | 0.004176159986 | 1.597450288 |
| ENSG0000016 | 120.1719967 | 2.167254844 | 0.5753610134 | 3.766773892 | 1.65E-04 | 0.00531568455 | 1.22159078 |
| ENSG0000006 | 183.8985349 | 5.017830456 | 1.332359419 | 3.766123752 | 1.66E-04 | 0.005320550867 | 1.22159078 |
| ENSG0000012 | 100.5636279 | 1.902585369 | 0.5053970536 | 3.764535933 | 1.67E-04 | 0.005345456867 | 1.22159078 |
| ENSG0000010 | 50.90898291 | 2.063284496 | 0.5485519102 | 3.761329525 | 1.69E-04 | 0.005004282739 | 1.337480197 |
| ENSG0000013 | 25.71420461 | 4.072319266 | 1.084127918 | 3.756308823 | 1.72E-04 | 0.004908587517 | 1.393563518 |
| ENSG0000016 | 846.1843374 | 1.149015451 | 0.3064777561 | 3.749099007 | 1.77E-04 | 0.004514943203 | 1.597450288 |
| ENSG0000014 | 38.05411853 | -3.684334866 | 0.9841867943 | -3.74353211 | 1.81E-04 | 0.005372409798 | 1.319539707 |
| ENSG0000011 | 3628.634231 | 1.263139841 | 0.3375953612 | 3.741579375 | 1.83E-04 | 0.004606211093 | 1.602205449 |
| ENSG0000015 | 1171.998391 | 1.327492598 | 0.3548911012 | 3.740563213 | 1.84E-04 | 0.00462259127 | 1.597450288 |
| ENSG0000013 | 427.1432147 | -1.092969147 | 0.2922134549 | -3.74031082 | 1.84E-04 | 0.005675145699 | 1.242301963 |
| ENSG0000026 | 3862.668966 | 3.347085771 | 0.896533996 | 3.733361797 | 1.89E-04 | 0.004724151815 | 1.597450288 |

| | | | | | | | |
|---|---|---|---|---|---|---|---|
| ENSG00000144[illegible] | 18.94221631 | 3.951064867 | 1.059915294 | 3.727717573 | 1.93E-04 | 0.005587819552 | 1.337480197 |
| ENSG00000134[illegible] | 50.8400627 | 2.05692784 | 0.5520252723 | 3.726147956 | 1.94E-04 | 0.005610215187 | 1.337480197 |
| ENSG00000249[illegible] | 473.9931848 | -1.255476125 | 0.3369597534 | -3.725893411 | 1.95E-04 | 0.004554894289 | 1.729500602 |
| ENSG00000182[illegible] | 9.915700236 | 5.766338414 | 1.547648323 | 3.725871264 | 1.95E-04 | 0.005610215187 | 1.337480197 |
| ENSG00000189[illegible] | 4149.515245 | -1.087620956 | 0.2919149272 | -3.7258148 | 1.95E-04 | 0.004740269375 | 1.634803475 |
| ENSG00000135[illegible] | 115.9064287 | 1.636296444 | 0.4396376941 | 3.721920268 | 1.98E-04 | 0.006064953308 | 1.242301963 |
| ENSG00000144[illegible] | 270.9234721 | 1.643275769 | 0.4415727913 | 3.721415361 | 1.98E-04 | 0.005714887788 | 1.327567552 |
| ENSG00000009[illegible] | 8.591159865 | 6.171518777 | 1.659870477 | 3.718072502 | 2.01E-04 | 0.005581139422 | 1.393563518 |
| ENSG00000137[illegible] | 1278.318539 | 1.330188022 | 0.3578738926 | 3.716918305 | 2.02E-04 | 0.004901883476 | 1.634803475 |
| ENSG00000164[illegible] | 435.9901964 | 1.397433341 | 0.377302558 | 3.703747329 | 2.12E-04 | 0.00521524202 | 1.602205449 |
| ENSG00000110[illegible] | 210.6472488 | -1.493167841 | 0.40319476 | -3.703341386 | 2.13E-04 | 0.006612693782 | 1.198705341 |
| ENSG00000050[illegible] | 24.59078784 | -3.265647606 | 0.8820425341 | -3.702369761 | 2.14E-04 | 0.00585053097 | 1.393563518 |
| ENSG00000197[illegible] | 32.40700587 | -3.03651703 | 0.8205459847 | -3.700605556 | 2.15E-04 | 0.00632742371 | 1.278627818 |
| ENSG00000183[illegible] | 259.5581416 | 6.326324826 | 1.709983599 | 3.699640645 | 2.16E-04 | 0.01574826892 | 0.3939394882 |
| ENSG00000197[illegible] | 161.6289419 | 1.533100702 | 0.4155946363 | 3.688932839 | 2.25E-04 | 0.00858759264 | 0.9247574895 |
| ENSG00000135[illegible] | 13.23299141 | 5.315504977 | 1.442490408 | 3.684949964 | 2.29E-04 | 0.006214548622 | 1.393563518 |
| ENSG00000101[illegible] | 761.4177962 | 1.023363801 | 0.2777840882 | 3.684025992 | 2.30E-04 | 0.005646388378 | 1.564848717 |
| ENSG00000206[illegible] | 65.42902942 | 2.021278989 | 0.5491546021 | 3.680710279 | 2.33E-04 | 0.006288083524 | 1.393563518 |
| ENSG00000172[illegible] | 22.13996406 | 3.839666314 | 1.043213218 | 3.680615093 | 2.33E-04 | 0.006423988161 | 1.357988055 |
| ENSG00000125[illegible] | 3568.594174 | 1.26127276 | 0.34301878 | 3.676978737 | 2.36E-04 | 0.005675145699 | 1.597450288 |
| ENSG00000068[illegible] | 274.032691 | 1.733143154 | 0.4715455997 | 3.675451866 | 2.37E-04 | 0.007029709672 | 1.242301963 |
| ENSG00000144[illegible] | 224.2780181 | -1.451265341 | 0.3950036241 | -3.674055761 | 2.39E-04 | 0.007057086457 | 1.242301963 |
| ENSG00000130[illegible] | 38.45231421 | -2.94918347 | 0.8031545049 | -3.67200016 | 2.41E-04 | 0.006751016881 | 1.319539707 |
| ENSG00000114[illegible] | 200.3474294 | 1.30534748 | 0.3555771568 | 3.671066758 | 2.42E-04 | 0.006745490712 | 1.327567552 |
| ENSG00000025[illegible] | 325.384016 | 1.168119162 | 0.3184095363 | 3.668606082 | 2.44E-04 | 0.007175198182 | 1.242301963 |
| ENSG00000006[illegible] | 51.33034087 | 2.191413739 | 0.597439238 | 3.66801107 | 2.44E-04 | 0.006122889403 | 1.516424702 |
| ENSG00000152[illegible] | 148.6009506 | 1.652339187 | 0.450623818 | 3.666781739 | 2.46E-04 | 0.007454228107 | 1.198705341 |
| ENSG00000164[illegible] | 76.62759367 | 1.727261117 | 0.4712123633 | 3.665568334 | 2.47E-04 | 0.006607859536 | 1.393563518 |
| ENSG00000177[illegible] | 5988.226103 | 1.424149171 | 0.3885281111 | 3.665498404 | 2.47E-04 | 0.005112647335 | 1.902376403 |
| ENSG00000287[illegible] | 30.70516422 | -3.843403073 | 1.050905162 | -3.657231129 | 2.55E-04 | 0.007028167484 | 1.337480197 |
| ENSG00000099[illegible] | 179.7176739 | 1.490745421 | 0.4077478902 | 3.656046926 | 2.56E-04 | 0.00942687464 | 0.9247574895 |
| ENSG00000274[illegible] | 11.75459794 | 4.63199982 | 1.267947895 | 3.653146819 | 2.59E-04 | 0.007662554734 | 1.227924265 |
| ENSG00000163[illegible] | 674.6101659 | 1.27748995 | 0.3497503692 | 3.652576415 | 2.60E-04 | 0.005528214639 | 1.825593782 |

| | | | | | | | |
|---|---|---|---|---|---|---|---|
| ENSG00000262 | 1204.505317 | 4.385007716 | 1.20085714 | 3.651564844 | 2.61E-04 | 0.006065891622 | 1.634803475 |
| ENSG00000185 | 939.1664707 | 1.361600943 | 0.3731056736 | 3.649370781 | 2.63E-04 | 0.005389751549 | 1.902376403 |
| ENSG00000065 | 305.978753 | 1.878478265 | 0.5150000216 | 3.64753046 | 2.65E-04 | 0.00766708916 | 1.249985076 |
| ENSG00000158 | 15.08388559 | 5.007468752 | 1.373398196 | 3.646042908 | 2.66E-04 | 0.006580723296 | 1.514850267 |
| ENSG00000268 | 62.68030165 | 11.01765623 | 3.021993632 | 3.645823775 | 2.67E-04 | 0.007175198182 | 1.357988055 |
| ENSG00000175 | 56.70679927 | 2.252181113 | 0.6180268957 | 3.644147413 | 2.68E-04 | 0.006607859536 | 1.516424702 |
| ENSG00000144 | 48.96111916 | 2.547733034 | 0.6996402848 | 3.641489904 | 2.71E-04 | 0.00664845022 | 1.516424702 |
| ENSG00000207 | 12.49608552 | 4.68291294 | 1.286149513 | 3.641033095 | 2.72E-04 | 0.006262032649 | 1.636424459 |
| ENSG00000135 | 825.8240857 | -1.093382205 | 0.3006060311 | -3.637259707 | 2.76E-04 | 0.006329929086 | 1.634803475 |
| ENSG00000244 | 307.8175444 | -2.177528681 | 0.5991951828 | -3.634089097 | 2.79E-04 | 0.00776491136 | 1.29680518 |
| ENSG00000288 | 88.08263785 | -11.06930387 | 3.046898395 | -3.6329744 | 2.80E-04 | 0.007088691288 | 1.449168615 |
| ENSG00000180 | 432.1321493 | -1.334238885 | 0.3677907121 | -3.627712285 | 2.86E-04 | 0.006249321279 | 1.729500602 |
| ENSG00000131 | 151.969318 | 1.660592721 | 0.4578577123 | 3.626875067 | 2.87E-04 | 0.00834704798 | 1.22159078 |
| ENSG00000013 | 1544.247469 | -1.040673755 | 0.2869649449 | -3.626483909 | 2.87E-04 | 0.006658884326 | 1.602205449 |
| ENSG00000046 | 99.46936869 | 2.558536024 | 0.7057965277 | 3.625033454 | 2.89E-04 | 0.008778481756 | 1.133466638 |
| ENSG00000140 | 1946.674819 | -2.959383214 | 0.8167893945 | -3.623190058 | 2.91E-04 | 0.005848291584 | 1.902376403 |
| ENSG00000135 | 263.0032272 | 1.322421493 | 0.3653781139 | 3.619323224 | 2.95E-04 | 0.02119642399 | 0.3706356727 |
| ENSG00000106 | 620.7895028 | -1.225661607 | 0.3386538553 | -3.619216459 | 2.95E-04 | 0.006138125892 | 1.825593782 |
| ENSG00000124 | 117.3224297 | -1.540516516 | 0.4258672357 | -3.617363317 | 2.98E-04 | 0.008469364285 | 1.242301963 |
| ENSG00000178 | 87.33037617 | -1.951717876 | 0.5397684217 | -3.615843013 | 2.99E-04 | 0.007215695133 | 1.514054369 |
| ENSG00000152 | 101.1876643 | -1.963651672 | 0.5433562723 | -3.613930255 | 3.02E-04 | 0.00858759264 | 1.22159078 |
| ENSG00000064 | 117.4565637 | 1.733990883 | 0.4801132924 | 3.611628569 | 3.04E-04 | 0.00858759264 | 1.242301963 |
| ENSG00000169 | 167.4677096 | 1.441046772 | 0.3990844216 | 3.610882044 | 3.05E-04 | 0.00858759264 | 1.242301963 |
| ENSG00000125 | 379.5761897 | 1.188097566 | 0.3292475768 | 3.608523341 | 3.08E-04 | 0.00858759264 | 1.242301963 |
| ENSG00000170 | 75.42525214 | -1.98354875 | 0.5500289003 | -3.606262777 | 3.11E-04 | 0.007996535008 | 1.393563518 |
| ENSG00000035 | 2391.594914 | 1.000981025 | 0.2776472017 | 3.605226413 | 3.12E-04 | 0.007028167484 | 1.634803475 |
| ENSG00000102 | 69.24425785 | 2.564403293 | 0.7116378441 | 3.603522935 | 3.14E-04 | 0.008068889778 | 1.393563518 |
| ENSG00000079 | 122.8870575 | 2.312740954 | 0.6419753176 | 3.602538744 | 3.15E-04 | 0.009435884187 | 1.130447911 |
| ENSG00000145 | 157.5787723 | 2.107789109 | 0.5855252311 | 3.59982627 | 3.18E-04 | 0.008814306519 | 1.242301963 |
| ENSG00000134 | 37.60778427 | 2.467117456 | 0.686390105 | 3.594337154 | 3.25E-04 | 0.00858759264 | 1.319539707 |
| ENSG00000123 | 281.1906351 | -1.344771888 | 0.3741938478 | -3.593784065 | 3.26E-04 | 0.00896829308 | 1.242301963 |
| ENSG00000248 | 9.80446049 | 4.881422051 | 1.358437312 | 3.593409875 | 3.26E-04 | 0.007266967691 | 1.636424459 |
| ENSG00000227 | 23.38372566 | 3.576937802 | 0.9955276324 | 3.59300705 | 3.27E-04 | 0.00858759264 | 1.319539707 |

| | | | | | | | |
|---|---|---|---|---|---|---|---|
| ENSG000002767 | 21.09988427 | 3.658114703 | 1.018129135 | 3.592977137 | 3.27E-04 | 0.00858759264 | 1.337480197 |
| ENSG000001777 | 39.68060649 | -2.439661327 | 0.6795200107 | -3.590271499 | 3.30E-04 | 0.00858759264 | 1.337480197 |
| ENSG000002768 | 56.69242609 | 3.330081208 | 0.9279038397 | 3.588821455 | 3.32E-04 | 0.007882489812 | 1.516424702 |
| ENSG000000736 | 7.824045532 | 6.187461099 | 1.724772238 | 3.587407638 | 3.34E-04 | 0.008469364285 | 1.393563518 |
| ENSG000001766 | 23.65869969 | 2.833150086 | 0.7898005972 | 3.587171365 | 3.34E-04 | 0.00858759264 | 1.357988055 |
| ENSG000001509 | 211.8374527 | 1.284171449 | 0.3582127879 | 3.584940272 | 3.37E-04 | 0.009466259118 | 1.198705341 |
| ENSG000002877 | 13.96708612 | 4.57815173 | 1.277217476 | 3.584473135 | 3.38E-04 | 0.008709802878 | 1.337480197 |
| ENSG000002586 | 896.3812119 | 1.031607016 | 0.2878621783 | 3.583683767 | 3.39E-04 | 0.00766708916 | 1.597450288 |
| ENSG000001571 | 266.8208962 | -1.283321383 | 0.3583179934 | -3.58151532 | 3.42E-04 | 0.02270035464 | 0.3939394882 |
| ENSG000001962 | 31.27619719 | 2.711048836 | 0.7571204143 | 3.580736677 | 3.43E-04 | 0.00858759264 | 1.393563518 |
| ENSG000001522 | 88.55717101 | 2.08496674 | 0.5825164376 | 3.57924104 | 3.45E-04 | 0.008127049139 | 1.514054369 |
| ENSG000002447 | 3698.190833 | 2.824284313 | 0.7892322754 | 3.578521052 | 3.46E-04 | 0.007665280589 | 1.634803475 |
| ENSG000001601 | 10664.20443 | -1.234289416 | 0.3451081953 | -3.576528849 | 3.48E-04 | 0.007832181635 | 1.602205449 |
| ENSG000001849 | 16.03059882 | 4.744682658 | 1.328155276 | 3.572385506 | 3.54E-04 | 0.008312939465 | 1.514850267 |
| ENSG000002504 | 78.19002663 | 1.85161333 | 0.5185247003 | 3.570925992 | 3.56E-04 | 0.00858759264 | 1.449168615 |
| ENSG000001652 | 1226.247213 | -1.23407963 | 0.3457390496 | -3.569396143 | 3.58E-04 | 0.006951004644 | 1.902376403 |
| ENSG000001825 | 823.6664283 | 1.062538289 | 0.2979501801 | 3.566160922 | 3.62E-04 | 0.008213410704 | 1.572476891 |
| ENSG000001113 | 901.9856648 | -1.303426619 | 0.365523104 | -3.56592129 | 3.63E-04 | 0.008093208773 | 1.602205449 |
| ENSG000001122 | 292.1240887 | 1.380076564 | 0.3876328632 | 3.560267188 | 3.70E-04 | 0.009466259118 | 1.32213648 |
| ENSG000000476 | 83.27099392 | 1.678511299 | 0.4718224572 | 3.557506162 | 3.74E-04 | 0.008866755553 | 1.449168615 |
| ENSG000001341 | 5495.665604 | -2.591711089 | 0.7285653874 | -3.557280011 | 3.75E-04 | 0.008457136918 | 1.572476891 |
| ENSG000001816 | 460.0599485 | -1.25529464 | 0.3529104927 | -3.556977382 | 3.75E-04 | 0.008469364285 | 1.564848717 |
| ENSG000001336 | 182.7591609 | -2.067401818 | 0.5813988297 | -3.555909837 | 3.77E-04 | 0.01265853831 | 0.9247574895 |
| ENSG000001786 | 64.11903133 | 2.27086531 | 0.6390096349 | 3.553726245 | 3.80E-04 | 0.008664890237 | 1.514054369 |
| ENSG000002722 | 9.091675859 | 6.176888442 | 1.740500194 | 3.548915687 | 3.87E-04 | 0.009527466429 | 1.357988055 |
| ENSG000001238 | 11.23549649 | -5.46742234 | 1.541025345 | -3.547912018 | 3.88E-04 | 0.01033665786 | 1.227924265 |
| ENSG000000888 | 44.98550894 | 2.564344877 | 0.723140838 | 3.546120952 | 3.91E-04 | 0.009748676789 | 1.337480197 |
| ENSG000001822 | 1194.063412 | -1.073736138 | 0.3028501826 | -3.545436655 | 3.92E-04 | 0.008622634283 | 1.572476891 |
| ENSG000001854 | 44.28223321 | 2.637331906 | 0.7438929985 | 3.545310833 | 3.92E-04 | 0.009644853854 | 1.357988055 |
| ENSG000001472 | 12.25507675 | 4.331904919 | 1.221920037 | 3.545162356 | 3.92E-04 | 0.009466259118 | 1.393563518 |
| ENSG000001983 | 30.70214302 | 3.0172281 | 0.8511364886 | 3.544940371 | 3.93E-04 | 0.009466259118 | 1.393563518 |
| ENSG000002818 | 30.3592549 | 3.084057585 | 0.8704003949 | 3.543263081 | 3.95E-04 | 0.009512928164 | 1.393563518 |
| ENSG000002446 | 157.7199557 | -1.506548586 | 0.4252322421 | -3.542884185 | 3.96E-04 | 0.0105186363 | 1.22159078 |

| | | | | | | | |
|---|---|---|---|---|---|---|---|
| ENSG000000598 | 832.8611442 | 1.460160249 | 0.4122139537 | 3.542238772 | 3.97E-04 | 0.00858759264 | 1.602205449 |
| ENSG000001966 | 10.13575534 | 5.343686044 | 1.508586461 | 3.542180832 | 3.97E-04 | 0.009527466429 | 1.393563518 |
| ENSG000002798 | 32.76897692 | -2.956401076 | 0.8350425038 | -3.540419874 | 3.99E-04 | 0.00991904237 | 1.337480197 |
| ENSG000000810 | 193.5932004 | -1.311398606 | 0.3704090286 | -3.540406698 | 4.00E-04 | 0.007909553317 | 1.814765822 |
| ENSG000001195 | 161.2477062 | 2.160603374 | 0.6103512565 | 3.539934342 | 4.00E-04 | 0.01315118477 | 0.9247574895 |
| ENSG000001141 | 102.9114016 | 1.670541642 | 0.4722658939 | 3.537290461 | 4.04E-04 | 0.01017074999 | 1.318051683 |
| ENSG000001369 | 165.7014366 | -1.433218701 | 0.4051753915 | -3.537279734 | 4.04E-04 | 0.01090384694 | 1.198705341 |
| ENSG000001660 | 1484.652253 | 1.1552037 | 0.3271815273 | 3.530772992 | 4.14E-04 | 0.008866755553 | 1.602205449 |
| ENSG000001066 | 869.0032584 | 1.275832414 | 0.3613787487 | 3.530457779 | 4.15E-04 | 0.00889063967 | 1.597450288 |
| ENSG000001757 | 416.4561879 | 1.200775674 | 0.3403433715 | 3.528130044 | 0.000418506497 | 0.01033665786 | 1.32213648 |
| ENSG000001805 | 12.13609804 | 4.418454415 | 1.253474148 | 3.524966528 | 4.24E-04 | 0.010240229 | 1.357988055 |
| ENSG000002543 | 16.83289738 | 3.795885095 | 1.077423162 | 3.523114436 | 4.27E-04 | 0.009466259118 | 1.514850267 |
| ENSG000000914 | 15.18701867 | 5.323766505 | 1.512059033 | 3.520872127 | 4.30E-04 | 0.01020896867 | 1.393563518 |
| ENSG000000992 | 59.90434257 | 2.401050561 | 0.6820344919 | 3.520423952 | 4.31E-04 | 0.01020896867 | 1.393563518 |
| ENSG000002150 | 93.95336875 | 1.771665422 | 0.5032776252 | 3.520254692 | 4.31E-04 | 0.0104801062 | 1.337480197 |
| ENSG000002034 | 114.5580487 | -1.971089616 | 0.5600074983 | -3.519755757 | 4.32E-04 | 0.01114835695 | 1.242301963 |
| ENSG000001187 | 118.1320947 | 1.581661418 | 0.4495345317 | 3.518442537 | 4.34E-04 | 0.01206267293 | 1.130447911 |
| ENSG000001800 | 13.61536042 | -5.00892243 | 1.424025051 | -3.517439829 | 4.36E-04 | 0.01044953893 | 1.357988055 |
| ENSG000001512 | 274.5233361 | 1.143265335 | 0.3250379628 | 3.517328637 | 4.36E-04 | 0.01064566953 | 1.327567552 |
| ENSG000001874 | 1283.210056 | 1.248433913 | 0.3549737637 | 3.51697517 | 4.36E-04 | 0.009433801795 | 1.572476891 |
| ENSG000001619 | 532.3227054 | -1.184868391 | 0.3370167925 | -3.515754756 | 4.39E-04 | 0.009435884187 | 1.572476891 |
| ENSG000001987 | 120.3895967 | 1.525058453 | 0.4337962593 | 3.515609967 | 4.39E-04 | 0.01145268086 | 1.22159078 |
| ENSG000000683 | 1792.298681 | 1.123722276 | 0.3197683012 | 3.514176583 | 4.41E-04 | 0.009384299939 | 1.602205449 |
| ENSG000000445 | 61.82879796 | 2.079366435 | 0.5927815274 | 3.507812472 | 4.52E-04 | 0.0109064077 | 1.337480197 |
| ENSG000001684 | 842.1773634 | -1.094280641 | 0.3121614254 | -3.50549604 | 4.56E-04 | 0.009435884187 | 1.634803475 |
| ENSG000001624 | 55.4281034 | 2.005754425 | 0.5722888318 | 3.504793933 | 4.57E-04 | 0.01022020535 | 1.470104381 |
| ENSG000001643 | 149.0226454 | 1.581868845 | 0.4515074391 | 3.503527755 | 4.59E-04 | 0.0120485658 | 1.198705341 |
| ENSG000001368 | 39.57311773 | 2.319423432 | 0.6622131984 | 3.502532776 | 4.61E-04 | 0.01094166169 | 1.357988055 |
| ENSG000002249 | 64.34976827 | 1.84771731 | 0.5278542283 | 3.50043101 | 4.65E-04 | 0.01114835695 | 1.337480197 |
| ENSG000001121 | 28.62507752 | 2.894679233 | 0.8270167772 | 3.500145719 | 4.65E-04 | 0.01158077996 | 1.278627818 |
| ENSG000000543 | 67.19469812 | 2.164852266 | 0.6188842385 | 3.497992244 | 4.69E-04 | 0.01020896867 | 1.514054369 |
| ENSG000002101 | 229.4171748 | -1.980402207 | 0.5663167072 | -3.496987079 | 4.71E-04 | 0.02963004923 | 0.3706356727 |
| ENSG000001043 | 162.3884798 | 1.68550397 | 0.4820456612 | 3.496564964 | 4.71E-04 | 0.01492065348 | 0.9247574895 |

| | | | | | | | |
|---|---|---|---|---|---|---|---|
| ENSG00000141 | 13.01318793 | 4.33262986 | 1.239858105 | 3.494456216 | 4.75E-04 | 0.01137218553 | 1.337480197 |
| ENSG00000172 | 82.9178269 | 1.839957908 | 0.5270851289 | 3.490817341 | 4.82E-04 | 0.01111016134 | 1.393563518 |
| ENSG00000101 | 1432.413791 | -1.020900704 | 0.2925468189 | -3.489700241 | 4.84E-04 | 0.009850992487 | 1.634803475 |
| ENSG00000204 | 510.8408723 | -1.160460288 | 0.3327864261 | -3.487102228 | 4.88E-04 | 0.01022020535 | 1.572476891 |
| ENSG00000175 | 197.3280302 | -1.875570341 | 0.5379103382 | -3.486771321 | 4.89E-04 | 0.03044962766 | 0.3706356727 |
| ENSG00000035 | 44.45531415 | -2.905688801 | 0.8336203507 | -3.485626039 | 4.91E-04 | 0.01165788223 | 1.337480197 |
| ENSG00000057 | 125.1093916 | -1.420024766 | 0.4074681856 | -3.484995433 | 4.92E-04 | 0.01319458635 | 1.130447911 |
| ENSG00000101 | 280.5177727 | -1.655343348 | 0.4750609622 | -3.484486159 | 4.93E-04 | 0.01234168874 | 1.249985076 |
| ENSG00000197 | 52.53748623 | 2.156348574 | 0.6189431159 | 3.483920442 | 4.94E-04 | 0.01088212293 | 1.470104381 |
| ENSG00000119 | 18.42624992 | 4.273585507 | 1.227738027 | 3.480861073 | 5.00E-04 | 0.01145223853 | 1.393563518 |
| ENSG00000069 | 298.6845792 | 1.048081686 | 0.3012722441 | 3.478852457 | 5.04E-04 | 0.01261411097 | 1.242301963 |
| ENSG00000169 | 40.40971022 | 2.406004982 | 0.6916118453 | 3.478837151 | 5.04E-04 | 0.011029196 | 1.470104381 |
| ENSG00000081 | 284.665006 | 1.356779044 | 0.3901819985 | 3.477297899 | 5.06E-04 | 0.01261411097 | 1.249985076 |
| ENSG00000113 | 464.5825067 | 1.104322457 | 0.3178787184 | 3.474037086 | 5.13E-04 | 0.009858374848 | 1.729500602 |
| ENSG00000163 | 689.9813422 | -1.042088342 | 0.3000345807 | -3.473227451 | 5.14E-04 | 0.01044953893 | 1.602205449 |
| ENSG00000143 | 107.2237007 | 2.092746193 | 0.6028743527 | 3.471280845 | 5.18E-04 | 0.01298659618 | 1.22159078 |
| ENSG00000181 | 9.294705229 | 5.714483159 | 1.646244805 | 3.471223199 | 5.18E-04 | 0.01033665786 | 1.636424459 |
| ENSG00000106 | 99.01479571 | 2.18627273 | 0.6302110455 | 3.469112046 | 5.22E-04 | 0.01305748145 | 1.22159078 |
| ENSG00000272 | 233.7373726 | 1.381685892 | 0.398580828 | 3.466513678 | 5.27E-04 | 0.03080477025 | 0.3939394882 |
| ENSG00000185 | 16.75014333 | 3.867452575 | 1.116498651 | 3.463911552 | 5.32E-04 | 0.0120485658 | 1.393563518 |
| ENSG00000283 | 43.34085554 | 2.198725752 | 0.6348311667 | 3.463481108 | 5.33E-04 | 0.0120485658 | 1.393563518 |
| ENSG00000182 | 735.3969 | -1.097886619 | 0.317058185 | -3.462729149 | 5.35E-04 | 0.01020896867 | 1.729500602 |
| ENSG00000273 | 26.93367972 | 3.223865243 | 0.93141569 | 3.461252884 | 5.38E-04 | 0.01208735666 | 1.393563518 |
| ENSG00000171 | 12.92767485 | 4.180407125 | 1.209380682 | 3.456651148 | 5.47E-04 | 0.01269149245 | 1.337480197 |
| ENSG00000129 | 274.1257224 | -1.128689697 | 0.3266559754 | -3.455285629 | 5.50E-04 | 0.03321524094 | 0.3706356727 |
| ENSG00000101 | 296.2269807 | -1.342948007 | 0.3891503728 | -3.450974484 | 5.59E-04 | 0.01292023795 | 1.327567552 |
| ENSG00000099 | 2519.047431 | 1.245772968 | 0.3610976266 | 3.449961661 | 5.61E-04 | 0.01140076265 | 1.572476891 |
| ENSG00000141 | 9.630212071 | 4.908263315 | 1.422873956 | 3.449541891 | 5.62E-04 | 0.01278241084 | 1.357988055 |
| ENSG00000064 | 24.40116801 | 3.256379929 | 0.9440875647 | 3.449235061 | 5.62E-04 | 0.01278241084 | 1.357988055 |
| ENSG00000203 | 42.30150143 | 2.300207472 | 0.6670284782 | 3.44843968 | 5.64E-04 | 0.01292837676 | 1.337480197 |
| ENSG00000226 | 47.98878056 | 2.588990334 | 0.7510089658 | 3.447349435 | 5.66E-04 | 0.01182068219 | 1.516424702 |
| ENSG00000091 | 255.660657 | 1.296973632 | 0.3763133098 | 3.446526067 | 5.68E-04 | 0.03404851745 | 0.3706356727 |
| ENSG00000082 | 145.7892033 | 1.741893868 | 0.5055282177 | 3.44569068 | 5.70E-04 | 0.01029088854 | 1.814765822 |

| | | | | | | | |
|---|---|---|---|---|---|---|---|
| ENSG00000183 | 314.1784039 | -1.7341245 | 0.5038562672 | -3.441704734 | 5.78E-04 | 0.01320074255 | 1.32213648 |
| ENSG00000102 | 154.6916421 | 1.256968109 | 0.3652359479 | 3.44152353 | 5.78E-04 | 0.01382368227 | 1.242301963 |
| ENSG00000142 | 105.7461432 | -1.492509268 | 0.4337200077 | -3.441181504 | 5.79E-04 | 0.01405788191 | 1.22159078 |
| ENSG00000178 | 28.61739159 | 3.358175245 | 0.9762474652 | 3.439881142 | 5.82E-04 | 0.01361706293 | 1.278627818 |
| ENSG00000183 | 15.04538499 | -4.716557462 | 1.371258606 | -3.43958276 | 5.83E-04 | 0.01308809555 | 1.357988055 |
| ENSG00000185 | 75.85060087 | 1.757552785 | 0.5110073196 | 3.439388669 | 5.83E-04 | 0.01255346437 | 1.449168615 |
| ENSG00000180 | 15.94178424 | 4.250750701 | 1.235967568 | 3.439208933 | 5.83E-04 | 0.01308900581 | 1.357988055 |
| ENSG00000144 | 42.73444185 | 2.094102985 | 0.6089269634 | 3.439005186 | 5.84E-04 | 0.0120785544 | 1.516424702 |
| ENSG00000164 | 354.9371628 | -1.318184195 | 0.3835241546 | -3.437030443 | 5.88E-04 | 0.01341002019 | 1.32213648 |
| ENSG00000176 | 131.3185675 | 1.526868264 | 0.4444885823 | 3.435112453 | 5.92E-04 | 0.01432448777 | 1.22159078 |
| ENSG00000197 | 82.41786886 | 10.46206044 | 3.046817299 | 3.433766915 | 5.95E-04 | 0.01273126917 | 1.449168615 |
| ENSG00000175 | 93.91911808 | 1.719868633 | 0.5011032974 | 3.43216387 | 5.99E-04 | 0.01235677666 | 1.514054369 |
| ENSG00000091 | 837.3264218 | -1.078349988 | 0.3142874469 | -3.431094684 | 6.01E-04 | 0.01166170603 | 1.634803475 |
| ENSG00000187 | 8.358891787 | -5.447285146 | 1.587817665 | -3.43067423 | 6.02E-04 | 0.01314572688 | 1.393563518 |
| ENSG00000171 | 43.91095227 | 2.080830515 | 0.6070622799 | 3.427705169 | 6.09E-04 | 0.01278241084 | 1.470104381 |
| ENSG00000166 | 222.1142987 | 1.311846567 | 0.3827211794 | 3.427682181 | 6.09E-04 | 0.03430966636 | 0.3939394882 |
| ENSG00000204 | 37.19494645 | 2.461082305 | 0.7180122907 | 3.427632559 | 6.09E-04 | 0.01348331325 | 1.357988055 |
| ENSG00000161 | 131.6693321 | 1.848078265 | 0.5391759964 | 3.42759744 | 6.09E-04 | 0.01464960838 | 1.22159078 |
| ENSG00000182 | 75.54254772 | 1.833593494 | 0.5349614705 | 3.427524401 | 6.09E-04 | 0.01292023795 | 1.449168615 |
| ENSG00000089 | 54.77523484 | 1.935041558 | 0.5645771147 | 3.427417632 | 6.09E-04 | 0.01320074255 | 1.393563518 |
| ENSG00000135 | 22.37525999 | 2.97848532 | 0.8691248213 | 3.426993738 | 6.10E-04 | 0.01200562086 | 1.605577974 |
| ENSG00000100 | 360.2805953 | 1.99277161 | 0.581559477 | 3.426599838 | 6.11E-04 | 0.0137946295 | 1.32213648 |
| ENSG00000124 | 38.83272586 | -10.35350996 | 3.021975225 | -3.426073739 | 6.12E-04 | 0.01368037799 | 1.337480197 |
| ENSG00000111 | 33.24243022 | 2.917902426 | 0.8517446523 | 3.42579483 | 6.13E-04 | 0.01353983432 | 1.357988055 |
| ENSG00000162 | 21.25771815 | -3.549156942 | 1.03730604 | -3.42151381 | 6.23E-04 | 0.01382368227 | 1.337480197 |
| ENSG00000118 | 8.533576203 | 6.078699489 | 1.777618919 | 3.419574028 | 6.27E-04 | 0.01351703571 | 1.393563518 |
| ENSG00000173 | 131.6173266 | 1.507143045 | 0.4417836858 | 3.411495476 | 6.46E-04 | 0.01531053519 | 1.22159078 |
| ENSG00000197 | 597.077237 | 1.022595887 | 0.2997621112 | 3.411358038 | 6.46E-04 | 0.01182068219 | 1.729500602 |
| ENSG00000174 | 11.80751425 | -4.641888524 | 1.360910456 | -3.410869909 | 6.48E-04 | 0.01382368227 | 1.393563518 |
| ENSG00000167 | 181.5176375 | -1.83139399 | 0.5371859732 | -3.40923643 | 6.51E-04 | 0.01908596559 | 0.9247574895 |
| ENSG00000168 | 616.4500358 | 1.18502655 | 0.3476436188 | 3.408739541 | 6.53E-04 | 0.0128580945 | 1.564848717 |
| ENSG00000123 | 679.5055766 | -1.11617399 | 0.3276523663 | -3.406579975 | 6.58E-04 | 0.01292023795 | 1.564848717 |
| ENSG00000153 | 379.5953916 | -1.090077654 | 0.3200021969 | -3.406469282 | 6.58E-04 | 0.01464636496 | 1.32213648 |

| | | | | | | | |
|---|---|---|---|---|---|---|---|
| ENSG000002592 | 80.66936722 | 1.98201736 | 0.5818869524 | 3.406189727 | 6.59E-04 | 0.01361706293 | 1.449168615 |
| ENSG000001817 | 377.7023575 | -1.05617182 | 0.3101515255 | -3.405341368 | 6.61E-04 | 0.01531053519 | 1.249985076 |
| ENSG000002137 | 84.51401669 | -1.910428795 | 0.5624789746 | -3.396444812 | 6.83E-04 | 0.01445133988 | 1.393563518 |
| ENSG000001287 | 137.8712752 | 1.700332771 | 0.5006902024 | 3.395977717 | 6.84E-04 | 0.01602815606 | 1.22159078 |
| ENSG000001305 | 97.16677658 | 1.66193836 | 0.489441751 | 3.395579467 | 6.85E-04 | 0.01511662721 | 1.318051683 |
| ENSG000001714 | 39.18164921 | 2.351724213 | 0.6926385516 | 3.395312328 | 6.86E-04 | 0.01511662721 | 1.319539707 |
| ENSG000001032 | 325.2172866 | 1.078445028 | 0.3176505237 | 3.395067686 | 6.86E-04 | 0.01511662721 | 1.32213648 |
| ENSG000001481 | 1587.652791 | 1.164591561 | 0.3433071965 | 3.392272497 | 6.93E-04 | 0.01158682805 | 1.902376403 |
| ENSG000001665 | 283.9657859 | 1.636363109 | 0.4824563769 | 3.39173278 | 6.95E-04 | 0.01518254986 | 1.327567552 |
| ENSG000002744 | 16.98648169 | 3.625678861 | 1.069303681 | 3.39069146 | 6.97E-04 | 0.01514630535 | 1.337480197 |
| ENSG000000731 | 911.0585633 | -1.269414132 | 0.3743826919 | -3.390685948 | 6.97E-04 | 0.01304408575 | 1.634803475 |
| ENSG000001428 | 2977.696774 | 2.490103639 | 0.73577577 | 3.384324056 | 7.14E-04 | 0.01341002019 | 1.602205449 |
| ENSG000001547 | 82.99625678 | 1.70448411 | 0.5039348545 | 3.382350108 | 7.19E-04 | 0.01506653086 | 1.393563518 |
| ENSG000002143 | 2361.352395 | 1.097249326 | 0.3247469563 | 3.37878248 | 7.28E-04 | 0.01379894928 | 1.572476891 |
| ENSG000001302 | 63.35608837 | 2.951204199 | 0.8735835023 | 3.378273732 | 7.29E-04 | 0.01424419696 | 1.516424702 |
| ENSG000001624 | 15.86600598 | 4.210578914 | 1.24655618 | 3.377769075 | 7.31E-04 | 0.01571906733 | 1.337480197 |
| ENSG000000680 | 261.9058591 | -1.130527973 | 0.3347026257 | -3.377708706 | 7.31E-04 | 0.0391745482 | 0.3939394882 |
| ENSG000002042 | 113.6536495 | 1.592840019 | 0.4716556208 | 3.377125066 | 7.32E-04 | 0.01788357824 | 1.133466638 |
| ENSG000001627 | 243.9187843 | 1.525189007 | 0.4517774571 | 3.375974128 | 7.36E-04 | 0.01672956443 | 1.242301963 |
| ENSG000001330 | 60.46264255 | 2.275074011 | 0.6749146595 | 3.370906202 | 7.49E-04 | 0.01492065348 | 1.470104381 |
| ENSG000002607 | 26.06169956 | -2.953993412 | 0.8777484799 | -3.365421279 | 7.64E-04 | 0.01626479593 | 1.337480197 |
| ENSG000001712 | 47.88823317 | -2.624629135 | 0.7799054055 | -3.365317276 | 7.65E-04 | 0.01626479593 | 1.337480197 |
| ENSG000001086 | 210.3023514 | 1.5346337 | 0.456129781 | 3.364467228 | 7.67E-04 | 0.01775416275 | 1.198705341 |
| ENSG000002248 | 12.92214158 | 4.652043937 | 1.383396571 | 3.362769601 | 7.72E-04 | 0.01622731838 | 1.357988055 |
| ENSG000001877 | 112.2789354 | -1.987710384 | 0.5911525209 | -3.362432391 | 7.73E-04 | 0.01743877011 | 1.242301963 |
| ENSG000001325 | 12.33298477 | 5.218832589 | 1.553354552 | 3.359717575 | 7.80E-04 | 0.01602815606 | 1.393563518 |
| ENSG000001295 | 13.44570474 | 5.411333888 | 1.611321723 | 3.35831995 | 7.84E-04 | 0.01664146531 | 1.337480197 |
| ENSG000001451 | 2202.239969 | -1.389726017 | 0.413987681 | -3.356926017 | 7.88E-04 | 0.01449297325 | 1.602205449 |
| ENSG000001530 | 26.35853141 | 3.444015679 | 1.026813302 | 3.354081675 | 7.96E-04 | 0.01676447555 | 1.337480197 |
| ENSG000001117 | 49.78459678 | 2.03809642 | 0.607674179 | 3.353929607 | 7.97E-04 | 0.01676447555 | 1.337480197 |
| ENSG000000631 | 12.45503429 | 4.100685868 | 1.22269494 | 3.353809469 | 7.97E-04 | 0.0179210008 | 1.227924265 |
| ENSG000000672 | 2960.244209 | -1.025298715 | 0.3057371025 | -3.353530553 | 7.98E-04 | 0.01465990397 | 1.597450288 |
| ENSG000001859 | 118.8631217 | 1.908207613 | 0.5692906906 | 3.351903772 | 8.03E-04 | 0.01807397281 | 1.22159078 |

| | | | | | | | |
|---|---|---|---|---|---|---|---|
| ENSG00000160 | 183.9221621 | -1.319300694 | 0.3940740832 | -3.347849428 | 8.14E-04 | 0.01805606157 | 1.242301963 |
| ENSG00000188 | 72.98698836 | 1.98081429 | 0.5916777258 | 3.347792563 | 8.15E-04 | 0.01607208751 | 1.449168615 |
| ENSG00000196 | 157.9355984 | 1.67806759 | 0.5021578584 | 3.341713292 | 8.33E-04 | 0.01857486217 | 1.22159078 |
| ENSG00000164 | 73.18665924 | -1.917501209 | 0.5740757207 | -3.340153816 | 8.37E-04 | 0.01641978532 | 1.449168615 |
| ENSG00000169 | 55.28420143 | 1.914407102 | 0.5733085913 | 3.339226257 | 8.40E-04 | 0.01589952832 | 1.516424702 |
| ENSG00000186 | 934.2565098 | 1.343706382 | 0.4025317465 | 3.33813766 | 8.43E-04 | 0.01506653086 | 1.634803475 |
| ENSG00000161 | 10.02366366 | 5.698576716 | 1.708061824 | 3.336282467 | 8.49E-04 | 0.01751124023 | 1.357988055 |
| ENSG00000177 | 78.47852901 | -1.520438478 | 0.455787671 | -3.335848192 | 8.50E-04 | 0.01751751112 | 1.357988055 |
| ENSG00000237 | 64.97680261 | 2.038230174 | 0.6110429206 | 3.335657946 | 8.51E-04 | 0.01771302433 | 1.337480197 |
| ENSG00000010 | 56.92710567 | 2.505294344 | 0.7514886009 | 3.333775577 | 8.57E-04 | 0.01760618332 | 1.357988055 |
| ENSG00000131 | 1723.2571 | -1.195923377 | 0.3587366167 | -3.333708691 | 8.57E-04 | 0.01547206226 | 1.597450288 |
| ENSG00000165 | 159.4649635 | 1.800718581 | 0.5406435434 | 3.330694694 | 8.66E-04 | 0.01919090377 | 1.22159078 |
| ENSG00000163 | 43.07050676 | 2.319535042 | 0.6967092879 | 3.329272456 | 8.71E-04 | 0.01672956443 | 1.470104381 |
| ENSG00000105 | 1313.810701 | -1.049542618 | 0.3156310302 | -3.325220013 | 8.83E-04 | 0.01556717252 | 1.634803475 |
| ENSG00000185 | 12.43230368 | 3.534329286 | 1.063038642 | 3.324742062 | 8.85E-04 | 0.01770111014 | 1.393563518 |
| ENSG00000214 | 24.49050507 | 4.001817777 | 1.204046358 | 3.323640947 | 8.89E-04 | 0.01772862757 | 1.393563518 |
| ENSG00000115 | 33.38593674 | 2.509346671 | 0.755242014 | 3.32257293 | 8.92E-04 | 0.01775416275 | 1.393563518 |
| ENSG00000088 | 237.4263315 | -1.066607829 | 0.3210360616 | -3.322392581 | 8.92E-04 | 0.01837501034 | 1.327567552 |
| ENSG00000128 | 42.03462758 | 2.055515742 | 0.6189954805 | 3.320728191 | 8.98E-04 | 0.01785061885 | 1.393563518 |
| ENSG00000171 | 72.00751247 | 2.260902588 | 0.6809098279 | 3.320414092 | 8.99E-04 | 0.01674796438 | 1.514054369 |
| ENSG00000135 | 25.91572697 | 3.955330227 | 1.191311379 | 3.320148113 | 9.00E-04 | 0.01837501034 | 1.337480197 |
| ENSG00000082 | 82.0838793 | 1.783071574 | 0.5371004701 | 3.319810117 | 9.01E-04 | 0.01822653033 | 1.357988055 |
| ENSG00000166 | 14.27634896 | 5.073654632 | 1.528574334 | 3.319207003 | 9.03E-04 | 0.01823157479 | 1.357988055 |
| ENSG00000223 | 11.89596601 | 4.949706538 | 1.491955868 | 3.317595811 | 9.08E-04 | 0.01830670597 | 1.357988055 |
| ENSG00000170 | 958.2729623 | -1.230221858 | 0.3710047631 | -3.315919309 | 9.13E-04 | 0.01626087384 | 1.602205449 |
| ENSG00000157 | 20.11531913 | 2.875761635 | 0.8676158805 | 3.314556244 | 9.18E-04 | 0.01868066264 | 1.337480197 |
| ENSG00000155 | 122.2618018 | -1.331359693 | 0.4017881446 | -3.313586305 | 9.21E-04 | 0.02154788394 | 1.130447911 |
| ENSG00000178 | 568.7962331 | -1.592113712 | 0.480579815 | -3.312901753 | 9.23E-04 | 0.01664596499 | 1.572476891 |
| ENSG00000126 | 144.9876394 | 1.357448632 | 0.4101206968 | 3.309875954 | 9.33E-04 | 0.0252285786 | 0.9247574895 |
| ENSG00000132 | 2033.995615 | 1.028318602 | 0.3108948195 | 3.307609319 | 9.41E-04 | 0.01667818496 | 1.597450288 |
| ENSG00000159 | 167.3594812 | -1.248834295 | 0.3776182962 | -3.307133969 | 9.43E-04 | 0.02093773246 | 1.198705341 |
| ENSG00000088 | 38.88826898 | 2.72514993 | 0.8241097548 | 3.306780334 | 9.44E-04 | 0.01930985312 | 1.319539707 |
| ENSG00000115 | 27.40097215 | -2.776638685 | 0.8396948858 | -3.306723349 | 9.44E-04 | 0.01983309789 | 1.278627818 |

| | | | | | | | |
|---|---|---|---|---|---|---|---|
| ENSG00000075[illegible] | 8.775614945 | 5.319575312 | 1.608973144 | 3.30619273 | 9.46E-04 | 0.0189079684 | 1.357988055 |
| ENSG00000175[illegible] | 131.3069859 | -4.960469811 | 1.502224413 | -3.302083077 | 9.60E-04 | 0.02093773246 | 1.22159078 |
| ENSG00000129[illegible] | 35.72632695 | 2.449615538 | 0.7418901551 | 3.301857454 | 9.60E-04 | 0.01873866107 | 1.393563518 |
| ENSG00000140[illegible] | 214.8369399 | 1.112562299 | 0.337186832 | 3.299542548 | 9.68E-04 | 0.02141422678 | 1.198705341 |
| ENSG00000082[illegible] | 24.49088034 | 3.179603289 | 0.9642006656 | 3.297657222 | 9.75E-04 | 0.01983309789 | 1.319539707 |
| ENSG00000185[illegible] | 14.99356798 | 4.539043615 | 1.376770207 | 3.296878153 | 9.78E-04 | 0.01711586091 | 1.605577974 |
| ENSG00000143[illegible] | 11.498156 | 5.207919792 | 1.580150062 | 3.295838741 | 9.81E-04 | 0.02123063477 | 1.227924265 |
| ENSG00000127[illegible] | 45.11181133 | -2.862555299 | 0.8689887032 | -3.294122568 | 9.87E-04 | 0.01958340244 | 1.357988055 |
| ENSG00000189[illegible] | 1737.492243 | -1.038085313 | 0.3152496512 | -3.292899165 | 9.92E-04 | 0.01742718321 | 1.597450288 |
| ENSG00000101[illegible] | 35.53850544 | 2.454795225 | 0.746053421 | 3.290374597 | 0.001000540879 | 0.02005749131 | 1.337480197 |
| ENSG00000136[illegible] | 33.98571514 | 2.19531875 | 0.6674467584 | 3.289129392 | 0.001004978111 | 0.02093773246 | 1.278627818 |
| ENSG00000133[illegible] | 184.6624103 | 1.281613689 | 0.3903940277 | 3.282872171 | 0.001027552489 | 0.02182474456 | 1.242301963 |
| ENSG00000154[illegible] | 823.6532308 | -1.060313321 | 0.3230277054 | -3.282422231 | 0.001029193704 | 0.01804818391 | 1.572476891 |
| ENSG00000008[illegible] | 272.806124 | 1.055649773 | 0.3219152902 | 3.279278135 | 0.00104073008 | 0.02093773246 | 1.327567552 |
| ENSG00000262[illegible] | 242.4042286 | 1.259383084 | 0.3841777217 | 3.278126276 | 0.001044986369 | 0.05087013435 | 0.3939394882 |
| ENSG00000170[illegible] | 275.2552485 | -1.792898444 | 0.5469324623 | -3.278098425 | 0.001045089481 | 0.0530317578 | 0.3706356727 |
| ENSG00000196[illegible] | 160.8440667 | -1.357008104 | 0.4141395182 | -3.276693106 | 0.001050304632 | 0.02750200926 | 0.9247574895 |
| ENSG00000127[illegible] | 121.2074311 | -1.466136068 | 0.447950384 | -3.272987635 | 0.00106417133 | 0.02141422678 | 1.318051683 |
| ENSG00000123[illegible] | 232.3475822 | -1.478743803 | 0.4520431425 | -3.271244853 | 0.001070751596 | 0.0540279661 | 0.3706356727 |
| ENSG00000158[illegible] | 158.531686 | 1.454452892 | 0.4449704659 | 3.268650401 | 0.001080617288 | 0.02305311115 | 1.22159078 |
| ENSG00000163[illegible] | 19.44337737 | 3.652216648 | 1.117580075 | 3.267968649 | 0.00108322364 | 0.01840752361 | 1.605577974 |
| ENSG00000277[illegible] | 1666.155564 | 1.235994809 | 0.3783371976 | 3.266913262 | 0.001087269873 | 0.01823157479 | 1.634803475 |
| ENSG00000106[illegible] | 125.4475113 | 1.351077979 | 0.4138223536 | 3.264874329 | 0.001095126531 | 0.02320654305 | 1.22159078 |
| ENSG00000186[illegible] | 215.9507842 | -1.797740688 | 0.5517856798 | -3.258041579 | 0.001121839677 | 0.05566532369 | 0.3706356727 |
| ENSG00000119[illegible] | 694.5841107 | -1.058958849 | 0.3250968581 | -3.257364144 | 0.00112452073 | 0.01930985312 | 1.572476891 |
| ENSG00000131[illegible] | 188.0828931 | 1.226963595 | 0.3769197228 | 3.255238504 | 0.001132971769 | 0.02417079851 | 1.198705341 |
| ENSG00000110[illegible] | 64.50960553 | 2.122172486 | 0.6519519199 | 3.255105816 | 0.001133501246 | 0.02153520207 | 1.393563518 |
| ENSG00000123[illegible] | 226.9282688 | 1.144025946 | 0.3515853123 | 3.253907105 | 0.00113829496 | 0.02249638291 | 1.327567552 |
| ENSG00000156[illegible] | 111.3375241 | -1.570593045 | 0.4830326373 | -3.251525722 | 0.001147873884 | 0.02527231213 | 1.130447911 |
| ENSG00000163[illegible] | 240.0588355 | 1.173201422 | 0.3608722817 | 3.251015614 | 0.001149935421 | 0.05656714806 | 0.3706356727 |
| ENSG00000116[illegible] | 250.8520054 | 1.165032992 | 0.3583907908 | 3.250733618 | 0.001151076539 | 0.05657890589 | 0.3706356727 |
| ENSG00000204[illegible] | 99.10064081 | 1.39652783 | 0.4296588502 | 3.250317851 | 0.001152760883 | 0.0239135537 | 1.242301963 |
| ENSG00000048[illegible] | 34.39689123 | 2.563838433 | 0.7891063163 | 3.249040566 | 0.001157949635 | 0.0218999211 | 1.393563518 |

| | | | | | | | |
|---|---|---|---|---|---|---|---|
| ENSG000000719 | 156.112791 | 1.604284962 | 0.4939379981 | 3.247948059 | 0.001162404881 | 0.02455559922 | 1.198705341 |
| ENSG000001455 | 39.39882836 | 3.334400279 | 1.026698719 | 3.247691088 | 0.001163455107 | 0.02300370006 | 1.319539707 |
| ENSG000001703 | 24.89505829 | 2.566014813 | 0.7907604434 | 3.244996427 | 0.001174520975 | 0.02266675971 | 1.357988055 |
| ENSG000002117 | 155.4588119 | 1.564977966 | 0.4828975806 | 3.24080722 | 0.0011919176 | 0.02437508846 | 1.242301963 |
| ENSG000001571 | 22.3972002 | 3.2556789 | 1.0046689 | 3.2405491 | 0.001192997247 | 0.02248628113 | 1.393563518 |
| ENSG000001227 | 1182.41275 | -2.792100495 | 0.8618020487 | -3.239839705 | 0.001195969119 | 0.01968289178 | 1.634803475 |
| ENSG000001493 | 114.301343 | -1.682405522 | 0.5195045831 | -3.238480615 | 0.001201681869 | 0.02610890727 | 1.130447911 |
| ENSG000001095 | 239.4841478 | 1.370573189 | 0.4234395754 | 3.236762146 | 0.00120894131 | 0.05603966512 | 0.3939394882 |
| ENSG000001182 | 238.3834688 | 1.272882658 | 0.3934531481 | 3.235156877 | 0.001215759124 | 0.02526979315 | 1.198705341 |
| ENSG000001373 | 169.8864433 | 1.268113367 | 0.3923319559 | 3.232245928 | 0.001228213031 | 0.03062905127 | 0.9247574895 |
| ENSG000000044 | 393.2459079 | -1.062175619 | 0.3287069098 | -3.231375998 | 0.001231957651 | 0.02396068852 | 1.32213648 |
| ENSG000001592 | 33.04447582 | 2.355909436 | 0.7295504884 | 3.229261681 | 0.001241102747 | 0.0239135537 | 1.337480197 |
| ENSG000002578 | 17.17930113 | 4.584938486 | 1.419944275 | 3.228956634 | 0.001242427333 | 0.0239135537 | 1.337480197 |
| ENSG000001987 | 26.78160595 | 2.778818614 | 0.8605980046 | 3.228939178 | 0.001242503171 | 0.0231836308 | 1.393563518 |
| ENSG000001496 | 649.2087519 | -1.096751407 | 0.3399951022 | -3.2257859 | 0.001256272943 | 0.01958340244 | 1.729500602 |
| ENSG000001274 | 214.9455806 | -1.324972983 | 0.4107541563 | -3.225708037 | 0.001256614733 | 0.02522942176 | 1.242301963 |
| ENSG000000071 | 218.975027 | -1.451322612 | 0.4500910163 | -3.224509177 | 0.00126188812 | 0.05986740222 | 0.3706356727 |
| ENSG000001300 | 9.899277889 | 4.70105403 | 1.458459376 | 3.223301319 | 0.001267221744 | 0.02552121903 | 1.227924265 |
| ENSG000002794 | 45.64409095 | -1.961848563 | 0.6089540851 | -3.221669106 | 0.001274462284 | 0.0243147318 | 1.337480197 |
| ENSG000000357 | 12.77972644 | 4.374851918 | 1.358626037 | 3.220055998 | 0.001281655585 | 0.02437508846 | 1.337480197 |
| ENSG000001855 | 69.94393822 | 1.549036227 | 0.4810727784 | 3.219962334 | 0.001282074406 | 0.0241701678 | 1.357988055 |
| ENSG000001718 | 55.53983626 | 2.577996681 | 0.8008271271 | 3.219167525 | 0.001285633521 | 0.02439397568 | 1.337480197 |
| ENSG000001643 | 208.3477281 | -1.96984147 | 0.6127544993 | -3.214731956 | 0.001305663792 | 0.02487754117 | 1.327567552 |
| ENSG000001047 | 298.2068844 | -1.112789961 | 0.3463984327 | -3.212456686 | 0.001316049915 | 0.0249488842 | 1.327567552 |
| ENSG000001027 | 148.8608927 | 1.244741504 | 0.3875690942 | 3.211663476 | 0.001319688635 | 0.02679809349 | 1.198705341 |
| ENSG000000182 | 26.70860364 | 3.362870148 | 1.047192639 | 3.211319505 | 0.001321269427 | 0.02493403866 | 1.337480197 |
| ENSG000002559 | 11.47043794 | 4.569663412 | 1.42322586 | 3.210778794 | 0.001323757914 | 0.0249488842 | 1.337480197 |
| ENSG000002659 | 2275.420064 | 1.006893246 | 0.3136296218 | 3.210453273 | 0.001325258131 | 0.0221621645 | 1.572476891 |
| ENSG000001287 | 72.40819483 | 4.922831821 | 1.534580712 | 3.207932814 | 0.001336927268 | 0.02502772276 | 1.337480197 |
| ENSG000000701 | 65.71811878 | 1.970347714 | 0.6147351322 | 3.205197833 | 0.001349696783 | 0.0231836308 | 1.514054369 |
| ENSG000000808 | 67.16853556 | 1.697718353 | 0.5297535258 | 3.204732522 | 0.001351880467 | 0.0231859346 | 1.514054369 |
| ENSG000001704 | 203.677553 | 1.207441313 | 0.3767709566 | 3.204709099 | 0.001351990478 | 0.0273395158 | 1.198705341 |
| ENSG000001228 | 310.6483409 | -1.329106087 | 0.4148216989 | -3.204041858 | 0.001355127736 | 0.0266637224 | 1.242301963 |

| | | | | | | | |
|---|---|---|---|---|---|---|---|
| ENSG000001961 | 609.4171686 | -1.141172636 | 0.3565745063 | -3.200376403 | 0.001372482199 | 0.0229082856 | 1.564848717 |
| ENSG000001604 | 50.73352993 | 2.16672458 | 0.6771478343 | 3.199780713 | 0.00137532184 | 0.02493403866 | 1.393563518 |
| ENSG000002856 | 27.90388538 | 2.685964939 | 0.8400824387 | 3.197263525 | 0.00138738115 | 0.0259185002 | 1.319539707 |
| ENSG000001173 | 186.1176618 | -1.792557003 | 0.5607431797 | -3.196752217 | 0.001389842603 | 0.02789788959 | 1.198705341 |
| ENSG000001842 | 45.4009484 | 2.230152343 | 0.6976910662 | 3.196475419 | 0.001391176797 | 0.0253937239 | 1.357988055 |
| ENSG000001326 | 10.60323709 | 5.107278783 | 1.59781589 | 3.196412562 | 0.001391479935 | 0.0253937239 | 1.357988055 |
| ENSG000002240 | 37.35547432 | -2.447286345 | 0.7657528721 | -3.195921862 | 0.001393848539 | 0.02502772276 | 1.393563518 |
| ENSG000001802 | 20.13198847 | 3.277775569 | 1.026644822 | 3.192706473 | 0.001409461395 | 0.02522942176 | 1.393563518 |
| ENSG000002462 | 21.07391402 | 3.159680221 | 0.989993489 | 3.191617173 | 0.001414787131 | 0.02398949177 | 1.514850267 |
| ENSG000001664 | 95.94840264 | 1.922218019 | 0.6025745915 | 3.190008417 | 0.001422686502 | 0.02643983812 | 1.318051683 |
| ENSG000000260 | 1262.052214 | 1.023384642 | 0.3209145719 | 3.188962832 | 0.001427842347 | 0.0231859346 | 1.597450288 |
| ENSG000000791 | 31.00775808 | 2.647924346 | 0.8306102965 | 3.187926224 | 0.001432970931 | 0.02657293623 | 1.319539707 |
| ENSG000000707 | 6129.544258 | -1.011341537 | 0.317251114 | -3.18782659 | 0.001433464759 | 0.0231859346 | 1.602205449 |
| ENSG000002614 | 11.19816437 | 5.050629741 | 1.58505502 | 3.186406577 | 0.001440520029 | 0.02553589847 | 1.393563518 |
| ENSG000002760 | 197.4519205 | -1.968589554 | 0.6181881662 | -3.184450402 | 0.001450291617 | 0.06629827703 | 0.3706356727 |
| ENSG000000696 | 15.47554033 | 3.870309893 | 1.215954729 | 3.182939135 | 0.001457882569 | 0.02581624689 | 1.393563518 |
| ENSG000002411 | 21.20038161 | 3.862307159 | 1.214479103 | 3.180217056 | 0.001471647758 | 0.02600478958 | 1.393563518 |
| ENSG000001524 | 54.95004952 | -2.356925588 | 0.7411479316 | -3.180101418 | 0.001472235168 | 0.02679809349 | 1.337480197 |
| ENSG000001073 | 170.9877331 | 1.337670277 | 0.4208264608 | 3.178674351 | 0.001479502088 | 0.02869915185 | 1.22159078 |
| ENSG000002270 | 7.129575913 | -5.268596883 | 1.658059775 | -3.177567518 | 0.001485161053 | 0.02667675097 | 1.357988055 |
| ENSG000001542 | 80.76194104 | 1.565974388 | 0.492927452 | 3.176886135 | 0.001488654707 | 0.02706430977 | 1.337480197 |
| ENSG000002052 | 2143.136986 | -1.116217295 | 0.3516846462 | -3.173915345 | 0.001503975488 | 0.02412983502 | 1.597450288 |
| ENSG000001362 | 671.3545919 | -1.000391183 | 0.3152141148 | -3.173687776 | 0.001505155063 | 0.02177923681 | 1.825593782 |
| ENSG000001167 | 134.6757967 | 1.537055711 | 0.4846041478 | 3.171775806 | 0.00151509923 | 0.02917380914 | 1.22159078 |
| ENSG000001051 | 118.7809298 | -1.670518855 | 0.5270662429 | -3.169466604 | 0.001527190074 | 0.02901201706 | 1.242301963 |
| ENSG000001830 | 384.779665 | 1.21135641 | 0.3822312026 | 3.169171961 | 0.001528739183 | 0.02785338799 | 1.32213648 |
| ENSG000000474 | 16.11704618 | 4.006688106 | 1.264612848 | 3.168312036 | 0.001533268591 | 0.0243411018 | 1.605577974 |
| ENSG000000260 | 497.8589789 | -1.208265127 | 0.3814229471 | -3.167782999 | 0.001536061278 | 0.02437508846 | 1.602205449 |
| ENSG000002045 | 25.87101795 | -3.430199239 | 1.082894577 | -3.167620664 | 0.001536919152 | 0.02851208856 | 1.278627818 |
| ENSG000002798 | 34.33822198 | -2.917644522 | 0.9211519757 | -3.167386706 | 0.001538156305 | 0.02789788959 | 1.319539707 |
| ENSG000002706 | 10.57798443 | 4.266782001 | 1.348207021 | 3.16478251 | 0.00155198913 | 0.02422699525 | 1.636424459 |
| ENSG000002240 | 588.4741876 | 1.064477773 | 0.3364985289 | 3.163395029 | 0.00155940577 | 0.0249488842 | 1.572476891 |
| ENSG000002731 | 335.1134108 | -1.099495586 | 0.3478511411 | -3.160822134 | 0.001573245351 | 0.02871828269 | 1.29680518 |

| | | | | | | | |
|---|---|---|---|---|---|---|---|
| ENSG00000111 | 299.7362821 | -1.260659135 | 0.3988968213 | -3.160363953 | 0.001575721736 | 0.02873422612 | 1.29680518 |
| ENSG00000205 | 68.37486594 | -1.647429037 | 0.5215162092 | -3.15892202 | 0.001583538551 | 0.02667675097 | 1.449168615 |
| ENSG00000184 | 281.7712063 | -1.42792574 | 0.452082024 | -3.158554563 | 0.001585536259 | 0.02847511138 | 1.32213648 |
| ENSG00000151 | 58.32402429 | 1.877675016 | 0.5945320497 | 3.158240195 | 0.001587247189 | 0.02755133478 | 1.393563518 |
| ENSG00000185 | 161.2466206 | 1.21302823 | 0.3841105328 | 3.158018661 | 0.001588453892 | 0.03717840104 | 0.9247574895 |
| ENSG00000166 | 421.8139904 | 1.183179649 | 0.3748909077 | 3.156063871 | 0.001599138386 | 0.02904306138 | 1.29680518 |
| ENSG00000185 | 34.45031288 | -3.260200689 | 1.033314909 | -3.155089181 | 0.001604490526 | 0.02940556778 | 1.278627818 |
| ENSG00000165 | 16.0950378 | 3.405813541 | 1.079730916 | 3.154316961 | 0.001608742583 | 0.02610890727 | 1.514850267 |
| ENSG00000180 | 226.055836 | 1.400565384 | 0.4441042402 | 3.153686133 | 0.001612223791 | 0.07108118798 | 0.3706356727 |
| ENSG00000221 | 15.57127733 | 3.399742065 | 1.07854342 | 3.152160591 | 0.001620671129 | 0.02623403442 | 1.514850267 |
| ENSG00000174 | 108.5366492 | 1.446002916 | 0.4587878319 | 3.151790034 | 0.001622729146 | 0.03237370565 | 1.133466638 |
| ENSG00000106 | 14.74211616 | 3.329837593 | 1.056525077 | 3.151688175 | 0.001623295277 | 0.02842958701 | 1.357988055 |
| ENSG00000274 | 10.07398734 | 5.654983878 | 1.794910636 | 3.150565696 | 0.001629546037 | 0.02499815285 | 1.636424459 |
| ENSG00000273 | 8.008206869 | 4.869167274 | 1.546602478 | 3.148299154 | 0.001642235335 | 0.03080477025 | 1.227924265 |
| ENSG00000287 | 18.44113559 | 3.3306513 | 1.057967197 | 3.148161217 | 0.001643010505 | 0.0253937239 | 1.605577974 |
| ENSG00000157 | 260.7924002 | -1.067596815 | 0.3394056503 | -3.145489222 | 0.00165809304 | 0.06958927194 | 0.3939394882 |
| ENSG00000115 | 2513.467376 | -5.231408918 | 1.663652232 | -3.144532744 | 0.00166352294 | 0.02286540069 | 1.902376403 |
| ENSG00000251 | 8.329312088 | -5.510982406 | 1.75358566 | -3.142693586 | 0.001674009776 | 0.0253937239 | 1.636424459 |
| ENSG00000205 | 8.037801337 | 4.962669647 | 1.579279522 | 3.142363069 | 0.001675900814 | 0.02939243255 | 1.337480197 |
| ENSG00000154 | 28.34074366 | 2.337675126 | 0.7440326241 | 3.141898689 | 0.001678561062 | 0.02963004923 | 1.319539707 |
| ENSG00000020 | 92.31281764 | 1.471115395 | 0.4685109923 | 3.139980533 | 0.001689590619 | 0.02789788959 | 1.449168615 |
| ENSG00000198 | 61.0222259 | 1.662698043 | 0.5298618486 | 3.137984076 | 0.001701141187 | 0.02963004923 | 1.337480197 |
| ENSG00000006 | 53.06412964 | -1.818868121 | 0.5798463106 | -3.136810717 | 0.001707963536 | 0.02941335019 | 1.357988055 |
| ENSG00000123 | 127.0103418 | -1.734021994 | 0.5528240388 | -3.136661708 | 0.001708831725 | 0.03141033601 | 1.242301963 |
| ENSG00000278 | 43.42582554 | -9.556146214 | 3.047191625 | -3.136050302 | 0.001712398302 | 0.02976328638 | 1.337480197 |
| ENSG00000270 | 7.843552163 | 4.70849566 | 1.502573989 | 3.133619839 | 0.001726643961 | 0.02997985924 | 1.337480197 |
| ENSG00000121 | 106.7941526 | 1.285077481 | 0.4106691334 | 3.129228316 | 0.001752660667 | 0.03238046662 | 1.22159078 |
| ENSG00000153 | 37.35216236 | 5.871930912 | 1.876733151 | 3.128804385 | 0.001755191141 | 0.03140760315 | 1.278627818 |
| ENSG00000164 | 15.63592479 | 3.272320795 | 1.046245602 | 3.127679379 | 0.001761922685 | 0.02789788959 | 1.514850267 |
| ENSG00000078 | 10.50378026 | 5.09750977 | 1.629844032 | 3.127605875 | 0.001762363321 | 0.03044796962 | 1.337480197 |
| ENSG00000113 | 47.34631498 | -2.438545658 | 0.779863597 | -3.126887403 | 0.001766675745 | 0.0301815409 | 1.357988055 |
| ENSG00000103 | 79.47073972 | 1.881972984 | 0.6023938727 | 3.124156916 | 0.001783153349 | 0.02813731263 | 1.514054369 |
| ENSG00000204 | 788.9197578 | 1.224220431 | 0.3921793303 | 3.121583255 | 0.001798813778 | 0.0273238302 | 1.597450288 |

| | | | | | | | |
|---|---|---|---|---|---|---|---|
| ENSG000001402 | 164.2822168 | 1.486195352 | 0.4763717341 | 3.119822705 | 0.001809599229 | 0.02500519872 | 1.814765822 |
| ENSG000001353 | 52.73143 | 2.25456061 | 0.7226944365 | 3.119659563 | 0.001810601666 | 0.02842958701 | 1.516424702 |
| ENSG000002614 | 15.19373681 | -4.133581154 | 1.325099066 | -3.119450658 | 0.001811886044 | 0.02842958701 | 1.514850267 |
| ENSG000001729 | 172.9577093 | -1.711790534 | 0.5491106742 | -3.117387103 | 0.001824618164 | 0.04102518673 | 0.9247574895 |
| ENSG000000930 | 51.37038331 | -2.377672128 | 0.7627160324 | -3.11737531 | 0.001824691157 | 0.02917380914 | 1.470104381 |
| ENSG000001657 | 712.7098335 | 1.088254455 | 0.3491596516 | 3.11678182 | 0.001828368314 | 0.02789788959 | 1.572476891 |
| ENSG000002835 | 8.584608928 | -4.893558139 | 1.570317836 | -3.116285141 | 0.00183145088 | 0.02718539258 | 1.636424459 |
| ENSG000002734 | 21.76970024 | -2.61309764 | 0.8385848596 | -3.116080155 | 0.001832724488 | 0.02758283761 | 1.605577974 |
| ENSG000001435 | 194.2511762 | -1.441367221 | 0.4626299364 | -3.115594361 | 0.001835746048 | 0.03346066675 | 1.22159078 |
| ENSG000001872 | 117.7374851 | 1.419887885 | 0.4558125397 | 3.115069818 | 0.001839013757 | 0.03318388912 | 1.242301963 |
| ENSG000001677 | 20.1696939 | -3.151288364 | 1.012292217 | -3.113022418 | 0.001851819511 | 0.02781239914 | 1.605577974 |
| ENSG000001321 | 105.4330366 | -1.456468221 | 0.4679652155 | -3.112342912 | 0.001856087649 | 0.03192700015 | 1.318051683 |
| ENSG000002331 | 25.33977873 | 2.670452785 | 0.8584861913 | 3.110653161 | 0.001866740593 | 0.03174636086 | 1.337480197 |
| ENSG000001771 | 15.69506396 | 3.273406787 | 1.052347844 | 3.110574898 | 0.001867235354 | 0.02904306138 | 1.514850267 |
| ENSG000001679 | 334.6718296 | -1.048433687 | 0.3371054619 | -3.110105904 | 0.001870202765 | 0.03192700015 | 1.327567552 |
| ENSG000001623 | 316.1302319 | -1.27838572 | 0.4110537565 | -3.11002077 | 0.001870741888 | 0.03192700015 | 1.327567552 |
| ENSG000002616 | 9.126808295 | 5.215843979 | 1.677623886 | 3.109066354 | 0.001876795642 | 0.02768490073 | 1.636424459 |
| ENSG000002154 | 72.35493753 | -1.732526105 | 0.5573035636 | -3.108765525 | 0.00187870749 | 0.03152907856 | 1.357988055 |
| ENSG000001018 | 66.16933046 | -2.120748397 | 0.6824096356 | -3.10773513 | 0.001885269498 | 0.03104455656 | 1.393563518 |
| ENSG000001266 | 33.89122244 | 2.782624605 | 0.8956736117 | 3.106739518 | 0.001891629983 | 0.03201301797 | 1.337480197 |
| ENSG000001707 | 67.62614731 | -2.067566259 | 0.6656637177 | -3.106022161 | 0.00189622504 | 0.03029601297 | 1.449168615 |
| ENSG000001280 | 892.7605099 | 1.026150733 | 0.3305388877 | 3.104478084 | 0.001906150487 | 0.02839410974 | 1.602205449 |
| ENSG000002324 | 92.94082169 | -2.198306194 | 0.7085096316 | -3.102718857 | 0.001917517072 | 0.03238046662 | 1.337480197 |
| ENSG000002736 | 7.154924887 | 5.885508117 | 1.897845797 | 3.10115191 | 0.001927693689 | 0.03152907856 | 1.393563518 |
| ENSG000002061 | 131.1433127 | 1.891747068 | 0.6100728739 | 3.100854257 | 0.001929632403 | 0.03291317946 | 1.318051683 |
| ENSG000001985 | 53.10852814 | 1.960168952 | 0.6323948059 | 3.099596857 | 0.001937842065 | 0.03166023267 | 1.393563518 |
| ENSG000002397 | 185.3419704 | -1.285187119 | 0.4152560046 | -3.094927238 | 0.001968611847 | 0.03494946608 | 1.242301963 |
| ENSG000001311 | 134.4943639 | 1.570366836 | 0.5074131081 | 3.094848774 | 0.001969132683 | 0.03541700232 | 1.22159078 |
| ENSG000001751 | 12.14098668 | 4.129614044 | 1.335923561 | 3.091205339 | 0.001993457262 | 0.02877823152 | 1.636424459 |
| ENSG000001292 | 41.44885414 | -2.028259009 | 0.6565213069 | -3.089403174 | 0.002005590683 | 0.03342095594 | 1.337480197 |
| ENSG000001683 | 102.5840357 | -1.607652091 | 0.5204207362 | -3.089139189 | 0.002007373693 | 0.0380799723 | 1.130447911 |
| ENSG000001746 | 105.1574089 | 1.386081462 | 0.4487527861 | 3.088741742 | 0.002010060876 | 0.03598308819 | 1.22159078 |
| ENSG000001548 | 54.41636718 | 1.688097579 | 0.546784441 | 3.087318241 | 0.002019712434 | 0.03324363332 | 1.357988055 |

| | | | | | | | |
|---|---|---|---|---|---|---|---|
| ENSG00000102 | 49.06400032 | 2.073837867 | 0.6718815323 | 3.086612398 | 0.002024513917 | 0.0332906535 | 1.357988055 |
| ENSG00000150 | 96.32928205 | 1.413635117 | 0.4580616858 | 3.086123902 | 0.002027843026 | 0.03839927896 | 1.130447911 |
| ENSG00000187 | 11.7157346 | 4.07745844 | 1.321358209 | 3.085808535 | 0.002029994933 | 0.03611852264 | 1.227924265 |
| ENSG00000128 | 39.53500955 | -2.620101775 | 0.8491072284 | -3.085713662 | 0.002030642702 | 0.03279099978 | 1.393563518 |
| ENSG00000011 | 943.6442859 | -1.140221052 | 0.3696157007 | -3.084882622 | 0.002036324998 | 0.02963004923 | 1.602205449 |
| ENSG00000088 | 598.3657698 | -1.368459109 | 0.4437174678 | -3.084077614 | 0.002041843198 | 0.03024108229 | 1.564848717 |
| ENSG00000105 | 367.5326446 | 1.227089934 | 0.3983225873 | 3.080643612 | 0.002065537179 | 0.03618952024 | 1.242301963 |
| ENSG00000235 | 35.28883243 | 2.694456307 | 0.8754020374 | 3.0779644 | 0.00208419812 | 0.03336512422 | 1.393563518 |
| ENSG00000176 | 165.5711079 | 1.240038447 | 0.4028790406 | 3.077942315 | 0.00208435258 | 0.04525437056 | 0.9247574895 |
| ENSG00000116 | 104.7956796 | -1.332213142 | 0.4331774726 | -3.07544419 | 0.002101892614 | 0.03922171749 | 1.130447911 |
| ENSG00000163 | 209.4543746 | 1.322106753 | 0.4300824955 | 3.074077106 | 0.002111548503 | 0.08499011101 | 0.3706356727 |
| ENSG00000184 | 13.6776927 | 3.397083925 | 1.105293153 | 3.073468714 | 0.002115858724 | 0.03377472653 | 1.393563518 |
| ENSG00000107 | 39.88329314 | 2.499609666 | 0.813287461 | 3.073463917 | 0.002115892737 | 0.03242007477 | 1.470104381 |
| ENSG00000135 | 191.4553908 | 1.186828882 | 0.3861721855 | 3.07331529 | 0.002116946946 | 0.02789788959 | 1.814765822 |
| ENSG00000258 | 22.96462611 | 2.502674644 | 0.8143266387 | 3.073305631 | 0.002117015473 | 0.03535064503 | 1.319539707 |
| ENSG00000280 | 16.14452516 | 4.284317323 | 1.394074186 | 3.073234815 | 0.002117517952 | 0.03451281865 | 1.357988055 |
| ENSG00000141 | 55.30710282 | 1.742575751 | 0.5670553045 | 3.0730261 | 0.002118999538 | 0.03451281865 | 1.357988055 |
| ENSG00000104 | 78.15862324 | 1.518812303 | 0.494455691 | 3.071685351 | 0.002128539675 | 0.03192700015 | 1.514054369 |
| ENSG00000166 | 116.2855947 | 1.555378455 | 0.5066008103 | 3.070224964 | 0.002138975903 | 0.03966342894 | 1.133466638 |
| ENSG00000115 | 462.0821645 | -1.006456755 | 0.3280009496 | -3.068456833 | 0.002151674119 | 0.02921250618 | 1.729500602 |
| ENSG00000137 | 219.2084681 | -1.598081665 | 0.5208259508 | -3.068360289 | 0.002152369457 | 0.03839927896 | 1.198705341 |
| ENSG00000149 | 506.8704656 | -1.095019852 | 0.3568815872 | -3.068300217 | 0.002152802216 | 0.03135465678 | 1.572476891 |
| ENSG00000280 | 15.04368338 | -3.926911837 | 1.280279412 | -3.067230324 | 0.00216052312 | 0.03090973185 | 1.605577974 |
| ENSG00000157 | 121.9303179 | 1.536267952 | 0.5008778833 | 3.067150703 | 0.002161098719 | 0.04010814009 | 1.130447911 |
| ENSG00000232 | 116.9227383 | -1.562518585 | 0.5094955733 | -3.066795213 | 0.002163670361 | 0.03596594745 | 1.318051683 |
| ENSG00000280 | 37.49773068 | -2.338389591 | 0.7625585301 | -3.06650506 | 0.002165771428 | 0.03447250176 | 1.393563518 |
| ENSG00000101 | 201.6036588 | -1.437287868 | 0.4688296305 | -3.065693324 | 0.00217165934 | 0.08316070756 | 0.3939394882 |
| ENSG00000114 | 88.85872853 | 1.692721588 | 0.5524544282 | 3.064002208 | 0.002183972979 | 0.03539333481 | 1.357988055 |
| ENSG00000276 | 13.54504957 | 4.159525704 | 1.357637207 | 3.063797664 | 0.002185466674 | 0.03539333481 | 1.357988055 |
| ENSG00000270 | 10.12072802 | 5.392240279 | 1.760972539 | 3.062080845 | 0.002198040783 | 0.03597262267 | 1.337480197 |
| ENSG00000163 | 105.806688 | 1.342187219 | 0.4393023485 | 3.055269847 | 0.00224858109 | 0.03702839402 | 1.318051683 |
| ENSG00000120 | 38.19421052 | -2.373823446 | 0.7770380441 | -3.054964251 | 0.002250873503 | 0.03545492686 | 1.393563518 |
| ENSG00000106 | 431.7281298 | -1.07188929 | 0.3508816993 | -3.054845245 | 0.002251766797 | 0.03242007477 | 1.564848717 |

| | | | | | | | |
|---|---|---|---|---|---|---|---|
| ENSG0000007[illegible] | 311.5126927 | -1.25057425 | 0.4096328933 | -3.052914622 | 0.002266304076 | 0.03872800473 | 1.242301963 |
| ENSG0000015[illegible] | 14.06674696 | 4.456226719 | 1.459683855 | 3.052871142 | 0.002266632462 | 0.03629574229 | 1.357988055 |
| ENSG0000012[illegible] | 374.8365804 | 1.39134026 | 0.4557723087 | 3.052709068 | 0.002267856921 | 0.03716104127 | 1.32213648 |
| ENSG0000016[illegible] | 95.41958502 | 2.553929842 | 0.83661581 | 3.052691344 | 0.002267990861 | 0.0414930328 | 1.133466638 |
| ENSG0000014[illegible] | 55.91904124 | 1.662092076 | 0.5450689021 | 3.049324718 | 0.002293564294 | 0.03716104127 | 1.337480197 |
| ENSG0000027[illegible] | 139.5102215 | -2.017773076 | 0.6620080608 | -3.047958469 | 0.002304017674 | 0.0391745482 | 1.242301963 |
| ENSG0000010[illegible] | 28.82558799 | 2.411297442 | 0.7912801658 | 3.047337146 | 0.002308785935 | 0.03612833345 | 1.393563518 |
| ENSG0000025[illegible] | 11.75260649 | 4.056012703 | 1.331940081 | 3.045191569 | 0.002325321458 | 0.03759546043 | 1.337480197 |
| ENSG0000012[illegible] | 74.53559044 | -1.458527976 | 0.479352184 | -3.042706438 | 0.002344609367 | 0.03780249569 | 1.337480197 |
| ENSG0000013[illegible] | 88.90567243 | 1.885885695 | 0.6200293816 | 3.04160698 | 0.002353189274 | 0.03668550887 | 1.393563518 |
| ENSG0000023[illegible] | 91.80951894 | -1.856824423 | 0.6105256757 | -3.041353536 | 0.002355171163 | 0.03786671701 | 1.337480197 |
| ENSG0000026[illegible] | 48.53445598 | -2.402297541 | 0.7900117305 | -3.040837811 | 0.002359208766 | 0.03447611084 | 1.516424702 |
| ENSG0000019[illegible] | 228.8656786 | -1.171200603 | 0.3854116847 | -3.038830035 | 0.00237498805 | 0.04112247064 | 1.198705341 |
| ENSG0000028[illegible] | 17.02554141 | 3.484242915 | 1.146973084 | 3.037772172 | 0.002383340697 | 0.03781089589 | 1.357988055 |
| ENSG0000024[illegible] | 47.16057754 | 1.916136673 | 0.6308198348 | 3.037533963 | 0.002385225246 | 0.03475690052 | 1.516424702 |
| ENSG0000014[illegible] | 15.25767121 | 3.122714206 | 1.028131186 | 3.03727214 | 0.002387298185 | 0.03324363332 | 1.605577974 |
| ENSG0000012[illegible] | 7.974455127 | 4.032657994 | 1.327841111 | 3.03700342 | 0.002389427448 | 0.03786671701 | 1.357988055 |
| ENSG0000021[illegible] | 93.38406679 | -2.128204853 | 0.7010100829 | -3.035911901 | 0.002398094225 | 0.03793447117 | 1.357988055 |
| ENSG0000000[illegible] | 323.8867971 | -1.002649874 | 0.3303705519 | -3.034925082 | 0.002405954431 | 0.04027325612 | 1.249985076 |
| ENSG0000017[illegible] | 9.919951754 | 4.400053823 | 1.450021332 | 3.034475236 | 0.002409545365 | 0.03844960181 | 1.337480197 |
| ENSG0000016[illegible] | 284.9045425 | -1.199535444 | 0.3954278182 | -3.033513042 | 0.002417242626 | 0.04042582228 | 1.249985076 |
| ENSG0000017[illegible] | 16.60940766 | 3.821314784 | 1.25973704 | 3.033422581 | 0.00241796744 | 0.03852002592 | 1.337480197 |
| ENSG0000011[illegible] | 300.2942403 | 1.280609527 | 0.4222453459 | 3.032856464 | 0.002422507966 | 0.03933413802 | 1.29680518 |
| ENSG0000020[illegible] | 12.28901412 | 3.648362036 | 1.203277012 | 3.03202172 | 0.002429217236 | 0.03859475976 | 1.337480197 |
| ENSG0000025[illegible] | 44.70068107 | 1.818805972 | 0.5999168628 | 3.031763374 | 0.00243129714 | 0.03769183081 | 1.393563518 |
| ENSG0000010[illegible] | 9.178234002 | 3.824810736 | 1.263137453 | 3.028024169 | 0.002461583987 | 0.03900038582 | 1.337480197 |
| ENSG0000015[illegible] | 600.1866444 | -1.059923353 | 0.3501887144 | -3.026720479 | 0.002472224537 | 0.03487670248 | 1.564848717 |
| ENSG0000016[illegible] | 200.5386104 | 1.226576357 | 0.4055340721 | 3.024595075 | 0.00248966207 | 0.09452680217 | 0.3706356727 |
| ENSG0000016[illegible] | 21.71331308 | -2.877572895 | 0.9521204222 | -3.022278304 | 0.002508797759 | 0.03844960181 | 1.393563518 |
| ENSG0000000[illegible] | 126.8418729 | 1.286067135 | 0.4258190142 | 3.020220075 | 0.002525910755 | 0.04492914835 | 1.133466638 |
| ENSG0000000[illegible] | 380.8182334 | -1.452053615 | 0.4808574963 | -3.019717123 | 0.002530108709 | 0.04065926796 | 1.29680518 |
| ENSG0000006[illegible] | 34.87806219 | 2.023331086 | 0.6701766513 | 3.019101131 | 0.002535258864 | 0.03933413802 | 1.357988055 |
| ENSG0000020[illegible] | 25.89212171 | 2.952674352 | 0.9792637058 | 3.015198393 | 0.002568112132 | 0.04159518378 | 1.278627818 |

| | | | | | | | |
|---|---|---|---|---|---|---|---|
| ENSG00000271 | 13.39643823 | 3.622331565 | 1.201505398 | 3.014827541 | 0.002571254129 | 0.03902732409 | 1.393563518 |
| ENSG00000172 | 214.6276329 | -1.313665651 | 0.4358075102 | -3.014325409 | 0.002575513996 | 0.0426865479 | 1.242301963 |
| ENSG00000103 | 406.0473042 | -1.072102892 | 0.3558893696 | -3.012461128 | 0.002591386223 | 0.04065926796 | 1.327567552 |
| ENSG00000145 | 12.81933088 | 3.982928416 | 1.322508735 | 3.011646208 | 0.002598352389 | 0.04057548862 | 1.337480197 |
| ENSG00000230 | 74.50407836 | -2.461500616 | 0.8174265004 | -3.011280665 | 0.002601482713 | 0.03839927896 | 1.449168615 |
| ENSG00000100 | 765.9288629 | 1.262876264 | 0.4194609898 | 3.010712067 | 0.002606358752 | 0.03567453801 | 1.602205449 |
| ENSG00000165 | 189.3255035 | -1.679159886 | 0.5579155528 | -3.009702594 | 0.002615036122 | 0.04311246876 | 1.242301963 |
| ENSG00000078 | 179.9977243 | 1.336277818 | 0.4443847639 | 3.007028879 | 0.002638146995 | 0.05356717234 | 0.9247574895 |
| ENSG00000219 | 20.74378437 | 2.404947121 | 0.8005691854 | 3.004046577 | 0.002664145365 | 0.03615016885 | 1.605577974 |
| ENSG00000172 | 509.9823401 | 3.919911206 | 1.306484872 | 3.000349478 | 0.002696700016 | 0.03314491944 | 1.825593782 |
| ENSG00000279 | 164.9284469 | 1.192992206 | 0.3977198546 | 2.999579207 | 0.002703528203 | 0.04525437056 | 1.198705341 |
| ENSG00000102 | 150.0648057 | -1.14603126 | 0.3820738598 | -2.999501877 | 0.002704214577 | 0.04420283543 | 1.242301963 |
| ENSG00000154 | 335.5816308 | 1.217042805 | 0.4059037441 | 2.998353237 | 0.002714428579 | 0.04416473149 | 1.249985076 |
| ENSG00000176 | 72.51533183 | 1.745004638 | 0.5820127139 | 2.998224259 | 0.00271557768 | 0.04065926796 | 1.393563518 |
| ENSG00000271 | 152.2107751 | -1.110062417 | 0.3703013622 | -2.997727069 | 0.002720011462 | 0.04440514005 | 1.242301963 |
| ENSG00000185 | 168.6110059 | 1.08816496 | 0.3630922361 | 2.996938111 | 0.002727060706 | 0.05457899601 | 0.9247574895 |
| ENSG00000273 | 37.81382164 | 2.104959145 | 0.703445045 | 2.992357626 | 0.002768317587 | 0.04265477791 | 1.337480197 |
| ENSG00000164 | 147.7926253 | -1.224317548 | 0.4093173949 | -2.991120248 | 0.002779560166 | 0.04542643812 | 1.22159078 |
| ENSG00000168 | 106.4310443 | 1.311343223 | 0.4384363989 | 2.990954278 | 0.0027810713 | 0.04542643812 | 1.22159078 |
| ENSG00000218 | 26.6345328 | 3.670205283 | 1.228129672 | 2.988450948 | 0.002803955053 | 0.04440514005 | 1.278627818 |
| ENSG00000105 | 127.2588621 | 1.262863527 | 0.4225975388 | 2.988336208 | 0.002805008036 | 0.04574298408 | 1.22159078 |
| ENSG00000183 | 25.78047026 | 3.139547354 | 1.050613555 | 2.988298922 | 0.002805350291 | 0.04163343568 | 1.393563518 |
| ENSG00000137 | 234.5415156 | 1.07396214 | 0.3594484399 | 2.987805819 | 0.002809880188 | 0.04528732245 | 1.242301963 |
| ENSG00000227 | 29.28464206 | -2.392690993 | 0.801196281 | -2.986398028 | 0.00282284965 | 0.04276235233 | 1.357988055 |
| ENSG00000232 | 129.7485212 | 1.166104621 | 0.390486872 | 2.986283802 | 0.002823904361 | 0.04368787551 | 1.318051683 |
| ENSG00000276 | 12.09585019 | 4.59213488 | 1.537858392 | 2.986058343 | 0.002825987218 | 0.04319869243 | 1.337480197 |
| ENSG00000006 | 55.77997557 | -1.782627871 | 0.5970250149 | -2.985851223 | 0.002827901885 | 0.04027325612 | 1.470104381 |
| ENSG00000224 | 16.23419939 | -3.799388195 | 1.273032104 | -2.98451876 | 0.002840247847 | 0.04287414458 | 1.357988055 |
| ENSG00000153 | 24.91670108 | 2.607380543 | 0.8743910652 | 2.981938685 | 0.002864293581 | 0.0424326939 | 1.393563518 |
| ENSG00000169 | 465.6552569 | 1.257295337 | 0.4216445951 | 2.981884154 | 0.002864803797 | 0.03612833345 | 1.729500602 |
| ENSG00000203 | 196.5436754 | -1.207558529 | 0.405122597 | -2.980723707 | 0.002875681152 | 0.04603066846 | 1.242301963 |
| ENSG00000104 | 103.9171467 | 1.478615037 | 0.4963398464 | 2.979037544 | 0.002891553462 | 0.04954234594 | 1.133466638 |
| ENSG00000087 | 221.3647943 | -1.595288486 | 0.5355662933 | -2.978694713 | 0.002894790392 | 0.0475338529 | 1.198705341 |

| | | | | | | | |
|---|---|---|---|---|---|---|---|
| ENSG00000171 | 7.561512636 | 5.041075838 | 1.693197 | 2.977252994 | 0.002908438983 | 0.04287414458 | 1.393563518 |
| ENSG00000165 | 227.5538184 | 1.135052414 | 0.3817625959 | 2.973189166 | 0.00294722751 | 0.04697498255 | 1.242301963 |
| ENSG00000134 | 117.5830429 | -1.300463063 | 0.4375334291 | -2.972259893 | 0.002956163316 | 0.05032214755 | 1.130447911 |
| ENSG00000226 | 23.91799118 | -3.099450095 | 1.043524337 | -2.970175189 | 0.002976299643 | 0.04440514005 | 1.357988055 |
| ENSG00000143 | 105.515387 | -1.331529892 | 0.4484861567 | -2.968943126 | 0.002988258987 | 0.04790601019 | 1.22159078 |
| ENSG00000175 | 366.1071224 | -1.028661553 | 0.3464813265 | -2.968880209 | 0.002988870885 | 0.0474366937 | 1.242301963 |
| ENSG00000282 | 47.30136203 | -1.832897818 | 0.6175455376 | -2.968036698 | 0.002997085452 | 0.04105787066 | 1.516424702 |
| ENSG00000131 | 161.6568537 | -1.938793014 | 0.6532913066 | -2.967731232 | 0.003000065321 | 0.04805479815 | 1.22159078 |
| ENSG00000267 | 31.60716442 | 2.70709603 | 0.9124957171 | 2.966694505 | 0.003010198923 | 0.04525437056 | 1.337480197 |
| ENSG00000179 | 25.26684348 | -3.001031748 | 1.011727699 | -2.966244526 | 0.003014607007 | 0.04525437056 | 1.337480197 |
| ENSG00000175 | 15.29218584 | -3.405666483 | 1.148684181 | -2.964841457 | 0.003028389559 | 0.03966342894 | 1.605577974 |
| ENSG00000185 | 20.24874676 | 2.736813367 | 0.9235828982 | 2.963256869 | 0.003044024305 | 0.04542643812 | 1.337480197 |
| ENSG00000196 | 391.0659287 | -3.999589897 | 1.350051024 | -2.962547211 | 0.003051050168 | 0.04788100588 | 1.249985076 |
| ENSG00000267 | 42.21085549 | -2.190707277 | 0.7398602289 | -2.960974508 | 0.003066673206 | 0.0452545023 | 1.357988055 |
| ENSG00000186 | 15.95607797 | 3.382692738 | 1.143181904 | 2.959015293 | 0.003086237847 | 0.04476648728 | 1.393563518 |
| ENSG00000087 | 469.0975189 | 1.111491853 | 0.376031446 | 2.955848148 | 0.003118105667 | 0.03844960181 | 1.729500602 |
| ENSG00000054 | 126.0867851 | -1.264173309 | 0.4277232939 | -2.955586771 | 0.003120748993 | 0.04957063796 | 1.22159078 |
| ENSG00000079 | 99.19736113 | 1.449782995 | 0.490524971 | 2.955574294 | 0.003120875227 | 0.05221823646 | 1.130447911 |
| ENSG00000135 | 53.03924622 | -1.841903022 | 0.6235735066 | -2.953786527 | 0.003139010597 | 0.04600470206 | 1.357988055 |
| ENSG00000153 | 164.9415591 | 1.551610435 | 0.5257451503 | 2.951259625 | 0.003164807737 | 0.06010113344 | 0.9247574895 |
| ENSG00000223 | 10.65041199 | 3.150365858 | 1.067503067 | 2.951153918 | 0.003165891102 | 0.04542643812 | 1.393563518 |
| ENSG00000155 | 114.653462 | 1.160516355 | 0.3934217647 | 2.949802118 | 0.003179775156 | 0.05284704932 | 1.133466638 |
| ENSG00000074 | 14.26854152 | 3.492447229 | 1.185629872 | 2.945647128 | 0.003222798351 | 0.04341981157 | 1.514850267 |
| ENSG00000158 | 83.59342936 | 1.29702706 | 0.4403405429 | 2.945509063 | 0.00322423702 | 0.04491287372 | 1.449168615 |
| ENSG00000144 | 13.24810554 | 3.507751979 | 1.191641985 | 2.943629062 | 0.003243885376 | 0.04620937238 | 1.393563518 |
| ENSG00000151 | 18.47647007 | 3.04367845 | 1.034293481 | 2.942760935 | 0.003252995161 | 0.04374982283 | 1.514850267 |
| ENSG00000147 | 22.39416372 | -2.513334868 | 0.8545679256 | -2.94105921 | 0.003270920041 | 0.04655472668 | 1.393563518 |
| ENSG00000100 | 25.2756436 | 2.470117853 | 0.8400665914 | 2.940383391 | 0.003278063614 | 0.0475338529 | 1.357988055 |
| ENSG00000165 | 331.8264022 | 1.364769828 | 0.464224661 | 2.939890839 | 0.00328327896 | 0.04831171188 | 1.327567552 |
| ENSG00000234 | 97.04087756 | -1.24219225 | 0.4228068474 | -2.937966254 | 0.003303729826 | 0.0488404744 | 1.318051683 |
| ENSG00000151 | 90.09813593 | -1.562897956 | 0.5330720311 | -2.931870113 | 0.003369276285 | 0.04842583825 | 1.357988055 |
| ENSG00000287 | 13.25332476 | 3.618541903 | 1.235062514 | 2.92984514 | 0.003391309667 | 0.04785752012 | 1.393563518 |
| ENSG00000197 | 7.809577833 | 4.218813868 | 1.440694245 | 2.928320068 | 0.003407990227 | 0.04885919388 | 1.357988055 |

| | | | | | | | |
|---|---|---|---|---|---|---|---|
| ENSG000002242 | 12.2166907 | 3.880365487 | 1.325691958 | 2.927049126 | 0.003421948243 | 0.04287414458 | 1.636424459 |
| ENSG000002290 | 11.41351801 | 4.756525367 | 1.625825317 | 2.925606654 | 0.003437853108 | 0.04823107334 | 1.393563518 |
| ENSG000000771 | 68.07802311 | -1.459338936 | 0.4989513004 | -2.924812371 | 0.003446639671 | 0.04983408169 | 1.337480197 |
| ENSG000001239 | 118.1368208 | -1.733575191 | 0.5927443092 | -2.924659358 | 0.003448334694 | 0.05590599454 | 1.133466638 |
| ENSG000001352 | 176.491819 | 1.063453863 | 0.3637056144 | 2.923941291 | 0.003456299284 | 0.06393111234 | 0.9247574895 |
| ENSG000001065 | 7401.085706 | -1.001477389 | 0.3425729558 | -2.923398863 | 0.003462326835 | 0.04411871344 | 1.597450288 |
| ENSG000001668 | 381.9760471 | -1.300468643 | 0.4450854843 | -2.92184016 | 0.003479700684 | 0.05039728849 | 1.327567552 |
| ENSG000001823 | 58.78449799 | -8.674881247 | 2.969225596 | -2.92159722 | 0.003482415716 | 0.05018734014 | 1.337480197 |
| ENSG000002368 | 604.5394623 | 1.141368056 | 0.39080787 | 2.920534983 | 0.003494309635 | 0.04012120102 | 1.825593782 |
| ENSG000001751 | 36.85738174 | 3.306954611 | 1.132599679 | 2.919791231 | 0.003502659457 | 0.05087013435 | 1.319539707 |
| ENSG000002789 | 11.83474775 | -3.757281606 | 1.288170336 | -2.916758367 | 0.003536896569 | 0.04937145425 | 1.393563518 |
| ENSG000002047 | 16.62702223 | -2.927962159 | 1.004021969 | -2.91623316 | 0.003542856321 | 0.04941329624 | 1.393563518 |
| ENSG000001513 | 23.61464379 | 2.476925981 | 0.8497494004 | 2.914889943 | 0.003558139949 | 0.05094073559 | 1.337480197 |
| ENSG000001601 | 163.6641411 | -3.357476028 | 1.1518422 | -2.914874996 | 0.003558310359 | 0.05432639076 | 1.22159078 |
| ENSG000001647 | 43.32908653 | 2.441478115 | 0.8377323638 | 2.914389154 | 0.003563853449 | 0.04958178057 | 1.393563518 |
| ENSG000001582 | 21.98105475 | 3.224594087 | 1.106673753 | 2.913771179 | 0.003570915428 | 0.0510142668 | 1.337480197 |
| ENSG000001381 | 15.00688331 | 3.10432151 | 1.065745012 | 2.912818239 | 0.00358183021 | 0.04685818997 | 1.514850267 |
| ENSG000002692 | 54.9703399 | 1.743235468 | 0.5985731632 | 2.912318118 | 0.003587570623 | 0.04788100588 | 1.470104381 |
| ENSG000000678 | 20.97813882 | 2.640846465 | 0.9070687669 | 2.911407119 | 0.003598048635 | 0.05133166672 | 1.337480197 |
| ENSG000001632 | 377.4397676 | 1.057127733 | 0.3633637513 | 2.909282309 | 0.00362259576 | 0.05432639076 | 1.242301963 |
| ENSG000000100 | 52.78104684 | -2.004227284 | 0.689297031 | -2.907639513 | 0.003641678642 | 0.04738967299 | 1.516424702 |
| ENSG000001003 | 56.65347721 | 1.585153386 | 0.5454172661 | 2.906313175 | 0.003657152165 | 0.05133166672 | 1.357988055 |
| ENSG000000735 | 66.87568618 | -1.559762824 | 0.536717574 | -2.906114685 | 0.003659472955 | 0.05040758184 | 1.393563518 |
| ENSG000001834 | 34.79133845 | 2.000113423 | 0.6883692861 | 2.905582023 | 0.003665707577 | 0.05195488736 | 1.337480197 |
| ENSG000001606 | 38.89835239 | 1.794032653 | 0.6174821058 | 2.905400232 | 0.003667837593 | 0.05195488736 | 1.337480197 |
| ENSG000001728 | 259.0721698 | -1.187099743 | 0.4086832683 | -2.904693769 | 0.003676125773 | 0.05603966512 | 1.198705341 |
| ENSG000001716 | 135.399981 | 1.316500105 | 0.4532956257 | 2.904285925 | 0.003680918329 | 0.05829427493 | 1.133466638 |
| ENSG000001641 | 128.3886243 | 1.124313675 | 0.3878299061 | 2.898986534 | 0.003743709728 | 0.05550906026 | 1.242301963 |
| ENSG000001438 | 41.95291659 | 2.252246511 | 0.7778901369 | 2.89532725 | 0.003787634417 | 0.04871011805 | 1.516424702 |
| ENSG000001660 | 15.29079164 | 3.72633205 | 1.287234443 | 2.894835568 | 0.003793571937 | 0.05171538424 | 1.393563518 |
| ENSG000002048 | 91.12726527 | -1.384444075 | 0.478355477 | -2.894174189 | 0.003801572064 | 0.05039728849 | 1.449168615 |
| ENSG000002441 | 31.3177362 | 3.215616182 | 1.111986847 | 2.891775375 | 0.00383071721 | 0.05362519782 | 1.337480197 |
| ENSG000001759 | 38.00239378 | 1.811674128 | 0.6265154355 | 2.891667188 | 0.003832036439 | 0.05362519782 | 1.337480197 |

| | | | | | | | |
|---|---|---|---|---|---|---|---|
| ENSG00000143 4 | 135.3434792 | 1.18494366 | 0.4098851514 | 2.890916288 | 0.003841204237 | 0.05641817495 | 1.242301963 |
| ENSG00000140 5 | 10.73245267 | -4.541002412 | 1.57142453 | -2.88973624 | 0.00385565181 | 0.05706947995 | 1.227924265 |
| ENSG00000171 5 | 33.09904666 | 2.272011549 | 0.7864701629 | 2.888871894 | 0.003866265484 | 0.05234828155 | 1.393563518 |
| ENSG00000189 4 | 154.1255298 | 1.280144813 | 0.4433002694 | 2.88776006 | 0.003879957198 | 0.05750219169 | 1.22159078 |
| ENSG00000163 5 | 32.39072332 | -2.468491874 | 0.8549034433 | -2.887451084 | 0.003883769907 | 0.05454368933 | 1.319539707 |
| ENSG00000148 3 | 79.82081092 | 1.450977447 | 0.5025328614 | 2.887328488 | 0.003885283655 | 0.05362519782 | 1.357988055 |
| ENSG00000167 7 | 78.28345303 | -1.81547614 | 0.6290232365 | -2.88618295 | 0.00389945412 | 0.05432639076 | 1.337480197 |
| ENSG00000235 6 | 16.17960068 | 4.264607286 | 1.477894727 | 2.885596118 | 0.003906731474 | 0.04983408169 | 1.514850267 |
| ENSG00000168 3 | 11.70687388 | 3.80228833 | 1.318016916 | 2.884855485 | 0.003915933741 | 0.04726189635 | 1.636424459 |
| ENSG00000285 9 | 14.05989521 | -3.381498994 | 1.172248111 | -2.884627378 | 0.003918771899 | 0.05293031672 | 1.393563518 |
| ENSG00000112 5 | 292.2947531 | 1.156055245 | 0.4014272027 | 2.879862742 | 0.003978483352 | 0.05753339233 | 1.249985076 |
| ENSG00000163 3 | 3903.5487 | 1.02234803 | 0.3550288311 | 2.87961974 | 0.003981550735 | 0.04785752012 | 1.634803475 |
| ENSG00000116 1 | 106.8422173 | -1.331500207 | 0.4625069148 | -2.878876325 | 0.003990948092 | 0.0579522016 | 1.242301963 |
| ENSG00000138 1 | 141.7918677 | 1.36892046 | 0.4756884251 | 2.877767018 | 0.004005008061 | 0.04464892509 | 1.814765822 |
| ENSG00000169 7 | 73.14000394 | -1.566759636 | 0.5445587387 | -2.877117793 | 0.004013257547 | 0.05465300387 | 1.357988055 |
| ENSG00000144 0 | 74.82397627 | 1.386243176 | 0.4821808053 | 2.874944753 | 0.004040981992 | 0.05485599691 | 1.357988055 |
| ENSG00000104 4 | 126.2449862 | 1.208024786 | 0.4202141171 | 2.874783919 | 0.004043040865 | 0.05832962937 | 1.242301963 |
| ENSG00000144 8 | 359.8921809 | -1.227864927 | 0.4274043673 | -2.872841321 | 0.004067983831 | 0.05705869849 | 1.29680518 |
| ENSG00000105 4 | 46.83453105 | 1.912825987 | 0.6658736088 | 2.872656255 | 0.004070367352 | 0.05232732771 | 1.470104381 |
| ENSG00000182 9 | 80.22325388 | 1.526054362 | 0.5314522331 | 2.871479819 | 0.004085548679 | 0.05447717079 | 1.393563518 |
| ENSG00000279 0 | 68.79626003 | 2.142695342 | 0.7462219446 | 2.871391491 | 0.004086690576 | 0.05603966512 | 1.337480197 |
| ENSG00000124 1 | 1643.700589 | -2.42430385 | 0.8443452898 | -2.871223278 | 0.00408886604 | 0.05016257211 | 1.572476891 |
| ENSG00000160 3 | 41.57106402 | 2.073934849 | 0.7226040172 | 2.870084859 | 0.004103616533 | 0.05454368933 | 1.393563518 |
| ENSG00000103 1 | 173.6972161 | 1.087450813 | 0.3789152758 | 2.869904917 | 0.004105952467 | 0.05869421963 | 1.242301963 |
| ENSG00000210 1 | 504.1331956 | -1.497964497 | 0.5221397639 | -2.868895649 | 0.004119076702 | 0.04525437056 | 1.825593782 |
| ENSG00000197 7 | 73.73673063 | 1.332164048 | 0.464662988 | 2.866946761 | 0.00414452727 | 0.05485599691 | 1.393563518 |
| ENSG00000091 6 | 92.69740026 | -1.888777155 | 0.6590991635 | -2.865694966 | 0.004160949646 | 0.05603966512 | 1.357988055 |
| ENSG00000172 5 | 317.5165468 | 1.023993239 | 0.3573572242 | 2.865461139 | 0.004164023773 | 0.05901996077 | 1.249985076 |
| ENSG00000254 9 | 152.5504022 | -1.325295626 | 0.4625486484 | -2.865202677 | 0.004167424176 | 0.04574298408 | 1.814765822 |
| ENSG00000102 3 | 17.22644614 | 3.348184532 | 1.169917484 | 2.861898021 | 0.004211123794 | 0.05561809674 | 1.393563518 |
| ENSG00000165 8 | 33.91818838 | 2.283598961 | 0.7981009315 | 2.861290936 | 0.004219196729 | 0.05868921124 | 1.278627818 |
| ENSG00000091 8 | 65.54593826 | -1.457561483 | 0.5098767574 | -2.858654492 | 0.004254418874 | 0.05603966512 | 1.393563518 |
| ENSG00000269 9 | 7.622747629 | 4.679429095 | 1.637804178 | 2.857135888 | 0.00427482782 | 0.05603966512 | 1.393563518 |

| | | | | | | | |
|---|---|---|---|---|---|---|---|
| ENSG000001357 | 38.58809354 | -2.120199525 | 0.7422740548 | -2.856356775 | 0.004285332966 | 0.05722012868 | 1.357988055 |
| ENSG000000199 | 336.2333902 | 1.120210177 | 0.3923237248 | 2.855321018 | 0.004299334813 | 0.05822296886 | 1.327567552 |
| ENSG000001434 | 11.61001578 | -4.013973057 | 1.407958637 | -2.850916889 | 0.004359336231 | 0.05833856297 | 1.337480197 |
| ENSG000001656 | 31.58372966 | -2.37964763 | 0.8349110219 | -2.850181119 | 0.004369433979 | 0.0579522016 | 1.357988055 |
| ENSG000001018 | 7.304746478 | -4.52644385 | 1.588212386 | -2.850024272 | 0.004371589293 | 0.05840135586 | 1.337480197 |
| ENSG000001703 | 65.57724503 | 1.397420698 | 0.4908355863 | 2.847024007 | 0.004413003422 | 0.05737584214 | 1.393563518 |
| ENSG000000216 | 77.57153606 | 1.970388727 | 0.6924389478 | 2.845577553 | 0.004433096308 | 0.05753339233 | 1.393563518 |
| ENSG000002871 | 8.649489363 | 4.857046497 | 1.707298316 | 2.844872774 | 0.004442916486 | 0.05841299161 | 1.357988055 |
| ENSG000001882 | 16.42892516 | -3.095073903 | 1.088400417 | -2.84369048 | 0.00445943452 | 0.05236408832 | 1.605577974 |
| ENSG000001641 | 61.07632112 | 1.92314298 | 0.6768203725 | 2.841437785 | 0.004491061392 | 0.05472625553 | 1.516424702 |
| ENSG000001518 | 262.9092281 | -1.052264019 | 0.3703421168 | -2.841329601 | 0.004492585349 | 0.06264189433 | 1.242301963 |
| ENSG000001838 | 53.5428849 | 2.48248861 | 0.8740920683 | 2.840076806 | 0.00451026729 | 0.05603966512 | 1.470104381 |
| ENSG000001756 | 23.50836463 | 2.809634587 | 0.9899885262 | 2.838047627 | 0.004539040927 | 0.059811669 | 1.337480197 |
| ENSG000002509 | 57.28729993 | -2.292319125 | 0.8077892366 | -2.83776884 | 0.004543007074 | 0.05833856297 | 1.393563518 |
| ENSG000001272 | 50.14871408 | 1.721979002 | 0.6068226717 | 2.837697209 | 0.004544026629 | 0.05983121324 | 1.337480197 |
| ENSG000001137 | 176.5827992 | 1.096444425 | 0.38645769 | 2.837165499 | 0.00455160122 | 0.07684196265 | 0.9247574895 |
| ENSG000000894 | 127.0273163 | 1.32834854 | 0.4682079595 | 2.837090897 | 0.004552664883 | 0.06313238265 | 1.242301963 |
| ENSG000000953 | 101.3008285 | -1.715540845 | 0.6047605573 | -2.836727403 | 0.004557850798 | 0.0675742635 | 1.130447911 |
| ENSG000001873 | 46.67566338 | 1.862376241 | 0.6569029518 | 2.835085815 | 0.004581337755 | 0.05959210207 | 1.357988055 |
| ENSG000001983 | 50.54908255 | 1.639783993 | 0.5787236281 | 2.833449186 | 0.004604862842 | 0.05869421963 | 1.393563518 |
| ENSG000001145 | 45.13783798 | -1.567310449 | 0.5532502387 | -2.832914184 | 0.004612576709 | 0.06045899874 | 1.337480197 |
| ENSG000001438 | 23.14735128 | 3.425221469 | 1.209267555 | 2.832476118 | 0.004618901641 | 0.06049627917 | 1.337480197 |
| ENSG000001823 | 11.60555877 | -4.988719984 | 1.761691593 | -2.831778276 | 0.004628993489 | 0.05894361119 | 1.393563518 |
| ENSG000001823 | 72.20702764 | 8.402459796 | 2.968348872 | 2.83068472 | 0.004644848156 | 0.05603966512 | 1.514054369 |
| ENSG000001408 | 96.25194391 | -1.207451122 | 0.426585949 | -2.830499047 | 0.004647544968 | 0.06855805758 | 1.130447911 |
| ENSG000001158 | 13.5852972 | 3.497953291 | 1.23583845 | 2.830429245 | 0.004648559178 | 0.06010113344 | 1.357988055 |
| ENSG000001970 | 134.7686679 | 1.203024264 | 0.4251897289 | 2.829382231 | 0.004663796118 | 0.06416847382 | 1.242301963 |
| ENSG000002604 | 30.44215737 | -2.240688439 | 0.7923109043 | -2.828041905 | 0.004683367563 | 0.06313238265 | 1.278627818 |
| ENSG000001732 | 94.40922006 | 1.319899924 | 0.4667831002 | 2.82765148 | 0.004689082509 | 0.06897288323 | 1.130447911 |
| ENSG000002101 | 86.47003165 | -1.720730295 | 0.608544084 | -2.827618147 | 0.004689570726 | 0.05822296886 | 1.449168615 |
| ENSG000001567 | 47.79686347 | 2.462870207 | 0.8715937087 | 2.825709023 | 0.00471760971 | 0.05663155603 | 1.516424702 |
| ENSG000000435 | 21.07713403 | 3.366461171 | 1.19212269 | 2.823921732 | 0.004743996801 | 0.05986740222 | 1.393563518 |
| ENSG000001608 | 10.66031982 | 3.922018023 | 1.389528677 | 2.822552775 | 0.004764297996 | 0.0617952356 | 1.337480197 |

| | | | | | | | |
|---|---|---|---|---|---|---|---|
| ENSG00000169 | 24.10717461 | -2.246505413 | 0.7961119436 | -2.821846137 | 0.004774807942 | 0.06392345236 | 1.278627818 |
| ENSG00000177 | 51.06587277 | 1.853703096 | 0.6569884022 | 2.821515706 | 0.004779729693 | 0.06126942449 | 1.357988055 |
| ENSG00000283 | 61.13833235 | -1.508378065 | 0.535167152 | -2.818517653 | 0.004824595801 | 0.05852266444 | 1.470104381 |
| ENSG00000196 | 29.11274216 | -2.293540102 | 0.8139303222 | -2.81785804 | 0.004834517965 | 0.06068057154 | 1.393563518 |
| ENSG00000066 | 305.7352784 | -1.052682257 | 0.3736387973 | -2.817379418 | 0.00484172914 | 0.06392345236 | 1.29680518 |
| ENSG00000241 | 16.91295741 | -3.132560404 | 1.112458009 | -2.815890917 | 0.004864217928 | 0.06209212594 | 1.357988055 |
| ENSG00000154 | 83.75754293 | -1.629112614 | 0.5785510317 | -2.815849466 | 0.004864845547 | 0.06092032529 | 1.393563518 |
| ENSG00000011 | 54.30715715 | -2.219722754 | 0.7883400151 | -2.815692102 | 0.004867228849 | 0.0579522016 | 1.516424702 |
| ENSG00000233 | 10.1486363 | -3.698311365 | 1.31354074 | -2.815528482 | 0.004869708015 | 0.06685213904 | 1.227924265 |
| ENSG00000128 | 28.05961791 | -2.444798935 | 0.8684644437 | -2.815082359 | 0.004876473487 | 0.06292151118 | 1.337480197 |
| ENSG00000108 | 364.1555487 | -1.295407535 | 0.4603004051 | -2.814265469 | 0.00488888368 | 0.0665088697 | 1.242301963 |
| ENSG00000211 | 77.16354812 | 1.794316937 | 0.6376803628 | 2.813818712 | 0.004895682901 | 0.0596289029 | 1.449168615 |
| ENSG00000196 | 36.89891306 | 2.395610549 | 0.8515188765 | 2.81333816 | 0.004903005988 | 0.06525530275 | 1.278627818 |
| ENSG00000276 | 11.58487222 | 3.924842848 | 1.396451543 | 2.810582915 | 0.004945184521 | 0.06163726374 | 1.393563518 |
| ENSG00000180 | 50.91910439 | -1.935673555 | 0.6887705921 | -2.810331303 | 0.004949052623 | 0.06362184408 | 1.337480197 |
| ENSG00000091 | 44.24783229 | -2.030435409 | 0.7226065895 | -2.809876687 | 0.004956048489 | 0.06366468917 | 1.337480197 |
| ENSG00000184 | 114.6940663 | 1.424855472 | 0.5071099343 | 2.809756575 | 0.004957898318 | 0.07131870037 | 1.130447911 |
| ENSG00000272 | 32.99137429 | 2.20404402 | 0.784438908 | 2.809707674 | 0.004958651621 | 0.06418465957 | 1.319539707 |
| ENSG00000017 | 150.1830734 | -1.383632996 | 0.492525893 | -2.809259402 | 0.004965561906 | 0.06723283423 | 1.242301963 |
| ENSG00000119 | 190.2177265 | -1.050089577 | 0.3738950119 | -2.808514538 | 0.004977063518 | 0.06729596068 | 1.242301963 |
| ENSG00000172 | 188.3619696 | 1.06517152 | 0.3794203519 | 2.807365274 | 0.004994856829 | 0.05210175104 | 1.814765822 |
| ENSG00000127 | 48.53294394 | 1.701421345 | 0.606066563 | 2.807317626 | 0.004995595773 | 0.05863696666 | 1.516424702 |
| ENSG00000061 | 29.01355534 | 2.227946472 | 0.7940994384 | 2.805626556 | 0.005021885639 | 0.06362184408 | 1.357988055 |
| ENSG00000118 | 32.00189924 | 2.575676117 | 0.9182890218 | 2.804864325 | 0.005033776355 | 0.0665088697 | 1.278627818 |
| ENSG00000142 | 1771.221251 | 1.037417794 | 0.369928553 | 2.8043734 | 0.005041448193 | 0.05714462248 | 1.602205449 |
| ENSG00000146 | 13.54736361 | -3.84826031 | 1.372620687 | -2.803586123 | 0.005053773292 | 0.05901996077 | 1.514850267 |
| ENSG00000097 | 714.9829059 | 1.025394145 | 0.3658066483 | 2.803104179 | 0.005061331724 | 0.05234828155 | 1.825593782 |
| ENSG00000259 | 11.4794168 | 3.537541828 | 1.2620497 | 2.803013089 | 0.005062761468 | 0.0628076549 | 1.393563518 |
| ENSG00000277 | 12.62250157 | 3.791764905 | 1.352881748 | 2.802731955 | 0.005067176393 | 0.06392345236 | 1.357988055 |
| ENSG00000101 | 209.7067358 | -1.05802606 | 0.3775475325 | -2.802365183 | 0.005072941429 | 0.06987036086 | 1.198705341 |
| ENSG00000123 | 15.353204 | 2.618511418 | 0.9344041774 | 2.802332739 | 0.005073451674 | 0.06469467642 | 1.337480197 |
| ENSG00000204 | 92.96926746 | 1.717086359 | 0.6129582177 | 2.80131061 | 0.005089550551 | 0.0648523824 | 1.337480197 |
| ENSG00000109 | 25.55707023 | 2.02348439 | 0.7228801596 | 2.799197575 | 0.005122978048 | 0.06592419219 | 1.319539707 |

| | | | | | | | |
|---|---|---|---|---|---|---|---|
| ENSG00000149 | 17.22658602 | 3.242027454 | 1.158669692 | 2.798060117 | 0.005141054301 | 0.059811669 | 1.514850267 |
| ENSG00000143 | 41.3074911 | 2.016432273 | 0.7206884463 | 2.797925072 | 0.005143204234 | 0.06096463842 | 1.470104381 |
| ENSG00000100 | 11.99722456 | 3.37479841 | 1.20627295 | 2.797707111 | 0.005146675906 | 0.0693469155 | 1.227924265 |
| ENSG00000130 | 27.75316547 | 2.547801215 | 0.9107614731 | 2.797440703 | 0.005150922124 | 0.06756563118 | 1.278627818 |
| ENSG00000079 | 179.2084716 | 1.255605402 | 0.4488927887 | 2.797116447 | 0.005156094625 | 0.08389049778 | 0.9247574895 |
| ENSG00000248 | 7.691191188 | 4.34959651 | 1.555278146 | 2.796667928 | 0.005163257112 | 0.05722012868 | 1.636424459 |
| ENSG00000099 | 288.987247 | -1.028726287 | 0.3678780667 | -2.796378421 | 0.00516788507 | 0.06605168239 | 1.327567552 |
| ENSG00000259 | 19.51076264 | -2.526355315 | 0.9036711612 | -2.795657783 | 0.005179421243 | 0.06580456298 | 1.337480197 |
| ENSG00000183 | 1277.842645 | -1.095286715 | 0.391788507 | -2.795607057 | 0.005180234148 | 0.05171538424 | 1.902376403 |
| ENSG00000279 | 103.3083571 | 1.484493038 | 0.5315009249 | 2.793020611 | 0.005221836456 | 0.07032400422 | 1.22159078 |
| ENSG00000213 | 27.89603627 | 2.278088123 | 0.8166777404 | 2.789457837 | 0.005279637037 | 0.06659102842 | 1.337480197 |
| ENSG00000139 | 399.8257565 | -1.017470686 | 0.3648942434 | -2.788398844 | 0.005296928667 | 0.06826066131 | 1.29680518 |
| ENSG00000114 | 228.0951465 | 1.13776663 | 0.4082987071 | 2.786603559 | 0.005326359619 | 0.07043606854 | 1.242301963 |
| ENSG00000204 | 7.929255695 | 4.541461019 | 1.631388313 | 2.783801369 | 0.005372592493 | 0.05852266444 | 1.636424459 |
| ENSG00000272 | 55.20264587 | 1.764002249 | 0.6344227618 | 2.780483858 | 0.005427795801 | 0.06790326806 | 1.337480197 |
| ENSG00000099 | 10.63093899 | -3.46865081 | 1.247832735 | -2.779740195 | 0.005440240392 | 0.05894780937 | 1.636424459 |
| ENSG00000131 | 74.7391278 | -1.525978774 | 0.5492214003 | -2.778440121 | 0.005462057945 | 0.06750855515 | 1.357988055 |
| ENSG00000105 | 14.80284389 | 4.374779358 | 1.574986096 | 2.777662209 | 0.005475150421 | 0.06647078194 | 1.393563518 |
| ENSG00000164 | 24.2472497 | 3.161981988 | 1.138558121 | 2.777181006 | 0.005483263373 | 0.06899805379 | 1.319539707 |
| ENSG00000186 | 9.821941734 | -4.211491811 | 1.517257401 | -2.775726655 | 0.005507849362 | 0.05949885012 | 1.636424459 |
| ENSG00000106 | 281.5478003 | -1.047152541 | 0.3772711534 | -2.775596627 | 0.005510052332 | 0.07174968809 | 1.242301963 |
| ENSG00000168 | 63.38470377 | 1.534677052 | 0.552934255 | 2.775514517 | 0.005511443872 | 0.06866762934 | 1.337480197 |
| ENSG00000250 | 26.28503307 | 2.500230203 | 0.9009451847 | 2.775119114 | 0.005518149337 | 0.0707737271 | 1.278627818 |
| ENSG00000182 | 289.3803249 | 1.16487138 | 0.4198360043 | 2.774586667 | 0.005527190497 | 0.07191245975 | 1.242301963 |
| ENSG00000137 | 28.01523975 | -2.192035886 | 0.7901655188 | -2.774147737 | 0.005534653763 | 0.06690103494 | 1.393563518 |
| ENSG00000215 | 20.4663335 | 3.68432234 | 1.328319498 | 2.773671805 | 0.005542756454 | 0.06895889014 | 1.337480197 |
| ENSG00000131 | 472.7010178 | -1.051731689 | 0.3792886347 | -2.77290589 | 0.00555581854 | 0.06163726374 | 1.564848717 |
| ENSG00000220 | 98.36605451 | -1.424171825 | 0.5136387308 | -2.772711129 | 0.005559144469 | 0.07193326115 | 1.242301963 |
| ENSG00000127 | 86.38590547 | 1.444435909 | 0.5209839825 | 2.772515006 | 0.005562495475 | 0.06840419224 | 1.357988055 |
| ENSG00000162 | 27.1481608 | 2.238391695 | 0.8074574904 | 2.772148035 | 0.005568770518 | 0.06843219785 | 1.357988055 |
| ENSG00000188 | 437.0398347 | -1.119225649 | 0.4037593491 | -2.772011724 | 0.005571103002 | 0.06159901142 | 1.572476891 |
| ENSG00000171 | 13.26142693 | 3.093527572 | 1.116851984 | 2.769863525 | 0.005607978502 | 0.0687659117 | 1.357988055 |
| ENSG00000140 | 201.3537014 | 1.024191022 | 0.3697641553 | 2.769849396 | 0.005608221768 | 0.07242322421 | 1.242301963 |

| | | | | | | | |
|---|---|---|---|---|---|---|---|
| ENSG000000998 | 85.76088616 | -1.361347146 | 0.491830936 | -2.767916872 | 0.005641584349 | 0.06899805379 | 1.357988055 |
| ENSG000001443 | 258.5834584 | 1.24375518 | 0.4494170158 | 2.767485734 | 0.005649051783 | 0.07010357188 | 1.327567552 |
| ENSG000001476 | 3383.709034 | -1.090838976 | 0.3946405257 | -2.764133192 | 0.005707423689 | 0.06092032529 | 1.634803475 |
| ENSG000001562 | 30.84736911 | 2.443542191 | 0.8842611778 | 2.763371561 | 0.005720760242 | 0.07193326115 | 1.278627818 |
| ENSG000001964 | 332.2876911 | 1.028520319 | 0.3724751406 | 2.761312655 | 0.005756953502 | 0.07108118798 | 1.32213648 |
| ENSG000002296 | 9.48892414 | -3.843160461 | 1.391941129 | -2.761007905 | 0.005762328164 | 0.06126942449 | 1.636424459 |
| ENSG000001309 | 101.8448502 | 1.436803722 | 0.5203991285 | 2.76096489 | 0.005763087162 | 0.07853506831 | 1.130447911 |
| ENSG000002724 | 56.29875915 | -1.806493745 | 0.6543062483 | -2.760929992 | 0.005763702986 | 0.06997368789 | 1.357988055 |
| ENSG000001240 | 93.06862699 | 1.339582819 | 0.4857121871 | 2.757976544 | 0.005816037101 | 0.06914980542 | 1.393563518 |
| ENSG000001713 | 153.8005598 | -1.426493025 | 0.5173060209 | -2.757541896 | 0.00582377499 | 0.0753490991 | 1.22159078 |
| ENSG000001241 | 69.32310931 | -1.469381681 | 0.5329282887 | -2.757184619 | 0.005830142401 | 0.07108947723 | 1.337480197 |
| ENSG000002061 | 55.44949303 | 1.67824099 | 0.6087444448 | 2.756889208 | 0.005835411981 | 0.07054407657 | 1.357988055 |
| ENSG000002798 | 27.5772732 | 2.368839825 | 0.8598091438 | 2.755076335 | 0.005867844347 | 0.07131870037 | 1.337480197 |
| ENSG000000753 | 11.74545904 | 3.562246936 | 1.294064405 | 2.752758613 | 0.005909545112 | 0.06996237184 | 1.393563518 |
| ENSG000001832 | 63.64732353 | -1.494993686 | 0.5432585578 | -2.751900847 | 0.005925045741 | 0.07108947723 | 1.357988055 |
| ENSG000002727 | 18.65362266 | -3.031528413 | 1.101638265 | -2.751836522 | 0.005926209635 | 0.07108947723 | 1.357988055 |
| ENSG000000697 | 145.8458266 | -1.292499391 | 0.4697368353 | -2.751539359 | 0.005931589143 | 0.05840135586 | 1.814765822 |
| ENSG000002261 | 154.3051441 | -8.214948719 | 2.98580704 | -2.751332759 | 0.005935331796 | 0.07631427341 | 1.22159078 |
| ENSG000001988 | 88.00528905 | 1.412046038 | 0.5142809971 | 2.745670258 | 0.006038742454 | 0.06913043605 | 1.449168615 |
| ENSG000001389 | 26.62896543 | 2.067557112 | 0.7534877308 | 2.743982453 | 0.006069878238 | 0.07193326115 | 1.357988055 |
| ENSG000001532 | 38.07594955 | 1.965989961 | 0.7167993306 | 2.742734092 | 0.006093000318 | 0.0711745055 | 1.393563518 |
| ENSG000001300 | 11.49179835 | 3.808696919 | 1.389452339 | 2.741149741 | 0.006122459841 | 0.07136869844 | 1.393563518 |
| ENSG000001469 | 43.09379697 | -1.834194248 | 0.6693592007 | -2.740224152 | 0.006139729588 | 0.07147104515 | 1.393563518 |
| ENSG000001871 | 37.63888757 | 1.972439571 | 0.7199167536 | 2.739816181 | 0.006147355478 | 0.07566794103 | 1.278627818 |
| ENSG000001173 | 24.65889128 | 2.181799686 | 0.7963465729 | 2.739761506 | 0.006148378128 | 0.07147104515 | 1.393563518 |
| ENSG000001864 | 185.0753791 | 1.256341318 | 0.4586779372 | 2.73904894 | 0.006161720054 | 0.07811152826 | 1.22159078 |
| ENSG000001646 | 374.6055879 | -1.359654303 | 0.496532055 | -2.738301161 | 0.006175749313 | 0.07416867612 | 1.32213648 |
| ENSG000002726 | 33.40402582 | 2.502378305 | 0.9138887649 | 2.738165082 | 0.006178305416 | 0.07375116266 | 1.337480197 |
| ENSG000001262 | 7.219873592 | -5.316441049 | 1.941735643 | -2.737983963 | 0.006181709038 | 0.07297814852 | 1.357988055 |
| ENSG000001724 | 66.61615357 | 1.49189984 | 0.5450153487 | 2.737353807 | 0.006193564161 | 0.07306771262 | 1.357988055 |
| ENSG000002137 | 9.036130953 | 4.274158092 | 1.562462194 | 2.735527368 | 0.006228040711 | 0.07193326115 | 1.393563518 |
| ENSG000001031 | 109.7018935 | -1.133693156 | 0.4146297662 | -2.734230024 | 0.00625263472 | 0.08316070756 | 1.133466638 |
| ENSG000001374 | 153.18718 | 1.030091091 | 0.3768489731 | 2.733432128 | 0.006267803974 | 0.07877928185 | 1.22159078 |

| | | | | | | | |
|---|---|---|---|---|---|---|---|
| ENSG000001773 | 161.4469549 | 1.144386531 | 0.4188562891 | 2.732169864 | 0.006291869266 | 0.08026700352 | 1.198705341 |
| ENSG000001695 | 66.79554106 | 1.442613394 | 0.5282511597 | 2.730923288 | 0.006315717059 | 0.07401419333 | 1.357988055 |
| ENSG000002552 | 51.13760812 | 1.422886932 | 0.5210530743 | 2.730790781 | 0.006318256775 | 0.07401419333 | 1.357988055 |
| ENSG000001721 | 31.69817537 | 6.637252435 | 2.43058603 | 2.73072105 | 0.006319593665 | 0.07547843813 | 1.319539707 |
| ENSG000000852 | 64.97757848 | 1.599729985 | 0.5860774626 | 2.729553834 | 0.006342009332 | 0.0693469155 | 1.514054369 |
| ENSG000001791 | 46.5154349 | 1.50665033 | 0.5520889441 | 2.728999278 | 0.006352684304 | 0.07079049348 | 1.470104381 |
| ENSG000001080 | 12.48874816 | 2.851531567 | 1.046932651 | 2.723701056 | 0.006455490951 | 0.08029551285 | 1.227924265 |
| ENSG000002346 | 16.33209936 | 3.249686023 | 1.193198452 | 2.723508413 | 0.006459257042 | 0.07019826791 | 1.514850267 |
| ENSG000001155 | 11.29467774 | 4.795879584 | 1.761439153 | 2.722705225 | 0.006474980329 | 0.0753490991 | 1.357988055 |
| ENSG000000125 | 19.83328779 | 3.110930358 | 1.142730888 | 2.72236481 | 0.006481654692 | 0.07401419333 | 1.393563518 |
| ENSG000001143 | 83.7725393 | -1.443977612 | 0.5311761608 | -2.718453346 | 0.006558790454 | 0.07596291209 | 1.357988055 |
| ENSG000002560 | 186.5714518 | 1.08680048 | 0.3998537064 | 2.717995264 | 0.006567877821 | 0.08286802545 | 1.198705341 |
| ENSG000001000 | 147.6248802 | -1.194233897 | 0.4394573906 | -2.717519201 | 0.006577333884 | 0.06273378883 | 1.814765822 |
| ENSG000002535 | 46.29202825 | 1.938106625 | 0.7132107488 | 2.717438889 | 0.006578930338 | 0.07102223364 | 1.516424702 |
| ENSG000001307 | 62.05051216 | -1.666704902 | 0.6142111528 | -2.713569909 | 0.006656252204 | 0.0772803075 | 1.337480197 |
| ENSG000000998 | 427.2565555 | 1.162885089 | 0.428604753 | 2.713187571 | 0.006663937452 | 0.08150055069 | 1.242301963 |
| ENSG000002787 | 21.39649551 | 3.314723811 | 1.222158591 | 2.712187956 | 0.006684068055 | 0.07684196265 | 1.357988055 |
| ENSG000001539 | 68.43486511 | 1.398229486 | 0.5160611406 | 2.709426027 | 0.006739973283 | 0.07191245975 | 1.514054369 |
| ENSG000001761 | 7.942688681 | -4.856507196 | 1.792506245 | -2.709339066 | 0.006741740301 | 0.07714232234 | 1.357988055 |
| ENSG000002765 | 93.77756359 | -1.873612682 | 0.6916474703 | -2.708912795 | 0.006750408 | 0.07401419333 | 1.449168615 |
| ENSG000001721 | 117.5166001 | -1.36582247 | 0.5042531882 | -2.70860453 | 0.006756682418 | 0.0876632029 | 1.133466638 |
| ENSG000001967 | 12.16621626 | 3.285485505 | 1.213151326 | 2.708223974 | 0.006764435474 | 0.08316070756 | 1.227924265 |
| ENSG000001720 | 15.40640673 | 3.21748961 | 1.189774053 | 2.704286248 | 0.006845129391 | 0.07811152826 | 1.357988055 |
| ENSG000002848 | 73.19637051 | 1.355228198 | 0.5012189613 | 2.703864584 | 0.006853821414 | 0.07811152826 | 1.357988055 |
| ENSG000001539 | 30.1912729 | 2.319528844 | 0.8579398238 | 2.70360319 | 0.006859214665 | 0.07877928185 | 1.337480197 |
| ENSG000002789 | 8.319889229 | 4.592839367 | 1.699094467 | 2.703110073 | 0.006869399375 | 0.0840149521 | 1.227924265 |
| ENSG000002065 | 51.37536555 | 1.803824006 | 0.6674697927 | 2.70248036 | 0.006882425027 | 0.0782291841 | 1.357988055 |
| ENSG000001723 | 40.0578943 | 1.747195182 | 0.6467482554 | 2.701507375 | 0.006902594943 | 0.07840638307 | 1.357988055 |
| ENSG000002376 | 77.53268162 | -2.107818474 | 0.7808185799 | -2.699498357 | 0.006944409864 | 0.07867253864 | 1.357988055 |
| ENSG000001717 | 16.2915935 | 2.793812457 | 1.035845592 | 2.697132159 | 0.006993950701 | 0.0778811254 | 1.393563518 |
| ENSG000001097 | 41.78320329 | 1.988714841 | 0.7378363775 | 2.695333141 | 0.007031828698 | 0.07545536453 | 1.470104381 |
| ENSG000001302 | 76.95858453 | 1.391386427 | 0.5164137506 | 2.694324901 | 0.007053137475 | 0.07817552047 | 1.393563518 |
| ENSG000001653 | 9.58368367 | 3.816770461 | 1.416884246 | 2.693777189 | 0.007064737447 | 0.0707737271 | 1.636424459 |

| | | | | | | | |
|---|---|---|---|---|---|---|---|
| ENSG00000280[illegible] | 34.19313793 | -1.88726777 | 0.7008212169 | -2.692937549 | 0.007082553411 | 0.08345032202 | 1.278627818 |
| ENSG00000232[illegible] | 159.3918935 | 1.231295995 | 0.4575069235 | 2.691316637 | 0.007117060995 | 0.06639842724 | 1.814765822 |
| ENSG00000150[illegible] | 11.24417359 | -3.320751721 | 1.234004306 | -2.691037385 | 0.007123021228 | 0.08107483519 | 1.337480197 |
| ENSG00000230[illegible] | 147.9135192 | 1.206530049 | 0.448598611 | 2.689553689 | 0.007154763697 | 0.0665562733 | 1.814765822 |
| ENSG00000129[illegible] | 74.27093028 | 1.318269422 | 0.4909008738 | 2.685408587 | 0.007244118808 | 0.07965654192 | 1.393563518 |
| ENSG00000164[illegible] | 28.61678866 | -2.152837683 | 0.8021000812 | -2.684001328 | 0.007274681727 | 0.08316070756 | 1.319539707 |
| ENSG00000255[illegible] | 63.80202853 | 1.795203824 | 0.6688750171 | 2.683915198 | 0.007276556057 | 0.08150055069 | 1.357988055 |
| ENSG00000129[illegible] | 11.1699754 | 4.212803862 | 1.570424141 | 2.682589851 | 0.007305452453 | 0.07193326115 | 1.636424459 |
| ENSG00000204[illegible] | 9.609260295 | 3.857421742 | 1.438092532 | 2.682318179 | 0.007311388372 | 0.07193326115 | 1.636424459 |
| ENSG00000106[illegible] | 107.089419 | 1.444437411 | 0.5385576264 | 2.682048012 | 0.007317295724 | 0.0870940845 | 1.242301963 |
| ENSG00000275[illegible] | 9.779241491 | 4.137493217 | 1.542925733 | 2.681589353 | 0.007327334357 | 0.07193326115 | 1.636424459 |
| ENSG00000162[illegible] | 386.8463535 | 1.002585705 | 0.3739017786 | 2.681414645 | 0.007331161403 | 0.08709902328 | 1.242301963 |
| ENSG00000005[illegible] | 71.82027008 | 1.429486726 | 0.5331120005 | 2.681400389 | 0.007331473769 | 0.07817552047 | 1.449168615 |
| ENSG00000155[illegible] | 16.29570751 | 3.49434948 | 1.303583781 | 2.68057146 | 0.007349657017 | 0.07328537054 | 1.605577974 |
| ENSG00000101[illegible] | 767.2481435 | -1.088217424 | 0.4062477335 | -2.678703988 | 0.007390769903 | 0.07369817224 | 1.602205449 |
| ENSG00000236[illegible] | 45.01414519 | -2.073457236 | 0.7742342805 | -2.678074697 | 0.007404670291 | 0.07659226363 | 1.516424702 |
| ENSG00000110[illegible] | 26.72485413 | 2.437927403 | 0.9105831684 | 2.677325353 | 0.007421253128 | 0.08429294304 | 1.319539707 |
| ENSG00000117[illegible] | 24.38315529 | 1.962298532 | 0.7330700322 | 2.676822739 | 0.007432394536 | 0.0828304123 | 1.357988055 |
| ENSG00000198[illegible] | 110.5598108 | -1.227370955 | 0.4586360806 | -2.676132575 | 0.007447717806 | 0.08787372789 | 1.242301963 |
| ENSG00000161[illegible] | 74.60802327 | -1.396692928 | 0.522212176 | -2.674569825 | 0.007482519276 | 0.07692061821 | 1.514054369 |
| ENSG00000162[illegible] | 303.4945612 | -1.107997153 | 0.4143385836 | -2.674134625 | 0.007492236816 | 0.08817296271 | 1.242301963 |
| ENSG00000280[illegible] | 10.21108435 | 4.058495009 | 1.517720447 | 2.674072829 | 0.007493617572 | 0.08164654283 | 1.393563518 |
| ENSG00000178[illegible] | 36.47507773 | 1.640016858 | 0.6134426986 | 2.673463816 | 0.007507237453 | 0.08174154699 | 1.393563518 |
| ENSG00000138[illegible] | 61.75932371 | -1.506384695 | 0.5636475782 | -2.672564831 | 0.007527382779 | 0.07714232234 | 1.516424702 |
| ENSG00000236[illegible] | 22.76043801 | -2.40715766 | 0.9007444347 | -2.67240914 | 0.007530876568 | 0.08499011101 | 1.319539707 |
| ENSG00000198[illegible] | 36.17076303 | 2.272424603 | 0.8509646427 | 2.670410131 | 0.007575864846 | 0.08473095018 | 1.337480197 |
| ENSG00000166[illegible] | 90.66042015 | 1.38631024 | 0.5193156259 | 2.669494564 | 0.007596550273 | 0.08255236158 | 1.393563518 |
| ENSG00000168[illegible] | 7.954926905 | 4.260887456 | 1.596785179 | 2.66841621 | 0.007620978493 | 0.08421929502 | 1.357988055 |
| ENSG00000130[illegible] | 14.76004371 | 2.798624818 | 1.048989864 | 2.667923603 | 0.007632161034 | 0.08499011101 | 1.337480197 |
| ENSG00000114[illegible] | 439.4438209 | 1.083412833 | 0.4062327992 | 2.666975278 | 0.007653730157 | 0.07666689428 | 1.564848717 |
| ENSG00000174[illegible] | 262.74556 | 1.265563925 | 0.4746878896 | 2.666096929 | 0.007673756446 | 0.09169069917 | 1.198705341 |
| ENSG00000139[illegible] | 25.16241899 | 2.511072868 | 0.9419249893 | 2.665894733 | 0.007678373134 | 0.08463500823 | 1.357988055 |
| ENSG00000278[illegible] | 19.18136218 | -2.812427759 | 1.055027547 | -2.665738698 | 0.007681937567 | 0.07545536453 | 1.605577974 |

| | | | | | | | |
|---|---|---|---|---|---|---|---|
| ENSG00000130 | 173.906061 | -1.100192348 | 0.412860572 | -2.664803624 | 0.007703329184 | 0.09003434032 | 1.242301963 |
| ENSG00000060 | 117.2823734 | 1.565449585 | 0.5876550396 | 2.663892044 | 0.007724234706 | 0.09106980747 | 1.22159078 |
| ENSG00000104 | 85.7704643 | -1.417667915 | 0.5323737946 | -2.66291829 | 0.007746622247 | 0.08499011101 | 1.357988055 |
| ENSG00000143 | 110.3480405 | 1.224353883 | 0.4602294582 | 2.660311854 | 0.007806833046 | 0.08736439464 | 1.318051683 |
| ENSG00000007 | 105.915749 | -1.734070846 | 0.6519549192 | -2.659801767 | 0.007818665432 | 0.09169069917 | 1.22159078 |
| ENSG00000112 | 11.92338491 | 3.871340543 | 1.455702495 | 2.659431139 | 0.007827272879 | 0.07545536453 | 1.636424459 |
| ENSG00000267 | 7.444925948 | 4.27044861 | 1.606009555 | 2.659043091 | 0.007836293968 | 0.08429294304 | 1.393563518 |
| ENSG00000157 | 10.78128671 | 3.828730105 | 1.440471556 | 2.657969947 | 0.007861290293 | 0.07565885207 | 1.636424459 |
| ENSG00000234 | 13.79312005 | 2.739228448 | 1.030623214 | 2.657836937 | 0.007864393397 | 0.07672698994 | 1.605577974 |
| ENSG00000185 | 17.70619525 | 3.686870593 | 1.387984692 | 2.65627612 | 0.007900889289 | 0.07684196265 | 1.605577974 |
| ENSG00000058 | 129.5679647 | 1.016441858 | 0.38268812 | 2.656058041 | 0.007906000595 | 0.09143224583 | 1.242301963 |
| ENSG00000277 | 14.55153283 | -8.032183777 | 3.024797224 | -2.655445368 | 0.007920376174 | 0.07692061821 | 1.605577974 |
| ENSG00000105 | 9.845580834 | -3.980729607 | 1.499384772 | -2.654908655 | 0.007932988701 | 0.08499011101 | 1.393563518 |
| ENSG00000277 | 11.24374281 | 4.381929892 | 1.650738331 | 2.654527256 | 0.007941962325 | 0.08499011101 | 1.393563518 |
| ENSG00000203 | 18.77725192 | 3.101417774 | 1.168610232 | 2.653936865 | 0.007955871111 | 0.08026700352 | 1.514850267 |
| ENSG00000116 | 108.9539666 | 1.103834577 | 0.4164929552 | 2.650307917 | 0.008041844121 | 0.09348231495 | 1.22159078 |
| ENSG00000123 | 43.51006076 | -1.752056631 | 0.6611565837 | -2.649987422 | 0.008049476755 | 0.08810042945 | 1.337480197 |
| ENSG00000231 | 133.6492142 | 1.25928919 | 0.4752809868 | 2.649567782 | 0.008059480349 | 0.09760386879 | 1.130447911 |
| ENSG00000172 | 51.23472352 | 1.54354408 | 0.5828317381 | 2.648352825 | 0.00808850593 | 0.08834535985 | 1.337480197 |
| ENSG00000181 | 10.91786197 | -3.682823436 | 1.390896545 | -2.647805439 | 0.008101613672 | 0.0876632029 | 1.357988055 |
| ENSG00000138 | 38.49720154 | 1.99035909 | 0.7524881864 | 2.645036993 | 0.008168198652 | 0.08678568668 | 1.393563518 |
| ENSG00000265 | 14.29813812 | 3.680198921 | 1.391756009 | 2.644284556 | 0.008186380273 | 0.08817296271 | 1.357988055 |
| ENSG00000148 | 32.08387025 | 1.866188177 | 0.70580578 | 2.644053407 | 0.008191972942 | 0.08936216705 | 1.337480197 |
| ENSG00000169 | 29.90727818 | 8.00405427 | 3.028058091 | 2.643296142 | 0.008210318984 | 0.09011932818 | 1.319539707 |
| ENSG00000112 | 27.1001748 | -2.169878398 | 0.8214810762 | -2.641422256 | 0.008255875247 | 0.0897378233 | 1.337480197 |
| ENSG00000111 | 27.07880379 | 2.10679761 | 0.7977427092 | 2.640948749 | 0.008267422464 | 0.08745001783 | 1.393563518 |
| ENSG00000152 | 7.764665562 | 4.254021075 | 1.611389584 | 2.639970569 | 0.008291322734 | 0.09003434032 | 1.337480197 |
| ENSG00000158 | 35.53050244 | 2.341545712 | 0.8872128525 | 2.639215275 | 0.008309819387 | 0.0876632029 | 1.393563518 |
| ENSG00000145 | 172.6839767 | -1.490926487 | 0.5650062287 | -2.63877885 | 0.008320523964 | 0.07335056893 | 1.814765822 |
| ENSG00000166 | 13.79069883 | 2.990005784 | 1.133194048 | 2.638564675 | 0.008325781753 | 0.09011932818 | 1.337480197 |
| ENSG00000184 | 194.5360107 | 1.442700358 | 0.5468443692 | 2.638228424 | 0.008334042355 | 0.09628141467 | 1.198705341 |
| ENSG00000237 | 8.481511461 | 3.629854948 | 1.375962939 | 2.638047033 | 0.008338501604 | 0.07853506831 | 1.636424459 |
| ENSG00000060 | 83.35767599 | 1.544233662 | 0.5856666692 | 2.636710851 | 0.008371415561 | 0.08799205073 | 1.393563518 |

| | | | | | | | |
|---|---|---|---|---|---|---|---|
| ENSG00000128 | 28.51148789 | 2.224869462 | 0.8447027092 | 2.633908282 | 0.008440828544 | 0.09357187138 | 1.278627818 |
| ENSG00000125 | 768.8952749 | -1.100703332 | 0.4181067013 | -2.632589548 | 0.008473668207 | 0.0807237968 | 1.602205449 |
| ENSG00000198 | 645.1352357 | 1.151895336 | 0.4376664656 | 2.631902205 | 0.008490829946 | 0.07401419333 | 1.825593782 |
| ENSG00000214 | 306.8253789 | 1.043544613 | 0.3967230127 | 2.630411092 | 0.008528167309 | 0.09591762548 | 1.242301963 |
| ENSG00000089 | 366.4412717 | -1.212334135 | 0.4610296531 | -2.629622903 | 0.008547962761 | 0.09600827935 | 1.242301963 |
| ENSG00000125 | 97.70453479 | -1.630797576 | 0.6208482784 | -2.626724809 | 0.008621102388 | 0.09690101633 | 1.22159078 |
| ENSG00000121 | 7.439572106 | 3.755553194 | 1.430154942 | 2.625976447 | 0.008640079574 | 0.09229773899 | 1.337480197 |
| ENSG00000178 | 125.991767 | -1.390476936 | 0.5298530249 | -2.624269128 | 0.008683514063 | 0.09748735708 | 1.22159078 |
| ENSG00000153 | 22.52703551 | 2.1287922 | 0.8112955807 | 2.62394157 | 0.008691869485 | 0.09016720698 | 1.393563518 |
| ENSG00000180 | 61.52583278 | -2.122497805 | 0.810349825 | -2.619236457 | 0.008812683784 | 0.09350937679 | 1.337480197 |
| ENSG00000162 | 27.85233088 | 2.163571292 | 0.8261820101 | 2.618758658 | 0.008825035871 | 0.09357187138 | 1.337480197 |
| ENSG00000146 | 12.97839206 | 4.074548026 | 1.556282454 | 2.618128872 | 0.008841340746 | 0.09120956684 | 1.393563518 |
| ENSG00000135 | 99.84954879 | -1.15263053 | 0.4402551128 | -2.618096865 | 0.008842170122 | 0.09452680217 | 1.318051683 |
| ENSG00000239 | 38.48094139 | 2.144760087 | 0.8192531055 | 2.617945629 | 0.008846089895 | 0.09369224389 | 1.337480197 |
| ENSG00000152 | 15.18745511 | 3.365713715 | 1.285683563 | 2.617839889 | 0.008848831414 | 0.08660009901 | 1.514850267 |
| ENSG00000268 | 11.75280048 | -3.368530716 | 1.28784102 | -2.615641733 | 0.008905995036 | 0.09421189908 | 1.337480197 |
| ENSG00000241 | 74.0486704 | 1.478052224 | 0.5651383843 | 2.615381054 | 0.00891279587 | 0.08956700827 | 1.449168615 |
| ENSG00000189 | 23.67568842 | 2.273290067 | 0.8696464023 | 2.614039523 | 0.008947868501 | 0.09185180452 | 1.393563518 |
| ENSG00000164 | 522.6895204 | 1.051001703 | 0.4022700371 | 2.612677072 | 0.008983614177 | 0.07684196265 | 1.825593782 |
| ENSG00000282 | 24.38887803 | -2.591548394 | 0.9922737405 | -2.611727276 | 0.009008608665 | 0.09566148218 | 1.319539707 |
| ENSG00000139 | 22.98988337 | 2.229227813 | 0.8539069062 | 2.610621599 | 0.009037783486 | 0.09585572814 | 1.319539707 |
| ENSG00000257 | 8.710010947 | 4.566949182 | 1.749702665 | 2.610128722 | 0.009050815898 | 0.09424063446 | 1.357988055 |
| ENSG00000159 | 287.0532259 | 1.11543493 | 0.4274835289 | 2.609305048 | 0.009072632531 | 0.09570107966 | 1.327567552 |
| ENSG00000161 | 335.3941386 | 1.12097143 | 0.4297585206 | 2.608375114 | 0.009097320085 | 0.09892259076 | 1.249985076 |
| ENSG00000180 | 23.8365097 | -2.410576683 | 0.9243783399 | -2.607781445 | 0.009113111943 | 0.09613716887 | 1.319539707 |
| ENSG00000153 | 468.0467095 | 1.07613851 | 0.4126806524 | 2.60767861 | 0.009115849891 | 0.08499011101 | 1.602205449 |
| ENSG00000100 | 53.84808268 | 1.761673261 | 0.6758706608 | 2.60652424 | 0.00914663496 | 0.09011932818 | 1.470104381 |
| ENSG00000006 | 11.30559723 | -3.654911154 | 1.402687686 | -2.605648563 | 0.009170049638 | 0.09348231495 | 1.393563518 |
| ENSG00000160 | 133.522823 | -1.305399108 | 0.5010101585 | -2.605534212 | 0.009173111212 | 0.09991128882 | 1.242301963 |
| ENSG00000143 | 77.26414576 | 1.919676042 | 0.7368307763 | 2.605314684 | 0.009178991276 | 0.09116995575 | 1.449168615 |
| ENSG00000183 | 14.15589504 | 3.109147006 | 1.193386473 | 2.605314436 | 0.009178997942 | 0.09482547951 | 1.357988055 |
| ENSG00000274 | 27.43539377 | -2.400315395 | 0.9214615072 | -2.604900342 | 0.009190098641 | 0.09827209926 | 1.278627818 |
| ENSG00000184 | 9.361995736 | 3.819396106 | 1.4675157 | 2.602627084 | 0.0092512521 | 0.09616678908 | 1.337480197 |

| | | | | | | | |
|---|---|---|---|---|---|---|---|
| ENSG00000125 | 426.7323281 | -1.031870736 | 0.3967105578 | -2.601066988 | 0.009293430492 | 0.09679950064 | 1.32213648 |
| ENSG00000173 | 120.3288322 | 1.142370642 | 0.4392013331 | 2.601018158 | 0.009294753426 | 0.09688427813 | 1.318051683 |
| ENSG00000203 | 59.35839814 | -1.392279821 | 0.5353355849 | -2.600760832 | 0.009301727758 | 0.0943233436 | 1.393563518 |
| ENSG00000071 | 53.47717077 | -2.066246134 | 0.7947329293 | -2.599925154 | 0.009324409507 | 0.09444182215 | 1.393563518 |
| ENSG00000171 | 23.03148553 | 2.587970551 | 0.9954065278 | 2.59991318 | 0.009324734856 | 0.09444182215 | 1.393563518 |
| ENSG00000287 | 20.44041595 | 3.41526766 | 1.313983238 | 2.599171405 | 0.009344909869 | 0.08645251004 | 1.605577974 |
| ENSG00000229 | 80.45415441 | -1.816691897 | 0.698998744 | -2.598991647 | 0.009349804838 | 0.0960106101 | 1.357988055 |
| ENSG00000111 | 37.3528979 | 1.871964965 | 0.7203843972 | 2.598564005 | 0.009361459092 | 0.09452680217 | 1.393563518 |
| ENSG00000078 | 79.56632506 | 1.359347594 | 0.5236602252 | 2.595858018 | 0.009435504621 | 0.09014798375 | 1.514054369 |
| ENSG00000198 | 24.66603308 | 3.171225109 | 1.221657924 | 2.59583722 | 0.009436075747 | 0.09628141467 | 1.357988055 |
| ENSG00000225 | 33.16990528 | 2.352088867 | 0.9062704213 | 2.595349921 | 0.00944946607 | 0.09628141467 | 1.357988055 |
| ENSG00000199 | 11.92799463 | -4.371687654 | 1.684572839 | -2.595131271 | 0.009455479793 | 0.09628141467 | 1.357988055 |
| ENSG00000188 | 59.91470101 | -1.800180691 | 0.6938069956 | -2.594641886 | 0.009468952165 | 0.09208342048 | 1.470104381 |
| ENSG00000164 | 62.32757074 | -1.873300421 | 0.7221140231 | -2.59418923 | 0.009481428642 | 0.09539172883 | 1.393563518 |
| ENSG00000214 | 25.30528939 | -2.51855446 | 0.9709438013 | -2.593924033 | 0.00948874503 | 0.09827209926 | 1.319539707 |
| ENSG00000053 | 12.9092331 | 3.153999813 | 1.216026874 | 2.593692525 | 0.009495136106 | 0.09045200516 | 1.514850267 |
| ENSG00000144 | 9.265854779 | 3.521108831 | 1.3583479 | 2.592199561 | 0.009536443553 | 0.08650614448 | 1.636424459 |
| ENSG00000066 | 78.35747054 | 1.704056515 | 0.6574676102 | 2.591848615 | 0.009546176793 | 0.09679950064 | 1.357988055 |
| ENSG00000163 | 40.14090417 | -1.577123206 | 0.6086928603 | -2.591000008 | 0.009569748916 | 0.09591762548 | 1.393563518 |
| ENSG00000014 | 92.23282486 | -1.088792396 | 0.4204322305 | -2.589697737 | 0.009606023623 | 0.09120956684 | 1.514054369 |
| ENSG00000084 | 1023.553812 | 6.879951495 | 2.657747764 | 2.588639746 | 0.009635584124 | 0.08936216705 | 1.572476891 |
| ENSG00000260 | 17.9101859 | 2.624506747 | 1.013869377 | 2.588604417 | 0.009636572621 | 0.09616678908 | 1.393563518 |
| ENSG00000132 | 546.1222426 | -1.037272913 | 0.4008986222 | -2.587369612 | 0.009671179013 | 0.0840149521 | 1.729500602 |
| ENSG00000175 | 18.7029972 | 3.203926447 | 1.238636971 | 2.586654946 | 0.009691258682 | 0.09169069917 | 1.514850267 |
| ENSG00000162 | 8.99672733 | 3.841942383 | 1.485455619 | 2.586373052 | 0.009699189167 | 0.09628141467 | 1.393563518 |
| ENSG00000198 | 20.43044517 | 2.982444479 | 1.153249902 | 2.586121597 | 0.009706268167 | 0.09892259076 | 1.337480197 |
| ENSG00000135 | 77.10528702 | -1.781174801 | 0.6887636568 | -2.586046438 | 0.009708384966 | 0.09628141467 | 1.393563518 |
| ENSG00000139 | 85.84098335 | 1.153639995 | 0.4461118921 | 2.585987989 | 0.00971003141 | 0.09892259076 | 1.337480197 |
| ENSG00000188 | 16.46444155 | -2.919545794 | 1.129020072 | -2.585911329 | 0.009712191213 | 0.09892259076 | 1.337480197 |
| ENSG00000173 | 46.01957063 | -2.252663834 | 0.8712548269 | -2.585539574 | 0.009722671088 | 0.09892259076 | 1.337480197 |
| ENSG00000064 | 44.89665861 | 1.502864705 | 0.5812986253 | 2.585357405 | 0.00972781016 | 0.09182405024 | 1.516424702 |
| ENSG00000162 | 45.26995204 | 1.845657616 | 0.7139429482 | 2.58516121 | 0.00973334761 | 0.09647162254 | 1.393563518 |
| ENSG00000196 | 37.30028001 | -1.5904475 | 0.6152632643 | -2.584986935 | 0.00973826873 | 0.09892259076 | 1.337480197 |

| | | | | | | | |
|---|---|---|---|---|---|---|---|
| ENSG000001656 | 18.42529971 | 2.467801605 | 0.954667794 | 2.584984662 | 0.009738332943 | 0.09892259076 | 1.337480197 |
| ENSG000002749 | 12.214521 | 3.117860939 | 1.20627893 | 2.584693193 | 0.009746568381 | 0.0876632029 | 1.636424459 |
| ENSG000002879 | 15.24606691 | 2.73953239 | 1.061673385 | 2.580390947 | 0.00986885208 | 0.08956700827 | 1.605577974 |
| ENSG000001286 | 27.1332597 | 2.097016133 | 0.8134048341 | 2.578071884 | 0.009935332727 | 0.09934155163 | 1.357988055 |
| ENSG000001961 | 20.5871052 | 2.266920092 | 0.8801064712 | 2.575733921 | 0.01000275874 | 0.09972448049 | 1.357988055 |
| ENSG000001149 | 67.46008512 | -1.452191325 | 0.5640541642 | -2.574560064 | 0.01003676581 | 0.09619369307 | 1.449168615 |
| ENSG000001745 | 11.62485619 | 3.482246675 | 1.353148989 | 2.573439217 | 0.01006933321 | 0.08956700827 | 1.636424459 |
| ENSG000002603 | 32.82933725 | -2.183008311 | 0.8484197285 | -2.573028699 | 0.01008128477 | 0.09874672472 | 1.393563518 |
| ENSG000001625 | 62.80490503 | -1.499990782 | 0.5840177938 | -2.56839911 | 0.01021694481 | 0.09469192067 | 1.516424702 |
| ENSG000002791 | 13.1109813 | -3.018050541 | 1.175185179 | -2.568148915 | 0.0102243223 | 0.09474421998 | 1.514850267 |
| ENSG000001629 | 83.52037986 | -1.709808688 | 0.6665939436 | -2.564992834 | 0.01031779359 | 0.09758662565 | 1.449168615 |
| ENSG000001872 | 70.47262775 | -7.482405431 | 2.921934609 | -2.560771007 | 0.01044401709 | 0.09613428269 | 1.514054369 |
| ENSG000001471 | 21.0354057 | 3.131794752 | 1.225918599 | 2.554651471 | 0.01062941706 | 0.09372449874 | 1.605577974 |
| ENSG000000930 | 78.4192381 | 1.183904693 | 0.4634833467 | 2.554362959 | 0.01063822975 | 0.09956068168 | 1.449168615 |
| ENSG000000970 | 62.39617917 | 1.447538045 | 0.5668891855 | 2.553476203 | 0.01066535666 | 0.09679950064 | 1.516424702 |
| ENSG000001332 | 8.90164748 | 3.298926155 | 1.292436941 | 2.552485194 | 0.01069574554 | 0.09304159544 | 1.636424459 |
| ENSG000001703 | 20.67199763 | 2.300616785 | 0.9050410404 | 2.542002718 | 0.01102193163 | 0.09892259076 | 1.514850267 |
| ENSG000002065 | 17.02886249 | 2.502243626 | 0.9899840563 | 2.52755952 | 0.01148583535 | 0.09787821473 | 1.605577974 |
| ENSG000001677 | 483.3948409 | -1.206070046 | 0.4818713603 | -2.502888001 | 0.01231845191 | 0.09474421998 | 1.825593782 |

| gene_name | genes_description |
|---|---|
| DPT | dermatopontin |
| MYH11 | myosin heavy chain 11 |
| DES | desmin |
| SYNPO2 | synaptopodin 2 |
| SYNM | synemin |
| CNN1 | calponin 1 |
| ACTG2 | actin gamma 2, smooth muscle |
| FABP4 | fatty acid binding protein 4 |
| MGP | matrix Gla protein |
| HSPB6 | heat shock protein family B (small) member 6 |
| TAGLN | transgelin |
| MYL9 | myosin light chain 9 |
| LMOD1 | leiomodin 1 |
| SPEG | striated muscle enriched protein kinase |
| PSD | pleckstrin and Sec7 domain containing |
| PRELP | proline and arginine rich end leucine rich repeat protein |
| PGM5 | phosphoglucomutase 5 |
| CLU | clusterin |
| ANK2 | ankyrin 2 |
| FXYD6 | FXYD domain containing ion transport regulator 6 |
| CCDC80 | coiled-coil domain containing 80 |
| FLNC | filamin C |
| PLN | phospholamban |
| DTNA | dystrobrevin alpha |
| NEXN | nexilin F-actin binding protein |
| PTGIS | prostaglandin I2 synthase |
| SORBS1 | sorbin and SH3 domain containing 1 |
| AP000892.6 | NA |
| SLIT3 | slit guidance ligand 3 |
| TPM2 | tropomyosin 2 |
| CEMIP | cell migration inducing hyaluronidase 1 |
| PLIN4 | perilipin 4 |

| | |
|---|---|
| PPP1R1A | protein phosphatase 1 regulatory inhibitor subunit 1A |
| TNS1 | tensin 1 |
| NIBAN1 | niban apoptosis regulator 1 |
| SFRP1 | secreted frizzled related protein 1 |
| RBPMS2 | RNA binding protein, mRNA processing factor 2 |
| MYLK | myosin light chain kinase |
| ADH1B | alcohol dehydrogenase 1B (class I), beta polypeptide |
| PDLIM3 | PDZ and LIM domain 3 |
| HAND2-AS1 | HAND2 antisense RNA 1 |
| ADIPOQ | adiponectin, C1Q and collagen domain containing |
| FLNA | filamin A |
| RAMP1 | receptor activity modifying protein 1 |
| LMO3 | LIM domain only 3 |
| AOC3 | amine oxidase copper containing 3 |
| APOD | apolipoprotein D |
| SFRP4 | secreted frizzled related protein 4 |
| GPM6A | glycoprotein M6A |
| ASB5 | ankyrin repeat and SOCS box containing 5 |
| HAND1 | heart and neural crest derivatives expressed 1 |
| CHRDL1 | chordin like 1 |
| CXCL12 | C-X-C motif chemokine ligand 12 |
| AF001548.3 | NA |
| MIR99AHG | mir-99a-let-7c cluster host gene |
| NXPH3 | neurexophilin 3 |
| KY | kyphoscoliosis peptidase |
| SLCO4A1 | solute carrier organic anion transporter family member 4A1 |
| FBXO32 | F-box protein 32 |
| CASQ2 | calsequestrin 2 |
| CARMN | NA |
| CCDC69 | coiled-coil domain containing 69 |
| JPH2 | junctophilin 2 |
| AHNAK2 | AHNAK nucleoprotein 2 |
| CTD-3014M21.1 | NA |

| | |
|---|---|
| RSPO2 | R-spondin 2 |
| DDR2 | discoidin domain receptor tyrosine kinase 2 |
| PCOLCE2 | procollagen C-endopeptidase enhancer 2 |
| REEP1 | receptor accessory protein 1 |
| IGFBP6 | insulin like growth factor binding protein 6 |
| FBLN1 | fibulin 1 |
| LDB3 | LIM domain binding 3 |
| RBFOX3 | RNA binding fox-1 homolog 3 |
| NRP2 | neuropilin 2 |
| ATP1A2 | ATPase Na+/K+ transporting subunit alpha 2 |
| ATP2B4 | ATPase plasma membrane Ca2+ transporting 4 |
| MSRB3 | methionine sulfoxide reductase B3 |
| RP11-248G5.8 | NA |
| KCNMB1 | potassium calcium-activated channel subfamily M regulatory beta subunit 1 |
| CALD1 | caldesmon 1 |
| CCBE1 | collagen and calcium binding EGF domains 1 |
| IL11 | interleukin 11 |
| APOA4 | apolipoprotein A4 |
| PPP1R12B | protein phosphatase 1 regulatory subunit 12B |
| MAP1B | microtubule associated protein 1B |
| DPYSL3 | dihydropyrimidinase like 3 |
| RNVU1-27 | NA |
| ITGA7 | integrin subunit alpha 7 |
| U2 | NA |
| SMTN | smoothelin |
| CPXM2 | carboxypeptidase X, M14 family member 2 |
| MRPS31P5 | mitochondrial ribosomal protein S31 pseudogene 5 |
| C3 | complement C3 |
| ACTA2 | actin alpha 2, smooth muscle |
| CFD | complement factor D |
| MEIS1 | Meis homeobox 1 |
| NOVA1 | NOVA alternative splicing regulator 1 |
| GRIN2D | glutamate ionotropic receptor NMDA type subunit 2D |

| | |
|---|---|
| LIMS2 | LIM zinc finger domain containing 2 |
| HSPB7 | heat shock protein family B (small) member 7 |
| THBS4 | thrombospondin 4 |
| JAM3 | junctional adhesion molecule 3 |
| PABPN1 | poly(A) binding protein nuclear 1 |
| SLC2A4 | solute carrier family 2 member 4 |
| PHLDB2 | pleckstrin homology like domain family B member 2 |
| CSRP1 | cysteine and glycine rich protein 1 |
| PRSS22 | serine protease 22 |
| DCN | decorin |
| PDZRN4 | PDZ domain containing ring finger 4 |
| RNU12 | NA |
| MMP12 | matrix metallopeptidase 12 |
| SVIL | supervillin |
| DACT3 | dishevelled binding antagonist of beta catenin 3 |
| FXYD1 | FXYD domain containing ion transport regulator 1 |
| LIFR | LIF receptor subunit alpha |
| AR | androgen receptor |
| MMP10 | matrix metallopeptidase 10 |
| SCN7A | sodium voltage-gated channel alpha subunit 7 |
| EFEMP1 | EGF containing fibulin extracellular matrix protein 1 |
| PHYHIP | phytanoyl-CoA 2-hydroxylase interacting protein |
| C4BPB | complement component 4 binding protein beta |
| SGCD | sarcoglycan delta |
| GPC3 | glypican 3 |
| PPP1R14A | protein phosphatase 1 regulatory inhibitor subunit 14A |
| TCEAL2 | transcription elongation factor A like 2 |
| TMEM35A | transmembrane protein 35A |
| IL1B | interleukin 1 beta |
| OGN | osteoglycin |
| ACACB | acetyl-CoA carboxylase beta |
| TNFSF12 | TNF superfamily member 12 |
| HSPH1 | heat shock protein family H (Hsp110) member 1 |

| | |
|---|---|
| SCARNA9 | small Cajal body-specific RNA 9 |
| NCKAP1L | NCK associated protein 1 like |
| CBX6 | chromobox 6 |
| ADCY5 | adenylate cyclase 5 |
| RAI2 | retinoic acid induced 2 |
| CCL14 | C-C motif chemokine ligand 14 |
| AKAP12 | A-kinase anchoring protein 12 |
| MFAP5 | microfibril associated protein 5 |
| CRYAB | crystallin alpha B |
| NPTXR | neuronal pentraxin receptor |
| FHL1 | four and a half LIM domains 1 |
| KCNB1 | potassium voltage-gated channel subfamily B member 1 |
| MUC5AC | mucin 5AC, oligomeric mucus/gel-forming |
| PDE5A | phosphodiesterase 5A |
| SPINK1 | serine peptidase inhibitor Kazal type 1 |
| PVRIG | PVR related immunoglobulin domain containing |
| SVEP1 | sushi, von Willebrand factor type A, EGF and pentraxin domain containing 1 |
| SLC26A10 | solute carrier family 26 member 10 |
| PTGER3 | prostaglandin E receptor 3 |
| CCN5 | cellular communication network factor 5 |
| SLC35F1 | solute carrier family 35 member F1 |
| MAPT | microtubule associated protein tau |
| CTSG | cathepsin G |
| DUSP1 | dual specificity phosphatase 1 |
| AMOTL1 | angiomotin like 1 |
| PPP1R3C | protein phosphatase 1 regulatory subunit 3C |
| CXCL1 | C-X-C motif chemokine ligand 1 |
| ADAMTSL3 | ADAMTS like 3 |
| GNAO1 | G protein subunit alpha o1 |
| SPARCL1 | SPARC like 1 |
| FN1 | fibronectin 1 |
| MAB21L2 | mab-21 like 2 |
| CNTNAP3B | contactin associated protein family member 3B |

| | |
|---|---|
| SGCA | sarcoglycan alpha |
| PRIMA1 | proline rich membrane anchor 1 |
| MEIS2 | Meis homeobox 2 |
| CAV1 | caveolin 1 |
| PI16 | peptidase inhibitor 16 |
| DMD | dystrophin |
| BOC | BOC cell adhesion associated, oncogene regulated |
| COL14A1 | collagen type XIV alpha 1 chain |
| CH17-3B23.3 | NA |
| FOXP2 | forkhead box P2 |
| MITF | melanocyte inducing transcription factor |
| BMP3 | bone morphogenetic protein 3 |
| SLMAP | sarcolemma associated protein |
| LL22NC03-2H8. | NA |
| RERG | RAS like estrogen regulated growth inhibitor |
| WDR74 | WD repeat domain 74 |
| ZBTB16 | zinc finger and BTB domain containing 16 |
| GNAZ | G protein subunit alpha z |
| RP11-799B12.4 | NA |
| NNAT | neuronatin |
| MYOC | myocilin |
| PLAC9 | placenta associated 9 |
| KCNAB1 | potassium voltage-gated channel subfamily A regulatory beta subunit 1 |
| ABI3BP | ABI family member 3 binding protein |
| SNORD3B-2 | NA |
| CADM3 | cell adhesion molecule 3 |
| RBPMS | RNA binding protein, mRNA processing factor |
| UCHL1 | ubiquitin C-terminal hydrolase L1 |
| ADAMTS9-AS2 | ADAMTS9 antisense RNA 2 |
| MIR100HG | mir-100-let-7a-2-mir-125b-1 cluster host gene |
| DNAJB5 | DnaJ heat shock protein family (Hsp40) member B5 |
| CILP2 | cartilage intermediate layer protein 2 |
| NCAM1 | neural cell adhesion molecule 1 |

| | |
|---|---|
| KANK2 | KN motif and ankyrin repeat domains 2 |
| MBNL1-AS1 | MBNL1 antisense RNA 1 |
| FAM157A | family with sequence similarity 157 member A |
| LEPR | leptin receptor |
| PRNP | prion protein |
| KCNMA1 | potassium calcium-activated channel subfamily M alpha 1 |
| CYS1 | cystin 1 |
| ABCA9 | ATP binding cassette subfamily A member 9 |
| ACKR1 | atypical chemokine receptor 1 (Duffy blood group) |
| CH17-373J23.1 | NA |
| MYOM1 | myomesin 1 |
| H2AC18 | H2A clustered histone 18 |
| MAMDC2 | MAM domain containing 2 |
| CXCL13 | C-X-C motif chemokine ligand 13 |
| PBX3 | PBX homeobox 3 |
| MYBL2 | MYB proto-oncogene like 2 |
| CILP | cartilage intermediate layer protein |
| FNBP1 | formin binding protein 1 |
| STARD9 | StAR related lipid transfer domain containing 9 |
| SERTAD4-AS1 | NA |
| CDKN1C | cyclin dependent kinase inhibitor 1C |
| CTSF | cathepsin F |
| RP11-327J17.7 | NA |
| ESR1 | estrogen receptor 1 |
| FLRT2 | fibronectin leucine rich transmembrane protein 2 |
| SNHG22 | small Cajal body-specific RNA 17 |
| PDK4 | pyruvate dehydrogenase kinase 4 |
| OMD | osteomodulin |
| TRPS1 | transcriptional repressor GATA binding 1 |
| CTA-363E6.3 | NA |
| CEACAM7 | CEA cell adhesion molecule 7 |
| CEACAM6 | CEA cell adhesion molecule 6 |
| SAA1 | serum amyloid A1 |

| | |
|---|---|
| ADAM33 | ADAM metallopeptidase domain 33 |
| SCARA3 | scavenger receptor class A member 3 |
| ARMCX4 | armadillo repeat containing X-linked 4 |
| SRPX | sushi repeat containing protein X-linked |
| IRAG1 | inositol 1,4,5-triphosphate receptor associated 1 |
| TMEM200B | transmembrane protein 200B |
| CDH19 | cadherin 19 |
| ZFHX4 | zinc finger homeobox 4 |
| MFAP4 | microfibril associated protein 4 |
| SLC17A9 | solute carrier family 17 member 9 |
| SSBP2 | single stranded DNA binding protein 2 |
| RUNX1T1 | RUNX1 partner transcriptional co-repressor 1 |
| LPP | LIM domain containing preferred translocation partner in lipoma |
| C14orf132 | chromosome 14 open reading frame 132 |
| SERPINB5 | serpin family B member 5 |
| RGMA | repulsive guidance molecule BMP co-receptor a |
| SPIB | Spi-B transcription factor |
| ANKRD22 | ankyrin repeat domain 22 |
| PHLDA3 | pleckstrin homology like domain family A member 3 |
| ARID5A | AT-rich interaction domain 5A |
| ASB2 | ankyrin repeat and SOCS box containing 2 |
| CDON | cell adhesion associated, oncogene regulated |
| CFL2 | cofilin 2 |
| GFRA3 | GDNF family receptor alpha 3 |
| LONRF2 | LON peptidase N-terminal domain and ring finger 2 |
| IL24 | interleukin 24 |
| CLDN11 | claudin 11 |
| ITIH5 | inter-alpha-trypsin inhibitor heavy chain 5 |
| SLC6A14 | solute carrier family 6 member 14 |
| PRRT2 | proline rich transmembrane protein 2 |
| OLFM4 | olfactomedin 4 |
| DUOXA2 | dual oxidase maturation factor 2 |
| PGM5-AS1 | PGM5 antisense RNA 1 |

| | |
|---|---|
| PCDH9 | protocadherin 9 |
| FILIP1 | filamin A interacting protein 1 |
| BHMT2 | betaine--homocysteine S-methyltransferase 2 |
| RIC3 | RIC3 acetylcholine receptor chaperone |
| CAVIN2 | caveolae associated protein 2 |
| ADIRF | adipogenesis regulatory factor |
| TWIST2 | twist family bHLH transcription factor 2 |
| RERGL | RERG like |
| MYOCD | myocardin |
| TMEM176A | transmembrane protein 176A |
| WWTR1 | WW domain containing transcription regulator 1 |
| TFPI2 | tissue factor pathway inhibitor 2 |
| GJC1 | gap junction protein gamma 1 |
| NKX3-2 | NK3 homeobox 2 |
| DHRS9 | dehydrogenase/reductase 9 |
| SOBP | sine oculis binding protein homolog |
| FERMT2 | FERM domain containing kindlin 2 |
| PCDHGB6 | protocadherin gamma subfamily B, 6 |
| PTH1R | parathyroid hormone 1 receptor |
| ANGPTL1 | angiopoietin like 1 |
| MAN1C1 | mannosidase alpha class 1C member 1 |
| CD8B2 | CD8b2 molecule |
| RP11-736K20.4 | NA |
| SNORD3B-1 | NA |
| SSPN | sarcospan |
| GEM | GTP binding protein overexpressed in skeletal muscle |
| GREM2 | gremlin 2, DAN family BMP antagonist |
| WTIP | WT1 interacting protein |
| U3 | NA |
| NBEAL2 | neurobeachin like 2 |
| AC053503.12 | NA |
| NRXN1 | neurexin 1 |
| TSHZ2 | teashirt zinc finger homeobox 2 |

| | |
|---|---|
| PNMA6A | PNMA family member 6A |
| IGFBP4 | insulin like growth factor binding protein 4 |
| SOX10 | SRY-box transcription factor 10 |
| CCNB1 | cyclin B1 |
| RSPO3 | R-spondin 3 |
| TRPC1 | transient receptor potential cation channel subfamily C member 1 |
| HSD17B6 | hydroxysteroid 17-beta dehydrogenase 6 |
| TGFBR3 | transforming growth factor beta receptor 3 |
| CXCL8 | C-X-C motif chemokine ligand 8 |
| C7 | complement C7 |
| PLEKHO1 | pleckstrin homology domain containing O1 |
| TPM1 | tropomyosin 1 |
| S100P | S100 calcium binding protein P |
| CAMSAP3 | calmodulin regulated spectrin associated protein family member 3 |
| RNF150 | ring finger protein 150 |
| ISLR | immunoglobulin superfamily containing leucine rich repeat |
| NFIC | nuclear factor I C |
| CLIP4 | CAP-Gly domain containing linker protein family member 4 |
| GHR | growth hormone receptor |
| GPX3 | glutathione peroxidase 3 |
| LYPD8 | LY6/PLAUR domain containing 8 |
| COLEC12 | collectin subfamily member 12 |
| COL7A1 | collagen type VII alpha 1 chain |
| FGFR1 | fibroblast growth factor receptor 1 |
| L1TD1 | LINE1 type transposase domain containing 1 |
| TMEM59L | transmembrane protein 59 like |
| LRIG1 | leucine rich repeats and immunoglobulin like domains 1 |
| MEOX2 | mesenchyme homeobox 2 |
| GFUS | GDP-L-fucose synthase |
| ZNF331 | zinc finger protein 331 |
| ESM1 | endothelial cell specific molecule 1 |
| SFRP2 | secreted frizzled related protein 2 |
| PYGM | glycogen phosphorylase, muscle associated |

| | |
|---|---|
| CCL5 | C-C motif chemokine ligand 5 |
| FAM180A | family with sequence similarity 180 member A |
| ZNF483 | zinc finger protein 483 |
| FAAH | fatty acid amide hydrolase |
| BHLHE22 | basic helix-loop-helix family member e22 |
| GFRA1 | GDNF family receptor alpha 1 |
| FBXL22 | F-box and leucine rich repeat protein 22 |
| PART1 | prostate androgen-regulated transcript 1 |
| TREM1 | triggering receptor expressed on myeloid cells 1 |
| RBMS3 | RNA binding motif single stranded interacting protein 3 |
| NBEA | neurobeachin |
| REG1B | regenerating family member 1 beta |
| PDZRN3 | PDZ domain containing ring finger 3 |
| P2RX1 | purinergic receptor P2X 1 |
| CAND2 | cullin associated and neddylation dissociated 2 (putative) |
| TEF | TEF transcription factor, PAR bZIP family member |
| TPSAB1 | tryptase alpha/beta 1 |
| NUDT10 | nudix hydrolase 10 |
| BEST4 | bestrophin 4 |
| HSPB8 | heat shock protein family B (small) member 8 |
| TFF1 | trefoil factor 1 |
| KCNH2 | potassium voltage-gated channel subfamily H member 2 |
| CRISPLD2 | cysteine rich secretory protein LCCL domain containing 2 |
| IGF1 | insulin like growth factor 1 |
| CXCL3 | C-X-C motif chemokine ligand 3 |
| DHRSX | NA |
| RGS1 | regulator of G protein signaling 1 |
| GLI3 | GLI family zinc finger 3 |
| ARL4D | ADP ribosylation factor like GTPase 4D |
| ATP2C2 | ATPase secretory pathway Ca2+ transporting 2 |
| RP4-565E6.1 | uncharacterized LOC101927468 |
| AQP1 | aquaporin 1 (Colton blood group) |
| BNC2 | basonuclin 2 |

| | |
|---|---|
| PALLD | palladin, cytoskeletal associated protein |
| CLEC3B | C-type lectin domain family 3 member B |
| PLAC8 | placenta associated 8 |
| UBE2C | ubiquitin conjugating enzyme E2 C |
| TUBB2B | tubulin beta 2B class IIb |
| LRP4 | LDL receptor related protein 4 |
| DIXDC1 | DIX domain containing 1 |
| NR3C1 | nuclear receptor subfamily 3 group C member 1 |
| RGS2 | regulator of G protein signaling 2 |
| CAP2 | cyclase associated actin cytoskeleton regulatory protein 2 |
| LHFPL6 | LHFPL tetraspan subfamily member 6 |
| PER3 | period circadian regulator 3 |
| TNFAIP8L3 | TNF alpha induced protein 8 like 3 |
| PLA2G2C | phospholipase A2 group IIC |
| FBLN2 | fibulin 2 |
| S100A11 | S100 calcium binding protein A11 |
| FER1L4 | fer-1 like family member 4 (pseudogene) |
| INMT | indolethylamine N-methyltransferase |
| SUSD2 | sushi domain containing 2 |
| ADAMTS1 | ADAM metallopeptidase with thrombospondin type 1 motif 1 |
| TSPAN2 | tetraspanin 2 |
| HIF3A | hypoxia inducible factor 3 subunit alpha |
| TGFB1I1 | transforming growth factor beta 1 induced transcript 1 |
| APLN | apelin |
| NFIX | nuclear factor I X |
| NEXMIF | neurite extension and migration factor |
| SPART | spartin |
| AC092168.2 | NA |
| PLAAT4 | phospholipase A and acyltransferase 4 |
| ZNF43 | zinc finger protein 43 |
| SSC5D | scavenger receptor cysteine rich family member with 5 domains |
| ZSCAN18 | zinc finger and SCAN domain containing 18 |
| ZNF239 | zinc finger protein 239 |

| | |
|---|---|
| COL6A1 | collagen type VI alpha 1 chain |
| HOXD10 | homeobox D10 |
| CES1 | carboxylesterase 1 |
| EML1 | EMAP like 1 |
| SLC23A1 | solute carrier family 23 member 1 |
| SERPING1 | serpin family G member 1 |
| PEG10 | paternally expressed 10 |
| FANCG | FA complementation group G |
| FBXO17 | F-box protein 17 |
| FAXC | failed axon connections homolog, metaxin like GST domain containing |
| PTCHD1 | patched domain containing 1 |
| NXN | nucleoredoxin |
| CIDEC | cell death inducing DFFA like effector c |
| BAG2 | BAG cochaperone 2 |
| ALDH1A3 | aldehyde dehydrogenase 1 family member A3 |
| PALM | paralemmin |
| NR1D2 | nuclear receptor subfamily 1 group D member 2 |
| LONRF1 | LON peptidase N-terminal domain and ring finger 1 |
| MRPS30-DT | NA |
| PLAAT3 | phospholipase A and acyltransferase 3 |
| DOCK6 | dedicator of cytokinesis 6 |
| KCNIP4 | potassium voltage-gated channel interacting protein 4 |
| PER1 | period circadian regulator 1 |
| AQP7 | aquaporin 7 |
| NR2F1-AS1 | NR2F1 antisense RNA 1 |
| MYOT | myotilin |
| FEZ1 | fasciculation and elongation protein zeta 1 |
| AMIGO2 | adhesion molecule with Ig like domain 2 |
| RP11-532F6.3 | NA |
| RP11-571M6.19 | NA |
| GNAL | G protein subunit alpha L |
| MUC2 | mucin 2, oligomeric mucus/gel-forming |
| CD34 | CD34 molecule |

| | |
|---|---|
| NSG2 | neuronal vesicle trafficking associated 2 |
| CHRM2 | cholinergic receptor muscarinic 2 |
| ADAMTS9-AS1 | ADAMTS9 antisense RNA 1 |
| FAXDC2 | fatty acid hydroxylase domain containing 2 |
| LINC00261 | long intergenic non-protein coding RNA 261 |
| SKA3 | spindle and kinetochore associated complex subunit 3 |
| ECM2 | extracellular matrix protein 2 |
| TUBB3 | tubulin beta 3 class III |
| NTRK3 | neurotrophic receptor tyrosine kinase 3 |
| CLIP3 | CAP-Gly domain containing linker protein 3 |
| TPPP | tubulin polymerization promoting protein |
| ABCA8 | ATP binding cassette subfamily A member 8 |
| NPR1 | natriuretic peptide receptor 1 |
| PLEKHA4 | pleckstrin homology domain containing A4 |
| RGN | regucalcin |
| ATP6V0C | ATPase H+ transporting V0 subunit c |
| HACD1 | 3-hydroxyacyl-CoA dehydratase 1 |
| BEST2 | bestrophin 2 |
| ESPL1 | extra spindle pole bodies like 1, separase |
| SLC29A2 | solute carrier family 29 member 2 |
| GYPC | glycophorin C (Gerbich blood group) |
| CACNA1C | calcium voltage-gated channel subunit alpha1 C |
| DPYD | dihydropyrimidine dehydrogenase |
| KRT17 | keratin 17 |
| PDLIM7 | PDZ and LIM domain 7 |
| CEACAM5 | CEA cell adhesion molecule 5 |
| MCM7 | minichromosome maintenance complex component 7 |
| LTB | lymphotoxin beta |
| HMGA1 | high mobility group AT-hook 1 |
| NR4A1 | nuclear receptor subfamily 4 group A member 1 |
| SELE | selectin E |
| NOS1 | nitric oxide synthase 1 |
| FAM110B | family with sequence similarity 110 member B |

| | |
|---|---|
| ZEB1 | zinc finger E-box binding homeobox 1 |
| TRIB3 | tribbles pseudokinase 3 |
| NEGR1 | neuronal growth regulator 1 |
| TRNP1 | TMF1 regulated nuclear protein 1 |
| PPA1 | inorganic pyrophosphatase 1 |
| GNG7 | G protein subunit gamma 7 |
| TUBB4B | tubulin beta 4B class IVb |
| DNER | delta/notch like EGF repeat containing |
| LEFTY1 | left-right determination factor 1 |
| AARD | alanine and arginine rich domain containing protein |
| PPP2R2B | protein phosphatase 2 regulatory subunit Bbeta |
| FMO2 | flavin containing dimethylaniline monoxygenase 2 |
| SERPINF1 | serpin family F member 1 |
| SLC7A5 | solute carrier family 7 member 5 |
| EGR1 | early growth response 1 |
| MANF | mesencephalic astrocyte derived neurotrophic factor |
| HLF | HLF transcription factor, PAR bZIP family member |
| GSTM2 | glutathione S-transferase mu 2 |
| IMPDH1 | inosine monophosphate dehydrogenase 1 |
| RADIL | Rap associating with DIL domain |
| IL1RN | interleukin 1 receptor antagonist |
| PDX1 | pancreatic and duodenal homeobox 1 |
| TMEM130 | transmembrane protein 130 |
| ZNF362 | zinc finger protein 362 |
| PMP22 | peripheral myelin protein 22 |
| MS4A2 | membrane spanning 4-domains A2 |
| TNFSF13B | TNF superfamily member 13b |
| GPNMB | glycoprotein nmb |
| MCRIP1 | MAPK regulated corepressor interacting protein 1 |
| CDK14 | cyclin dependent kinase 14 |
| NRARP | NOTCH regulated ankyrin repeat protein |
| JCAD | junctional cadherin 5 associated |
| APCDD1L | APC down-regulated 1 like |

| | |
|---|---|
| TACR2 | tachykinin receptor 2 |
| CRACR2B | calcium release activated channel regulator 2B |
| RHOB | ras homolog family member B |
| MMP3 | matrix metallopeptidase 3 |
| SLC52A2 | solute carrier family 52 member 2 |
| RTN1 | reticulon 1 |
| HUNK | hormonally up-regulated Neu-associated kinase |
| MASP1 | MBL associated serine protease 1 |
| PCSK9 | proprotein convertase subtilisin/kexin type 9 |
| NCS1 | neuronal calcium sensor 1 |
| CCDC9B | coiled-coil domain containing 9B |
| VAT1 | vesicle amine transport 1 |
| ZNF580 | zinc finger protein 580 |
| HSPB2 | heat shock protein family B (small) member 2 |
| ANKRD35 | ankyrin repeat domain 35 |
| BCHE | butyrylcholinesterase |
| FOXA2 | forkhead box A2 |
| LYVE1 | lymphatic vessel endothelial hyaluronan receptor 1 |
| AUNIP | aurora kinase A and ninein interacting protein |
| RP4-569M23.4 | NA |
| GPRASP1 | G protein-coupled receptor associated sorting protein 1 |
| NOP2 | NOP2 nucleolar protein |
| LYNX1 | Ly6/neurotoxin 1 |
| MUSK | muscle associated receptor tyrosine kinase |
| COPZ2 | COPI coat complex subunit zeta 2 |
| RFC3 | replication factor C subunit 3 |
| TNS2 | tensin 2 |
| TSC22D3 | TSC22 domain family member 3 |
| PLP1 | proteolipid protein 1 |
| PDE2A | phosphodiesterase 2A |
| CNBP | CCHC-type zinc finger nucleic acid binding protein |
| ZC3H6 | zinc finger CCCH-type containing 6 |
| L3MBTL2-AS1 | NA |

| | |
|---|---|
| PLIN1 | perilipin 1 |
| ZFPM2 | zinc finger protein, FOG family member 2 |
| RYR3 | ryanodine receptor 3 |
| NAALAD2 | N-acetylated alpha-linked acidic dipeptidase 2 |
| FFAR4 | free fatty acid receptor 4 |
| HAND2 | heart and neural crest derivatives expressed 2 |
| ZFP36L1 | ZFP36 ring finger protein like 1 |
| TESC | tescalcin |
| AKT3 | AKT serine/threonine kinase 3 |
| CD109 | CD109 molecule |
| OSR2 | odd-skipped related transciption factor 2 |
| SLC5A6 | solute carrier family 5 member 6 |
| ACAN | aggrecan |
| UNC13B | unc-13 homolog B |
| RP11-21N3.2 | NA |
| CXCL5 | C-X-C motif chemokine ligand 5 |
| ZNF329 | zinc finger protein 329 |
| CORO6 | coronin 6 |
| MAPK10 | mitogen-activated protein kinase 10 |
| ZNF547 | zinc finger protein 547 |
| HBA2 | hemoglobin subunit alpha 2 |
| SLC2A1 | solute carrier family 2 member 1 |
| CXCL6 | C-X-C motif chemokine ligand 6 |
| ENAH | ENAH actin regulator |
| GZMK | granzyme K |
| SERPINB8 | serpin family B member 8 |
| TMEM255A | transmembrane protein 255A |
| CLCA4 | chloride channel accessory 4 |
| ASRGL1 | asparaginase and isoaspartyl peptidase 1 |
| FOS | Fos proto-oncogene, AP-1 transcription factor subunit |
| NAALADL1 | N-acetylated alpha-linked acidic dipeptidase like 1 |
| CHRDL2 | chordin like 2 |
| BAIAP2L1 | BAR/IMD domain containing adaptor protein 2 like 1 |

| | |
|---|---|
| BST2 | bone marrow stromal cell antigen 2 |
| ANKRD33B | ankyrin repeat domain 33B |
| JUN | Jun proto-oncogene, AP-1 transcription factor subunit |
| SALL1 | spalt like transcription factor 1 |
| RAB5IF | RAB5 interacting factor |
| ZBTB47 | zinc finger and BTB domain containing 47 |
| BMP8A | bone morphogenetic protein 8a |
| FAM107A | family with sequence similarity 107 member A |
| LDOC1 | LDOC1 regulator of NFKB signaling |
| PPP1R12A | protein phosphatase 1 regulatory subunit 12A |
| ACTC1 | actin alpha cardiac muscle 1 |
| WBP1L | WW domain binding protein 1 like |
| METTL7A | methyltransferase like 7A |
| SIX4 | SIX homeobox 4 |
| RP11-236L14.2 | NA |
| ID4 | inhibitor of DNA binding 4, HLH protein |
| ZNF512 | zinc finger protein 512 |
| ARHGAP8 | Rho GTPase activating protein 8 |
| TUBA1A | tubulin alpha 1a |
| METTL24 | methyltransferase like 24 |
| PLK1 | polo like kinase 1 |
| RBMS1 | RNA binding motif single stranded interacting protein 1 |
| MAP1A | microtubule associated protein 1A |
| MAOB | monoamine oxidase B |
| AIF1L | allograft inflammatory factor 1 like |
| SYNGR1 | synaptogyrin 1 |
| DUSP26 | dual specificity phosphatase 26 |
| PKIG | cAMP-dependent protein kinase inhibitor gamma |
| KLK3 | kallikrein related peptidase 3 |
| CCN2 | cellular communication network factor 2 |
| C1R | complement C1r |
| HELZ2 | helicase with zinc finger 2 |
| SNORD3A | NA |

| | |
|---|---|
| AGTR1 | angiotensin II receptor type 1 |
| ADAMTS8 | ADAM metallopeptidase with thrombospondin type 1 motif 8 |
| TMEM158 | transmembrane protein 158 |
| PLCXD3 | phosphatidylinositol specific phospholipase C X domain containing 3 |
| CLDN4 | claudin 4 |
| CD36 | CD36 molecule |
| ITGA9 | integrin subunit alpha 9 |
| TENM1 | teneurin transmembrane protein 1 |
| IL11RA | interleukin 11 receptor subunit alpha |
| GUCY1A1 | guanylate cyclase 1 soluble subunit alpha 1 |
| NECTIN1 | nectin cell adhesion molecule 1 |
| LAMC3 | laminin subunit gamma 3 |
| ZNF675 | zinc finger protein 675 |
| PCP4 | Purkinje cell protein 4 |
| CYRIA | CYFIP related Rac1 interactor A |
| HRK | harakiri, BCL2 interacting protein |
| ANKRD12 | ankyrin repeat domain 12 |
| VGLL3 | vestigial like family member 3 |
| MAL | mal, T cell differentiation protein |
| DSTN | destrin, actin depolymerizing factor |
| RASGRP2 | RAS guanyl releasing protein 2 |
| HES6 | hes family bHLH transcription factor 6 |
| CEP85 | centrosomal protein 85 |
| GNB4 | G protein subunit beta 4 |
| HSF2 | heat shock transcription factor 2 |
| CX3CL1 | C-X3-C motif chemokine ligand 1 |
| SPOCK1 | SPARC (osteonectin), cwcv and kazal like domains proteoglycan 1 |
| CITED2 | Cbp/p300 interacting transactivator with Glu/Asp rich carboxy-terminal domain 2 |
| CAVIN1 | caveolae associated protein 1 |
| CTD-2126E3.6 | NA |
| ATP5F1D | ATP synthase F1 subunit delta |
| RP11-77K12.9 | NA |
| PBXIP1 | PBX homeobox interacting protein 1 |

| | |
|---|---|
| RP11-160E2.6 | NA |
| PBX1 | PBX homeobox 1 |
| NTN1 | netrin 1 |
| DYNC1I1 | dynein cytoplasmic 1 intermediate chain 1 |
| FAM156A | family with sequence similarity 156 member A |
| SCUBE2 | signal peptide, CUB domain and EGF like domain containing 2 |
| GASK1A | golgi associated kinase 1A |
| RNVU1-15 | RNA, variant U1 small nuclear 15 |
| HILPDA | hypoxia inducible lipid droplet associated |
| UBE2V1 | ubiquitin conjugating enzyme E2 V1 |
| F8A1 | coagulation factor VIII associated 1 |
| FAM83H | family with sequence similarity 83 member H |
| PDLIM4 | PDZ and LIM domain 4 |
| GPRC5A | G protein-coupled receptor class C group 5 member A |
| GPM6B | glycoprotein M6B |
| DUOX2 | dual oxidase 2 |
| ANXA1 | annexin A1 |
| BUD31 | BUD31 homolog |
| SPDEF | SAM pointed domain containing ETS transcription factor |
| RPP25 | ribonuclease P and MRP subunit p25 |
| SPC25 | SPC25 component of NDC80 kinetochore complex |
| LIMCH1 | LIM and calponin homology domains 1 |
| PCDH7 | protocadherin 7 |
| SEPTIN6 | septin 6 |
| ZNF296 | zinc finger protein 296 |
| TIMP2 | TIMP metallopeptidase inhibitor 2 |
| MEDAG | mesenteric estrogen dependent adipogenesis |
| SCGN | secretagogin, EF-hand calcium binding protein |
| SLIT2 | slit guidance ligand 2 |
| GSTM5 | glutathione S-transferase mu 5 |
| DDX39A | DExD-box helicase 39A |
| LINC01197 | lymphatic endothelial transcriptional regulator lncRNA 1 |
| MIR663AHG | MIR663A host gene |

| | |
|---|---|
| AC142472.6 | NA |
| RP5-1061H20.4 | uncharacterized LOC105373159 |
| CH17-360D5.2 | NA |
| ADAM11 | ADAM metallopeptidase domain 11 |
| NANOGP1 | NA |
| ITPR1 | inositol 1,4,5-trisphosphate receptor type 1 |
| RP11-554A11.11 | NA |
| BCL2L2-PABPN | BCL2L2-PABPN1 readthrough |
| LRP8 | LDL receptor related protein 8 |
| ZNF471 | zinc finger protein 471 |
| SETBP1 | SET binding protein 1 |
| HBB | hemoglobin subunit beta |
| TFF3 | trefoil factor 3 |
| SORCS2 | sortilin related VPS10 domain containing receptor 2 |
| RP11-631M6.2 | NA |
| CLDN3 | claudin 3 |
| MXRA7 | matrix remodeling associated 7 |
| SCNN1A | sodium channel epithelial 1 subunit alpha |
| RAB23 | RAB23, member RAS oncogene family |
| ARHGAP6 | Rho GTPase activating protein 6 |
| REG4 | regenerating family member 4 |
| PHLDA2 | pleckstrin homology like domain family A member 2 |
| CHI3L1 | chitinase 3 like 1 |
| ADAMTSL1 | ADAMTS like 1 |
| RP11-489E7.4 | NA |
| C4BPA | complement component 4 binding protein alpha |
| CPXM1 | carboxypeptidase X, M14 family member 1 |
| BACE2 | beta-secretase 2 |
| ROR1 | receptor tyrosine kinase like orphan receptor 1 |
| IGSF1 | immunoglobulin superfamily member 1 |
| HOXC4 | homeobox C4 |
| LINC00891 | NA |
| LINC02086 | NA |

| | |
|---|---|
| SLC2A3 | solute carrier family 2 member 3 |
| FAM180B | family with sequence similarity 180 member B |
| RP11-58O9.2 | NA |
| CXCL2 | C-X-C motif chemokine ligand 2 |
| NR4A3 | nuclear receptor subfamily 4 group A member 3 |
| KAT2B | lysine acetyltransferase 2B |
| DPM2 | dolichyl-phosphate mannosyltransferase subunit 2, regulatory |
| HTRA1 | HtrA serine peptidase 1 |
| AEBP1 | AE binding protein 1 |
| NR2F1 | nuclear receptor subfamily 2 group F member 1 |
| SKIDA1 | SKI/DACH domain containing 1 |
| RP11-760H22.2 | NA |
| SMPX | small muscle protein X-linked |
| PALMD | palmdelphin |
| CD27-AS1 | CD27 antisense RNA 1 |
| IQANK1 | IQ motif and ankyrin repeat containing 1 |
| PKD2 | polycystin 2, transient receptor potential cation channel |
| LINC02870 | NA |
| DIP2C | disco interacting protein 2 homolog C |
| C11orf96 | chromosome 11 open reading frame 96 |
| WDR90 | WD repeat domain 90 |
| ECI2 | enoyl-CoA delta isomerase 2 |
| INPP5A | inositol polyphosphate-5-phosphatase A |
| EPHA3 | EPH receptor A3 |
| STIP1 | stress induced phosphoprotein 1 |
| PRKAA2 | protein kinase AMP-activated catalytic subunit alpha 2 |
| EBF1 | EBF transcription factor 1 |
| TMOD1 | tropomodulin 1 |
| AP001347.6 | NA |
| MDGA1 | MAM domain containing glycosylphosphatidylinositol anchor 1 |
| HHAT | hedgehog acyltransferase |
| MT-TN | NA |
| CPQ | carboxypeptidase Q |

| | |
|---|---|
| ZNF287 | zinc finger protein 287 |
| MRGPRF | MAS related GPR family member F |
| WFDC2 | WAP four-disulfide core domain 2 |
| SPIRE2 | spire type actin nucleation factor 2 |
| ETV4 | ETS variant transcription factor 4 |
| DEPDC1B | DEP domain containing 1B |
| PRDM1 | PR/SET domain 1 |
| E2F1 | E2F transcription factor 1 |
| DPP4 | dipeptidyl peptidase 4 |
| ECRG4 | ECRG4 augurin precursor |
| MAST4 | microtubule associated serine/threonine kinase family member 4 |
| MN1 | MN1 proto-oncogene, transcriptional regulator |
| MEF2C | myocyte enhancer factor 2C |
| ARRDC3 | arrestin domain containing 3 |
| CCT3 | chaperonin containing TCP1 subunit 3 |
| KCNN3 | potassium calcium-activated channel subfamily N member 3 |
| SOX2 | SRY-box transcription factor 2 |
| ASPN | asporin |
| H2AC19 | H2A clustered histone 19 |
| NAT8L | N-acetyltransferase 8 like |
| RP11-208N14.5 | uncharacterized LOC101929130 |
| KPNA2 | karyopherin subunit alpha 2 |
| RP11-686O6.2 | NA |
| LURAP1 | leucine rich adaptor protein 1 |
| RIPK3 | receptor interacting serine/threonine kinase 3 |
| NELFCD | negative elongation factor complex member C/D |
| CIRBP | cold inducible RNA binding protein |
| ASXL3 | ASXL transcriptional regulator 3 |
| NGFR | nerve growth factor receptor |
| FAM229B | family with sequence similarity 229 member B |
| FAM66C | family with sequence similarity 66 member C |
| NLRP1 | NLR family pyrin domain containing 1 |
| SERTAD4 | SERTA domain containing 4 |

| | |
|---|---|
| IQGAP3 | IQ motif containing GTPase activating protein 3 |
| SMARCA1 | SWI/SNF related, matrix associated, actin dependent regulator of chromatin, subfamily a, member 1 |
| CNKSR1 | connector enhancer of kinase suppressor of Ras 1 |
| SHISA3 | shisa family member 3 |
| SMG1P5 | SMG1 pseudogene 5 |
| IFIT1 | interferon induced protein with tetratricopeptide repeats 1 |
| GAS1 | growth arrest specific 1 |
| IL17RD | interleukin 17 receptor D |
| ITGA2 | integrin subunit alpha 2 |
| PNMA1 | PNMA family member 1 |
| AKAP17A | A-kinase anchoring protein 17A |
| LPL | lipoprotein lipase |
| ITGA6 | integrin subunit alpha 6 |
| FOXD4L3 | forkhead box D4 like 3 |
| COL8A2 | collagen type VIII alpha 2 chain |
| CLMP | CXADR like membrane protein |
| GGTA1 | glycoprotein alpha-galactosyltransferase 1 (inactive) |
| COX7A1 | cytochrome c oxidase subunit 7A1 |
| UBE2E2 | ubiquitin conjugating enzyme E2 E2 |
| P2RX7 | purinergic receptor P2X 7 |
| EPHA7 | EPH receptor A7 |
| CBX7 | chromobox 7 |
| VAMP7 | NA |
| ADGRD1 | adhesion G protein-coupled receptor D1 |
| TEDC2 | tubulin epsilon and delta complex 2 |
| CA14 | carbonic anhydrase 14 |
| CSPG4 | chondroitin sulfate proteoglycan 4 |
| PARVA | parvin alpha |
| GLIS1 | GLIS family zinc finger 1 |
| CDT1 | chromatin licensing and DNA replication factor 1 |
| TNXB | tenascin XB |
| DDX54 | DEAD-box helicase 54 |
| RPIA | ribose 5-phosphate isomerase A |

| | |
|---|---|
| CSPG4P12 | chondroitin sulfate proteoglycan 4 pseudogene |
| COPG1 | COPI coat complex subunit gamma 1 |
| ZNF888 | zinc finger protein 888 |
| HOXD9 | homeobox D9 |
| LSP1 | lymphocyte specific protein 1 |
| HOPX | HOP homeobox |
| FOXF1 | forkhead box F1 |
| GSN | gelsolin |
| PRKCB | protein kinase C beta |
| CTD-2036P10.6 | NA |
| MCM2 | minichromosome maintenance complex component 2 |
| CCN1 | cellular communication network factor 1 |
| JAM2 | junctional adhesion molecule 2 |
| NEURL1B | neuralized E3 ubiquitin protein ligase 1B |
| DPP6 | dipeptidyl peptidase like 6 |
| RBP7 | retinol binding protein 7 |
| HYAL2 | hyaluronidase 2 |
| HLA-DOA | major histocompatibility complex, class II, DO alpha |
| ATF3 | activating transcription factor 3 |
| DCLK1 | doublecortin like kinase 1 |
| RP11-1348G14.5 | NA |
| SHCBP1 | SHC binding and spindle associated 1 |
| CCL2 | C-C motif chemokine ligand 2 |
| THRB-IT1 | NA |
| FANCA | FA complementation group A |
| REEP2 | receptor accessory protein 2 |
| CDO1 | cysteine dioxygenase type 1 |
| MUC4 | mucin 4, cell surface associated |
| CPB1 | carboxypeptidase B1 |
| ST8SIA1 | ST8 alpha-N-acetyl-neuraminide alpha-2,8-sialyltransferase 1 |
| SLC6A16 | solute carrier family 6 member 16 |
| PKM | pyruvate kinase M1/2 |
| RASA3 | RAS p21 protein activator 3 |

| | |
|---|---|
| WDR4 | WD repeat domain 4 |
| ZNF793 | zinc finger protein 793 |
| CACNA1H | calcium voltage-gated channel subunit alpha1 H |
| TNFRSF11B | TNF receptor superfamily member 11b |
| ROR2 | receptor tyrosine kinase like orphan receptor 2 |
| THBS2 | thrombospondin 2 |
| CD300LG | CD300 molecule like family member g |
| PUS1 | pseudouridine synthase 1 |
| NALT1 | NA |
| SEMA3G | semaphorin 3G |
| PPP1R1B | protein phosphatase 1 regulatory inhibitor subunit 1B |
| DEPP1 | DEPP1 autophagy regulator |
| ICA1L | islet cell autoantigen 1 like |
| LIG1 | DNA ligase 1 |
| ANKS1B | ankyrin repeat and sterile alpha motif domain containing 1B |
| UPF3AP2 | NA |
| PDE1A | phosphodiesterase 1A |
| PDRG1 | p53 and DNA damage regulated 1 |
| CCDC136 | coiled-coil domain containing 136 |
| NPTX1 | neuronal pentraxin 1 |
| PRPH | peripherin |
| C1QTNF3 | C1q and TNF related 3 |
| CRABP1 | cellular retinoic acid binding protein 1 |
| SNORD71 | small nucleolar RNA, C/D box 71 |
| SMAGP | small cell adhesion glycoprotein |
| SNX22 | sorting nexin 22 |
| FBXL18 | F-box and leucine rich repeat protein 18 |
| AURKB | aurora kinase B |
| HCST | hematopoietic cell signal transducer |
| PNISR | PNN interacting serine and arginine rich protein |
| STC1 | stanniocalcin 1 |
| EBF4 | EBF family member 4 |
| IL1RL2 | interleukin 1 receptor like 2 |

| | |
|---|---|
| WSCD2 | WSC domain containing 2 |
| TOMM5 | translocase of outer mitochondrial membrane 5 |
| ATP1B2 | ATPase Na+/K+ transporting subunit beta 2 |
| FBLN5 | fibulin 5 |
| PGR | progesterone receptor |
| MORN5 | MORN repeat containing 5 |
| OSR1 | odd-skipped related transcription factor 1 |
| F2RL3 | F2R like thrombin or trypsin receptor 3 |
| JPT1 | Jupiter microtubule associated homolog 1 |
| SLA2 | Src like adaptor 2 |
| CCN3 | cellular communication network factor 3 |
| TCEAL4 | transcription elongation factor A like 4 |
| TRIM11 | tripartite motif containing 11 |
| PKD1 | polycystin 1, transient receptor potential channel interacting |
| SPON1 | spondin 1 |
| CDK1 | cyclin dependent kinase 1 |
| LONP1 | lon peptidase 1, mitochondrial |
| TRAF2 | TNF receptor associated factor 2 |
| NDUFAF4 | NADH:ubiquinone oxidoreductase complex assembly factor 4 |
| ADAMTS4 | ADAM metallopeptidase with thrombospondin type 1 motif 4 |
| PYHIN1 | pyrin and HIN domain family member 1 |
| TRAC | T cell receptor alpha constant |
| PRUNE2 | prune homolog 2 with BCH domain |
| SAPCD2 | suppressor APC domain containing 2 |
| HDHD3 | haloacid dehalogenase like hydrolase domain containing 3 |
| SYNE1 | spectrin repeat containing nuclear envelope protein 1 |
| NRXN2 | neurexin 2 |
| BHLHE41 | basic helix-loop-helix family member e41 |
| UTP14A | UTP14A small subunit processome component |
| CDC42EP3 | CDC42 effector protein 3 |
| RRAGC | Ras related GTP binding C |
| AIF1 | allograft inflammatory factor 1 |
| CC2D2A | coiled-coil and C2 domain containing 2A |

| | |
|---|---|
| CYBRD1 | cytochrome b reductase 1 |
| ADAMTS16 | ADAM metallopeptidase with thrombospondin type 1 motif 16 |
| SEMA3E | semaphorin 3E |
| TRBC2 | T cell receptor beta constant 2 |
| TIMP4 | TIMP metallopeptidase inhibitor 4 |
| SPINK4 | serine peptidase inhibitor Kazal type 4 |
| GLB1L2 | galactosidase beta 1 like 2 |
| WFS1 | wolframin ER transmembrane glycoprotein |
| C1orf54 | chromosome 1 open reading frame 54 |
| RNF121 | ring finger protein 121 |
| FKBP4 | FKBP prolyl isomerase 4 |
| CBR3 | carbonyl reductase 3 |
| RP1-78O14.1 | NA |
| COLGALT2 | collagen beta(1-O)galactosyltransferase 2 |
| YTHDF1 | YTH N6-methyladenosine RNA binding protein 1 |
| FGFRL1 | fibroblast growth factor receptor like 1 |
| NOS2 | nitric oxide synthase 2 |
| PRRG3 | proline rich and Gla domain 3 |
| AC007191.4 | NA |
| STAP1 | signal transducing adaptor family member 1 |
| LSAMP | limbic system associated membrane protein |
| PRND | prion like protein doppel |
| FOXQ1 | forkhead box Q1 |
| MCM4 | minichromosome maintenance complex component 4 |
| KATNAL1 | katanin catalytic subunit A1 like 1 |
| CNTN1 | contactin 1 |
| CCND2-AS1 | CCND2 antisense RNA 1 |
| TXNIP | thioredoxin interacting protein |
| HOXD13 | homeobox D13 |
| FGF10 | fibroblast growth factor 10 |
| IGSF9B | immunoglobulin superfamily member 9B |
| DENND5B | DENN domain containing 5B |
| PLAU | plasminogen activator, urokinase |

| | |
|---|---|
| CTSE | cathepsin E |
| BRSK1 | BR serine/threonine kinase 1 |
| RP5-934G17.7 | NA |
| CDC20 | cell division cycle 20 |
| POU6F1 | POU class 6 homeobox 1 |
| SNAP25 | synaptosome associated protein 25 |
| SLC44A3-AS1 | SLC44A3 antisense RNA 1 |
| HERC2P3 | HERC2 pseudogene 3 |
| SBF2-AS1 | SBF2 antisense RNA 1 |
| ZMAT1 | zinc finger matrin-type 1 |
| VIM | vimentin |
| OSBPL6 | oxysterol binding protein like 6 |
| PABPC1 | poly(A) binding protein cytoplasmic 1 |
| RP11-1024P17.1 | NA |
| UHRF1 | ubiquitin like with PHD and ring finger domains 1 |
| DRD4 | dopamine receptor D4 |
| HLA-DOB | major histocompatibility complex, class II, DO beta |
| RPP38 | ribonuclease P/MRP subunit p38 |
| PTGDS | prostaglandin D2 synthase |
| EPS15-AS1 | NA |
| ABCA6 | ATP binding cassette subfamily A member 6 |
| MUC12 | mucin 12, cell surface associated |
| TTYH3 | tweety family member 3 |
| OLFML3 | olfactomedin like 3 |
| CCNE1 | cyclin E1 |
| GAS6 | growth arrest specific 6 |
| CP | ceruloplasmin |
| RTEL1-TNFRSF | RTEL1-TNFRSF6B readthrough (NMD candidate) |
| LINC01123 | long intergenic non-protein coding RNA 1123 |
| RP4-635A23.6 | NA |
| NAV2-AS6 | NA |
| SNHG14 | NA |
| RP4-583P15.15 | NA |

| | |
|---|---|
| TIMELESS | timeless circadian regulator |
| MFSD2B | major facilitator superfamily domain containing 2B |
| LRRC26 | leucine rich repeat containing 26 |
| SLC25A4 | solute carrier family 25 member 4 |
| SLC24A3 | solute carrier family 24 member 3 |
| GREM1 | gremlin 1, DAN family BMP antagonist |
| KRT6B | keratin 6B |
| LRFN5 | leucine rich repeat and fibronectin type III domain containing 5 |
| MTURN | maturin, neural progenitor differentiation regulator homolog |
| PLXNA4 | plexin A4 |
| GTF2IRD2B | GTF2I repeat domain containing 2B |
| CNTNAP3 | contactin associated protein family member 3 |
| RP5-864K19.6 | NA |
| RP11-973D8.5 | NA |
| RP11-373D17.4 | NA |
| SLC35B2 | solute carrier family 35 member B2 |
| REG1A | regenerating family member 1 alpha |
| ZNF346-IT1 | NA |
| VIT | vitrin |
| ANKRD29 | ankyrin repeat domain 29 |
| SAMD4A | sterile alpha motif domain containing 4A |
| SHISA4 | shisa family member 4 |
| YBX2 | Y-box binding protein 2 |
| HJURP | Holliday junction recognition protein |
| CH507-513H4.1 | NA |
| RP11-92C4.6 | NA |
| TSHZ3 | teashirt zinc finger homeobox 3 |
| C16orf89 | chromosome 16 open reading frame 89 |
| CMYA5 | cardiomyopathy associated 5 |
| ADCYAP1R1 | ADCYAP receptor type I |
| STC2 | stanniocalcin 2 |
| NDRG4 | NDRG family member 4 |
| HSPA1A | heat shock protein family A (Hsp70) member 1A |

| | |
|---|---|
| FGF7 | fibroblast growth factor 7 |
| LMO2 | LIM domain only 2 |
| CTD-2576D5.4 | uncharacterized LOC102723692 |
| MYEOV | myeloma overexpressed |
| CDC45 | cell division cycle 45 |
| NDRG2 | NDRG family member 2 |
| RP11-456N14.6 | NA |
| RP11-166O4.6 | NA |
| EFNA3 | ephrin A3 |
| DYNC2H1 | dynein cytoplasmic 2 heavy chain 1 |
| KLK11 | kallikrein related peptidase 11 |
| TRIM21 | tripartite motif containing 21 |
| LINC00702 | NA |
| DSCAML1 | DS cell adhesion molecule like 1 |
| ZNF598 | zinc finger protein 598, E3 ubiquitin ligase |
| PDZK1IP1 | PDZK1 interacting protein 1 |
| RP11-6O2.3 | NA |
| MIR17HG | miR-17-92a-1 cluster host gene |
| GPR143 | G protein-coupled receptor 143 |
| GLIS2 | GLIS family zinc finger 2 |
| SDR16C5 | short chain dehydrogenase/reductase family 16C member 5 |
| ZFP36 | ZFP36 ring finger protein |
| EMSLR | NA |
| RP11-2N1.3 | NA |
| HBA1 | hemoglobin subunit alpha 1 |
| ITGBL1 | integrin subunit beta like 1 |
| MRPS17 | mitochondrial ribosomal protein S17 |
| NFATC1 | nuclear factor of activated T cells 1 |
| CADM2 | cell adhesion molecule 2 |
| LRRC6 | dynein axonemal assembly factor 11 |
| DTYMK | deoxythymidylate kinase |
| CMKLR1 | chemerin chemokine-like receptor 1 |
| PIEZO2 | piezo type mechanosensitive ion channel component 2 |

| | |
|---|---|
| CAB39L | calcium binding protein 39 like |
| ADRA2A | adrenoceptor alpha 2A |
| ZNF385C | zinc finger protein 385C |
| VGF | VGF nerve growth factor inducible |
| PLAUR | plasminogen activator, urokinase receptor |
| TPX2 | TPX2 microtubule nucleation factor |
| ERF | ETS2 repressor factor |
| ZSCAN31 | zinc finger and SCAN domain containing 31 |
| FIBIN | fin bud initiation factor homolog |
| KIF21B | kinesin family member 21B |
| GPR155 | G protein-coupled receptor 155 |
| CNTN2 | contactin 2 |
| CDH23 | cadherin related 23 |
| DNAJB2 | DnaJ heat shock protein family (Hsp40) member B2 |
| LINC01579 | uncharacterized LOC105369203 |
| RP11-33E12.2 | NA |
| CACNA1A | calcium voltage-gated channel subunit alpha1 A |
| DMPK | DM1 protein kinase |
| ADAMTS15 | ADAM metallopeptidase with thrombospondin type 1 motif 15 |
| MLPH | melanophilin |
| TRIM29 | tripartite motif containing 29 |
| HSD17B12 | hydroxysteroid 17-beta dehydrogenase 12 |
| RP11-229D13.3 | NA |
| TSPAN18 | tetraspanin 18 |
| SNHG15 | small nucleolar RNA host gene 15 |
| PCDH20 | protocadherin 20 |
| PABPC1L | poly(A) binding protein cytoplasmic 1 like |
| ARHGEF26 | Rho guanine nucleotide exchange factor 26 |
| CTD-2270P14.5 | NA |
| RP11-342K6.1 | NA |
| FYCO1 | FYVE and coiled-coil domain autophagy adaptor 1 |
| MSX2 | msh homeobox 2 |
| AIMP2 | aminoacyl tRNA synthetase complex interacting multifunctional protein 2 |

| | |
|---|---|
| RAP1GAP | RAP1 GTPase activating protein |
| CTF1 | cardiotrophin 1 |
| KLF2 | Kruppel like factor 2 |
| TM4SF4 | transmembrane 4 L six family member 4 |
| PRCC | proline rich mitotic checkpoint control factor |
| CCL3L1 | C-C motif chemokine ligand 3 like 1 |
| NKG7 | natural killer cell granule protein 7 |
| AC100830.3 | NA |
| RPP40 | ribonuclease P/MRP subunit p40 |
| CCL21 | C-C motif chemokine ligand 21 |
| LINC02595 | NA |
| CYP4F35P | cytochrome P450 family 4 subfamily F member 35, pseudogene |
| SNHG17 | small nucleolar RNA host gene 17 |
| CTA-215D11.5 | NA |
| RTL5 | retrotransposon Gag like 5 |
| SNCA | synuclein alpha |
| NRK | Nik related kinase |
| MT-TY | NA |
| POLR2J | RNA polymerase II subunit J |
| CSDC2 | cold shock domain containing C2 |
| TK1 | thymidine kinase 1 |
| LEP | leptin |
| KLF9 | Kruppel like factor 9 |
| PLPP4 | phospholipid phosphatase 4 |
| CTB-131B5.5 | uncharacterized LOC101929719 |
| DOK5 | docking protein 5 |
| LAD1 | ladinin 1 |
| IL6R | interleukin 6 receptor |
| PITX2 | paired like homeodomain 2 |
| PRKAR2B | protein kinase cAMP-dependent type II regulatory subunit beta |
| SYT7 | synaptotagmin 7 |
| ABCC9 | ATP binding cassette subfamily C member 9 |
| RNU4-1 | RNA, U4 small nuclear 1 |

| | |
|---|---|
| AC098614.2 | NA |
| MYD88 | MYD88 innate immune signal transduction adaptor |
| TMC5 | transmembrane channel like 5 |
| DDIT4L | DNA damage inducible transcript 4 like |
| TRPM2-AS | NA |
| NFATC4 | nuclear factor of activated T cells 4 |
| MELK | maternal embryonic leucine zipper kinase |
| ITM2A | integral membrane protein 2A |
| FAM228B | family with sequence similarity 228 member B |
| CCL19 | C-C motif chemokine ligand 19 |
| RP11-342K2.1 | NA |
| GDPD3 | glycerophosphodiester phosphodiesterase domain containing 3 |
| SORBS2 | sorbin and SH3 domain containing 2 |
| TCEANC | transcription elongation factor A N-terminal and central domain containing |
| GS1-393G12.13 | NA |
| PRKG1 | protein kinase cGMP-dependent 1 |
| RP11-7F17.8 | NA |
| DCAF13 | DDB1 and CUL4 associated factor 13 |
| PXDC1 | PX domain containing 1 |
| TENM3 | teneurin transmembrane protein 3 |
| NUMBL | NUMB like endocytic adaptor protein |
| FAM133CP | NA |
| RDX | radixin |
| ZNF731P | NA |
| RP11-1114A5.4 | NA |
| RP11-157L3.13 | NA |
| TMEM132A | transmembrane protein 132A |
| LINC01357 | NA |
| JAZF1 | JAZF zinc finger 1 |
| CD52 | CD52 molecule |
| CAPN8 | calpain 8 |
| SLC30A4 | solute carrier family 30 member 4 |
| AURKA | aurora kinase A |

| | |
|---|---|
| ECEL1 | endothelin converting enzyme like 1 |
| VSTM4 | V-set and transmembrane domain containing 4 |
| LOXL2 | lysyl oxidase like 2 |
| ZNF503-AS1 | ZNF503 antisense RNA 1 |
| NUF2 | NUF2 component of NDC80 kinetochore complex |
| PCCA | propionyl-CoA carboxylase subunit alpha |
| RP11-295P9.13 | NA |
| TNS4 | tensin 4 |
| AC138430.4 | NA |
| WDR97 | WD repeat domain 97 |
| CCNE2 | cyclin E2 |
| ZNF829 | zinc finger protein 829 |
| MMP1 | matrix metallopeptidase 1 |
| AC005306.3 | NA |
| ZNF17 | zinc finger protein 17 |
| MMP2 | matrix metallopeptidase 2 |
| PLEKHH1 | pleckstrin homology, MyTH4 and FERM domain containing H1 |
| LIPE | lipase E, hormone sensitive type |
| USP30 | ubiquitin specific peptidase 30 |
| LGI4 | leucine rich repeat LGI family member 4 |
| RNU2-6P | RNA, U2 small nuclear 6, pseudogene |
| LSM11 | LSM11, U7 small nuclear RNA associated |
| GLI2 | GLI family zinc finger 2 |
| MRAS | muscle RAS oncogene homolog |
| FBLN7 | fibulin 7 |
| NPAS3 | neuronal PAS domain protein 3 |
| GCNA | germ cell nuclear acidic peptidase |
| TPTEP1 | TPTE pseudogene 1 |
| ALDH1A1 | aldehyde dehydrogenase 1 family member A1 |
| JRK | Jrk helix-turn-helix protein |
| SLC7A11 | solute carrier family 7 member 11 |
| RP11-486M3.4 | uncharacterized LOC101928868 |
| ZNF257 | zinc finger protein 257 |

| | |
|---|---|
| SLC25A5-AS1 | SLC25A5 antisense RNA 1 |
| HECW2-AS1 | HECW2 antisense RNA 1 |
| UBE2T | ubiquitin conjugating enzyme E2 T |
| CKS2 | CDC28 protein kinase regulatory subunit 2 |
| MDFIC | MyoD family inhibitor domain containing |
| AGR2 | anterior gradient 2, protein disulphide isomerase family member |
| PCLAF | PCNA clamp associated factor |
| PLCXD1 | phosphatidylinositol specific phospholipase C X domain containing 1 |
| BCYRN1 | brain cytoplasmic RNA 1 |
| TVP23C | trans-golgi network vesicle protein 23 homolog C |
| LA16c-431H6.7 | NA |
| RANBP17 | RAN binding protein 17 |
| AKAP6 | A-kinase anchoring protein 6 |
| TFF2 | trefoil factor 2 |
| SOX17 | SRY-box transcription factor 17 |
| TENT5B | terminal nucleotidyltransferase 5B |
| LOXL4 | lysyl oxidase like 4 |
| ZSCAN16-AS1 | ZSCAN16 antisense RNA 1 |
| PDZD4 | PDZ domain containing 4 |
| ANTXR2 | ANTXR cell adhesion molecule 2 |
| SPRTN | SprT-like N-terminal domain |
| GRAP2 | GRB2 related adaptor protein 2 |
| NLE1 | notchless homolog 1 |
| GPR132 | G protein-coupled receptor 132 |
| SAFB | scaffold attachment factor B |
| DHCR7 | 7-dehydrocholesterol reductase |
| PIK3CD | phosphatidylinositol-4,5-bisphosphate 3-kinase catalytic subunit delta |
| PRMT9 | protein arginine methyltransferase 9 |
| WNT9A | Wnt family member 9A |
| CMTM5 | CKLF like MARVEL transmembrane domain containing 5 |
| AC021218.2 | uncharacterized LOC389602 |
| LINC01322 | NA |
| DENND2C | DENN domain containing 2C |

| | |
|---|---|
| DYRK3 | dual specificity tyrosine phosphorylation regulated kinase 3 |
| RHCG | Rh family C glycoprotein |
| MAP6 | microtubule associated protein 6 |
| MMP23B | matrix metallopeptidase 23B |
| CIP2A | cellular inhibitor of PP2A |
| HMCN2 | hemicentin 2 |
| KRT80 | keratin 80 |
| FBXO30-DT | EPM2A divergent transcript |
| CX3CR1 | C-X3-C motif chemokine receptor 1 |
| RP11-167J8.5 | NA |
| PHF1 | PHD finger protein 1 |
| COL6A3 | collagen type VI alpha 3 chain |
| TPSG1 | tryptase gamma 1 |
| DUSP5 | dual specificity phosphatase 5 |
| RAC3 | Rac family small GTPase 3 |
| EXOC6B | exocyst complex component 6B |
| NCALD | neurocalcin delta |
| NFKBIZ | NFKB inhibitor zeta |
| PLA2G4C | phospholipase A2 group IVC |
| CADM1 | cell adhesion molecule 1 |
| CTD-3126B10.5 | NA |
| PI3 | peptidase inhibitor 3 |
| S100B | S100 calcium binding protein B |
| WFDC1 | WAP four-disulfide core domain 1 |
| MT-TE | NA |
| ZNF780A | zinc finger protein 780A |
| ORC6 | origin recognition complex subunit 6 |
| PDE3A | phosphodiesterase 3A |
| DPP3 | dipeptidyl peptidase 3 |
| ZDHHC15 | zinc finger DHHC-type palmitoyltransferase 15 |
| SALL2 | spalt like transcription factor 2 |
| ANGPT2 | angiopoietin 2 |
| RMRP | RNA component of mitochondrial RNA processing endoribonuclease |

| | |
|---|---|
| URB2 | URB2 ribosome biogenesis homolog |
| ZRANB1 | zinc finger RANBP2-type containing 1 |
| ANXA9 | annexin A9 |
| QSOX2 | quiescin sulfhydryl oxidase 2 |
| VSIG1 | V-set and immunoglobulin domain containing 1 |
| DCLK2 | doublecortin like kinase 2 |
| NRXN3 | neurexin 3 |
| CH17-353B19.2 | NA |
| AGAP4 | ArfGAP with GTPase domain, ankyrin repeat and PH domain 4 |
| MAP9 | microtubule associated protein 9 |
| CENPJ | centromere protein J |
| PNMA3 | PNMA family member 3 |
| SUGCT | succinyl-CoA:glutarate-CoA transferase |
| RP11-297P16.4 | NA |
| ABHD8 | abhydrolase domain containing 8 |
| DBN1 | drebrin 1 |
| CDIP1 | cell death inducing p53 target 1 |
| TBC1D2 | TBC1 domain family member 2 |
| MAGI2 | membrane associated guanylate kinase, WW and PDZ domain containing 2 |
| PEG3 | paternally expressed 3 |
| C3orf52 | chromosome 3 open reading frame 52 |
| ETNK2 | ethanolamine kinase 2 |
| DCAF4L1 | DDB1 and CUL4 associated factor 4 like 1 |
| PLCXD1 | NA |
| KATNB1 | katanin regulatory subunit B1 |
| PLCL1 | phospholipase C like 1 (inactive) |
| GAL3ST4 | galactose-3-O-sulfotransferase 4 |
| RP3-406A7.7 | NA |
| ABLIM3 | actin binding LIM protein family member 3 |
| MT-TM | NA |
| MS4A1 | membrane spanning 4-domains A1 |
| ADRB1 | adrenoceptor beta 1 |
| MYL3 | myosin light chain 3 |

| | |
|---|---|
| PUSL1 | pseudouridine synthase like 1 |
| LRRN4CL | LRRN4 C-terminal like |
| RP11-681B3.4 | NA |
| XRCC2 | X-ray repair cross complementing 2 |
| SLC9A3 | solute carrier family 9 member A3 |
| CDRT1 | CMT1A duplicated region transcript 1 |
| PITPNC1 | phosphatidylinositol transfer protein cytoplasmic 1 |
| ANLN | anillin actin binding protein |
| AC006042.8 | NA |
| ALDH1A2 | aldehyde dehydrogenase 1 family member A2 |
| RNF43 | ring finger protein 43 |
| TRBC1 | T cell receptor beta constant 1 |
| ZFP28 | ZFP28 zinc finger protein |
| CTD-3128G10.7 | NA |
| ANKRD18A | ankyrin repeat domain 18A |
| PUS7 | pseudouridine synthase 7 |
| PROS1 | protein S |
| CTA-390C10.10 | NA |
| SLC38A5 | solute carrier family 38 member 5 |
| HEATR1 | HEAT repeat containing 1 |
| EFEMP2 | EGF containing fibulin extracellular matrix protein 2 |
| SGCE | sarcoglycan epsilon |
| PRDM6 | PR/SET domain 6 |
| SLC8A2 | solute carrier family 8 member A2 |
| COL6A2 | collagen type VI alpha 2 chain |
| RAB19 | RAB19, member RAS oncogene family |
| ABL1 | ABL proto-oncogene 1, non-receptor tyrosine kinase |
| RP11-999E24.3 | NA |
| RP11-102G14.1 | NA |
| DNTTIP1 | deoxynucleotidyltransferase terminal interacting protein 1 |
| PDE1B | phosphodiesterase 1B |
| FOXO6 | forkhead box O6 |
| MYH3 | myosin heavy chain 3 |

| | |
|---|---|
| SCN2B | sodium voltage-gated channel beta subunit 2 |
| SYPL2 | synaptophysin like 2 |
| CPNE6 | copine 6 |
| STK33 | serine/threonine kinase 33 |
| MOXD1 | monooxygenase DBH like 1 |
| RP11-259O2.1 | uncharacterized LOC105374618 |
| ZDHHC8 | zinc finger DHHC-type palmitoyltransferase 8 |
| RP11-304L19.1 | uncharacterized LOC105371049 |
| ASCL2 | achaete-scute family bHLH transcription factor 2 |
| RP11-517I3.2 | NA |
| MLLT11 | MLLT11 transcription factor 7 cofactor |
| NUP58 | nucleoporin 58 |
| PLSCR4 | phospholipid scramblase 4 |
| ATP6V0E2-AS1 | ATP6V0E2 antisense RNA 1 |
| XXbac-BPGBPC | NA |
| IGFALS | insulin like growth factor binding protein acid labile subunit |
| ZNF341 | zinc finger protein 341 |
| AURKC | aurora kinase C |
| GPER1 | G protein-coupled estrogen receptor 1 |
| KLK12 | kallikrein related peptidase 12 |
| MOGAT3 | monoacylglycerol O-acyltransferase 3 |
| MTCL1 | microtubule crosslinking factor 1 |
| PKD1P6 | polycystin 1, transient receptor potential channel interacting pseudogene 6 |
| AP1S2 | adaptor related protein complex 1 subunit sigma 2 |
| CASP5 | caspase 5 |
| FAM72D | family with sequence similarity 72 member D |
| ACLY | ATP citrate lyase |
| RP3-399J4.2 | NA |
| STEAP4 | STEAP4 metalloreductase |
| DUSP19 | dual specificity phosphatase 19 |
| RALGAPA2 | Ral GTPase activating protein catalytic subunit alpha 2 |
| COL24A1 | collagen type XXIV alpha 1 chain |
| CRTC3 | CREB regulated transcription coactivator 3 |

| | |
|---|---|
| TRMT2A | tRNA methyltransferase 2 homolog A |
| GULP1 | GULP PTB domain containing engulfment adaptor 1 |
| MAL2 | mal, T cell differentiation protein 2 |
| TIAM1 | TIAM Rac1 associated GEF 1 |
| EVL | Enah/Vasp-like |
| AC009237.8 | NA |
| HABP4 | hyaluronan binding protein 4 |
| RP3-395M20.12 | NA |
| OBSL1 | obscurin like cytoskeletal adaptor 1 |
| ZDHHC16 | zinc finger DHHC-type palmitoyltransferase 16 |
| PARD6B | par-6 family cell polarity regulator beta |
| GOLGA8O | golgin A8 family member O |
| RP11-1334A24.5 | NA |
| ADD2 | adducin 2 |
| GDPGP1 | GDP-D-glucose phosphorylase 1 |
| RP11-534C12.1 | NA |
| PLA2G10 | phospholipase A2 group X |
| LINC00685 | NA |
| CD3E | CD3e molecule |
| SHISAL1 | shisa like 1 |
| CD96 | CD96 molecule |
| KCNA5 | potassium voltage-gated channel subfamily A member 5 |
| NCAPG2 | non-SMC condensin II complex subunit G2 |
| AKR1C1 | aldo-keto reductase family 1 member C1 |
| CR2 | complement C3d receptor 2 |
| GNG2 | G protein subunit gamma 2 |
| PTTG1 | PTTG1 regulator of sister chromatid separation, securin |
| RP11-190A12.1( | uncharacterized LOC107985216 |
| PROZ | protein Z, vitamin K dependent plasma glycoprotein |
| TCEAL1 | transcription elongation factor A like 1 |
| RP5-849H19.2 | NA |
| NPRL3 | NPR3 like, GATOR1 complex subunit |
| FCHSD2 | FCH and double SH3 domains 2 |

| | |
|---|---|
| HIC1 | HIC ZBTB transcriptional repressor 1 |
| GPR183 | G protein-coupled receptor 183 |
| AP006621.5 | uncharacterized LOC171391 |
| CALB2 | calbindin 2 |
| FCN1 | ficolin 1 |
| GIMAP7 | GTPase, IMAP family member 7 |
| EBF3 | EBF transcription factor 3 |
| SDCBP2-AS1 | SDCBP2 antisense RNA 1 |
| SMYD1 | SET and MYND domain containing 1 |
| NR1H4 | nuclear receptor subfamily 1 group H member 4 |
| HYAL1 | hyaluronidase 1 |
| CTSO | cathepsin O |
| SLC35E4 | solute carrier family 35 member E4 |
| HOXA-AS2 | NA |
| FIBCD1 | fibrinogen C domain containing 1 |
| GADD45B | growth arrest and DNA damage inducible beta |
| Metazoa_SRP | NA |
| CACNA2D1 | calcium voltage-gated channel auxiliary subunit alpha2delta 1 |
| UFSP1 | UFM1 specific peptidase 1 (inactive) |
| HERC2P2 | HERC2 pseudogene 2 |
| ISG20 | interferon stimulated exonuclease gene 20 |
| DAPK1 | death associated protein kinase 1 |
| GAP43 | growth associated protein 43 |
| SEPT5-GP1BB | SEPT5-GP1BB readthrough |
| SEMA3D | semaphorin 3D |
| bP-2189O9.2 | NA |
| XKR4 | XK related 4 |
| ARNT2 | aryl hydrocarbon receptor nuclear translocator 2 |
| KIFC1 | kinesin family member C1 |
| LRRC34 | leucine rich repeat containing 34 |
| BST1 | bone marrow stromal cell antigen 1 |
| APOC1 | apolipoprotein C1 |
| FAT3 | FAT atypical cadherin 3 |

| | |
|---|---|
| CH507-154B10.1 | NA |
| LYRM9 | LYR motif containing 9 |
| LYPD1 | LY6/PLAUR domain containing 1 |
| HOXB-AS1 | HOXB cluster antisense RNA 1 |
| MAP7D3 | MAP7 domain containing 3 |
| NOS3 | nitric oxide synthase 3 |
| RP3-403A15.5 | SOGA family member 3 |
| ACKR4 | atypical chemokine receptor 4 |
| LINC02731 | long intergenic non-protein coding RNA 2731 |
| CPED1 | cadherin like and PC-esterase domain containing 1 |
| RP13-1032I1.11 | NA |
| DHRS3 | dehydrogenase/reductase 3 |
| FAM214B | family with sequence similarity 214 member B |
| DKK2 | dickkopf WNT signaling pathway inhibitor 2 |
| FERMT1 | FERM domain containing kindlin 1 |
| CTAGE9 | CTAGE family member 9 |
| CD69 | CD69 molecule |
| KCNQ4 | potassium voltage-gated channel subfamily Q member 4 |
| SFT2D1 | SFT2 domain containing 1 |
| RACGAP1 | Rac GTPase activating protein 1 |
| PDPN | podoplanin |
| RP11-119F19.5 | NA |
| CALHM5 | calcium homeostasis modulator family member 5 |
| SLC49A4 | solute carrier family 49 member 4 |
| ARHGEF38 | Rho guanine nucleotide exchange factor 38 |
| CSF2RA | NA |
| APBB1 | amyloid beta precursor protein binding family B member 1 |
| PLAAT5 | phospholipase A and acyltransferase 5 |
| LRCH2 | leucine rich repeats and calponin homology domain containing 2 |
| ABTB1 | ankyrin repeat and BTB domain containing 1 |
| RGMB | repulsive guidance molecule BMP co-receptor b |
| VAMP1 | vesicle associated membrane protein 1 |
| Metazoa_SRP | NA |

| | |
|---|---|
| CDC16 | cell division cycle 16 |
| PTPRU | protein tyrosine phosphatase receptor type U |
| TRPA1 | transient receptor potential cation channel subfamily A member 1 |
| TOR1AIP1 | torsin 1A interacting protein 1 |
| E2F2 | E2F transcription factor 2 |
| GPLD1 | glycosylphosphatidylinositol specific phospholipase D1 |
| KDSR-DT | NA |
| NMNAT2 | nicotinamide nucleotide adenylyltransferase 2 |
| LINC01934 | NA |
| IL3RA | interleukin 3 receptor subunit alpha |
| USP13 | ubiquitin specific peptidase 13 |
| ENSG00000277[illegible] | proline dehydrogenase 1, mitochondrial |
| STEAP1B | STEAP family member 1B |
| RP11-616M22.1[illegible] | NA |
| STUM | stum, mechanosensory transduction mediator homolog |
| PARD3B | par-3 family cell polarity regulator beta |
| KCNJ2 | potassium inwardly rectifying channel subfamily J member 2 |
| LINC-PINT | long intergenic non-protein coding RNA, p53 induced transcript |
| KLF11 | Kruppel like factor 11 |
| CFAP65 | cilia and flagella associated protein 65 |
| HERC5 | HECT and RLD domain containing E3 ubiquitin protein ligase 5 |
| RN7SL73P | RNA, 7SL, cytoplasmic 73, pseudogene |
| FAM13C | family with sequence similarity 13 member C |
| DHRSX | dehydrogenase/reductase X-linked |
| COL9A1 | collagen type IX alpha 1 chain |
| ACSS3 | acyl-CoA synthetase short chain family member 3 |
| IGSF10 | immunoglobulin superfamily member 10 |
| MPZ | myelin protein zero |
| NKD2 | NKD inhibitor of WNT signaling pathway 2 |
| TUB | TUB bipartite transcription factor |
| CLDN5 | claudin 5 |
| FLG-AS1 | FLG antisense RNA 1 |
| BCAT1 | branched chain amino acid transaminase 1 |

| | |
|---|---|
| WDFY4 | WDFY family member 4 |
| SDCBP2 | syndecan binding protein 2 |
| ZNF358 | zinc finger protein 358 |
| TSPAN4 | tetraspanin 4 |
| BIRC5 | baculoviral IAP repeat containing 5 |
| SRMS | src-related kinase lacking C-terminal regulatory tyrosine and N-terminal myristylation sites |
| LRAT | lecithin retinol acyltransferase |
| P4HTM | prolyl 4-hydroxylase, transmembrane |
| FBXL2 | F-box and leucine rich repeat protein 2 |
| ZNF816 | zinc finger protein 816 |
| NEUROD1 | neuronal differentiation 1 |
| AGBL3 | AGBL carboxypeptidase 3 |
| MRPL44 | mitochondrial ribosomal protein L44 |
| SNORD13 | small nucleolar RNA, C/D box 13 |
| GRIA4 | glutamate ionotropic receptor AMPA type subunit 4 |
| MIR4453HG | MIR4453 host gene |
| EGFL8 | EGF like domain multiple 8 |
| RELN | reelin |
| GASK1B | golgi associated kinase 1B |
| LINC01002 | NA |
| PIANP | PILR alpha associated neural protein |
| OR7E47P | olfactory receptor family 7 subfamily E member 47 pseudogene |
| CCDC107 | coiled-coil domain containing 107 |
| TAMALIN | trafficking regulator and scaffold protein tamalin |
| LINC01559 | long intergenic non-protein coding RNA 1559 |
| RASSF3 | Ras association domain family member 3 |
| CHADL | chondroadherin like |
| CCL26 | C-C motif chemokine ligand 26 |
| FGFR4 | fibroblast growth factor receptor 4 |
| CRABP2 | cellular retinoic acid binding protein 2 |
| CFAP91 | cilia and flagella associated protein 91 |
| ZNF658 | zinc finger protein 658 |
| KCNJ12 | potassium inwardly rectifying channel subfamily J member 12 |

| | |
|---|---|
| SOX9 | SRY-box transcription factor 9 |
| ENDOV | endonuclease V |
| ARRDC1-AS1 | ARRDC1 antisense RNA 1 |
| TRIP13 | thyroid hormone receptor interactor 13 |
| SOX7 | SRY-box transcription factor 7 |
| RP11-245P10.9 | NA |
| ORM1 | orosomucoid 1 |
| KLRB1 | killer cell lectin like receptor B1 |
| ZCWPW1 | zinc finger CW-type and PWWP domain containing 1 |
| BHLHB9 | basic helix-loop-helix family member b9 |
| DBH-AS1 | DBH antisense RNA 1 |
| Y_RNA | NA |
| GJB3 | gap junction protein beta 3 |
| ORC5 | origin recognition complex subunit 5 |
| OVOS | NA |
| NRIP2 | nuclear receptor interacting protein 2 |
| STAC | SH3 and cysteine rich domain |
| FGFR2 | fibroblast growth factor receptor 2 |
| FBXO41 | F-box protein 41 |
| ZC3H3 | zinc finger CCCH-type containing 3 |
| APOB | apolipoprotein B |
| RP11-48B3.4 | NA |
| RAB25 | RAB25, member RAS oncogene family |
| FAM182B | NA |
| BEND5 | BEN domain containing 5 |
| SULT1C4 | sulfotransferase family 1C member 4 |
| PSAT1 | phosphoserine aminotransferase 1 |
| PRICKLE1 | prickle planar cell polarity protein 1 |
| INSC | INSC spindle orientation adaptor protein |
| CBX2 | chromobox 2 |
| SNX24 | sorting nexin 24 |
| NBPF20 | NBPF member 20 |
| SCN8A | sodium voltage-gated channel alpha subunit 8 |

| | |
|---|---|
| SLC18A2 | solute carrier family 18 member A2 |
| RNU11 | RNA, U11 small nuclear |
| CH17-408M7.2 | uncharacterized LOC100996756 |
| LRRC17 | leucine rich repeat containing 17 |
| COLCA1 | colorectal cancer associated 1 |
| NEK4 | NIMA related kinase 4 |
| NPAS4 | neuronal PAS domain protein 4 |
| LINC00562 | long intergenic non-protein coding RNA 562 |
| WNT4 | Wnt family member 4 |
| CTD-3126B10.2 | NA |
| MAIP1 | matrix AAA peptidase interacting protein 1 |
| MAGED4B | MAGE family member D4B |
| ITGB1BP2 | integrin subunit beta 1 binding protein 2 |
| ADA2 | adenosine deaminase 2 |
| SYDE2 | synapse defective Rho GTPase homolog 2 |
| PDE6B | phosphodiesterase 6B |
| LRRN2 | leucine rich repeat neuronal 2 |
| ZCWPW2 | zinc finger CW-type and PWWP domain containing 2 |
| GPT | glutamic--pyruvic transaminase |

| ID | Description | setSize | enrichmentS | NES | pvalue | p.adjust | qvalues | rank | leading_edg |
|---|---|---|---|---|---|---|---|---|---|
| hsa04270 | Vascular smooth m | 119 | 0.59581469 | 2.28412857 | 1.08E-10 | 2.66E-08 | 1.73E-08 | 2328 | tags=33% |
| hsa04110 | Cell cycle | 123 | -0.5157526 | -2.3154016 | 1.59E-10 | 2.66E-08 | 1.73E-08 | 4353 | tags=53% |
| hsa04022 | cGMP-PKG signal | 149 | 0.54706662 | 2.15524236 | 5.26E-09 | 5.87E-07 | 3.82E-07 | 2653 | tags=34% |
| hsa04510 | Focal adhesion | 194 | 0.48037868 | 1.95372946 | 6.81E-08 | 5.71E-06 | 3.71E-06 | 2136 | tags=29% |
| hsa04514 | Cell adhesion mol | 142 | 0.49902277 | 1.95427531 | 9.86E-07 | 6.01E-05 | 3.91E-05 | 3460 | tags=45% |
| hsa05414 | Dilated cardiomyo | 87 | 0.55957122 | 2.04326191 | 1.12E-06 | 6.01E-05 | 3.91E-05 | 1737 | tags=29% |
| hsa04261 | Adrenergic signali | 135 | 0.50238314 | 1.94804215 | 1.43E-06 | 6.01E-05 | 3.91E-05 | 2663 | tags=29% |
| hsa04923 | Regulation of lipol | 51 | 0.63409988 | 2.13019816 | 1.44E-06 | 6.01E-05 | 3.91E-05 | 2653 | tags=41% |
| hsa03013 | Nucleocytoplasmic | 104 | -0.4548858 | -1.9646573 | 2.00E-06 | 7.43E-05 | 4.83E-05 | 5357 | tags=55% |
| hsa04921 | Oxytocin signaling | 139 | 0.48716329 | 1.89888035 | 5.39E-06 | 1.81E-04 | 1.18E-04 | 2867 | tags=31% |
| hsa03050 | Proteasome | 42 | -0.5893928 | -2.1355247 | 8.78E-06 | 2.57E-04 | 1.67E-04 | 5480 | tags=71% |
| hsa04020 | Calcium signaling | 208 | 0.43813512 | 1.79146834 | 9.20E-06 | 2.57E-04 | 1.67E-04 | 3315 | tags=35% |
| hsa03030 | DNA replication | 36 | -0.6106173 | -2.1417729 | 1.20E-05 | 2.92E-04 | 1.90E-04 | 2179 | tags=69% |
| hsa00970 | Aminoacyl-tRNA b | 43 | -0.5803867 | -2.1245871 | 1.29E-05 | 2.92E-04 | 1.90E-04 | 4655 | tags=58% |
| hsa05014 | Amyotrophic later | 329 | -0.3088698 | -1.5573307 | 1.38E-05 | 2.92E-04 | 1.90E-04 | 4038 | tags=32% |
| hsa04024 | cAMP signaling pa | 188 | 0.44036956 | 1.78623185 | 1.40E-05 | 2.92E-04 | 1.90E-04 | 4121 | tags=37% |
| hsa05205 | Proteoglycans in ca | 193 | 0.43268107 | 1.75748670 | 2.06E-05 | 4.06E-04 | 2.64E-04 | 3955 | tags=38% |
| hsa04658 | Th1 and Th2 cell d | 83 | 0.53457300 | 1.93308079 | 2.48E-05 | 4.62E-04 | 3.01E-04 | 2968 | tags=37% |
| hsa04728 | Dopaminergic syna | 124 | 0.47361341 | 1.82163885 | 3.27E-05 | 5.60E-04 | 3.64E-04 | 2663 | tags=32% |
| hsa05410 | Hypertrophic cardi | 83 | 0.53003954 | 1.91668727 | 3.34E-05 | 5.60E-04 | 3.64E-04 | 2329 | tags=30% |
| hsa03040 | Spliceosome | 137 | -0.3867820 | -1.7844416 | 3.86E-05 | 5.99E-04 | 3.90E-04 | 4868 | tags=40% |
| hsa03460 | Fanconi anemia pa | 54 | -0.5228323 | -2.0005239 | 3.93E-05 | 5.99E-04 | 3.90E-04 | 5222 | tags=57% |
| hsa04713 | Circadian entrainm | 85 | 0.52490445 | 1.91093840 | 4.36E-05 | 6.35E-04 | 4.13E-04 | 2820 | tags=41% |
| hsa04145 | Phagosome | 140 | 0.45374926 | 1.77425501 | 4.66E-05 | 6.50E-04 | 4.23E-04 | 2602 | tags=31% |
| hsa03008 | Ribosome biogene | 85 | -0.4483896 | -1.8724615 | 5.97E-05 | 8.00E-04 | 5.20E-04 | 3629 | tags=44% |
| hsa05310 | Asthma | 22 | 0.72749962 | 2.03770607 | 7.15E-05 | 9.21E-04 | 5.99E-04 | 2677 | tags=59% |
| hsa04371 | Apelin signaling pa | 125 | 0.46463416 | 1.78949024 | 7.58E-05 | 9.41E-04 | 6.12E-04 | 2328 | tags=28% |
| hsa05150 | Staphylococcus au | 70 | 0.52006265 | 1.83937514 | 9.08E-05 | 0.00108647 | 7.07E-04 | 2452 | tags=34% |
| hsa01250 | Biosynthesis of nu | 36 | -0.5602527 | -1.9651163 | 1.38E-04 | 0.00159420 | 0.00103692 | 3985 | tags=47% |
| hsa04310 | Wnt signaling path | 158 | 0.42997508 | 1.70099163 | 1.46E-04 | 0.00162622 | 0.00105775 | 2328 | tags=25% |
| hsa04925 | Aldosterone synthe | 88 | 0.49535209 | 1.80864566 | 2.09E-04 | 0.00226207 | 0.00147132 | 4257 | tags=44% |
| hsa04360 | Axon guidance | 175 | 0.40968853 | 1.64009247 | 2.31E-04 | 0.00241797 | 0.00157272 | 3252 | tags=31% |
| hsa01230 | Biosynthesis of am | 69 | -0.4507732 | -1.8066199 | 3.20E-04 | 0.00324387 | 0.00210991 | 3894 | tags=39% |
| hsa04010 | MAPK signaling p | 280 | 0.37789644 | 1.58863613 | 3.34E-04 | 0.00328924 | 0.00213943 | 3587 | tags=31% |
| hsa04725 | Cholinergic synaps | 98 | 0.47295059 | 1.75102872 | 3.72E-04 | 0.00355860 | 0.00231463 | 2663 | tags=33% |
| hsa05416 | Viral myocarditis | 55 | 0.53642846 | 1.81682625 | 3.93E-04 | 0.00365710 | 0.00237869 | 2452 | tags=38% |
| hsa04970 | Salivary secretion | 75 | 0.48798146 | 1.74109168 | 4.94E-04 | 0.00447657 | 0.00291170 | 2328 | tags=32% |
| hsa04657 | IL-17 signaling pa | 87 | -0.3985348 | -1.6792792 | 5.34E-04 | 0.00470987 | 0.00306345 | 4513 | tags=41% |

| | | | | | | | | | | | |
|---|---|---|---|---|---|---|---|---|---|---|---|
| hsa04810 | Regulation of actir | | 202 | 0.39427597 | 1.60778799 | 5.71E-04 | 0.00480083 | 0.00312261 | | 2419 | tags=22% |
| hsa04115 | p53 signaling path | | 73 | -0.4349003 | -1.7637993 | 5.73E-04 | 0.00480083 | 0.00312261 | | 3806 | tags=44% |
| hsa01523 | Antifolate resistan | | 28 | -0.5845685 | -1.9047734 | 7.65E-04 | 0.00624741 | 0.00406351 | | 3459 | tags=61% |
| hsa00051 | Fructose and mann | | 31 | -0.5516984 | -1.8666560 | 8.11E-04 | 0.00646482 | 0.00420492 | | 3057 | tags=61% |
| hsa04961 | Endocrine and oth | | 48 | 0.55230730 | 1.82850915 | 8.38E-04 | 0.00652700 | 0.00424537 | | 2543 | tags=35% |
| hsa03440 | Homologous recor | | 41 | -0.5021758 | -1.8069826 | 8.95E-04 | 0.00681467 | 0.00443248 | | 5684 | tags=63% |
| hsa04924 | Renin secretion | | 63 | 0.51202083 | 1.76600013 | 9.95E-04 | 0.00740365 | 0.00481557 | | 2820 | tags=41% |
| hsa05144 | Malaria | | 46 | 0.55751737 | 1.83207213 | 0.00108061 | 0.00777174 | 0.00505499 | | 2299 | tags=46% |
| hsa05160 | Hepatitis C | | 134 | -0.3438281 | -1.5693805 | 0.00109036 | 0.00777174 | 0.00505499 | | 3319 | tags=32% |
| hsa04974 | Protein digestion a | | 92 | 0.45240760 | 1.67287846 | 0.00148965 | 0.01039657 | 0.00676226 | | 1524 | tags=26% |
| hsa04260 | Cardiac muscle co | | 73 | 0.48217771 | 1.71409699 | 0.00159386 | 0.01070460 | 0.00696261 | | 1819 | tags=23% |
| hsa04740 | Olfactory transduc | | 76 | 0.47227497 | 1.68337509 | 0.00162206 | 0.01070460 | 0.00696261 | | 6127 | tags=42% |
| hsa04512 | ECM-receptor inte | | 79 | 0.47444910 | 1.71295210 | 0.00162965 | 0.01070460 | 0.00696261 | | 3175 | tags=43% |
| hsa04141 | Protein processing | | 163 | -0.31178741 | -1.4644498 | 0.00168904 | 0.01088132 | 0.00707756 | | 4666 | tags=38% |
| hsa04672 | Intestinal immune | | 41 | 0.55448785 | 1.77883705 | 0.00180626 | 0.01141693 | 0.00742593 | | 2452 | tags=32% |
| hsa04926 | Relaxin signaling | | 120 | 0.41630125 | 1.59961557 | 0.00189510 | 0.01157613 | 0.00752948 | | 2653 | tags=28% |
| hsa04659 | Th17 cell different | | 99 | 0.43726647 | 1.62225942 | 0.00190055 | 0.01157613 | 0.00752948 | | 4112 | tags=39% |
| hsa03320 | PPAR signaling pa | | 66 | 0.49469423 | 1.72379470 | 0.00195553 | 0.01169826 | 0.00760892 | | 3804 | tags=39% |
| hsa01200 | Carbon metabolism | | 106 | -0.3660424 | -1.5924001 | 0.00209504 | 0.01231300 | 0.00800877 | | 3894 | tags=33% |
| hsa04726 | Serotonergic synap | | 92 | 0.44149840 | 1.63253924 | 0.00241598 | 0.01395438 | 0.00907637 | | 2328 | tags=27% |
| hsa05140 | Leishmaniasis | | 70 | 0.46083512 | 1.62989720 | 0.00259002 | 0.01470608 | 0.00956530 | | 2616 | tags=34% |
| hsa05222 | Small cell lung car | | 91 | -0.3705376 | -1.5750711 | 0.00267962 | 0.01496124 | 0.00973127 | | 2033 | tags=31% |
| hsa04151 | PI3K-Akt signalin | | 311 | 0.33736546 | 1.43577240 | 0.00290264 | 0.01594078 | 0.01036839 | | 3512 | tags=29% |
| hsa04080 | Neuroactive ligand | | 251 | 0.34833059 | 1.45148295 | 0.00302013 | 0.01631848 | 0.01061406 | | 3117 | tags=24% |
| hsa03410 | Base excision repa | | 33 | -0.5178400 | -1.7815176 | 0.00328677 | 0.01745827 | 0.01135542 | | 3842 | tags=48% |
| hsa01232 | Nucleotide metabo | | 80 | -0.3849216 | -1.5877314 | 0.00333531 | 0.01745827 | 0.01135542 | | 3746 | tags=48% |
| hsa04929 | GnRH secretion | | 58 | 0.48895554 | 1.66891748 | 0.00338903 | 0.01746654 | 0.01136080 | | 2736 | tags=34% |
| hsa05010 | Alzheimer disease | | 344 | -0.2585057 | -1.3002969 | 0.00354058 | 0.01797113 | 0.01168900 | | 4056 | tags=33% |
| hsa00900 | Terpenoid backbo | | 21 | -0.5782841 | -1.7585917 | 0.00378661 | 0.01893309 | 0.01231469 | | 2842 | tags=57% |
| hsa05016 | Huntington diseas | | 275 | -0.2670819 | -1.3288690 | 0.00414143 | 0.02018257 | 0.01312739 | | 4038 | tags=31% |
| hsa04933 | AGE-RAGE signa | | 100 | 0.42027694 | 1.56117027 | 0.00415700 | 0.02018257 | 0.01312739 | | 3343 | tags=39% |
| hsa00520 | Amino sugar and n | | 48 | -0.4335390 | -1.6194666 | 0.00428460 | 0.02032451 | 0.01321971 | | 3985 | tags=38% |
| hsa05130 | Pathogenic Escher | | 185 | -0.2901477 | -1.3757062 | 0.00430758 | 0.02032451 | 0.01321971 | | 3308 | tags=30% |
| hsa04014 | Ras signaling path | | 215 | 0.35769971 | 1.46747079 | 0.00457251 | 0.02075669 | 0.01350081 | | 3559 | tags=31% |
| hsa00240 | Pyrimidine metabo | | 53 | -0.41116068 | -1.5639755 | 0.00457785 | 0.02075669 | 0.01350081 | | 4017 | tags=45% |
| hsa04928 | Parathyroid hormo | secretion ar | | 95 | 0.41837783 | 1.54431230 | 0.00458506 | 0.02075669 | 0.01350081 | | 2583 |
| hsa04340 | Hedgehog signalin | | 53 | 0.48484616 | 1.63452128 | 0.00486018 | 0.02145714 | 0.01395641 | | 2620 | tags=32% |
| hsa04611 | Platelet activation | | 115 | 0.39709635 | 1.51466742 | 0.00486788 | 0.02145714 | 0.01395641 | | 2653 | tags=30% |
| hsa05032 | Morphine addictio | | 78 | 0.44907986 | 1.61036702 | 0.00527217 | 0.02293739 | 0.01491921 | | 1780 | tags=22% |

| | | | | | | | | | |
|---|---|---|---|---|---|---|---|---|---|
| hsa04920 | Adipocytokine sig | 64 | 0.46833129 | 1.62202595 | 0.00538547 | 0.02297789 | 0.01494555 | 2653 | tags=30% |
| hsa00670 | One carbon pool b | 19 | -0.5825998 | -1.7321546 | 0.00541866 | 0.02297789 | 0.01494555 | 2610 | tags=47% |
| hsa05330 | Allograft rejection | 30 | 0.56996780 | 1.71754603 | 0.00551771 | 0.02310541 | 0.01502850 | 2452 | tags=33% |
| hsa04911 | Insulin secretion | 77 | 0.43552716 | 1.55972444 | 0.00565801 | 0.02317973 | 0.01507684 | 2663 | tags=29% |
| hsa04912 | GnRH signaling p | 83 | 0.43842317 | 1.58539136 | 0.00567384 | 0.02317973 | 0.01507684 | 2663 | tags=25% |
| hsa03430 | Mismatch repair | 23 | -0.5510441 | -1.7086385 | 0.00591197 | 0.02386159 | 0.01552034 | 4598 | tags=61% |
| hsa05219 | Bladder cancer | 41 | -0.4493607 | -1.6169378 | 0.00640582 | 0.02554705 | 0.01661662 | 3279 | tags=41% |
| hsa03020 | RNA polymerase | 33 | -0.4960180 | -1.70644361 | 0.00650199 | 0.02562549 | 0.01666764 | 4944 | tags=48% |
| hsa04152 | AMPK signaling p | 112 | 0.39698352 | 1.51033188 | 0.00779667 | 0.03037075 | 0.01975411 | 2026 | tags=21% |
| hsa05320 | Autoimmune thyro | 33 | 0.55309167 | 1.70059717 | 0.00927570 | 0.03571681 | 0.02323136 | 3564 | tags=39% |
| hsa00220 | Arginine biosynthe | 19 | -0.55802261 | -1.6590829 | 0.00956504 | 0.03641237 | 0.02368377 | 3654 | tags=68% |
| hsa05412 | Arrhythmogenic ri | 73 | 0.44393119 | 1.57813415 | 0.00968401 | 0.03645105 | 0.02370893 | 1737 | tags=22% |
| hsa05171 | Coronavirus diseas | 204 | 0.34995759 | 1.43158119 | 0.00999223 | 0.03710487 | 0.02413419 | 3147 | tags=26% |
| hsa04640 | Hematopoietic cell | 83 | 0.42249497 | 1.52779305 | 0.01015036 | 0.03710487 | 0.02413419 | 2795 | tags=37% |
| hsa04730 | Long-term depress | 56 | 0.47676533 | 1.61883728 | 0.01018999 | 0.03710487 | 0.02413419 | 2820 | tags=30% |
| hsa04724 | Glutamatergic syn | 99 | 0.40291675 | 1.49482189 | 0.01073013 | 0.03865155 | 0.02514021 | 2901 | tags=29% |
| hsa04972 | Pancreatic secretio | 86 | 0.41138669 | 1.49870546 | 0.01224441 | 0.04363700 | 0.02838290 | 2507 | tags=26% |
| hsa00512 | Mucin type O-glyc | 32 | -0.46661641 | -1.5881600 | 0.01348636 | 0.04755719 | 0.03093272 | 4762 | tags=50% |
| hsa04727 | GABAergic synap | 76 | 0.42238311 | 1.50554075 | 0.01380185 | 0.04816272 | 0.03132658 | 2583 | tags=24% |
| hsa05322 | Systemic lupus ery | 93 | 0.39920358 | 1.47628499 | 0.01414658 | 0.04885675 | 0.03177799 | 2471 | tags=26% |
| hsa04723 | Retrograde endoca | 124 | 0.36802386 | 1.41551433 | 0.01613412 | 0.05515237 | 0.03587287 | 2594 | tags=23% |
| hsa05169 | Epstein-Barr virus | 183 | -0.2684187 | -1.2758519 | 0.01698940 | 0.05703774 | 0.03709917 | 3360 | tags=29% |
| hsa04015 | Rap1 signaling pat | 198 | 0.33744062 | 1.37465456 | 0.01702619 | 0.05703774 | 0.03709917 | 3827 | tags=32% |
| hsa04950 | Maturity onset dia | 23 | -0.5111385 | -1.58490201 | 0.01836350 | 0.06090864 | 0.03961693 | 2375 | tags=35% |
| hsa05321 | Inflammatory bow | 56 | 0.45673850 | 1.55083699 | 0.01872829 | 0.06150959 | 0.04000781 | 2616 | tags=32% |
| hsa00350 | Tyrosine metabolis | 30 | 0.51633448 | 1.55592689 | 0.01902184 | 0.06186716 | 0.04024038 | 4044 | tags=37% |
| hsa04610 | Complement and c | 71 | 0.41389518 | 1.46733597 | 0.01959141 | 0.06310696 | 0.04104678 | 2491 | tags=27% |
| hsa04910 | Insulin signaling p | 126 | 0.36371585 | 1.39834278 | 0.02069002 | 0.06569258 | 0.04272856 | 2249 | tags=17% |
| hsa04935 | Growth hormone s | secretion ar | 108 | 0.38357304 | 1.44591000 | 0.02078631 | 0.06569258 | 0.04272856 | 2653 |
| hsa05134 | Legionellosis | 55 | -0.3757862 | -1.4375084 | 0.02108636 | 0.06601806 | 0.04294026 | 1510 | tags=18% |
| hsa03450 | Non-homologous e | 12 | -0.6164601 | -1.6259224 | 0.02280205 | 0.07072860 | 0.04600415 | 4081 | tags=67% |
| hsa04971 | Gastric acid secret | 64 | 0.42687491 | 1.47844529 | 0.02520144 | 0.07745397 | 0.05037854 | 2663 | tags=27% |
| hsa04660 | T cell receptor sig | 98 | 0.38466310 | 1.42415752 | 0.02574432 | 0.07840315 | 0.05099592 | 4222 | tags=34% |
| hsa00510 | N-Glycan biosynth | 49 | -0.3742176 | -1.4013624 | 0.02600884 | 0.07849516 | 0.05105577 | 2738 | tags=29% |
| hsa05203 | Viral carcinogenes | 179 | -0.2615994 | -1.2377673 | 0.03009497 | 0.09001621 | 0.05854943 | 3070 | tags=24% |
| hsa04931 | Insulin resistance | 103 | 0.38182888 | 1.43289079 | 0.03293874 | 0.09765026 | 0.06351486 | 2841 | tags=25% |
| hsa05142 | Chagas disease | 97 | 0.37855870 | 1.40004491 | 0.03429355 | 0.10077491 | 0.06554723 | 2839 | tags=30% |
| hsa00100 | Steroid biosynthes | 19 | -0.5074985 | -1.5088674 | 0.03711626 | 0.10812130 | 0.07032555 | 5724 | tags=63% |
| hsa05031 | Amphetamine addi | 62 | 0.42079855 | 1.44918811 | 0.03890489 | 0.11235466 | 0.07307907 | 2663 | tags=32% |

| | | | | | | | | | |
|---|---|---|---|---|---|---|---|---|---|
| hsa05417 | Lipid and atherosc | 194 | -0.2514379 | -1.1946328 | 0.0399215 | 0.1143052 | 0.0743477 | 3319 | tags=28% |
| hsa04211 | Longevity regulati | 83 | 0.38788582 | 1.4026421 | 0.0405594 | 0.1145474 | 0.0745053 | 2653 | tags=20% |
| hsa04960 | Aldosterone-regula | 34 | 0.49762907 | 1.53993331 | 0.0406899 | 0.1145474 | 0.0745053 | 1506 | tags=26% |
| hsa05022 | Pathways of neuro | 431 | -0.2146872 | -1.1122867 | 0.04177742 | 0.1166286 | 0.0758590 | 4047 | tags=29% |
| hsa05200 | Pathways in cance | 487 | 0.2789321 | 1.22462651 | 0.0440386 | 0.1210554 | 0.0787383 | 3512 | tags=26% |
| hsa04064 | NF-kappa B signal | 102 | -0.2898743 | -1.2548466 | 0.0440858 | 0.1210554 | 0.0787383 | 1242 | tags=18% |
| hsa05202 | Transcriptional mi | 170 | 0.3332323 | 1.3271340 | 0.0455696 | 0.1241123 | 0.0807266 | 3348 | tags=25% |
| hsa04670 | Leukocyte transen | 104 | 0.3658963 | 1.37505938 | 0.0467914 | 0.1260331 | 0.0819760 | 2583 | tags=26% |
| hsa03015 | mRNA surveillanc | 91 | -0.3073734 | -1.3065747 | 0.0470272 | 0.1260331 | 0.0819760 | 5633 | tags=46% |
| hsa05145 | Toxoplasmosis | 106 | 0.3617376 | 1.3631062 | 0.0482573 | 0.1283033 | 0.0834525 | 2653 | tags=28% |
| hsa04550 | Signaling pathway | 125 | 0.3451588 | 1.3293435 | 0.0495436 | 0.12974906 | 0.0843929 | 3955 | tags=30% |
| hsa04114 | Oocyte meiosis | 109 | -0.2928894 | -1.2859762 | 0.0495757 | 0.12974906 | 0.0843929 | 3089 | tags=27% |
| hsa04662 | B cell receptor sig | 78 | 0.3833877 | 1.37479981 | 0.0518934 | 0.13450972 | 0.0874894 | 4167 | tags=37% |
| hsa05235 | PD-L1 expression | 87 | 0.3752539 | 1.3702313 | 0.0521978 | 0.13450972 | 0.0874894 | 2839 | tags=24% |
| hsa00410 | beta-Alanine metal | 28 | 0.4929811 | 1.4711624 | 0.0535987 | 0.1370655 | 0.0891518 | 3941 | tags=32% |
| hsa04630 | JAK-STAT signali | 124 | 0.3443119 | 1.32431208 | 0.0549019 | 0.13933452 | 0.0906276 | 3084 | tags=26% |
| hsa05017 | Spinocerebellar ata | 131 | -0.2704029 | -1.2334539 | 0.0565110 | 0.1423398 | 0.0925824 | 4038 | tags=34% |
| hsa04976 | Bile secretion | 66 | 0.3987612 | 1.3895098 | 0.0569800 | 0.1424501 | 0.0926541 | 2687 | tags=29% |
| hsa01240 | Biosynthesis of co | 135 | -0.2671086 | -1.2246338 | 0.0574150 | 0.1424744 | 0.0926699 | 3746 | tags=30% |
| hsa00511 | Other glycan degra | 15 | -0.5242988 | -1.4674821 | 0.0585480 | 0.1442175 | 0.0938036 | 4584 | tags=47% |
| hsa04722 | Neurotrophin signa | 116 | 0.3478543 | 1.3249527 | 0.0608465 | 0.1482035 | 0.0963963 | 2818 | tags=23% |
| hsa00601 | Glycosphingolipid | 26 | -0.4418917 | -1.4261666 | 0.0613742 | 0.1482035 | 0.0963963 | 2054 | tags=38% |
| hsa04927 | Cortisol synthesis | 58 | 0.4116182 | 1.4049474 | 0.06149341 | 0.1482035 | 0.0963963 | 3260 | tags=31% |
| hsa04392 | Hippo signaling pa | 29 | 0.4767684 | 1.42687798 | 0.06832298 | 0.1634871 | 0.10633727 | 4358 | tags=48% |
| hsa05152 | Tuberculosis | 154 | 0.3255881 | 1.2876937 | 0.0691823 | 0.1643695 | 0.1069111 | 2663 | tags=23% |
| hsa00071 | Fatty acid degradat | 39 | 0.4421942 | 1.4114637 | 0.0708898 | 0.1672402 | 0.1087784 | 3804 | tags=36% |
| hsa04213 | Longevity regulati | 57 | 0.4016467 | 1.36710884 | 0.0718475 | 0.1683140 | 0.1094768 | 1566 | tags=18% |
| hsa05418 | Fluid shear stress | 132 | 0.3357686 | 1.2980700 | 0.0737386 | 0.1705407 | 0.1109251 | 3329 | tags=30% |
| hsa04540 | Gap junction | 79 | 0.3671272 | 1.3254771 | 0.0738161 | 0.1705407 | 0.1109251 | 2820 | tags=28% |
| hsa04964 | Proximal tubule bi | 21 | 0.5216326 | 1.4447582 | 0.07580645 | 0.17308189 | 0.1125780 | 1240 | tags=24% |
| hsa05207 | Chemical carcinog | 171 | 0.3220460 | 1.28340302 | 0.0759493 | 0.17308189 | 0.1125780 | 3700 | tags=26% |
| hsa04720 | Long-term potenti | 59 | 0.3966692 | 1.36104529 | 0.0793650 | 0.1796439 | 0.1168461 | 4667 | tags=41% |
| hsa00590 | Arachidonic acid | 55 | 0.4047756 | 1.3709321 | 0.0810014 | 0.1821174 | 0.1184549 | 2903 | tags=25% |
| hsa03060 | Protein export | 23 | -0.4477527 | -1.3883598 | 0.0844327 | 0.1885664 | 0.1226496 | 3552 | tags=30% |
| hsa04930 | Type II diabetes m | 44 | 0.4193010 | 1.3618312 | 0.0851063 | 0.1885730 | 0.1226539 | 2736 | tags=27% |
| hsa00980 | Metabolism of xen | 56 | 0.39746038 | 1.3495605 | 0.08638360 | 0.1885730 | 0.1226539 | 4745 | tags=46% |
| hsa00020 | Citrate cycle (TCA | 29 | -0.4068016 | -1.3561621 | 0.0865921 | 0.1885730 | 0.1226539 | 5188 | tags=52% |
| hsa04710 | Circadian rhythm | 33 | 0.4575532 | 1.4068439 | 0.0866873 | 0.1885730 | 0.1226539 | 3404 | tags=33% |
| hsa04973 | Carbohydrate dige | 41 | 0.4284928 | 1.3746360 | 0.0881458 | 0.1905088 | 0.1239130 | 2653 | tags=27% |

| | | | | | | | | | | |
|---|---|---|---|---|---|---|---|---|---|---|
| hsa04068 | FoxO signaling pa | | 124 | 0.32964398 | 1.26789544 | 0.08888888 | 0.19088319 | 0.12415654 | 3586 | tags=25% |
| hsa05012 | Parkinson disease | | 239 | -0.2285685 | -1.1265052 | 0.08951330 | 0.19099973 | 0.12423234 | 3773 | tags=31% |
| hsa04922 | Glucagon signalin | | 92 | 0.34376366 | 1.27114315 | 0.09239130 | 0.19589295 | 0.12741505 | 3484 | tags=28% |
| hsa05415 | Diabetic cardiomy | | 181 | 0.31275622 | 1.25576834 | 0.09331651 | 0.19661027 | 0.12788162 | 2663 | tags=21% |
| hsa00250 | Alanine | aspartate ar | 34 | -0.3767452 | -1.3094046 | 0.0977011 | 0.20456178 | 0.1330535 | 1341 | |
| hsa04218 | Cellular senescenc | | 149 | -0.2538934 | -1.1788153 | 0.10376018 | 0.21589852 | 0.14042731 | 2468 | tags=20% |
| hsa04625 | C-type lectin recep | | 97 | 0.34300173 | 1.26854258 | 0.10699588 | 0.22125692 | 0.14391259 | 2662 | tags=26% |
| hsa00982 | Drug metabolism - | | 51 | 0.39447918 | 1.32521526 | 0.10787172 | 0.22169954 | 0.14420049 | 3107 | tags=31% |
| hsa04750 | Inflammatory med | | 90 | 0.34288841 | 1.25797219 | 0.10926694 | 0.22319771 | 0.14517495 | 2663 | tags=28% |
| hsa04623 | Cytosolic DNA-se | | 47 | -0.3428425 | -1.2799469 | 0.11550151 | 0.23408219 | 0.15225456 | 5448 | tags=47% |
| hsa00360 | Phenylalanine met | | 14 | 0.54581473 | 1.36454828 | 0.11599297 | 0.23408219 | 0.15225456 | 1715 | tags=21% |
| hsa04979 | Cholesterol metab | | 46 | 0.39332273 | 1.29250789 | 0.12162162 | 0.24201871 | 0.15741672 | 4516 | tags=39% |
| hsa04916 | Melanogenesis | | 91 | 0.33611812 | 1.23913496 | 0.12191780 | 0.24201871 | 0.15741672 | 4577 | tags=34% |
| hsa00514 | Other types of O-g | | 44 | -0.3401417 | -1.2514443 | 0.12209302 | 0.24201871 | 0.15741672 | 4895 | tags=45% |
| hsa01210 | 2-Oxocarboxylic a | | 18 | -0.4477015 | -1.3278393 | 0.12437810 | 0.24509803 | 0.15941962 | 5773 | tags=61% |
| hsa04915 | Estrogen signaling | | 117 | 0.32500000 | 1.23930776 | 0.12796833 | 0.24993631 | 0.16256658 | 2653 | tags=23% |
| hsa05332 | Graft-versus-host | | 31 | 0.44042739 | 1.33450226 | 0.12832550 | 0.24993631 | 0.16256658 | 2452 | tags=32% |
| hsa04668 | TNF signaling pat | | 111 | -0.2716767 | -1.1977666 | 0.13061224 | 0.25291966 | 0.16450705 | 2010 | tags=18% |
| hsa04917 | Prolactin signaling | | 64 | 0.36248230 | 1.25542689 | 0.13467048 | 0.25927938 | 0.16864362 | 2653 | tags=20% |
| hsa04060 | Cytokine-cytokine | | 232 | 0.28545229 | 1.18205455 | 0.14130434 | 0.27049689 | 0.17593984 | 2712 | tags=22% |
| hsa04062 | Chemokine signali | | 179 | 0.29905029 | 1.19830493 | 0.14501891 | 0.27603032 | 0.17953896 | 2712 | tags=23% |
| hsa00340 | Histidine metaboli | | 22 | 0.47505869 | 1.33062611 | 0.1488 | 0.28159552 | 0.18315875 | 3870 | tags=41% |
| hsa01521 | EGFR tyrosine kin | | 78 | 0.34298733 | 1.22992709 | 0.15147265 | 0.28159552 | 0.18315875 | 3062 | tags=24% |
| hsa04217 | Necroptosis | | 129 | -0.2518361 | -1.1509216 | 0.15189873 | 0.28159552 | 0.18315875 | 4487 | tags=30% |
| hsa05100 | Bacterial invasion | | 76 | 0.34499909 | 1.22971342 | 0.15211267 | 0.28159552 | 0.18315875 | 2419 | tags=22% |
| hsa04380 | Osteoclast differen | | 122 | 0.31287168 | 1.20156692 | 0.15214564 | 0.28159552 | 0.18315875 | 4112 | tags=34% |
| hsa04061 | Viral protein intera | | 93 | -0.2765383 | -1.1753734 | 0.15355805 | 0.28264806 | 0.18384335 | 1586 | tags=19% |
| hsa04120 | Ubiquitin mediate | | 141 | -0.2546386 | -1.1759272 | 0.15513660 | 0.28399323 | 0.18471830 | 5029 | tags=32% |
| hsa05218 | Melanoma | | 66 | 0.35207501 | 1.22682862 | 0.15669515 | 0.28528737 | 0.18556005 | 3060 | tags=27% |
| hsa04140 | Autophagy - anima | | 135 | 0.30747409 | 1.19226234 | 0.15891472 | 0.28776450 | 0.18717125 | 4033 | tags=26% |
| hsa04066 | HIF-1 signaling pa | | 101 | -0.2651843 | -1.1465833 | 0.16030534 | 0.28872198 | 0.18779403 | 3279 | tags=26% |
| hsa00830 | Retinol metabolism | | 47 | 0.37155903 | 1.22579309 | 0.16493313 | 0.29318098 | 0.19069431 | 3167 | tags=30% |
| hsa04530 | Tight junction | | 153 | 0.30070505 | 1.18701625 | 0.16561314 | 0.29318098 | 0.19069431 | 2377 | tags=18% |
| hsa04919 | Thyroid hormone s | | 114 | 0.31289904 | 1.19575682 | 0.16601307 | 0.29318098 | 0.19069431 | 3866 | tags=27% |
| hsa00730 | Thiamine metaboli | | 14 | -0.4662070 | -1.2812161 | 0.16628175 | 0.29318098 | 0.19069431 | 5050 | tags=50% |
| hsa04918 | Thyroid hormone s | | 68 | 0.34509149 | 1.21201611 | 0.18169014 | 0.31867118 | 0.20727395 | 2546 | tags=25% |
| hsa05166 | Human T-cell leuk | | 213 | 0.28151925 | 1.15481391 | 0.18359853 | 0.31881146 | 0.20736519 | 2833 | tags=23% |
| hsa04978 | Mineral absorptior | | 51 | 0.36032708 | 1.21048453 | 0.18367346 | 0.31881146 | 0.20736519 | 1520 | tags=18% |
| hsa00513 | Various types of N | | 37 | -0.3346264 | -1.1829003 | 0.18713450 | 0.32261947 | 0.20984204 | 1286 | tags=19% |

| | | | | | | | | | |
|---|---|---|---|---|---|---|---|---|---|
| hsa05167 | Kaposi sarcoma-as | 169 | -0.23772554 | -1.11615994 | 0.18779342 | 0.32261947 | 0.20984204 | 3094 | tags=26% |
| hsa04216 | Ferroptosis | 39 | -0.32509557 | -1.15092329 | 0.18879056 | 0.32267774 | 0.20987994 | 2586 | tags=49% |
| hsa04977 | Vitamin digestion a | 23 | 0.44988291 | 1.27007066 | 0.19101123 | 0.32298118 | 0.21007731 | 2826 | tags=39% |
| hsa04612 | Antigen processing | 62 | 0.34909912 | 1.20226248 | 0.19164265 | 0.32298118 | 0.21007731 | 2602 | tags=26% |
| hsa03420 | Nucleotide excisio | 44 | -0.31958191 | -1.17580099 | 0.19186046 | 0.32298118 | 0.21007731 | 4226 | tags=36% |
| hsa05143 | African trypanosor | 33 | 0.39784469 | 1.22325756 | 0.19504643 | 0.32520325 | 0.21152261 | 2328 | tags=33% |
| hsa03018 | RNA degradation | 75 | -0.28319932 | -1.15808947 | 0.19512195 | 0.32520325 | 0.21152261 | 4411 | tags=29% |
| hsa00534 | Glycosaminoglyca | 23 | -0.39580800 | -1.22729326 | 0.19788918 | 0.32818255 | 0.21346045 | 3783 | tags=57% |
| hsa04137 | Mitophagy - anima | 69 | 0.33606898 | 1.18570156 | 0.19887955 | 0.32820024 | 0.21347195 | 4033 | tags=32% |
| hsa00533 | Glycosaminoglyca | 14 | -0.44362711 | -1.21916272 | 0.21016166 | 0.34511841 | 0.22447607 | 6523 | tags=71% |
| hsa00531 | Glycosaminoglyca | 18 | -0.41761951 | -1.23861888 | 0.21144278 | 0.34552845 | 0.22474278 | 4148 | tags=44% |
| hsa00380 | Tryptophan metabo | 39 | 0.38026138 | 1.21377682 | 0.21417797 | 0.34829914 | 0.22654492 | 5006 | tags=38% |
| hsa04070 | Phosphatidylinosit | 92 | 0.31196611 | 1.15356459 | 0.22554347 | 0.36500997 | 0.23741418 | 4701 | tags=34% |
| hsa00330 | Arginine and proli | 46 | -0.30137399 | -1.12562374 | 0.23511904 | 0.37867731 | 0.24630386 | 1395 | tags=22% |
| hsa03250 | Viral life cycle - H | 62 | 0.33534199 | 1.15488428 | 0.23631123 | 0.37877638 | 0.24636830 | 2099 | tags=18% |
| hsa01522 | Endocrine resistan | 94 | 0.30774307 | 1.13647147 | 0.24523160 | 0.39070599 | 0.25412769 | 1776 | tags=15% |
| hsa00591 | Linoleic acid meta | 25 | 0.40955185 | 1.18277588 | 0.24722662 | 0.39070599 | 0.25412769 | 2903 | tags=32% |
| hsa05323 | Rheumatoid arthrit | 85 | 0.31245946 | 1.13752280 | 0.24725274 | 0.39070599 | 0.25412769 | 2602 | tags=28% |
| hsa04975 | Fat digestion and a | 38 | 0.37122071 | 1.17764851 | 0.25188536 | 0.39449603 | 0.25659286 | 3604 | tags=32% |
| hsa00592 | alpha-Linolenic ac | 23 | 0.41879423 | 1.18230379 | 0.25200642 | 0.39449603 | 0.25659286 | 4242 | tags=43% |
| hsa05162 | Measles | 122 | -0.24762341 | -1.10693589 | 0.25321888 | 0.39455035 | 0.25662819 | 3077 | tags=33% |
| hsa05163 | Human cytomegalo | 200 | 0.27271609 | 1.11030356 | 0.25870646 | 0.40123456 | 0.26097582 | 2653 | tags=20% |
| hsa05212 | Pancreatic cancer | 76 | -0.26762702 | -1.10575335 | 0.26369863 | 0.40636067 | 0.26431000 | 2033 | tags=21% |
| hsa00620 | Pyruvate metabolis | 41 | 0.35947705 | 1.15322833 | 0.26443769 | 0.40636067 | 0.26431000 | 3722 | tags=27% |
| hsa05164 | Influenza A | 149 | 0.28049186 | 1.10503533 | 0.27122940 | 0.41409147 | 0.26933836 | 2672 | tags=21% |
| hsa04614 | Renin-angiotensin | 21 | 0.42016634 | 1.16372850 | 0.27258064 | 0.41409147 | 0.26933836 | 1684 | tags=24% |
| hsa05170 | Human immunode | 189 | 0.27246225 | 1.10377269 | 0.27317676 | 0.41409147 | 0.26933836 | 2653 | tags=21% |
| hsa00562 | Inositol phosphate | 70 | 0.32154192 | 1.13724031 | 0.27552447 | 0.41437487 | 0.26952269 | 4522 | tags=31% |
| hsa00430 | Taurine and hypota | 13 | 0.48488788 | 1.17669102 | 0.27667269 | 0.41437487 | 0.26952269 | 614 | tags=15% |
| hsa05133 | Pertussis | 72 | 0.31786446 | 1.12744065 | 0.27707454 | 0.41437487 | 0.26952269 | 2662 | tags=28% |
| hsa05221 | Acute myeloid leul | 65 | 0.32074948 | 1.11833798 | 0.27982954 | 0.41663510 | 0.27099282 | 3586 | tags=22% |
| hsa04350 | TGF-beta signaling | 89 | 0.30429154 | 1.11434176 | 0.28116343 | 0.41676880 | 0.27107979 | 3358 | tags=26% |
| hsa05230 | Central carbon me | 67 | -0.27408977 | -1.09443067 | 0.28571428 | 0.41854072 | 0.27223230 | 3870 | tags=33% |
| hsa05161 | Hepatitis B | 146 | -0.23274920 | -1.07390286 | 0.28638497 | 0.41854072 | 0.27223230 | 3182 | tags=23% |
| hsa00480 | Glutathione metab | 52 | -0.29128074 | -1.09486391 | 0.28662420 | 0.41854072 | 0.27223230 | 2827 | tags=31% |
| hsa04664 | Fc epsilon RI signa | 63 | 0.32293664 | 1.11383389 | 0.28735632 | 0.41854072 | 0.27223230 | 3065 | tags=25% |
| hsa00650 | Butanoate metabol | 24 | -0.35961093 | -1.12871649 | 0.29442970 | 0.42571787 | 0.27690054 | 6942 | tags=58% |
| hsa05132 | Salmonella infectio | 242 | 0.25901567 | 1.07546754 | 0.29482551 | 0.42571787 | 0.27690054 | 2928 | tags=17% |
| hsa04962 | Vasopressin-regula | 40 | 0.35011783 | 1.12139392 | 0.29713423 | 0.42721017 | 0.27787118 | 3942 | tags=30% |

| | | | | | | | | | | |
|---|---|---|---|---|---|---|---|---|---|---|
| hsa00190 | Oxidative phospho | | 113 | -0.2411164 | -1.0641108 | 0.3034188 | 0.4343816 | 0.2825357 | 6004 | tags=42% |
| hsa02010 | ABC transporters | | 43 | 0.34534483 | 1.11807632 | 0.3068702 | 0.43745330 | 0.2845336 | 1725 | tags=21% |
| hsa04936 | Alcoholic liver dis | | 123 | 0.28190477 | 1.0825319 | 0.3111979 | 0.44174280 | 0.2873236 | 3817 | tags=27% |
| hsa00563 | Glycosylphosphati | | 26 | -0.3363401 | -1.0855081 | 0.3178082 | 0.4492225 | 0.2921887 | 1561 | tags=54% |
| hsa05146 | Amoebiasis | | 88 | 0.2968286 | 1.08379041 | 0.3259361 | 0.4587757 | 0.2984024 | 2350 | tags=27% |
| hsa05110 | Vibrio cholerae inf | | 47 | -0.2870804 | -1.0717683 | 0.3343465 | 0.4686446 | 0.30482152 | 2778 | tags=34% |
| hsa04650 | Natural killer cell r | | 99 | 0.2897319 | 1.0749059 | 0.3374316 | 0.4709984 | 0.3063524 | 4112 | tags=30% |
| hsa00053 | Ascorbate and alda | | 18 | 0.4218135 | 1.12109855 | 0.34 | 0.4726141 | 0.3074033 | 5437 | tags=44% |
| hsa05220 | Chronic myeloid le | | 76 | -0.2566403 | -1.0603595 | 0.34589041 | 0.4788152 | 0.3114367 | 3077 | tags=25% |
| hsa05204 | Chemical carcinog | | 49 | 0.3239570 | 1.0802244 | 0.3515509 | 0.4846484 | 0.3152308 | 4671 | tags=39% |
| hsa04613 | Neutrophil extrace | | 145 | 0.2689274 | 1.0573859 | 0.3549618 | 0.4873451 | 0.3169848 | 2925 | tags=21% |
| hsa04071 | Sphingolipid signa | | 117 | 0.27412975 | 1.0453265 | 0.3588390 | 0.4906574 | 0.3191393 | 3473 | tags=26% |
| hsa01212 | Fatty acid metabol | | 54 | -0.2753880 | -1.0537227 | 0.3625 | 0.4936483 | 0.3210847 | 4309 | tags=33% |
| hsa00630 | Glyoxylate and dic | | 28 | -0.3218517 | -1.0487302 | 0.3667621 | 0.49743048 | 0.3235447 | 3406 | tags=25% |
| hsa05020 | Prion disease | | 243 | 0.2499921 | 1.0380918 | 0.3762019 | 0.5081759 | 0.3305339 | 2213 | tags=16% |
| hsa04072 | Phospholipase D si | | 139 | 0.2673155 | 1.0419510 | 0.3814432 | 0.51114690 | 0.3324663 | 2928 | tags=22% |
| hsa05224 | Breast cancer | | 135 | 0.2684293 | 1.0408623 | 0.3824289 | 0.51114690 | 0.3324663 | 3671 | tags=25% |
| hsa04940 | Type I diabetes me | | 36 | 0.3356001 | 1.0509463 | 0.38297872 | 0.51114690 | 0.3324663 | 2703 | tags=28% |
| hsa05231 | Choline metabolism | | 95 | 0.2842013 | 1.0490414 | 0.3857729 | 0.5128330 | 0.33356304 | 2809 | tags=22% |
| hsa01524 | Platinum drug resis | | 70 | -0.2574969 | -1.0325185 | 0.4006968 | 0.5303190 | 0.3449365 | 4042 | tags=37% |
| hsa05030 | Cocaine addiction | | 45 | 0.3173942 | 1.0415547 | 0.40209267 | 0.5303190 | 0.3449365 | 4283 | tags=38% |
| hsa05206 | MicroRNAs in can | | 158 | 0.2579308 | 1.0203806 | 0.40533672 | 0.5325011 | 0.3463558 | 3437 | tags=27% |
| hsa04913 | Ovarian steroidoge | | 42 | 0.3198841 | 1.0314557 | 0.4106870 | 0.53742247 | 0.3495567 | 1323 | tags=12% |
| hsa00532 | Glycosaminoglyca | | 20 | 0.38196164 | 1.0394993 | 0.4155629 | 0.5401430 | 0.3513263 | 4338 | tags=40% |
| hsa05165 | Human papillomav | | 301 | 0.2411851 | 1.0203378 | 0.41599073 | 0.5401430 | 0.3513263 | 3175 | tags=20% |
| hsa04150 | mTOR signaling pa | | 147 | 0.2597880 | 1.0222199 | 0.4183673 | 0.5411315 | 0.3519692 | 3971 | tags=24% |
| hsa05226 | Gastric cancer | | 137 | -0.2199344 | -1.0146803 | 0.4235807 | 0.54576755 | 0.3549847 | 3398 | tags=26% |
| hsa05120 | Epithelial cell sign | | 68 | -0.2532295 | -1.0133487 | 0.42808219 | 0.5485554 | 0.3567980 | 2777 | tags=25% |
| hsa04621 | NOD-like receptor | | 155 | -0.2187014 | -1.0132672 | 0.4292452 | 0.5485554 | 0.3567980 | 4565 | tags=30% |
| hsa04914 | Progesterone-medi | | 85 | -0.2427433 | -1.0136889 | 0.4306569 | 0.5485554 | 0.3567980 | 3666 | tags=28% |
| hsa05217 | Basal cell carcinom | | 58 | 0.30010934 | 1.02434207 | 0.43631039 | 0.55219010 | 0.35916214 | 3955 | tags=34% |
| hsa00920 | Sulfur metabolism | | 10 | -0.4102177 | -1.014513 | 0.43680709 | 0.55219010 | 0.35916214 | 5408 | tags=70% |
| hsa04721 | Synaptic vesicle cy | | 71 | 0.2817888 | 0.9989944 | 0.4565826 | 0.57501948 | 0.3740111 | 2177 | tags=17% |
| hsa00280 | Valine | leucine and | 48 | -0.2639259 | -0.9858838 | 0.4589665 | 0.5758569 | 0.37455580 | 2796 | |
| hsa04966 | Collecting duct aci | | 22 | 0.35260285 | 0.98763071 | 0.464 | 0.58 | 0.3772505 | 2218 | tags=18% |
| hsa04932 | Non-alcoholic fatty | | 145 | 0.2521699 | 0.9914979 | 0.4758269 | 0.5894810 | 0.3834173 | 2653 | tags=19% |
| hsa05135 | Yersinia infection | | 134 | 0.2549943 | 0.9880180 | 0.4767441 | 0.5894810 | 0.3834173 | 2662 | tags=19% |
| hsa05034 | Alcoholism | | 140 | 0.2529444 | 0.9890661 | 0.4768637 | 0.5894810 | 0.3834173 | 4283 | tags=29% |
| hsa04934 | Cushing syndrome | | 143 | 0.2494071 | 0.9797162 | 0.4879594 | 0.6009794 | 0.3908963 | 3980 | tags=28% |

| | | | | | | | | | |
|---|---|---|---|---|---|---|---|---|---|
| hsa00030 | Pentose phosphate | 27 | -0.3025689( | -0.9824381{ | 0.49428571 | 0.60654107 | 0.39451375 | 4589 | tags=37% |
| hsa00760 | Nicotinate and nic | 32 | 0.30990844 | 0.94543265 | 0.50773993 | 0.62026688 | 0.40344146 | 4738 | tags=38% |
| hsa05340 | Primary immunod | 34 | 0.30720533 | 0.95065934 | 0.50917431 | 0.62026688 | 0.40344146 | 2267 | tags=18% |
| hsa00260 | Glycine | serine and t | 37 | 0.30717855 | 0.96495409 | 0.51212121 | 0.62159639 | 0.40430622 | 2738 |
| hsa00061 | Fatty acid biosynth | 16 | -0.33272077 | -0.9481538 | 0.51566265 | 0.62363533 | 0.40563240 | 3918 | tags=44% |
| hsa05131 | Shigellosis | 229 | 0.23574345 | 0.97785175 | 0.51985559 | 0.62644469 | 0.40745970 | 2662 | tags=18% |
| hsa03010 | Ribosome | 135 | 0.24850463 | 0.96360219 | 0.53359173 | 0.64069258 | 0.41672698 | 3129 | tags=20% |
| hsa00230 | Purine metabolism | 117 | 0.25187281 | 0.96045516 | 0.54089709 | 0.64714474 | 0.42092368 | 3542 | tags=22% |
| hsa04622 | RIG-I-like recepto | 52 | -0.25373179 | -0.9537251 | 0.54777070 | 0.65303624 | 0.42475570 | 5432 | tags=40% |
| hsa05168 | Herpes simplex vi | 457 | -0.1907356 | -0.9855723 | 0.55072463 | 0.65422962 | 0.42553191 | 3011 | tags=20% |
| hsa04210 | Apoptosis | 131 | 0.24625100 | 0.95222330 | 0.56144890 | 0.66461265 | 0.43228537 | 2653 | tags=17% |
| hsa04390 | Hippo signaling pa | 151 | 0.24036183 | 0.94947688 | 0.56855345 | 0.67065284 | 0.43621410 | 3955 | tags=28% |
| hsa00010 | Glycolysis / Gluco | 58 | -0.24185012 | -0.9372260 | 0.57366771 | 0.67431116 | 0.43859359 | 2691 | tags=22% |
| hsa00062 | Fatty acid elongati | 26 | -0.2820114{ | -0.9101671( | 0.58082191 | 0.68033336 | 0.44251062 | 4309 | tags=35% |
| hsa04744 | Phototransduction | 18 | 0.33336250 | 0.88601292 | 0.60333333 | 0.70423925 | 0.45805978 | 4257 | tags=39% |
| hsa00140 | Steroid hormone b | 45 | 0.27388682 | 0.89878156 | 0.62929745 | 0.73199530 | 0.47611320 | 4469 | tags=29% |
| hsa05225 | Hepatocellular car | 159 | -0.2030060{ | -0.9469482 | 0.64055299 | 0.74250952 | 0.48295199 | 3398 | tags=25% |
| hsa04215 | Apoptosis - multip | 31 | -0.2625321{ | -0.8882702 | 0.65289256 | 0.75289631 | 0.48970789 | 3420 | tags=35% |
| hsa05214 | Glioma | 73 | 0.25327298 | 0.90036193 | 0.65400843 | 0.75289631 | 0.48970789 | 2833 | tags=21% |
| hsa03022 | Basal transcription | 43 | -0.2452099 | -0.8976253( | 0.65706051 | 0.75381943 | 0.49030831 | 7446 | tags=53% |
| hsa05033 | Nicotine addiction | 32 | -0.2584625( | -0.8796943 | 0.66011235 | 0.75473597 | 0.49090446 | 1038 | tags=9% |
| hsa05215 | Prostate cancer | 94 | 0.24230659 | 0.89481958 | 0.67029972 | 0.76377690 | 0.49678497 | 2833 | tags=21% |
| hsa04742 | Taste transduction | 47 | 0.26373189 | 0.87006558 | 0.67607726 | 0.76774875 | 0.49936839 | 2328 | tags=19% |
| hsa04370 | VEGF signaling pa | 57 | 0.25087572 | 0.85392047 | 0.69648093 | 0.78824700 | 0.51270111 | 2653 | tags=21% |
| hsa04620 | Toll-like receptor s | 87 | 0.23918350 | 0.87337327 | 0.70054945 | 0.79018204 | 0.51395972 | 2653 | tags=17% |
| hsa00565 | Ether lipid metabo | 47 | 0.25927755 | 0.85537050 | 0.70579494 | 0.79342720 | 0.51607048 | 2903 | tags=19% |
| hsa00052 | Galactose metabol | 29 | -0.2623006 | -0.8744363 | 0.71508379 | 0.80118084 | 0.52111369 | 3984 | tags=24% |
| hsa00770 | Pantothenate and C | 19 | 0.29869009 | 0.80656491 | 0.72577996 | 0.80718586 | 0.52501955 | 4972 | tags=42% |
| hsa04012 | ErbB signaling pat | 83 | 0.23823531 | 0.86148777 | 0.72587412 | 0.80718586 | 0.52501955 | 3051 | tags=17% |
| hsa05213 | Endometrial cance | 58 | 0.24482998 | 0.83566093 | 0.72767203 | 0.80718586 | 0.52501955 | 1967 | tags=16% |
| hsa00515 | Mannose type O-g | 22 | -0.2641503 | -0.8091855( | 0.73740053 | 0.81527781 | 0.53028282 | 4118 | tags=36% |
| hsa04520 | Adherens junction | 71 | 0.23807948 | 0.84403638 | 0.74229691 | 0.81557377 | 0.53047531 | 2825 | tags=18% |
| hsa05208 | Chemical carcinog | 198 | 0.21550230 | 0.87790621 | 0.74253731 | 0.81557377 | 0.53047531 | 2653 | tags=16% |
| hsa05210 | Colorectal cancer | 86 | 0.23282333 | 0.84818885 | 0.74552957 | 0.81618433 | 0.53087245 | 1928 | tags=14% |
| hsa05211 | Renal cell carcino | 68 | 0.23734274 | 0.83358541 | 0.75633802 | 0.82270645 | 0.53511464 | 3586 | tags=22% |
| hsa00310 | Lysine degradation | 61 | -0.2192797 | -0.8524431 | 0.75907590 | 0.82270645 | 0.53511464 | 4339 | tags=28% |
| hsa04136 | Autophagy - other | 30 | 0.26691031 | 0.80430990 | 0.75975039 | 0.82270645 | 0.53511464 | 3996 | tags=27% |
| hsa00640 | Propanoate metabo | 30 | 0.26668000 | 0.80361588 | 0.76131045 | 0.82270645 | 0.53511464 | 3941 | tags=20% |
| hsa00564 | Glycerophospholip | 93 | 0.22475784 | 0.83117145 | 0.77006802 | 0.82949449 | 0.53952980 | 2903 | tags=19% |

| | | | | | | | | | |
|---|---|---|---|---|---|---|---|---|---|
| hsa04666 | Fc gamma R-medi | 95 | 0.22724186 | 0.83879301 | 0.77975376 | 0.83489784 | 0.54304431 | 3860 | tags=25% |
| hsa05223 | Non-small cell lun | 72 | -0.2094207 | -0.8492681 | 0.78006872 | 0.83489784 | 0.54304431 | 2033 | tags=22% |
| hsa00604 | Glycosphingolipid | 15 | 0.29838785 | 0.75532865 | 0.78260869 | 0.83494876 | 0.54307743 | 3314 | tags=20% |
| hsa00270 | Cysteine and meth | 47 | -0.2196766 | -0.8201270 | 0.79027355 | 0.84044965 | 0.54665539 | 5063 | tags=38% |
| hsa00983 | Drug metabolism - | 62 | 0.23389975 | 0.80552736 | 0.79827089 | 0.84626819 | 0.55043995 | 2535 | tags=19% |
| hsa04714 | Thermogenesis | 208 | 0.20253145 | 0.82812051 | 0.83065512 | 0.87782166 | 0.57096334 | 2173 | tags=12% |
| hsa00450 | Selenocompound r | 17 | -0.2480196 | -0.7207840 | 0.84444444 | 0.88958770 | 0.57861635 | 2142 | tags=24% |
| hsa00910 | Nitrogen metabolis | 14 | 0.27721655 | 0.69304720 | 0.86643233 | 0.90988975 | 0.59182146 | 4411 | tags=29% |
| hsa00860 | Porphyrin metabol | 30 | 0.23647661 | 0.71260073 | 0.87363494 | 0.91458658 | 0.59487642 | 5437 | tags=43% |
| hsa04144 | Endocytosis | 244 | -0.1764765 | -0.8683607 | 0.90116279 | 0.94046584 | 0.61170912 | 3513 | tags=22% |
| hsa00600 | Sphingolipid metal | 48 | 0.21330135 | 0.70617115 | 0.90638930 | 0.94298265 | 0.61334614 | 5803 | tags=42% |
| hsa01040 | Biosynthesis of un | 26 | -0.2197928 | -0.7093619 | 0.90958904 | 0.94338182 | 0.61360577 | 4309 | tags=31% |
| hsa05216 | Thyroid cancer | 37 | -0.2040130 | -0.7211835 | 0.92982456 | 0.95697359 | 0.62244629 | 3092 | tags=43% |
| hsa04130 | SNARE interaction | 33 | 0.21964236 | 0.67533683 | 0.93034055 | 0.95697359 | 0.62244629 | 1945 | tags=12% |
| hsa00130 | Ubiquinone and ot | 10 | -0.2336654 | -0.5778799 | 0.93126385 | 0.95697359 | 0.62244629 | 4617 | tags=30% |
| hsa04146 | Peroxisome | 80 | -0.1813645 | -0.7480958 | 0.95340501 | 0.97672991 | 0.63529643 | 4436 | tags=32% |
| hsa04330 | Notch signaling pa | 56 | 0.19352635 | 0.65711087 | 0.95754026 | 0.97797557 | 0.63610665 | 4383 | tags=23% |
| hsa00500 | Starch and sucrose | 31 | 0.19570788 | 0.59299812 | 0.96557120 | 0.98318040 | 0.63949204 | 5764 | tags=35% |
| hsa00561 | Glycerolipid metal | 56 | 0.18123373 | 0.61537178 | 0.97803806 | 0.99285682 | 0.64578590 | 2021 | tags=12% |
| hsa00603 | Glycosphingolipid | 14 | 0.19989354 | 0.49973805 | 0.98769771 | 0.99903598 | 0.64980502 | 4704 | tags=29% |
| hsa04142 | Lysosome | 127 | 0.16496348 | 0.63464416 | 0.99082568 | 0.99903598 | 0.64980502 | 3967 | tags=17% |
| hsa00120 | Primary bile acid b | 14 | -0.1814863 | -0.4987552 | 0.99307159 | 0.99903598 | 0.64980502 | 15710 | tags=100% |
| hsa00790 | Folate biosynthesis | 25 | -0.1577634 | -0.5042039 | 0.99730458 | 1 | 0.65043205 | 7292 | tags=48% |
| hsa00040 | Pentose and glucur | 24 | -0.1368292 | -0.4294679 | 1 | 1 | 0.65043205 | 6889 | tags=42% |

| core_enrichme | sig. | genes |
|---|---|---|
| list=12% | signal=29% | MYH11/ACTG2/MYL9/MYLK/RAMP1/KCNMB1/CALD1/PPP1R12B/ACTA2/PPP1R14A/ADCY5/KCN |
| list=23% | signal=41% | CDC14B/HDAC2/ORC2/STAG2/E2F5/GSK3B/ATR/YWHAE/E2F4/ANAPC13/ANAPC4/ZBTB17/ORC1/ |
| list=14% | signal=29% | MYL9/PLN/MYLK/ATP1A2/ATP2B4/KCNMB1/ADCY5/PDE5A/KCNMA1/IRAG1/RGS2/NPR1/CACNA |
| list=11% | signal=26% | MYL9/FLNC/MYLK/FLNA/PPP1R12B/ITGA7/THBS4/FN1/CAV1/IGF1/COL6A1/AKT3/MAPK10/JUN/ |
| list=18% | signal=37% | JAM3/CADM3/NCAM1/CLDN11/CD8B2/NRXN1/CD34/SELE/NEGR1/ITGA9/JAM2/HLA-DOA/NRXN |
| list=9% | signal=26% | DES/PLN/TPM2/ITGA7/SGCD/ADCY5/SGCA/DMD/TPM1/IGF1/CACNA1C/ACTC1/ITGA9/ADRB1/M |
| list=14% | signal=25% | PLN/TPM2/PPP1R1A/ATP1A2/ATP2B4/SCN7A/ADCY5/TPM1/CACNA1C/PPP2R2B/AKT3/ACTC1/AG |
| list=14% | signal=36% | FABP4/ADCY5/PTGER3/CIDEC/PLAAT3/AQP7/NPR1/PLIN1/AKT3/PRKG1/LIPE/PIK3CD/ADRB1/PT |
| list=28% | signal=40% | KPNA7/IPO7/KPNA6/THOC3/NUP88/SAP18/NUP93/TNPO1/IPO9/KPNA3/NCBP1/UPF3A/XPO1/RAE1 |
| list=15% | signal=27% | MYL9/MYLK/PPP1R12B/ADCY5/GNAO1/RGS2/NPR1/CACNA1C/RYR3/FOS/JUN/PPP1R12A/GUCY1 |
| list=29% | signal=51% | PSMB9/PSMB8/PSMA3/PSMD8/PSMD11/PSMD13/PSME1/PSMB5/SEM1/PSMD9/PSMD4/PSMB3/PSM |
| list=17% | signal=29% | PLN/MYLK/CASQ2/ATP2B4/PTGER3/FGFR1/P2RX1/GNAL/CHRM2/NTRK3/CACNA1C/NOS1/TACR |
| list=11% | signal=62% | RPA2/POLA2/MCM6/RNASEH1/RFC2/PRIM2/POLD4/PRIM1/RFC5/RPA3/FEN1/POLD3/MCM5/RFC4/ |
| list=24% | signal=44% | IARS1/DARS2/CARS1/PARS2/SARS2/EPRS1/FARSB/YARS1/QARS1/EARS2/QRSL1/NARS1/GARS1/ |
| list=21% | signal=26% | ATG13/NDUFB11/PSMD9/ACTR1B/NRBF2/NDUFAB1/ATP5F1B/COX6B1/SRSF7/DCTN5/KLC3/PSM |
| list=21% | signal=30% | MYL9/PLN/ATP1A2/ATP2B4/FXYD1/ADCY5/PTGER3/GLI3/CHRM2/NPR1/CACNA1C/AKT3/MAPK1 |
| list=21% | signal=30% | ANK2/FLNC/FLNA/PPP1R12B/DCN/GPC3/FN1/CAV1/ESR1/TWIST2/FGFR1/IGF1/HOXD10/HSPB2/A |
| list=15% | signal=32% | MAPK10/FOS/JUN/HLA-DOA/HLA-DOB/HLA-DRA/NFATC1/CD3E/HLA-DQB1/JAK2/HLA-DMB/NF |
| list=14% | signal=28% | ADCY5/GNAO1/GNAL/CACNA1C/GNG7/PPP2R2B/AKT3/MAPK10/FOS/MAOB/GNB4/ITPR1/PRKC |
| list=12% | signal=27% | DES/TPM2/ITGA7/SGCD/SGCA/DMD/TPM1/IGF1/CACNA1C/ACTC1/ITGA9/PRKAA2/MYL3/SLC8A |
| list=25% | signal=30% | CDC5L/SF3B6/ISY1/MAGOH/SF3A2/TRA2A/MAGOHB/LSM5/DHX15/SNRNP40/SF3B2/PRPF6/SNRN |
| list=27% | signal=42% | RPA2/TOP3A/FAAP24/FANCF/USP1/TOP3B/CENPS-CORT/CENPS/RMI1/ATR/EME1/ERCC4/FANCE/ |
| list=15% | signal=35% | ADCY5/GNAO1/PER3/PER1/CACNA1C/NOS1/GNG7/RYR3/FOS/GUCY1A1/GNB4/ITPR1/PRKCB/CA |
| list=14% | signal=27% | C3/THBS4/COLEC12/TUBB2B/ATP6V0C/NOS1/TUBA1A/C1R/CD36/DYNC1I1/HLA-DOA/THBS2/HL |
| list=19% | signal=35% | WDR43/RIOK1/NOP58/UTP6/FBL/DKC1/IMP3/RPP30/UTP4/RRP7A/GTPBP4/MDN1/RAN/BMS1/NXF |
| list=14% | signal=51% | MS4A2/HLA-DOA/HLA-DOB/HLA-DRA/HLA-DQB1/HLA-DMB/HLA-DPA1/HLA-DRB5/HLA-DMA/ |
| list=12% | signal=25% | MYLK/ACTA2/ADCY5/NOS1/GNG7/EGR1/PLIN1/RYR3/AKT3/CCN2/AGTR1/GNB4/ITPR1/PRKAA2/ |
| list=13% | signal=30% | C3/CFD/MASP1/C1R/HLA-DOA/HLA-DOB/HLA-DRA/CFH/C1S/C1QA/HLA-DQB1/C5/HLA-DMB/HL |
| list=21% | signal=37% | GALK2/PGM1/FCSK/NANS/NANP/HKDC1/GALE/TGDS/GMPPB/HK2/GPI/GNPNAT1/PMM2/GFPT1/ |
| list=12% | signal=22% | SFRP1/SFRP4/RSPO2/RSPO3/SFRP2/SERPINF1/APCDD1L/MAPK10/JUN/ROR1/PRKCB/ROR2/NFAT |
| list=22% | signal=35% | ATP1A2/ATP2B4/ADCY5/NPR1/CACNA1C/NR4A1/PDE2A/AGTR1/ITPR1/PRKCB/CACNA1H/ATP1B |
| list=17% | signal=26% | MYL9/SLIT3/CXCL12/BOC/RGMA/CFL2/TRPC1/ENAH/NTN1/SLIT2/EPHA3/EPHA7/SEMA3G/SEMA |
| list=20% | signal=31% | PC/IDH2/RPE/GOT2/ABHD14A-ACY1/ENO3/PGK1/TALDO1/TPI1/SDS/PFKP/NAGS/ACY1/PSPH/SH |
| list=19% | signal=25% | FLNC/FLNA/MAPT/DUSP1/FGFR1/IGF1/CACNA1C/NR4A1/AKT3/MAPK10/FOS/JUN/RASGRP2/ME |
| list=14% | signal=28% | ADCY5/GNAO1/CHRM2/CACNA1C/GNG7/AKT3/FOS/GNB4/ITPR1/PRKCB/CACNA1A/PIK3CD/GN |
| list=13% | signal=33% | SGCD/SGCA/CAV1/DMD/HLA-DOA/HLA-DOB/HLA-DRA/ABL1/HLA-DQB1/HLA-DMB/LAMA2/HL |
| list=12% | signal=28% | ATP1A2/ATP2B4/ADCY5/KCNMA1/NOS1/RYR3/GUCY1A1/ITPR1/PRKCB/ATP1B2/PRKG1/ADRB1/ |
| list=24% | signal=32% | IL17A/DEFB4A/TBK1/CHUK/MMP9/HSP90B1/GSK3B/HSP90AB1/PTGS2/IL17RB/TNF/IL17B/IKBKE |

| | | |
|---|---|---|
| list=13% | signal=19% | MYH11/MYL9/MYLK/CXCL12/PPP1R12B/ITGA7/NCKAP1L/FN1/CFL2/FGFR1/CHRM2/ENAH/PPP1 |
| list=20% | signal=35% | ATR/CD82/PMAIP1/APAF1/BCL2L1/FAS/RCHY1/TP53I3/PERP/TP53/BID/AIFM2/IGFBP3/GTSE1/RR |
| list=18% | signal=50% | FPGS/SLC46A1/TNF/ABCC4/TYMS/GGH/FOLR1/ATIC/SLC19A1/ABCC5/SHMT1/ABCC1/GART/IKB |
| list=16% | signal=52% | PFKFB3/TKFC/ALDOA/PFKFB2/FCSK/KHK/HKDC1/SORD/TPI1/PFKP/ENOSF1/PFKFB4/TIGAR/GM |
| list=13% | signal=31% | ATP1A2/ATP2B4/ESR1/PTH1R/PRKCB/ATP1B2/SLC8A2/ATP1B3/FXYD2/PRKACA/SLC8A1/DNM1/G |
| list=30% | signal=45% | RAD51B/BABAM1/MRE11/RAD54B/RPA2/TOP3A/SYCP3/XRCC3/RBBP8/SEM1/TOP3B/POLD4/EME |
| list=15% | signal=35% | ADCY5/KCNMA1/AQP1/NPR1/CACNA1C/AGTR1/GUCY1A1/ITPR1/PDE1A/ADCYAP1R1/PDE3A/A |
| list=12% | signal=40% | THBS4/ACKR1/GYPC/SELE/HBA2/CD36/HBB/CCL2/THBS2/HBA1/THBS1/KLRB1/LRP1/COMP/TGF |
| list=17% | signal=27% | APAF1/EGFR/TNF/YWHAZ/FAS/NRAS/IKBKE/CLDN23/EIF2AK3/OCLN/TP53/CLDN15/BID/PPP2R1 |
| list=8% | signal=24% | ATP1A2/COL14A1/COL6A1/DPP4/COL8A2/CPB1/ATP1B2/COL6A3/COL1A2/SLC8A2/COL6A2/COL2 |
| list=9% | signal=21% | TPM2/CASQ2/ATP1A2/TPM1/CACNA1C/ACTC1/COX7A1/ATP1B2/MYL3/SLC8A2/CACNA2D1/ATP1 |
| list=32% | signal=29% | RGS2/GNAL/GNG7/PDE2A/PDE1A/PRKG1/NCALD/SLC8A2/PDE1B/PRKACA/SLC8A1/CAMK2A/O |
| list=17% | signal=36% | ITGA7/THBS4/FN1/COL6A1/CD36/ITGA9/TNXB/THBS2/LAMA4/THBS1/COL6A3/COL1A2/COL6A2/ |
| list=24% | signal=29% | UBQLN1/EIF2AK1/UBE2J2/SEC24C/EDEM1/CAPN1/RAD23A/EIF2S1/RRBP1/HSP90B1/LMAN1/BAK |
| list=13% | signal=28% | CXCL12/TNFSF13B/HLA-DOA/HLA-DOB/HLA-DRA/HLA-DQB1/CXCR4/HLA-DMB/HLA-DPA1/HL |
| list=14% | signal=24% | ACTA2/ADCY5/GNAO1/NOS1/GNG7/AKT3/MAPK10/FOS/JUN/GNB4/MMP2/PIK3CD/COL1A2/GNG |
| list=21% | signal=31% | MAPK10/FOS/JUN/HLA-DOA/HLA-DOB/HLA-DRA/NFATC1/IL6R/CD3E/HLA-DQB1/JAK2/HLA-DM |
| list=20% | signal=32% | FABP4/SORBS1/PLIN4/ADIPOQ/AQP7/PLIN1/CD36/LPL/ILK/CPT1B/SLC27A6/SCD5/CPT1C/ME3/A |
| list=20% | signal=26% | PC/IDH2/ME1/RPE/GOT2/MDH1/ENO3/OGDH/PGK1/HKDC1/MCEE/TALDO1/TPI1/SDS/PFKP/SUCL |
| list=12% | signal=24% | ADCY5/DUSP1/GNAO1/TRPC1/CACNA1C/GNG7/MAOB/GNB4/ITPR1/PRKCB/CACNA1A/PLA2G4C |
| list=14% | signal=30% | C3/FOS/JUN/PRKCB/HLA-DOA/HLA-DOB/HLA-DRA/CYBB/HLA-DQB1/EEF1A1/JAK2/HLA-DMB/T |
| list=11% | signal=28% | RB1/CDK6/ITGA3/ITGA2B/CYCS/CDK4/PIK3R3/LAMA3/CASP3/CCND1/IKBKG/LAMC2/TRAF4/MY |
| list=18% | signal=24% | ITGA7/THBS4/FN1/GHR/FGFR1/IGF1/COL6A1/CHRM2/NR4A1/GNG7/PPP2R2B/AKT3/ITGA9/GNB4/ |
| list=16% | signal=20% | C3/PTGER3/CTSG/LEPR/PTH1R/GHR/P2RX1/NR3C1/CHRM2/TACR2/LYNX1/AGTR1/P2RX7/DRD4/ |
| list=20% | signal=39% | NTHL1/MPG/POLD4/LIG3/NEIL3/PARP4/UNG/FEN1/POLD3/APEX2/POLD2/SMUG1/POLE/MUTYH/ |
| list=20% | signal=38% | NT5C3B/ENTPD5/AK7/HPRT1/NT5C2/NTPCR/IMPDH2/UPP1/NME1-NME2/CTPS2/ENPP3/NME6/CM |
| list=14% | signal=30% | TRPC1/CACNA1C/AKT3/ITPR1/KCNN3/PRKCB/CACNA1H/PIK3CD/GPER1/GABBR1/KCNJ3/TRPC4 |
| list=21% | signal=26% | IRS2/PPID/ATG13/NDUFB11/GSK3B/PSMD9/NRBF2/NDUFAB1/ATP5F1B/COX6B1/KLC3/PSMD4/PS |
| list=15% | signal=49% | PDSS1/DHDDS/FDPS/NUS1/IDI1/HMGCS1/HMGCS2/FNTB/RCE1/MVD/HMGCR/ZMPSTE24 |
| list=21% | signal=25% | ATG13/NDUFB11/PSMD9/ACTR1B/NRBF2/MAP3K10/NDUFAB1/ATP5F1B/CREB3L4/COX6B1/DCT |
| list=17% | signal=32% | FN1/SELE/EGR1/AKT3/MAPK10/JUN/AGTR1/PRKCB/CCL2/NFATC1/MMP2/CYBB/PIK3CD/COL1A |
| list=21% | signal=30% | GALK2/PGM1/FCSK/NANS/NANP/HKDC1/GALE/CYB5R4/CYB5R2/GMPPB/HK2/GPI/GNPNAT1/PM |
| list=17% | signal=25% | TUBB/ACTR2/ARHGEF11/TNF/CASP7/PYCARD/FAS/ABI1/CLDN23/MYH14/ACTR3C/OCLN/MYO1 |
| list=19% | signal=26% | FGFR1/IGF1/PLA2G2C/PLAAT3/GNG7/AKT3/MAPK10/RASGRP2/GNB4/NGFR/PRKCB/RASA3/FGF |
| list=21% | signal=36% | NME1-NME2/CTPS2/ENPP3/NME6/CMPK1/NT5C/DCTD/DUT/ENTPD8/TYMS/DHODH/RRM2/ENTP |
| tags=24% | list=13% | signal=21% ADCY5/PTH1R/FGFR1/EGR1/FOS/ITPR1/MEF2C/PRKCB/JUND/PRKACA/PDE4D/CREB5 |
| list=14% | signal=28% | BOC/CDON/GLI3/SCUBE2/HHAT/GAS1/SPOP/GLI2/CCND2/MEGF8/PRKACA/DHH/GLI1/PTCH2/KI |
| list=14% | signal=26% | MYLK/ADCY5/P2RX1/AKT3/PPP1R12A/GUCY1A1/RASGRP2/ITPR1/PRKG1/PIK3CD/PLA2G4C/CO |
| list=9% | signal=20% | ADCY5/GNAO1/GNG7/PDE2A/GNB4/PRKCB/PDE1A/CACNA1A/PDE3A/PDE1B/GNG2/GABBR1/KC |

| | | |
|---|---|---|
| list=14% | signal=26% | ADIPOQ/SLC2A4/ACACB/LEPR/AKT3/MAPK10/CD36/PRKAA2/LEP/JAK2/PPARGC1A/CPT1B/NPY/ |
| list=14% | signal=41% | TYMS/ATIC/MTHFD2/MTHFD1L/SHMT1/MTHFS/GART/ALDH1L1/SHMT2 |
| list=13% | signal=29% | HLA-DOA/HLA-DOB/HLA-DRA/HLA-DQB1/HLA-DMB/HLA-DPA1/HLA-DRB5/HLA-DMA/CD40LG/ |
| list=14% | signal=25% | ATP1A2/KCNMB1/ADCY5/KCNMA1/CACNA1C/KCNN3/PRKCB/ATP1B2/SNAP25/ADCYAP1R1/ATP |
| list=14% | signal=22% | ADCY5/CACNA1C/EGR1/MAPK10/JUN/ITPR1/PRKCB/MMP2/PLA2G4C/PRKACA/MAP3K3/CAMK2 |
| list=24% | signal=46% | RFC2/EXO1/POLD4/MSH2/RFC5/MSH6/RPA3/POLD3/RFC4/POLD2/SSBP1/PCNA/LIG1/RFC3 |
| list=17% | signal=34% | EGFR/NRAS/CDH1/DAPK2/TP53/SRC/RB1/VEGFA/CDK4/CCND1/ERBB2/MYC/E2F3/E2F2/MMP1/E2 |
| list=26% | signal=36% | POLR2F/POLR1C/POLR1E/POLR2I/POLR2G/POLR2H/POLR2E/POLR2K/POLR2L/POLR1B/POLR1H/ |
| list=11% | signal=18% | ADIPOQ/SLC2A4/ACACB/LEPR/IGF1/PPP2R2B/AKT3/CD36/PRKAA2/CAB39L/LEP/CIDEA/LIPE/PIK |
| list=19% | signal=32% | HLA-DOA/HLA-DOB/HLA-DRA/HLA-DQB1/HLA-DMB/HLA-DPA1/HLA-DRB5/HLA-DMA/CD40LG/ |
| list=19% | signal=55% | GLS/CPS1/GLS2/GOT2/ABHD14A-ACY1/NAGS/ACY1/ARG2/GPT2/ASL/GPT/NOS3/NOS2 |
| list=9% | signal=20% | DES/ITGA7/SGCD/SGCA/DMD/CACNA1C/ITGA9/SLC8A2/CACNA2D1/ITGA5/TCF7L1/SLC8A1/CTN |
| list=16% | signal=22% | C3/CFD/C7/MASP1/MAPK10/FOS/JUN/C1R/AGTR1/PRKCB/CCL2/RPL36/RPL34/IL6R/CYBB/RPL17/ |
| list=15% | signal=32% | CD8B2/CD34/CD36/IL11RA/HLA-DOA/HLA-DOB/HLA-DRA/IL6R/MS4A1/CD3E/CR2/CSF2RA/IL3RA |
| list=15% | signal=26% | GNAO1/GNAZ/IGF1/NOS1/GUCY1A1/ITPR1/PRKCB/CACNA1A/PRKG1/PLA2G4C/RYR1/GUCY1B1/ |
| list=15% | signal=25% | ADCY5/GNAO1/TRPC1/CACNA1C/GNG7/GNB4/ITPR1/PRKCB/CACNA1A/PLA2G4C/GNG2/GRIA4/I |
| list=13% | signal=22% | ATP1A2/ATP2B4/ADCY5/KCNMA1/TRPC1/PLA2G2C/ITPR1/PRKCB/CPB1/ATP1B2/BST1/PLA2G12B |
| list=25% | signal=38% | GALNT1/B4GALT5/GALNTL6/GALNT4/ST3GAL2/ST3GAL1/C1GALT1/GALNT6/B3GNT6/B3GNT3/F |
| list=13% | signal=21% | ADCY5/GNAO1/CACNA1C/GNG7/GNB4/PRKCB/CACNA1A/PLCL1/GNG2/GABBR1/PRKACA/GABA |
| list=13% | signal=23% | C3/CTSG/C7/C1R/H2AC19/HLA-DOA/HLA-DOB/HLA-DRA/C1S/C1QA/HLA-DQB1/H4C5/ACTN1/H2 |
| list=14% | signal=20% | ADCY5/GNAO1/CACNA1C/GNG7/MAPK10/GNB4/ITPR1/PRKCB/CACNA1A/CNR1/GNG2/GRIA4/KC |
| list=18% | signal=24% | PSMD14/APAF1/TNF/PSMD7/PSMD2/PSMC1/FAS/ENTPD8/IKBKE/ADRM1/MAPK14/TP53/CCNA2/P |
| list=20% | signal=26% | ADCY5/GNAO1/FGFR1/IGF1/AKT3/ENAH/RASGRP2/NGFR/PRKCB/FGF10/FGF7/THBS1/MRAS/PIK |
| list=12% | signal=31% | MNX1/PAX4/NKX2-2/HNF1B/BHLHA15/FOXA3/FOXA2/PDX1 |
| list=14% | signal=28% | JUN/HLA-DOA/HLA-DOB/HLA-DRA/NFATC1/HLA-DQB1/HLA-DMB/TGFB3/MAF/HLA-DPA1/GATA |
| list=21% | signal=29% | ADH1B/AOC3/MAOB/AOX1/MAOA/ADH4/ADH5/ADH1C/TPO/GSTZ1/DDC |
| list=13% | signal=23% | CLU/C3/CFD/C7/SERPING1/MASP1/C1R/CFH/PROS1/CR2/C1S/C1QA/A2M/C5/C1QC/TFPI/F8/ITGB2/ |
| list=12% | signal=15% | SORBS1/SLC2A4/ACACB/PPP1R3C/PYGM/AKT3/MAPK10/INPP5A/PRKAA2/PRKAR2B/LIPE/PIK3C |
| tags=24% | list=14% | signal=21% ADCY5/GHR/IGF1/CACNA1C/AKT3/MAPK10/FOS/ITPR1/PRKCB/PIK3CD/JUNB/PRKAC |
| list=8% | signal=17% | CYCS/HSPA6/CASP3/HSPD1/MYD88/CXCL2/CXCL3/CXCL8/CXCL1/IL1B |
| list=21% | signal=53% | DCLRE1C/MRE11/POLM/LIG4/XRCC6/FEN1/PRKDC/XRCC5 |
| list=14% | signal=23% | MYLK/ATP1A2/ADCY5/ITPR1/PRKCB/ATP1B2/ATP1B3/PRKACA/CAMK2A/SSTR2/GNAQ/PLCB2/K |
| list=22% | signal=26% | CD8B2/AKT3/MAPK10/FOS/JUN/NFATC1/GRAP2/PIK3CD/CD3E/CD8A/PIK3R1/NFATC3/PPP3CB/MA |
| list=14% | signal=25% | DPM3/ALG11/MOGS/DOLPP1/DAD1/MAN2A1/RPN2/ALG1/ALG5/B4GALT3/ALG2/DPM1/ALG3/DPM |
| list=16% | signal=20% | YWHAZ/PSMC1/NRAS/GTF2H3/ATF6B/ATP6V0D2/ACTN4/TP53/CCNA2/SRC/KAT2A/DNAJA3/RB1/ |
| list=15% | signal=22% | SLC2A4/ACACB/PPP1R3C/PYGM/AKT3/MAPK10/CD36/PRKAA2/PRKCB/PIK3CD/PPP1CB/CREB5/F |
| list=15% | signal=26% | C3/GNAO1/CCL5/GNAL/PPP2R2B/AKT3/MAPK10/FOS/JUN/CCL2/PIK3CD/CD3E/C1QA/PIK3R1/TGI |
| list=30% | signal=44% | HSD17B7/FDFT1/CEL/CYP2R1/LBR/SQLE/CYP51A1/CYP27B1/NSDHL/DHCR24/MSMO1/DHCR7 |
| list=14% | signal=28% | ADCY5/CACNA1C/FOS/JUN/MAOB/PRKCB/GRIA4/SLC18A2/PRKACA/PPP1CB/CAMK2A/CREB5/P |

| | | |
|---|---|---|
| list=17% | signal=23% | APAF1/VAV3/TNF/MAP3K5/RAP1B/BCL2L1/CASP7/PYCARD/FAS/CALML4/NRAS/IKBKE/PLCB3/E |
| list=14% | signal=18% | ADIPOQ/ADCY5/IGF1/AKT3/PRKAA2/PIK3CD/PRKACA/CREB5/PIK3R1/PPARGC1A/FOXO1/APPL1 |
| list=8% | signal=24% | ATP1A2/IGF1/PRKCB/ATP1B2/PIK3CD/ATP1B3/FXYD2/NR3C2/PIK3R1 |
| list=21% | signal=24% | PPID/ATG13/NDUFB11/GSK3B/PSMD9/ACTR1B/BAK1/GPR37/NRBF2/MAP3K10/NDUFAB1/ATP5F1 |
| list=18% | signal=22% | CXCL12/AR/ADCY5/PTGER3/FN1/MITF/ZBTB16/ESR1/RUNX1T1/FGFR1/IGF1/GLI3/GNG7/GSTM2/ |
| list=6% | signal=17% | LYN/TRADD/PLCG1/BCL10/IKBKG/CCL13/SYK/IRAK1/TRIM25/UBE2I/MYD88/PLAU/TRAF2/CXCL |
| list=17% | signal=21% | MEIS1/MITF/ZBTB16/PBX3/RUNX1T1/IGF1/ZEB1/CDK14/SIX4/PBX1/NR4A3/MEF2C/NGFR/LMO2/I |
| list=13% | signal=23% | MYL9/CXCL12/JAM3/CLDN11/PRKCB/JAM2/MMP2/CYBB/PIK3CD/CLDN5/CXCR4/ACTN1/CTNNA |
| list=29% | signal=33% | PELO/DAZAP1/SAP18/PPP2R5A/PPP1CA/PPP2R5B/CPSF2/NCBP1/UPF3A/FIP1L1/CPSF3/CSTF3/MA( |
| list=14% | signal=25% | GNAO1/AKT3/MAPK10/HLA-DOA/HLA-DOB/HLA-DRA/HSPA1A/LAMA4/HLA-DQB1/CCR5/LAMB |
| list=21% | signal=24% | HAND1/MEIS1/LIFR/FGFR1/IGF1/AKT3/ID4/SOX2/PIK3CD/WNT9A/FGFR2/FZD3/BMPR1B/FZD7/JA |
| list=16% | signal=22% | ANAPC10/YWHAZ/CALML4/ADCY1/SGO1/MAPK14/SPDYA/PPP2R1A/BUB1/MAPK13/MAD2L1/YW |
| list=22% | signal=29% | AKT3/FOS/JUN/PRKCB/NFATC1/PIK3CD/CR2/BTK/PIK3R1/NFATC3/PPP3CB/CD19/CD22/PPP3CC/R |
| list=15% | signal=21% | AKT3/FOS/JUN/NFATC1/PIK3CD/CD3E/JAK2/MAP3K3/PIK3R1/NFATC3/PPP3CB/MAPK11/IFNGR2/N |
| list=21% | signal=26% | AOC3/DPYD/CNDP1/CARNS1/UPB1/CSAD/ALDH1B1/ACOX3/ALDH6A1 |
| list=16% | signal=22% | LIFR/FHL1/LEPR/GHR/AKT3/IL11RA/CTF1/LEP/IL6R/PIK3CD/CSF2RA/IL3RA/CCND2/PIM1/AOX1/. |
| list=21% | signal=27% | ATG13/PSMD9/NRBF2/PSMD4/SPTBN2/PSMB3/PSMA1/PSMD14/SLC25A5/PSMA2/MAP3K5/PSMD7 |
| list=14% | signal=25% | ATP1A2/ADCY5/AQP1/ATP1B2/NR1H4/ABCB1/ATP1B3/FXYD2/PRKACA/EPHX1/ABCG2/ABCC2/U( |
| list=20% | signal=25% | CTPS2/GGCX/GCH1/EPRS1/PHOSPHO2/FPGS/NME6/CMPK1/EARS2/NADK/GGH/GSS/DHODH/NFS |
| list=24% | signal=36% | FUCA1/GLB1/NEU1/FUCA2/NEU4/ENGASE/NEU3 |
| list=15% | signal=20% | NTRK3/AKT3/MAPK10/JUN/NGFR/PIK3CD/ABL1/MAP3K3/CAMK2A/PIK3R1/RPS6KA5/MAPK11/R |
| list=11% | signal=34% | FUT6/B3GALT1/B3GNT2/B3GNT3/ST3GAL4/B4GALT3/FUT2/FUT3/FUT4/ABO |
| list=17% | signal=26% | ADCY5/CACNA1C/NR4A1/AGTR1/PBX1/ITPR1/CACNA1H/KCNK3/PRKACA/CREB5/GNAQ/PLCB2/ |
| list=23% | signal=37% | WWTR1/WTIP/FAT4/TEAD3/DCHS1/DCHS2/FRMD6/LATS2/SAV1/MOB1B/LIMD1/TEAD2/RASSF2/R |
| list=14% | signal=20% | C3/ATP6V0C/AKT3/MAPK10/LSP1/HLA-DOA/HLA-DOB/HLA-DRA/HLA-DQB1/JAK2/TLR1/MRC2/C |
| list=20% | signal=29% | ADH1B/ECI2/CPT1B/ADH4/CPT1C/ACADSB/ADH5/CYP2U1/HADHB/ADH1C/ACADM/CPT1A/ALDI |
| list=8% | signal=16% | ADCY5/CRYAB/IGF1/AKT3/PRKAA2/HSPA1A/PIK3CD/PRKACA/PIK3R1/FOXO1 |
| list=17% | signal=25% | DUSP1/CAV1/SELE/GSTM2/AKT3/MAPK10/FOS/JUN/GSTM5/PRKAA2/MEF2C/CCL2/KLF2/MMP2/P |
| list=15% | signal=24% | ADCY5/TUBB2B/TUBA1A/GUCY1A1/ITPR1/PRKCB/PRKG1/ADRB1/TUBB6/PRKACA/HTR2A/PDGI |
| list=6% | signal=22% | ATP1A2/AQP1/ATP1B2/ATP1B3/FXYD2 |
| list=19% | signal=21% | AR/ADCY5/ESR1/CACNA1C/GSTM2/AKT3/FOS/JUN/GSTM5/PRKCB/PGR/FGF10/FGF7/CACNA1A/I |
| list=24% | signal=31% | PPP1R1A/CACNA1C/ITPR1/PRKCB/PRKACA/PPP1CB/CAMK2A/PPP3CB/RAP1A/PPP3CC/GNAQ/PL |
| list=15% | signal=22% | PTGIS/PLA2G2C/PLAAT3/CBR3/PTGDS/PLA2G4C/PLA2G12B/PTGS1/PLB1/PLA2G2D/CYP2E1/EPH |
| list=19% | signal=25% | SEC61B/SEC61A1/SRPRB/SPCS1/SRPRA/SRP9/HSPA5 |
| list=14% | signal=23% | ADIPOQ/SLC2A4/CACNA1C/MAPK10/CACNA1A/PIK3CD/PIK3R1/PRKCE/SOCS3/PKLR/KCNJ11/CA |
| list=25% | signal=35% | ADH1B/GSTM2/GSTM5/CBR3/AKR1C1/CYP1B1/EPHX1/GSTT2B/UGT1A10/ADH4/MGST3/GSTM4/( |
| list=27% | signal=38% | FH/IDH1/IDH3B/IDH3G/PDHA1/SUCLA2/PC/IDH2/MDH1/OGDH/SUCLG1/DLAT/SDHB/MDH2/ACLY |
| list=18% | signal=27% | PER3/NR1D2/PER1/PRKAA2/BHLHE41/CRY2/RORB/BHLHE40/NFIL3/NR1D1/DBP |
| list=14% | signal=23% | ATP1A2/AKT3/PRKCB/ATP1B2/PIK3CD/ATP1B3/FXYD2/PIK3R1/PLCB2/PLCB1/AKT1 |

| | | |
|---|---|---|
| list=19% | signal=20% | FBXO32/SLC2A4/IGF1/AKT3/MAPK10/PRKAA2/KLF2/PIK3CD/FOXO6/GADD45B/CCND2/PIK3R1/T |
| list=20% | signal=26% | NDUFAB1/ATP5F1B/COX6B1/ADORA2A/KLC3/PSMD4/PSMB3/SLC39A8/ATP5F1E/NDUFS3/PSMA1 |
| list=18% | signal=23% | ACACB/PYGM/AKT3/ITPR1/PRKAA2/PRKACA/CAMK2A/CREB5/PPARGC1A/PPP3CB/FOXO1/CPT |
| list=14% | signal=18% | PLN/SLC2A4/PDK4/AKT3/MAPK10/AGTR1/CD36/ATP5F1D/COX7A1/PRKCB/SLC25A4/MMP2/CYBI |
| tags=24% | list=7% | signal=22% ALDH4A1/PPAT/GPT2/CAD/ASL/GPT/GFPT1/ASRGL1 |
| list=13% | signal=18% | CACNA1D/MAPK14/TP53/CCNA2/TRPV4/VDAC1/CDC25A/IGFBP3/RB1/MAPK13/CDK6/HUS1/CCN |
| list=14% | signal=22% | AKT3/MAPK10/JUN/ITPR1/LSP1/NFATC1/NFATC4/MRAS/PIK3CD/PIK3R1/NFATC3/EGR2/PPP3CB/M |
| list=16% | signal=26% | ADH1B/FMO2/GSTM2/MAOB/GSTM5/AOX1/MAOA/GSTT2B/UGT1A10/ADH4/MGST3/GSTM4/CYP |
| list=14% | signal=24% | ADCY5/IGF1/MAPK10/ITPR1/PRKCB/PIK3CD/PLA2G4C/PRKACA/PPP1CB/HTR2A/CAMK2A/PIK3R |
| list=28% | signal=34% | IL18/CCL4/RIPK1/POLR2F/POLR1C/IRF7/TBK1/CHUK/POLR2H/POLR2E/ZBP1/PYCARD/IKBKE/CC |
| list=9% | signal=20% | AOC3/MAOB/MAOA |
| list=24% | signal=30% | APOA4/CD36/LPL/APOC1/APOB/LRP1/APOE/NPC2/APOC3/LIPA/STAR/CYP27A1/LIPC/LCAT/PLTP/ |
| list=24% | signal=26% | ADCY5/GNAO1/MITF/PRKCB/WNT9A/FZD3/TCF7L1/PRKACA/FZD7/CAMK2A/FZD4/KIT/GNAQ/PI |
| list=26% | signal=34% | POMT2/GALNT1/GALNTL6/GALNT4/COLGALT1/POGLUT1/ST3GAL3/OGT/B3GLCT/C1GALT1/XX |
| list=30% | signal=43% | IDH1/AADAT/IDH3B/IDH3G/IDH2/GOT2/ABHD14A-ACY1/NAGS/ACY1/GPT2/GPT |
| list=14% | signal=20% | ADCY5/GNAO1/ESR1/AKT3/FOS/JUN/ITPR1/PGR/HSPA1A/MMP2/PIK3CD/GPER1/GABBR1/KCNJ3/ |
| list=13% | signal=28% | HLA-DOA/HLA-DOB/HLA-DRA/HLA-DQB1/KLRD1/HLA-DMB/HLA-DPA1/HLA-DRB5/HLA-DMA/H |
| list=10% | signal=16% | MAPK13/BCL3/TRADD/PIK3R3/MLKL/CASP3/CCL20/IKBKG/PGAM5/PIK3R2/CREB3L1/TRAF2/RIP |
| list=14% | signal=18% | ESR1/AKT3/MAPK10/FOS/PIK3CD/CCND2/JAK2/PIK3R1/MAPK11/MAPK12/SOCS3/RELA/AKT1 |
| list=14% | signal=19% | CXCL12/LIFR/TNFSF12/CCL14/BMP3/LEPR/CXCL13/GHR/CCL5/LTB/TNFSF13B/IL11RA/CX3CL1/N |
| list=14% | signal=20% | CXCL12/ADCY5/CCL14/CXCL13/CCL5/GNG7/AKT3/RASGRP2/GNB4/CX3CL1/PRKCB/CCL2/CCL21 |
| list=20% | signal=33% | MAOB/CNDP1/ASPA/MAOA/CARNS1/HNMT/HAL/ALDH1B1/HDC |
| list=16% | signal=21% | IGF1/AKT3/PRKCB/GAS6/IL6R/PIK3CD/FGFR2/AXL/JAK2/PIK3R1/PDGFC/PDGFD/PTEN/PDGFRB/I |
| list=23% | signal=23% | PLA2G4A/CAPN1/JAK1/H2AX/CHMP4B/STAT6/PPID/TICAM2/PLA2G4D/HSP90AB1/CAPN2/CHMP2 |
| list=13% | signal=20% | FN1/CAV1/SEPTIN6/ARHGEF26/PIK3CD/ITGA5/ILK/CTNNA3/CAV2/PIK3R1/DNM1/VCL/ITGB1/AR |
| list=21% | signal=27% | MITF/AKT3/MAPK10/FOS/JUN/NFATC1/PIK3CD/JUND/BTK/JUNB/PIK3R1/PPP3CB/MAPK11/IFNGR |
| list=8% | signal=18% | IL18RAP/CCL15/CCL3/TNFRSF10D/CCL20/CCL13/CCR1/IL20RA/CCL26/CCL24/CCL3L1/CXCL2/CX |
| list=26% | signal=24% | KLHL9/ANAPC5/UBE2G1/UBE2Q1/UBA3/CUL5/UBE2J2/ANAPC7/ELOB/UBA6/UBE2W/CUL4A/BTI |
| list=16% | signal=23% | MITF/FGFR1/IGF1/AKT3/FGF10/FGF7/PIK3CD/GADD45B/POLK/PIK3R1/PDGFC/PDGFD/PTEN/PDG |
| list=21% | signal=21% | AKT3/MAPK10/ITPR1/PRKAA2/RRAGC/MRAS/PIK3CD/DAPK1/ATG2B/PRKACA/PIK3R1/GABARA |
| list=17% | signal=21% | EGFR/ENO3/PRKCG/EGLN3/HIF1A/PGK1/HKDC1/PFKP/PDK1/VEGFA/HMOX1/PLCG1/PIK3R3/GAI |
| list=17% | signal=25% | ADH1B/HSD17B6/ALDH1A3/ALDH1A1/DHRS4L2/DHRS3/LRAT/AOX1/UGT1A10/ADH4/CYP27C1/A |
| list=12% | signal=16% | MYH11/MYL9/JAM3/AMOTL1/CLDN11/PPP2R2B/MAPK10/JUN/TUBA1A/PRKAA2/JAM2/RDX/TIAI |
| list=20% | signal=22% | PLN/ATP1A2/ESR1/AKT3/KAT2B/PRKCB/ATP1B2/PIK3CD/ATP1B3/FXYD2/PRKACA/PIK3R1/SLC16 |
| list=26% | signal=37% | AK7/NTPCR/ALPI/NFS1/ACP1/TPK1/AK1 |
| list=13% | signal=22% | ATP1A2/ADCY5/GPX3/ITPR1/PRKCB/ATP1B2/ATP1B3/FXYD2/PRKACA/CREB5/TTR/GPX7/GNAQ/I |
| list=15% | signal=19% | ADCY5/EGR1/AKT3/MAPK10/FOS/JUN/KAT2B/HLA-DOA/HLA-DOB/SLC25A4/HLA-DRA/ZFP36/NI |
| list=8% | signal=16% | ATP1A2/ATP2B4/ATP1B2/CYBRD1/SLC8A2/ATP1B3/FXYD2/SLC8A1/TRPM7 |
| list=7% | signal=18% | DAD1/MAN2A1/RPN2/ALG1/B4GALT3/ALG2/ALG3 |

| | | |
|---|---|---|
| list=16% | signal=22% | GNG4/TCF7/LEF1/FAS/CALML4/NRAS/IKBKE/HIF1A/GNG5/GNGT2/MAPK14/TP53/SRC/BID/RB1/R |
| list=13% | signal=42% | GCLC/GCLM/SLC40A1/ATG5/MAP1LC3B2/ACSL6/SLC39A8/GSS/TP53/ACSL3/LPCAT3/HMOX1/SL |
| list=15% | signal=33% | APOA4/SLC23A1/LRAT/APOB/TCN2/PLB1/SLC19A2/RBP2/SLC19A3 |
| list=14% | signal=22% | CD8B2/HLA-DOA/HLA-DOB/HLA-DRA/HSPA1A/HLA-DQB1/CD8A/KLRD1/HLA-DMB/HLA-DPA1/N |
| list=22% | signal=28% | RAD23A/CUL4A/POLD4/GTF2H1/ERCC4/RFC5/GTF2H3/CDK7/RPA3/POLD3/RFC4/POLD2/POLE/PC |
| list=12% | signal=29% | SELE/HBA2/HBB/PRKCB/HBA1/LAMA4/IDO1/GNAQ/PLCB2/APOL1/PLCB1 |
| list=23% | signal=23% | LSM5/EDC3/PAN3/EXOSC1/EXOSC4/PAN2/TOB1/SKIV2L/ENO3/CNOT1/DIS3/TENT4B/PFKP/EXOS |
| list=20% | signal=45% | HS3ST3A1/HS2ST1/B3GALT6/B3GAT3/HS3ST3B1/EXT2/EXTL2/EXT1/NDST2/EXTL3/GLCE/HS3ST1 |
| list=21% | signal=25% | MITF/MAPK10/JUN/CITED2/MRAS/GABARAP/RAB7B/GABARAPL1/PINK1/CALCOCO2/OPTN/ATF |
| list=34% | signal=47% | B4GALT4/FUT8/CHST4/CHST2/ST3GAL2/B3GNT7/ST3GAL1/ST3GAL3/B3GNT2/B4GALT3 |
| list=22% | signal=35% | HPSE/GLB1/NAGLU/HYAL3/GUSB/IDUA/HYAL1/HYAL2 |
| list=26% | signal=28% | INMT/MAOB/AOX1/CYP1B1/MAOA/IDO1/KYNU/AFMID/ALDH1B1/TDO2/DDC/HAAO/IDO2/ALDH |
| list=24% | signal=26% | ITPR1/INPP5A/PRKCB/PIK3CD/PI4K2A/ITPKB/PIK3R1/DGKA/DGKB/MTMR8/PTEN/PLCB2/CALM3 |
| list=7% | signal=20% | CKMT1B/SAT1/ALDH4A1/ARG2/CKMT1A/PYCR1/SRM/NOS3/NOS2/ODC1 |
| list=11% | signal=16% | MAP1B/FEZ1/BST2/MAP1A/KAT2B/SAMHD1/CXCR4/CCR5/APOBEC3C/PSIP1/AFF4 |
| list=9% | signal=14% | ADCY5/ESR1/IGF1/AKT3/MAPK10/FOS/JUN/MMP2/PIK3CD/GPER1/PRKACA/PIK3R1/MAPK11/MA |
| list=15% | signal=27% | PLA2G2C/PLAAT3/PLA2G4C/PLA2G12B/PLB1/PLA2G2D/CYP2E1/PLA2G5 |
| list=14% | signal=25% | CXCL12/CCL5/ATP6V0C/LTB/TNFSF13B/FOS/JUN/HLA-DOA/CCL2/HLA-DOB/HLA-DRA/ATP6V1G |
| list=19% | signal=26% | APOA4/PLA2G2C/CD36/PLPP3/APOB/PLA2G12B/PNLIPRP2/PLA2G2D/PLPP1/PLA2G5/MOGAT2/SL |
| list=22% | signal=34% | PLA2G2C/PLAAT3/PLA2G4C/PLA2G12B/PLB1/PLA2G2D/PLA2G5/FADS2/ACOX3/PLA2G12A |
| list=16% | signal=28% | EIF2AK1/CSNK2A1/IRF7/JAK1/TBK1/EIF2S1/CHUK/CD46/GSK3B/BAK1/IL2RG/APAF1/BCL2L1/FAS |
| list=14% | signal=18% | CXCL12/ADCY5/PTGER3/GNAO1/CCL5/GNG7/AKT3/GNB4/CX3CL1/ITPR1/PRKCB/CCL2/NFATC1/ |
| list=11% | signal=19% | RB1/RAC1/CDK6/VEGFA/CDK4/PIK3R3/CCND1/IKBKG/ERBB2/BRCA2/RAD51/PIK3R2/E2F3/E2F2/ |
| list=19% | signal=22% | ADH1B/ACACB/ADH4/ME3/PKLR/ACSS1/ADH5/ADH1C/PCK1/ACSS2/ALDH1B1 |
| list=14% | signal=19% | PABPN1/CCL5/TPSAB1/AKT3/BCL2L2-PABPN1/PRKCB/HLA-DOA/CCL2/HLA-DOB/SLC25A4/HLA- |
| list=9% | signal=22% | CTSG/AGTR1/PRCP/CPA3/ENPEP |
| list=14% | signal=18% | GNAO1/CFL2/GNG7/AKT3/MAPK10/FOS/BST2/JUN/GNB4/ITPR1/PRKCB/NFATC1/NFATC4/PIK3CD |
| list=24% | signal=24% | INPP5A/PIK3CD/PI4K2A/ITPKB/MTMR8/PTEN/PLCB2/PIK3C3/IPMK/PLCB1/PLCG2/MTMR3/ISYNA |
| list=3% | signal=15% | FMO2/CDO1 |
| list=14% | signal=24% | C3/CFL2/SERPING1/MAPK10/FOS/JUN/C1R/ITGA5/C1S/C1QA/C5/MAPK11/C1QC/MAPK12/ITGB1/C |
| list=19% | signal=18% | ZBTB16/RUNX1T1/AKT3/PIK3CD/TCF7L1/PIM1/PIK3R1/KIT/PML/RELA/AKT1/FLT3/CSF1R/BRAF |
| list=17% | signal=21% | DCN/RGMA/GREM2/ID4/GREM1/THBS1/RGMB/GDF5/BMPR1B/TGFB3/LEFTY2/ID2/FBN1/GDF6/C |
| list=20% | signal=26% | IDH2/MET/GLS2/SLC16A3/EGFR/NRAS/HIF1A/HKDC1/PFKP/TP53/PDK1/PIK3R3/TIGAR/ERBB2/HK |
| list=17% | signal=20% | TNF/YWHAZ/FAS/PRKCG/NRAS/IKBKE/ATF6B/MAPK14/TP53/CCNA2/SRC/BID/RB1/MAPK13/YW |
| list=15% | signal=26% | PRDX6/GSS/GSTP1/GGCT/OPLAH/GPX2/RRM2/CHAC2/TXNDC12/MGST2/PGD/CHAC1/GSTT2/SRM |
| list=16% | signal=21% | MS4A2/AKT3/MAPK10/PIK3CD/PLA2G4C/BTK/PIK3R1/MAPK11/MAPK12/RAC2/FYN/PLCG2/FCER |
| list=36% | signal=37% | EHHADH/ECHS1/ACAT1/OXCT1/ACSM2B/L2HGDH/BDH1/GAD1/HMGCL/HMGCS1/HMGCS2/ACS |
| list=15% | signal=15% | MYL9/FLNC/FLNA/AHNAK2/NCKAP1L/TUBB2B/RHOB/AKT3/MAPK10/FOS/JUN/TUBA1A/DYNC1 |
| list=21% | signal=24% | DYNC1I1/DYNC2H1/DYNC1H1/PRKACA/CREB5/DYNC1I2/DCTN6/DCTN4/RAB5B/ARHGDIB/DYN |

| | | |
|---|---|---|
| list=31% | signal=29% | NDUFS7/COX10/NDUFB9/ATP6V1H/NDUFV2/NDUFA12/UQCRC1/NDUFA2/NDUFA10/NDUFC2/ATI |
| list=9% | signal=19% | ABCA9/ABCA8/ABCA6/ABCC9/ABCB1/ABCA3/ABCA10/ABCG2/ABCC2 |
| list=20% | signal=22% | ADH1B/ADIPOQ/C3/ACACB/AKT3/MAPK10/PRKAA2/C1QA/TCF7L1/C5/PPARGC1A/FOXO1/MAPK |
| list=8% | signal=50% | PIGN/PIGK/PIGP/PIGQ/PIGM/GPAA1/PIGZ/PIGT/PIGW/PIGH/PIGU/PIGG/PIGO/DPM2 |
| list=12% | signal=24% | CTSG/FN1/GNAL/PRKCB/LAMA4/PIK3CD/COL1A2/COL1A1/PRKACA/LAMB2/COL4A6/ACTN1/CD |
| list=14% | signal=29% | GNAS/ARF1/ATP6V1F/SLC12A2/SEC61B/SEC61A1/ATP6V1C2/ATP6V0D2/ATP6V1E2/KCNQ1/PLCG |
| list=21% | signal=24% | PRKCB/HCST/NFATC1/PIK3CD/KLRD1/PIK3R1/PPP3CB/IFNGR2/ITGAL/PPP3CC/RAC2/FYN/SH2D1 |
| list=28% | signal=32% | RGN/UGT1A10/KL/ALDH1B1/UGT1A1/UGT2A3/ALDH7A1/UGT1A7 |
| list=16% | signal=21% | BCL2L1/NRAS/SHC1/TP53/CRKL/RB1/CTBP2/CDK6/CDK4/PIK3R3/CCND1/IKBKG/MYC/HDAC1/PI |
| list=24% | signal=29% | GSTM2/GSTM5/AKR1C2/CYP1B1/EPHX1/GSTT2B/UGT1A10/MGST3/GSTM4/CYP2E1/HSD11B1L/SU |
| list=15% | signal=18% | C3/CTSG/AKT3/H2AC19/PRKCB/SLC25A4/CYBB/PIK3CD/H4C5/HDAC9/H2BC21/PIK3R1/C5/MAPK |
| list=18% | signal=22% | PPP2R2B/MS4A2/AKT3/MAPK10/PRKCB/PIK3CD/S1PR3/PPP2R3A/PIK3R1/MAPK11/PRKCE/MAPK |
| list=22% | signal=26% | ELOVL1/TECR/ACSF3/ACSL6/HACD3/PPT2/ELOVL5/ACACA/ACSL3/SCD/HSD17B4/FASN/ACADV |
| list=18% | signal=21% | MDH1/MCEE/PGP/MDH2/SHMT1/SHMT2/PCCA |
| list=12% | signal=15% | CAV1/NCAM1/PRNP/C7/CCL5/TUBB2B/CACNA1C/EGR1/RYR3/MAPK10/TUBA1A/ATP5F1D/ITPR1/ |
| list=15% | signal=18% | ADCY5/MS4A2/AKT3/AGTR1/MRAS/PIK3CD/PLA2G4C/PLPP3/PIK3R1/DGKA/PDGFC/PDGFD/DNM |
| list=19% | signal=21% | ESR1/FGFR1/IGF1/AKT3/FOS/JUN/PGR/FGF10/FGF7/PIK3CD/WNT9A/GADD45B/FZD3/TCF7L1/FZD |
| list=14% | signal=24% | HLA-DOA/HLA-DOB/HLA-DRA/HLA-DQB1/HLA-DMB/HLA-DPA1/HLA-DRB5/HLA-DMA/HLA-DP |
| list=15% | signal=19% | AKT3/MAPK10/FOS/JUN/PRKCB/PIK3CD/PLA2G4C/PLPP3/WASF3/PIK3R1/DGKA/PDGFC/PDGFD/I |
| list=21% | signal=29% | MGST1/ATP7B/BAK1/MSH2/PMAIP1/APAF1/MAP3K5/BCL2L1/FAS/BIRC2/GSTP1/MSH6/TP53/BRC |
| list=22% | signal=29% | ADCY5/JUN/MAOB/SLC18A2/PRKACA/CREB5/MAOA/FOSB/ATF4/GNAI2/RELA/DLG4/GRIN2C/DI |
| list=18% | signal=22% | TPM1/HOXD10/ZEB1/ZFPM2/TNXB/PRKCB/VIM/THBS1/RDX/PIK3CD/ZEB2/ABL1/ABCB1/ITGA5/ |
| list=7% | signal=11% | ADCY5/IGF1/PLA2G4C/PRKACA/CYP1B1 |
| list=23% | signal=31% | UST/CHST15/CHST12/CSGALNACT1/CHST11/XYLT2/CHST14/CHST3 |
| list=17% | signal=17% | ITGA7/THBS4/FN1/COL6A1/ATP6V0C/PPP2R2B/AKT3/ITGA9/TNXB/THBS2/LAMA4/THBS1/PIK3CI |
| list=21% | signal=19% | IGF1/AKT3/PRKAA2/PRKCB/RRAGC/CAB39L/PIK3CD/WNT9A/FZD3/ATP6V1G2/CLIP1/FZD7/RNF1 |
| list=18% | signal=21% | FZD9/APC2/EGFR/AXIN1/FZD5/TCF7/LEF1/NRAS/TERC/CDH1/SHC1/FZD2/WNT3/LRP5/TP53/RB1/ |
| list=14% | signal=21% | ATP6V1C2/ATP6V0D2/MAPK14/PAK1/SRC/RAC1/ATP6V1E2/MAPK13/LYN/PLCG1/F11R/CASP3/IKE |
| list=24% | signal=23% | NOD1/IRF7/DEFB4A/JAK1/TBK1/MAP1LC3B2/ATG16L1/CHUK/ATG12/MEFV/NLRC4/HSP90AB1/Y |
| list=19% | signal=23% | HSP90AB1/ANAPC13/ANAPC4/GNAI3/ANAPC10/ADCY1/MAPK14/CCNA2/SPDYA/CDC25A/BUB1/ |
| list=21% | signal=27% | GLI3/GLI2/WNT9A/GADD45B/FZD3/TCF7L1/FZD7/POLK/FZD4/GLI1/PTCH2/KIF7/CDKN1A/WNT6/ |
| list=28% | signal=50% | SELENBP1/BPNT2/PAPSS2/TST/SQOR/MPST/ETHE1 |
| list=11% | signal=15% | ATP6V0C/SNAP25/CACNA1A/SLC18A2/ATP6V1G2/SLC1A3/SYT1/SLC6A4/DNM1/ATP6V1B2/SLC17 |
| tags=21% | list=15% | signal=18% HMGCS1/DBT/MCEE/HSD17B10/HMGCS2/HIBADH/AACS/ACAD8/HADH/PCCA |
| list=12% | signal=16% | ATP6V0C/ATP6V1G2/ATP6V1B2/CLCNKB |
| list=14% | signal=16% | ADIPOQ/LEPR/AKT3/MAPK10/FOS/JUN/PRKAA2/COX7A1/LEP/IL6R/PIK3CD/NDUFA5/NDUFS4/PI |
| list=14% | signal=16% | FN1/CD8B2/AKT3/MAPK10/FOS/JUN/CCL2/NFATC1/PIK3CD/ITGA5/CD8A/PIK3R1/NFATC3/MAPK1 |
| list=22% | signal=22% | ADCY5/GNAO1/GNG7/MAOB/GNB4/H2AC19/GNG2/SLC18A2/PRKACA/H4C5/HDAC9/H2BC21/PPP |
| list=21% | signal=22% | ADCY5/CACNA1C/NR4A1/AGTR1/PBX1/ITPR1/CACNA1H/WNT9A/FZD3/KCNK3/TCF7L1/PRKACA |

| | | |
|---|---|---|
| list=24% | signal=28% | ALDOA/G6PD/PGM1/RPE/TALDO1/PFKP/PGD/ALDOC/GPI/RPIA |
| list=25% | signal=28% | BST1/NMNAT2/AOX1/NMNAT3/NNMT/SIRT4/NMRK1/NAPRT/NADK2/NT5M/NUDT12/CD38 |
| list=12% | signal=16% | CD8B2/CD3E/BTK/CD8A/CD19/CD40LG |
| tags=16% | list=14% | signal=14⁄ AOC3/MAOB/SARDH/MAOA/CBS/BHMT |
| list=20% | signal=35% | ACSF3/ACSL6/ACACA/ACSL3/FASN/ACSL4/OXSM |
| list=14% | signal=16% | MYL9/C3/FNBP1/CCL5/AKT3/MAPK10/JUN/SEPTIN6/ITPR1/RRAGC/PIK3CD/FOXO6/ITGA5/ILK/A[illegible] |
| list=16% | signal=17% | RPL36/RPL34/RPL17/RPL36AL/RPL17-C18orf32/RPL26/RPL9/RPS4X/RPS12/RPL22/RPL3/MRPL2/RP[illegible] |
| list=18% | signal=18% | ADCY5/PDE5A/NPR1/PDE2A/GUCY1A1/PDE1A/PDE3A/PDE1B/NME4/ADA2/PDE6B/PDE4D/ADSL/[illegible] |
| list=28% | signal=29% | DHX58/MAP3K7/DDX3X/RIPK1/TKFC/ATG5/IRF7/TBK1/CHUK/ATG12/TANK/TNF/SIKE1/IKBKE/M |
| list=16% | signal=17% | ZNF613/ZNF680/FAS/BIRC2/ZNF616/ZNF649/ZNF587/ZNF155/IKBKE/RBAK/ZNF398/ZNF114/ZNF19 |
| list=14% | signal=15% | CTSF/AKT3/MAPK10/FOS/JUN/TUBA1A/HRK/ITPR1/PIK3CD/CTSO/GADD45B/IL3RA/PIK3R1/NGF/ |
| list=21% | signal=23% | WWTR1/WTIP/PPP2R2B/CCN2/SOX2/GLI2/WNT9A/CCND2/FZD3/GDF5/BMPR1B/DLG5/TCF7L1/TE |
| list=14% | signal=19% | PGK1/HKDC1/TPI1/PFKP/DLAT/ALDH3A1/GAPDH/G6PC3/ALDOC/HK2/GPI/ENO1/PKM |
| list=22% | signal=27% | ELOVL1/TECR/HACD3/PPT2/ELOVL5/THEM4/HADH/ELOVL6/HSD17B12 |
| list=22% | signal=30% | PDE6B/CALM3/CALML3/SLC24A1/CNGB1/RGS9/CALM1 |
| list=23% | signal=22% | HSD17B6/AKR1C1/AKR1C2/CYP1B1/UGT1A10/CYP2E1/HSD11B1L/CYP21A2/HSD17B11/CYP7B1/H |
| list=18% | signal=21% | FZD9/APC2/EGFR/AXIN1/FZD5/TCF7/BCL2L1/LEF1/PRKCG/SMARCD2/NRAS/TERC/SHC1/FZD2/G |
| list=18% | signal=29% | BAK1/PMAIP1/DIABLO/APAF1/BCL2L1/CASP7/BIRC2/BID/CYCS/CASP3/BIRC5 |
| list=15% | signal=18% | IGF1/AKT3/PRKCB/PIK3CD/GADD45B/POLK/CAMK2A/PIK3R1/PTEN/CALM3/PDGFRB/PLCG2/AK |
| list=39% | signal=33% | TAF5/TAF1L/TBPL2/TAF13/TAF15/GTF2A1L/MNAT1/TAF6/TAF7L/GTF2F1/GTF2F2/TBPL1/TAF12/G[illegible] |
| list=5% | signal=9% | GABRE/GRIN2B/GRIN2D |
| list=15% | signal=18% | AR/FGFR1/IGF1/ZEB1/AKT3/PIK3CD/FGFR2/TCF7L1/CREB5/PIK3R1/PDGFC/PDGFD/FOXO1/NKX3 |
| list=12% | signal=17% | CACNA1C/PDE1A/CACNA1A/PDE1B/GABBR1/PRKACA/SCN9A/PLCB2/PLCB1 |
| list=14% | signal=18% | AKT3/PRKCB/PIK3CD/PLA2G4C/PIK3R1/PPP3CB/MAPK11/MAPK12/PPP3CC/RAC2/PLCG2/AKT1 |
| list=14% | signal=15% | CCL5/AKT3/MAPK10/FOS/JUN/PIK3CD/TLR1/PIK3R1/MAPK11/MAPK12/SPP1/TOLLIP/TAB1/RELA |
| list=15% | signal=16% | PLA2G2C/PLAAT3/PLA2G4C/PLPP3/PLA2G12B/PLB1/PLA2G2D/PLPP1/PLA2G5 |
| list=21% | signal=19% | GLB1/PGM1/HKDC1/GALE/PFKP/G6PC3/HK2 |
| list=26% | signal=31% | DPYD/BCAT1/UPB1/CSAD/ALDH1B1/PPCS/PANK2/PANK1 |
| list=16% | signal=14% | AKT3/MAPK10/JUN/PRKCB/PIK3CD/ABL1/CAMK2A/PIK3R1/PLCG2/AKT1/CAMK2G/CDKN1A/CD |
| list=10% | signal=14% | AKT3/PIK3CD/GADD45B/ILK/TCF7L1/POLK/CTNNA3/PIK3R1/PTEN |
| list=21% | signal=29% | POMT2/POMK/ST3GAL3/RXYLT1/FKTN/LARGE2/B4GALT3/FUT4 |
| list=15% | signal=16% | SORBS1/FGFR1/TCF7L1/ACTN1/CTNNA3/WASF3/PTPRM/RAC2/VCL/FYN/WASF1/NECTIN3/TGFB[illegible] |
| list=14% | signal=14% | GSTM2/AKT3/MAPK10/FOS/JUN/ATP5F1D/GSTM5/COX7A1/SLC25A4/PIK3CD/ABL1/AKR1C1/AKR |
| list=10% | signal=13% | AKT3/MAPK10/FOS/JUN/PIK3CD/GADD45B/TCF7L1/POLK/PIK3R1/TGFB3/APPL1/RAC2 |
| list=19% | signal=18% | AKT3/JUN/PRCC/PIK3CD/ARNT2/PIK3R1/TGFB3/RAP1A/AKT1/CDKN1A/FLCN/TFE3/TGFB2/ARN[illegible] |
| list=23% | signal=22% | COLGALT1/SUV39H1/SMYD3/KMT5B/EZH2/NSD2/EHMT2/PLOD2/KMT5A/PLOD1/DOT1L/SUV39H |
| list=21% | signal=21% | ATG2B/GABARAP/GABARAPL1/PIK3C3/ATG10/RPTOR/IGBP1/ATG4D |
| list=21% | signal=16% | ACACB/ACSS3/ACSS1/ACSS2/ACOX3/ALDH6A1 |
| list=15% | signal=17% | PLA2G2C/PLAAT3/PLPP4/PLA2G4C/ETNK2/PLPP3/PLA2G12B/DGKA/DGKB/AGPAT4/PCYT1A/PNP |

| | | |
|---|---|---|
| list=20% | signal=20% | CFL2/AKT3/GSN/PRKCB/PIK3CD/PLA2G4C/PLPP3/AMPH/WASF3/PIK3R1/PRKCE/RAC2/PLCG2/WA |
| list=11% | signal=20% | EGFR/PRKCG/NRAS/TP53/RB1/CDK6/PDPK1/PLCG1/CDK4/PIK3R3/CCND1/ERBB2/PIK3R2/E2F3/E |
| list=17% | signal=17% | ST8SIA1/ST8SIA5/B4GALNT1 |
| list=26% | signal=28% | GCLC/KYAT3/MPST/GCLM/LDHA/APIP/GOT2/MDH1/LACC1/GSS/SDS/AMD1/KYAT1/DNMT3B/EN |
| list=13% | signal=17% | CES1/DPYD/GSTM2/GSTM5/NME4/GSTT2B/UGT1A10/MGST3/GSTM4/CYP2E1/UPB1/UCK1 |
| list=11% | signal=10% | ADCY5/FGFR1/NPR1/PLIN1/ATP5F1D/PRKAA2/COX7A1/PRKG1/LIPE/CNR1/NDUFAF8/NDUFA5/P |
| list=11% | signal=21% | KYAT1/TXNRD3/MARS2/SCLY |
| list=23% | signal=22% | CA14/CA5B/CA7/CA1 |
| list=28% | signal=31% | CP/BLVRA/UGT1A10/UROS/UROD/PPOX/BLVRB/UGT1A1/UGT2A3/HCCS/FECH/MMAB/UGT1A7 |
| list=18% | signal=18% | ZFYVE27/CHMP2B/WWP1/GIT1/ARFGEF2/RAB10/SPG21/EGFR/ACTR2/AGAP5/SNX5/RAB8A/SNX |
| list=30% | signal=29% | PLPP3/CERK/KDSR/CERS1/SGMS1/PLPP1/SPTLC3/DEGS1/ASAH1/B4GALNT1/ASAH2/SPTLC2/PSA |
| list=22% | signal=24% | ELOVL1/TECR/HACD3/ELOVL5/SCD/HSD17B4/ELOVL6/HSD17B12 |
| list=16% | signal=36% | RXRB/RET/MAPK1/PAX8/TFG/TPR/BAK1/TCF7/LEF1/NRAS/CDH1/TP53/PPARG/TPM3/CCND1/MY |
| list=10% | signal=11% | VAMP1/VAMP7/VTI1B/VAMP5 |
| list=24% | signal=23% | VKORC1L1/GGCX/NQO1 |
| list=23% | signal=25% | IDH1/PEX5L/SOD1/HAO2/ACOT8/PXMP4/FAR1/PEX10/GNPAT/SOD2/PRDX1/PEX11B/IDH2/ACSL6/ |
| list=23% | signal=18% | KAT2B/NUMBL/DTX3/DTX1/RBPJ/HEYL/TLE3/NOTCH2/DVL2/ADAM17/TLE1/CIR1/NOTCH3 |
| list=30% | signal=25% | PYGM/AGL/TREH/GBA3/HK1/UGP2/GAA/SI/GYG1/GCK/ENPP1 |
| list=11% | signal=11% | LPL/PLPP4/PLPP3/DGKA/PNLIPRP2/DGKB/AGPAT4 |
| list=25% | signal=22% | ST8SIA1/A4GALT/GBGT1/FUT9 |
| list=21% | signal=14% | CTSG/CTSF/ATP6V0C/AP1S2/CTSO/IDS/NPC2/CTSB/AP4M1/LAPTM5/LAPTM4A/CTSL/LIPA/MCOL |
| list=82% | signal=18% | CYP7B1/CYP46A1/CYP39A1/CH25H/HSD3B7/SCP2/SLC27A5/AKR1D1/ACOX2/AMACR/ACOT8/AK |
| list=38% | signal=30% | ALPL/DHFR/PAH/MOCOS/SPR/AKR1B10/ALPI/GCH1/FPGS/PCBD1/GGH/PTS |
| list=36% | signal=27% | DCXR/UGT2B7/XYLB/AKR1B10/FGGY/AKR1A1/RPE/SORD/GUSB/CRYL1 |

MA1/IRAG1/PLA2G2C/NPR1/CACNA1C/PPP1R12A/AG
'BUB3/ANAPC10/YWHAZ/CDK7/TP53/TTK/CCNA2/RA
\1C/PDE2A/AKT3/PPP1R12A/AGTR1/GUCY1A1/ITPR1/
PPP1R12A/ITGA9/PARVA/TNXB/PRKCB/THBS2/LAMA
2/CNTN1/HLA-DOB/HLA-DRA/CNTN2/CADM1/NRXN
YL3/SLC8A2/CACNA2D1/ITGA5/PRKACA/SLC8A1/ITC
TR1/ATP1B2/ADRB1/MYL3/SLC8A2/CACNA2D1/ATP1
GS1/PRKACA/PIK3R1/NPY/ADRB3/TSHR/GNAI2/AKT
./MAGOH/NUP214/TPR/MAGOHB/NUP210/UPF1/SUMC
.A1/ITPR1/PRKAA2/MEF2C/PRKCB/NFATC1/NFATC4/P
1A1/PSMD14/PSMA2/PSMD7/PSMD2/PSMB6/PSMC1/PS
2/RYR3/AGTR1/ITPR1/P2RX7/PRKCB/CACNA1H/PDE1
'POLD2/POLE/DNA2/SSBP1/RNASEH2A/PCNA/MCM4/
'ARS2/AARS1/HARS2/MARS2/RARS2/TARS2/HARS1/K
)4/NUP188/PSMB3/NUP62/ATP5F1E/NDUFS3/PSMA1/C
0/FOS/JUN/PPP1R12A/ATP1B2/ADCYAP1R1/NFATC1/L
KT3/PPP1R12A/ITPR1/PRKCB/THBS1/RDX/MMP2/MR
ATC3/PPP3CB/MAF/MAPK11/HLA-DPA1/GATA3/IFNGR
3/DRD4/CACNA1A/GNG2/GRIA4/SLC18A2/KCNJ3/PPP
2/CACNA2D1/ITGA5/SLC8A1/ITGA1/TGFB3/LAMA2/T
IP70/PRPF19/HNRNPM/SNRNP27/SRSF7/PRPF38A/SF3
3LM/FANCC/ATRIP/EME2/BRCA1/RPA3/FANCL/FANC
CNA1H/ADCYAP1R1/PRKG1/GNG2/GRIA4/KCNJ3/PRK
A-DOB/HLA-DRA/DYNC2H1/DYNC1H1/THBS1/CYBB/
'1/SNU13/GAR1/NOL6/TBL3/LSG1/POP5/NXT1/GNL3/N
CD40LG/HLA-DPB1/FCER1G/PRG2
MEF2C/KLF2/LIPE/MRAS/MYL3/SLC8A2/GNG2/PRKA
.A-DPA1/C1QC/ITGAL/KRT24/SELP/HLA-DRB5/HLA-D
GMDS/PGM3/GFUS
C1/NFATC4/DAAM1/SOX17/WNT9A/DKK2/CCND2/PRI
2/LIPE/ATP1B3/KCNK3/PRKACA/CAMK2A/CREB5/PRK
\3E/PLXNA4/NFATC4/PIK3CD/ABLIM3/ABL1/SEMA3D
MT1/GAPDH/ARG2/ALDOC/GPT2/PYCR1/ASL/ENO1/G
F2C/NGFR/PRKCB/CACNA1H/FGF10/HSPA1A/FGF7/NF
G2/KCNQ4/KCNJ12/KCNJ3/PRKACA/JAK2/CAMK2A/C
.A-DPA1/ITGAL/RAC2/HLA-DRB5/FYN/HLA-DMA/CD4
3ST1/ATP1B3/FXYD2/PRKACA/GUCY1B1/GNAQ/ADRE
/IL17RE/MAPK14/MUC5B/FOSL1/MAPK13/S100A8/TRA

12A/ITGA9/GSN/FGF10/FGF7/RDX/MRAS/PIK3CD/TI
12/CDK6/SHISA5/CCNB2/CYCS/CDK4/SFN/CASP3/CC
KG/ABCC3/SHMT2/IL1B
PPB/ALDOC/HK2/PMM2/GMDS/GFUS
NAQ/PLCB2/AP2A2/PLCB1/KL
1/BLM/UIMC1/RAD54L/BARD1/BRCA1/RPA3/POLD3/
RB1/PDE1B/PRKACA/PPP3CB/PPP3CC/GUCY1B1/GN
B3/ITGAL/THBS3/SDC2/SELP/CD40LG/ITGB2
A/RNASEL/RB1/CDK6/OAS1/YWHAB/TRADD/CYCS/C
4A1/COL16A1/ELN/ATP1B3/XPNPEP2/COL1A1/FXYD2
B3/FXYD2/SLC8A1/UQCRB/SLC9A6/SLC9A7
8D4/CALM3/CAMK2G/OR2T2/CALML3/PRKG2/CNGB
ITGA5/RELN/COL1A1/LAMB2/COL4A6/COMP/ITGA1/
1/P4HB/HSP90AB1/CAPN2/DNAJA1/SEC61B/DNAJC5/
-DRB5/HLA-DMA/CD40LG/HLA-DPB1
2/VEGFB/COL1A1/PRKACA/COL4A6/VEGFD/CREB5/P
B/NFATC3/PPP3CB/MAPK11/HLA-DPA1/GATA3/IFNGR
OC3/OLR1/PCK1/ACADM/CYP27A1/CPT1A/RXRG/FA
G1/PGP/DLAT/SDHB/MDH2/PSPH/SHMT1/GAPDH/PG
/GNG2/SLC18A2/KCNJ3/PTGS1/PRKACA/HTR2A/SLC
GFB3/MAPK11/HLA-DPA1/IFNGR2/MAPK12/HLA-DR
C/SKP2/PIK3R2/CDK2/E2F3/E2F2/CKS2/CCNE2/CCNE
/PRKAA2/NGFR/TNXB/THBS2/FGF10/FGF7/LAMA4/T
DCYAP1R1/ADRA2A/LEP/ADRB1/CNR1/GABBR1/VI
PCNA/LIG1
PK1/NT5C/DCTD/PNP/LACC1/DUT/ENTPD8/TYMS/RR
/PIK3R1/SPP1/GNAQ/PLCB2/PLCB1/KCNJ11/AKT1/CA
MB3/CAPN2/SLC39A8/ATP5F1E/NAE1/NDUFS3/PTGS2

5/POLR2H/KLC3/TAF4/PSMD4/PSMB3/BDNF/ATP5F1
/VEGFB/COL1A1/PIM1/COL4A6/JAK2/VEGFD/PIK3R1
M2/GFPT1/GMDS/PGM3/GFUS
/MAPK14/ARF6/CASP4/PAK1/SRC/CLDN15/ACTR3/A
0/FGF7/MRAS/PIK3CD/PLA2G4C/ABL1/TIAM1/GNG2/
D6/DCTPP1/UCKL1/CDA/NME1/CANT1/CAD/UMPS/U
/NACA/GATA3/NR4A2/MMP16/GNAQ/PLCB2/MAFB/P
3A/EVC/KIF7
1A2/BTK/COL1A1/PTGS1/PRKACA/PPP1CB/PIK3R1/
NJ3/PRKACA/PDE4D/GNB5/GNG11

| |
|---|
| CPT1C/SOCS3/TNFRSF1A/STK11/RELA/AKT1 |
| /HLA-DPB1 |
| 1B3/FXYD2/PRKACA/CAMK2A/CREB5/GNAQ/PLCB2/ |
| A/MAPK11/MAPK12/MAPK7/GNAQ/PLCB2/CALM3/PI |
| 2F1/CXCL8 |
| POLR1A/POLR1D/POLR3K/POLR1G/POLR2J |
| 3CD/TBC1D1/PPP2R3A/CREB5/PIK3R1/PPARGC1A/FC |
| /TSHR/HLA-DPB1/FASLG/TPO |
| NA3/ITGA1/LAMA2/ITGB5 |
| PIK3CD/C1S/RPL36AL/C1QA/RPL17-C18orf32/RPL26/R |
| A/ITGA5/HLA-DQB1/CD8A/CD1C/ITGA1/HLA-DMB/HI |
| GNAQ/PLCB2/PLCB1/GNAI2/GUCY1A2 |
| KCNJ3/PRKACA/PPP3CB/SLC1A3/GNB5/HOMER2/GN |
| /ATP1B3/FXYD2/CPA3/RAP1A/PNLIPRP2/GNAQ/PLCB |
| OC1B-GALNT4/GALNT2/GALNT5/GALNT3/ST6GALN |
| ARAP/GNB5/GNG11/GABARAPL1/GPHN/GABRG2/GN |
| BC21/C5/HLA-DMB/HLA-DPA1/C1QC/H2AC8/HLA-DR |
| CNJ3/NDUFA5/PRKACA/NDUFS4/MAPK11/GNB5/MAP |
| SMD1/BID/PSMD12/RB1/RAC1/MAPK13/LYN/CDK6/PS |
| 3CD/MAGI2/CNR1/TIAM1/EVL/VEGFB/FGFR2/ANGPT |
| A3/IFNGR2/TBX21/IL12RB1/HLA-DRB5/HLA-DMA/HL |
| /SERPINA5 |
| D/PRKACA/PPP1CB/PIK3R1/PPARGC1A/FOXO1/SOCS |
| A/JAK2/CREB5/PIK3R1/SSTR2/MAPK11/MAPK12/GNA |
| CNK2/CALM3/PLCB1/GNAI2/CAMK2G |
| APK11/MAPK12/PPP3CC/FYN/CD3G/CD40LG/ITK/REL |
| M2 |
| RAC1/LYN/CDK6/YWHAB/TRADD/RASA2/CDK4/PIK |
| IK3R1/PPARGC1A/FOXO1/CPT1B/PRKCE/SLC27A6/GI |
| FB3/MAPK11/C1QC/IFNGR2/MAPK12/GNAQ/CD3G/PL |
| PP3CB/MAOA/ARC/PPP3CC/CALM3/FOSB/ATF4/CAM |

IF2AK3/NOX1/MAPK14/TP53/PLCB4/SRC/BID/HSPA8/(
./STK11/KL/ATF4/RELA/AKT1

B/COX6B1/DCTN5/KLC3/PSMD4/SPTBN2/PSMB3/CAP
AKT3/MAPK10/FOS/JUN/AGTR1/RASGRP2/GNB4/GST
L2/CXCL3/CXCL8/CXCL1/IL1B
PRCC/GADD45B/ARNT2/CCND2/TSPAN7/FLI1/POLK/S
.3/PIK3R1/MAPK11/RAP1A/RASSF5/ITGAL/MAPK12/C
GOH/PPP2R1B/MAGOHB/UPF1/SMG5/CPSF7/CSTF2/DI
2/JAK2/HLA-DMB/TGFB3/LAMA2/MAPK11/HLA-DPA1
.K2/PIK3R1/MAPK11/LEFTY2/ZFHX3/ID2/FZD4/MAPK
VHAB/ANAPC15/CCNB2/SMC1A/PKMYT1/CDK2/YWH
AC2/PLCG2/RELA/AKT1/LILRB5/DAPP1/VAV1/CD81/F
MAPK12/PPP3CC/PTEN/CD3G/RELA/AKT1/PDCD1/CD2

JAK2/PIK3R1/IFNGR2/IL12RB1/PIAS2/MPL/SOCS3/IL2
/PSMD2/PSMB6/PSMC1/PRKCG/ATP2A2/SP1/PSMA5/P
GT1A10/NR0B2/SLC22A1/SLC4A5/SCT/ABCB4/SULT2A
1/PTS/MTHFD2/ALAS1/LIPT2/GUSB/MTHFD1L/KMO/I

AP1A/NGF/MAPK12/SH2B1/MAPK7/NTF3/CALM3/BEX

/KCNK2/PLCB1/ATF4/CACNA1G/CYP21A2/STAR
RASSF4
CAMK2A/HLA-DMB/TGFB3/PPP3CB/MAPK11/HLA-DP.
H1B1/ACOX3

PIK3CD/BMPR1B/CAV2/PIK3R1/MAPK11/GPC1/MAPK1
FC/PDGFD/LPAR1/GUCY1B1/MAPK7/GNAQ/PLCB2/PL

PIK3CD/ADRB1/PRKACA/CYP1B1/JAK2/EPHX1/CREB
.CB2/CALM3/PLCB1/ATF4/CAMK2G/CALML3/GRIN2C
X2/CYP2U1/PLA2G5

ACNA1G
CYP2E1/HSD11B1L/SULT2A1/ADH5/ADH1C/GSTM3/GS
Y

GFB3/FOXO1/MAPK11/GABARAP/HOMER2/MAPK12/
/NDUFA4L2/PSMD14/APAF1/TUBB/SLC25A5/GNAI3/P
1B/PPP3CC/GNAQ/CPT1C/PLCB2/CALM3/PLCB1/ATF4/
3/PIK3CD/COL1A2/COL1A1/NDUFA5/NDUFS4/PPP1CB

B2/CDK4/PIK3R3/CCND1/MYC/PIK3R2/CDK2/CHEK1/
APK11/MAPK12/PPP3CC/CYLD/CALM3/PLCG2/EGR3
2E1/ADH5/ADH1C/GSTM3
1/MAPK11/PRKCE/NGF/MAPK12/GNAQ/PLCB2/CAL
L4L2/CGAS/POLR2K/POLR2L/IKBKG/POLR1D/POLR3

ANGPTL4/CIDEB/NPC1
LCB2/CALM3/PLCB1/GNAI2/CAMK2G/KITLG/WNT6/
YLT1/GALNT6/POC1B-GALNT4/GALNT2/GALNT5/PO

PRKACA/CREB5/PIK3R1/KRT24/GNAQ/PLCB2/CALM
HLA-DPB1
K3/CXCL2/CXCL6/CXCL5/MMP3/CXCL3/CXCL1/IL1B

GFR/CCL2/CTF1/CCL21/LEP/IL6R/CCL19/CX3CR1/AC
/CCL19/PIK3CD/CX3CR1/TIAM1/GNG2/CXCR4/CCR5/

PLCG2/AKT1/NRG2/FGF2/FGFR3
B/SLC25A5/ZBP1/TNF/RBCK1/H2AW/PYCARD/FAS/B
HGAP10/SEPTIN1/HCLS1/WASF1
2/MAPK12/PPP3CC/FYN/CYLD/SOCS3/FOSB/IL1R1/T
CL6/CXCL5/CXCL3/CXCL8/IL24/CXCL1
C/PRPF19/TRIM37/ANAPC13/ANAPC4/WWP1/SYVN1
FRB/AKT1/FGF9/CDKN1A/FGF2
P/RAB7B/GABARAPL1/PTEN/RRAGD/CTSB/UVRAG/
DH/ERBB2/ALDOC/TFRC/FLT1/HK2/ELOC/PIK3R2/E
DH5/ADH1C/RETSAT
1/CLDN5/PRKACA/ACTN1/CD1C/RAP1A/PRKCE/CG
A2/THRB/FOXO1/PLCB2/RCAN2/PLCB1/PLCG2/AKT1

LCB2/PLCB1/TSHR/ATF4
ATC1/NFATC4/PIK3CD/CRTC3/CD3E/CCND2/HLA-DQ

AC1/MAPK13/LYN/CDK6/VEGFA/TRADD/PLCG1/CY
C3A2/SAT1/STEAP3/TFRC/SLC11A2/ACSL4/SLC7A11

NFYB/HLA-DRB5/CTSB/HLA-DMA/HLA-DPB1/CTSL
NA/LIG1/RFC3

C5/LSM2/LSM8/HSPD1/ENO1/LSM7/LSM4/PABPC1L/P
/XYLT1
4/RELA/AMBRA1/RRAS/ULK1/TFE3/BCL2L13/FIS1/R

7A1/ASMT
/PIK3C3/CDS2/IPMK/PLCB1/PLCG2/MTMR3/CALML3/

PK12

2/HLA-DQB1/ANGPT1/HLA-DMB/TGFB3/HLA-DPA1/I
C27A1

S/RCHY1/IKBKE/IL1A/EIF2AK3/TP53/BID/HSPA8/CDK
IL6R/NFATC4/PIK3CD/GNG2/CXCR4/CCR5/PRKACA/C
RAC3/E2F1

DRA/PIK3CD/TPSB2/HLA-DQB1/JAK2/PIK3R1/HLA-D

/AP1S2/CD3E/GNG2/SAMHD1/CXCR4/CCR5/APOBEC3
1/PLCD4/PIP5KL1/SACM1L/PIP5K1A/ALDH6A1/MTM

ALM3/ITGB2/GNAI2/RELA/NLRP3

HRD/BMPR1A/TGFBR2/AMHR2/HAMP/SMAD4/BMPR
2/MYC/PIK3R2/SLC1A5/SIRT6/PKM/SLC2A1/SLC7A5
HAB/CYCS/PIK3R3/CASP3/IKBKG/MYC/IRAK1/PIK3R
M/GSTK1/ODC1
1G/AKT1/GAB2/VAV1
M3/AACS/HADH
I1/AHNAK/DYNC2H1/DYNC1H1/ARHGEF26/FYCO1/P
C2LI1/DCTN2

6V0B/ATP5PD/COX8A/ATP5F1A/COX7B/COX17/ATP6

11/C1QC/CPT1B/MAPK12/SCD5/ADH4/CPT1C/TNFRS

1C/PIK3R1/TGFB3/LAMA2/RAB7B/COL3A1/VCL/GNA
1/TJP2/PDIA4/CFTR/ERO1A/KDELR2
B/CD48/SH2D1A/ITGB2/PLCG2/FCER1G/CD247/VAV1/

K3R2/MECOM/E2F3/E2F2/E2F1
ULT2A1/GSTM3/GSTO2/GSTA2/UGT1A1/HPGDS/UGT2
11/ITGAL/MAPK12/H2AC8/SELP/RAC2/HDAC7/HDAC
12/RAC2/PTEN/FYN/GNAQ/PLCB2/PLCB1/TNFRSF1A/
L/HADH/ACSL4/ELOVL6/OXSM/HSD17B12

COX7A1/SLC25A4/HSPA1A/CYBB/PIK3CD/C1QA/TUB
1/LPAR1/KIT/DGKB/FYN/AGPAT4/CYTH3/PLCB2/PLC
7/POLK/PIK3R1/FLT4/FZD4/KIT/PTEN/AKT1/HEYL/F
B1/CPE
DGKB/RAC2/PCYT1A/PDGFRB/WASF1/SLC22A1/AKT1
1/BID/PDPK1/SLC31A1/CYCS/MGST2/PIK3R3/CASP3/
D1/DDC/RGS9/GNAI1
CND2/FZD3/RECK/TIMP3/PIM1/CYP1B1/PIK3R1/RPS

/WNT9A/COL6A3/COL1A2/COL6A2/ITGA5/CCND2/R
52/PIK3R1/FZD4/ATP6V1B2/PTEN/RRAGD/TNFRSF1A
WNT2/FRAT2/PIK3R3/FZD10/CCND1/ERBB2/WNT5A/
KG/CXCL2/CXCL3/CXCL8/CXCL1
WHAE/TANK/CARD6/TNF/RBCK1/BCL2L1/PKN2/PYC
MAPK13/MAD2L1/ANAPC15/CCNB2/PIK3R3/PIK3R2/P
FZD1/DVL2/WNT10A/PTCH1/WNT10B/FZD8

A7/AP2A2

K3R1/MAPK11/UQCRB/MAPK12/SOCS3/DDIT3/NDUF
1/WIPF3/MAPK12/WIPF1/RAC2/GNAQ/FYB1/ITGB1/T
1CB/CREB5/GNB5/NPY/MAOA/GNG11/H2AC8/HDAC7
/FZD7/CAMK2A/CREB5/RAP1A/FZD4/GNAQ/PLCB2/

CTN1/PIK3R1/FOXO1/RPS6KA5/MAPK11/PRKCE/TLN1
L13A/RPS8/RPS3A/RPL13/RPLP2/RPS23/RPS25/RPS24/I
GUCY1B1/GMPR/AK6/HDDC2/GUCY1A2/NUDT16/PDI
APK14/MAPK13/TRADD/IKBKG/TRIM25/TRAF2/CXCI
5/ZNF630/SRSF3/CGAS/ZNF74/ZNF764/ZNF689/EIF2AI
/CTSB/DDIT3/NTRK1/TNFRSF1A/ATF4/CTSL/RELA/AK
AD3/FZD7/CTNNA3/PPP1CB/TGFB3/ID2/FZD4/DLG2/F

ISD17B1/UGT1A1/UGT2A3
STP1/WNT3/LRP5/TP53/SMARCA4/TXNRD3/RB1/CDK

T1/CAMK2G/CDKN1A
TF2A2/GTF2A1/TAF4/GTF2H1/TAF5L/GTF2H3/CDK7/G

-1/PTEN/PDGFRB/ATF4/RELA/AKT1/CDKN1A

/AKT1

KN1B/NRG2

R2
1C2/NDUFA5/CYP1B1/NDUFS4/EPHX1/PIK3R1/MAPK

T/BRAF
I2/SMYD2/HADH/CAMKMT/MECOM/PLOD3

LA7/PLB1/PLA2G2D/CDS2/PTDSS2/PLPP1/PLA2G5

ASF1/AKT1/PLPP1/GAB2/ASAP3/VAV1/PTPRC/FCGR2B
2F2/E2F1

OPH1/MDH2/PSAT1/SRM

RKACA/NDUFS4/CREB5/PPARGC1A/MAPK11/CPT1B/U

32/PARD6A/IGF2R/RAB22A/AP2S1/WASHC5/VPS25/LD
AP/GALC/CERS2/DEGS2/CERS4/CERS5/B4GALT6/ENPP

C

HMGCL/SLC25A17/PEX7/ACSL3/SLC27A2/PECR/HSD1

N1/ASAH1/LGMN/GNS/ATP6AP1/PSAP/SUMF1/CTSW/
R1C4/HSD17B4

| morphologic feature | group |
|---|---|
| nuclear pleomorphism | nuclear & cellular morphology |
| nuclear size | |
| light nuclei | |
| nucleoli | |
| N/C ratio | |
| vacuolization | |
| tumor necrosis | tumor architecture |
| tumor cell count | |
| tubule formation | |
| crowded glands | |
| cribriform pattern | |
| solid pattern | |
| mucin amount | TME characteristics |
| ECM amount | |
| desmoplasia | |
| fibroblast infiltration | |
| immune infiltration | |
