## Supplemental Tables Description for "Conventional therapy induces tumor immunoediting and modulates the immune contexture in colorectal cancer"

### Supplementary Tables

**Table S1 Clonal composition and prevalences**

The table outlines the prevalence (column “clonal_prev”) of the clonal populations (column “clone”) found in the sequenced samples, as determined by the tumor evolutionary trajectory inference. The term clonal prevalence refers to the proportion of tumor cells harboring a specific mutation.

**Table S2 Clonal mutations**

The table describes the mutations (column “variation”) unique to each clonal population (column “clone_ID”). For each mutation, the chromosome (column “chrom”), genomic coordinates (column “coord”), and variant allele frequency (column “VAF”) are provided, respectively. VAF represents the fraction of variant sequencing reads within a genetic locus.

**Table S3 Differentially expressed genes**

The table demonstrates the results of the differential gene expression analysis (p.adjusted < 0.1) that was performed with DESeq2.

**Table S4 KEGG Gene Set Enrichment Analysis (GSEA) results**

The table presents the results of the KEGG GSEA analysis. Each enriched term (column "description") is listed alongside its corresponding p-value (column "pvalue"), adjusted p-value (column "p.adjust"), and the genes (column "genes") contributing to the enrichment of the KEGG term.

**Table S3 Histopathology - Morphologic features**

The table lists the features evaluated on selected tiles for semi-quantitative scoring.
